# Small molecule photocatalysis enables drug target identification via energy transfer

**DOI:** 10.1101/2021.08.02.454797

**Authors:** Aaron D. Trowbridge, Ciaran P. Seath, Frances P. Rodriguez-Rivera, Beryl X. Li, Barbara E. Dul, Adam G. Schwaid, Jacob B. Geri, James V. Oakley, Olugbeminiyi O. Fadeyi, Rob C. Oslund, Keun Ah Ryu, Cory White, Tamara Reyes-Robles, Paul Tawa, Dann L. Parker, David W. C. MacMillan

**Affiliations:** Merck Center for Catalysis at Princeton University, Princeton, NJ 08544, USA; Discovery Chemistry, Merck & Co., Inc, Kenilworth NJ 07033, USA; Department of Chemistry, Princeton University, Princeton, NJ 08544, USA; Discovery Chemistry, Merck & Co., Inc., Boston, MA 02115, USA; Merck Exploratory Science Center, Merck & Co., Inc., Cambridge, MA 02141, USA; Pharmacology, Merck & Co., Inc., Kenilworth, NJ 07033, USA

## Abstract

The identification of cellular targets that can be exploited for therapeutic benefit, broadly known as target ID, remains a fundamental goal in drug discovery. In recent years, the application of new chemical and biological technologies that accelerate target ID has become commonplace within drug discovery programs, as a complete understanding of how molecules react in a cellular environment can lead to increased binding selectivity, improved safety profiles, and clinical efficacy. Established approaches using photoaffinity labelling (PAL) are often costly and time-consuming due to poor signal-to-noise coupled with extensive probe optimization. Such challenges are exacerbated when dealing with low abundance membrane proteins or multiple protein target engagement, typically rendering target ID unfeasible. Herein, we describe a general platform for photocatalytic small molecule target ID, which hinges upon the generation of high-energy carbene intermediates via visible light-mediated Dexter energy transfer. By decoupling the reactive warhead from the drug, catalytic signal amplification results in multiple labelling events per drug, leading to unprecedented levels of target enrichment. Through the development of cell permeable photo-catalyst conjugates, this method has enabled the quantitative target and off target identification of several drugs including (+)-JQ1, paclitaxel, and dasatinib. Moreover, this methodology has led to the target ID of two GPCRs – ADORA2A and GPR40 – a class of drug target seldom successfully uncovered in small molecule PAL campaigns.

## Main text

The identification of biological targets and understanding of their interactions at the molecular level (target ID) is essential for the successful design of new therapeutic candidates and their progression into the clinic^1,2^. In recent years however, the intrinsic challenges associated with fully characterizing drug targets has manifested in low success rates and lengthy timelines, resulting in an industry-wide bottleneck within the developmental pipeline^3,4^. Therefore, the development of new methods to elucidate small molecule targets has the potential to significantly increase the success of therapeutic target selections, which should in turn lead to a reduction in clinical attrition and ultimately patient morbidity (Scheme 1a)^1,5,6^.

**Scheme 1.**
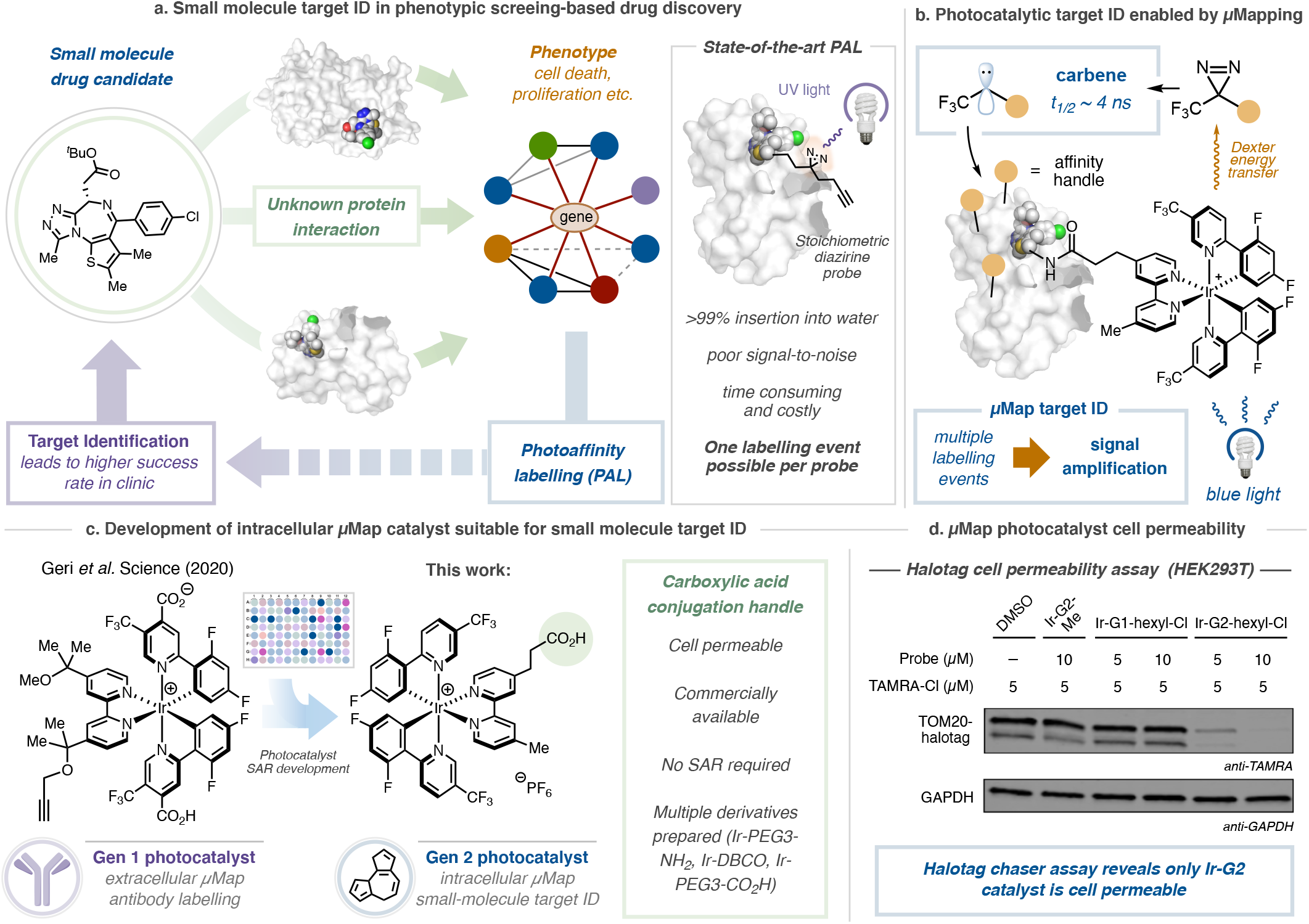
Photoaffinity labelling comprises a critical component of small molecule target ID. **a**, Target ID campaigns are critical for the development of successful drugs, though often rely on challenging photoaffinity labelling campaigns that employ the stoichiometric activation of diazirine small-molecule conjugates with UV light. **b**, Our approach separates the warhead from the small molecule probe, instead employing the photocatalytic activation of diazirines using visible light, giving rise to significant signal enhancement. **c**, Development of cell-penetrating, generation 2 photocatalyst suitable for small-molecule conjugation and target ID. **d**, Cell permeability of Ir-photocatalysts determined by halotag chaser assay; photocatalyst PEG-hexyl chloride conjugate and TAMRA hexyl chloride incubated with HEK293T cells expressing TOM20-halotag. Western blot analysis and immunostaining TOM20 with anti-TAMRA reveals off-compete only in the presence of Ir-G2 catalyst.

Over the last two decades, technological advancements in the fields of mass spectrometry^7^, chemical genetics^8^, and bioinformatics^9^ have transformed drug target identification leading to improvements in our understanding of biological pathways and cellular signalling^2,10^. However, while this information has provided a more focused route to the complex process of drug discovery, there remains a demand for target identification technologies for proteins without a well-described mechanism-of-action^11^. To address this need, affinity-based approaches^12^, and photoaffinity labelling (PAL) in particular, have now become routinely used tools in drug discovery (Scheme 1a)^13^. PAL works by the incorporation of a stoichiometric photoactivatable group, such as a diazirine, and an affinity handle, such as biotin, into the small-molecule architecture^14^. Following UV-activation and affinity-based enrichment, immunoblotting and proteomic analysis can be used to gather information regarding the identity of the target protein^15^.

While these methods have been extremely empowering for a number of protein classes^16–18^, practically they remain capricious and typically struggle to determine the complete interactome due to low receptor and protein abundance and short half-lives, leading to low cross-linking yields and high background^15,19^. The use of diazirine-based probes in particular has been challenging in this context as >99% of the carbenes generated upon UV irradiation react with water and not the target^20^. These spent probes serve to further block the binding of unreacted molecules, further hampering labelling efficiency. As a result, costly and time-consuming structural optimization campaigns are often required to overcome these shortfalls.

Indeed, the inherent difficulties associated with PAL have inspired the development of several elegant methods that hinge upon the use of stoichiometric activated electrophiles^12,21–24^, single-electron transfer events^25^, or specific oxidizable residues^26^ to identify target proteins. However, many of these technologies are limited to a single labelling event per drug molecule, often require extensive structural optimization (linker length and composition), and require low-yielding downstream ‘click’ processing^19^. Moreover, intracellular labelling technologies are often hampered by low cell-permeability leading to high background signal^5^. We therefore reasoned that the development of a catalytic target ID technology that separated the drug molecule from the reactive warhead could overcome these challenges through multiple labelling events leading to signal amplification (Scheme 1b).

We recently disclosed a novel antibody-based proximity labelling platform for cell surface microenvironment elucidation, termed µMap^27^. This method relies upon the activation of diazirine molecules in close proximity to a set of photocatalysts appended to an antibody via Dexter energy transfer. Inspired by this unique activation mode, we questioned whether such a tactic could be leveraged for small molecule target ID through the incorporation of a photocatalyst onto a bioactive small molecule. However, at the outset of the investigation, we were cognizant of several challenges inherent in developing such a technology, such as catalyst cell permeability and biocompatibility, ease of chemical manipulation, retention of biological activity, and labelling efficiency (given each antibody contained an average of 6–8 photocatalysts). However, we reasoned that by ‘switching on’ catalysis through visible light activation, labelling could be controlled both spatially and temporally, bypassing intrinsic reactivity problems and enabling the identification of novel targets across numerous drug discovery programs.

We began by investigating cell permeability: employing a halotag-based chaser assay off-competing a TAMRA dye in HEK293T cells, we identified that our previous catalyst design (Gen 1) was impermeable by virtue of its neutral net charge and two carboxylic acid residues (Scheme 1d). Through screening different photocatalyst structures, we realized that Ir-catalysts containing both the dFCF3-phenyl pyridine moiety and 4,4-dialkyl bpy ligand were crucial in achieving the necessary triplet energy^27^. Pleasingly, by removing the carboxylic acid groups, the cationic photocatalyst (Gen 2) was rendered cell permeable (Scheme 1d). With this in mind, we evaluated conjugation handles based around the 4,4-dMebpy ligand, opting for a distal carboxylic acid to enable facile amide coupling. Importantly, our G2-iridium catalyst can be accessed on gram-scale and be readily conjugated to a range of linkers and complex small molecules (*vide infra*).

Confident in our ability to access almost any Ir-drug conjugate, we initiated our target ID campaign with the validated epigenetic tool compound (+)-JQ1^28^. A potent inhibitor of the BET family of bromodomain proteins (BRD2/3/4), several JQ1 structural analogues are in clinical trials for a variety of cancers including NUT midline carcinoma^29^. We prepared the corresponding (+)-JQ1-G2 conjugate (**1**) (Scheme 2) and validated target engagement *in vitro* with recombinant BRD4 in a competition assay vs. bovine carbonic anhydrase (CA). An equimolar amount of CA and BRD4 was treated with (+)-JQ1-G2 probe (**1**) and an excess of diazirine-PEG3-biotin prior to irradiation at 450 nm. Labelling intensity was measured by western blotting with a streptavidin stain. Pleasingly, these preliminary experiments revealed a 20-fold increase in labelling for BRD4 vs. CA compared to the unconjugated (free) photocatalyst (Scheme 2a). Importantly, the (–)-JQ1-G2 conjugate, which is known to not bind BRD4^28^, showed significantly reduced labelling, demonstrating that labelling is as a result of a ligand/protein binding event (Scheme 2c, A). In addition, we were able to confirm this through microscale thermophoresis (MST), where the addition of the Ir-catalyst made only a minor impact on the binding constant (Figure S1).

**Scheme 2.**
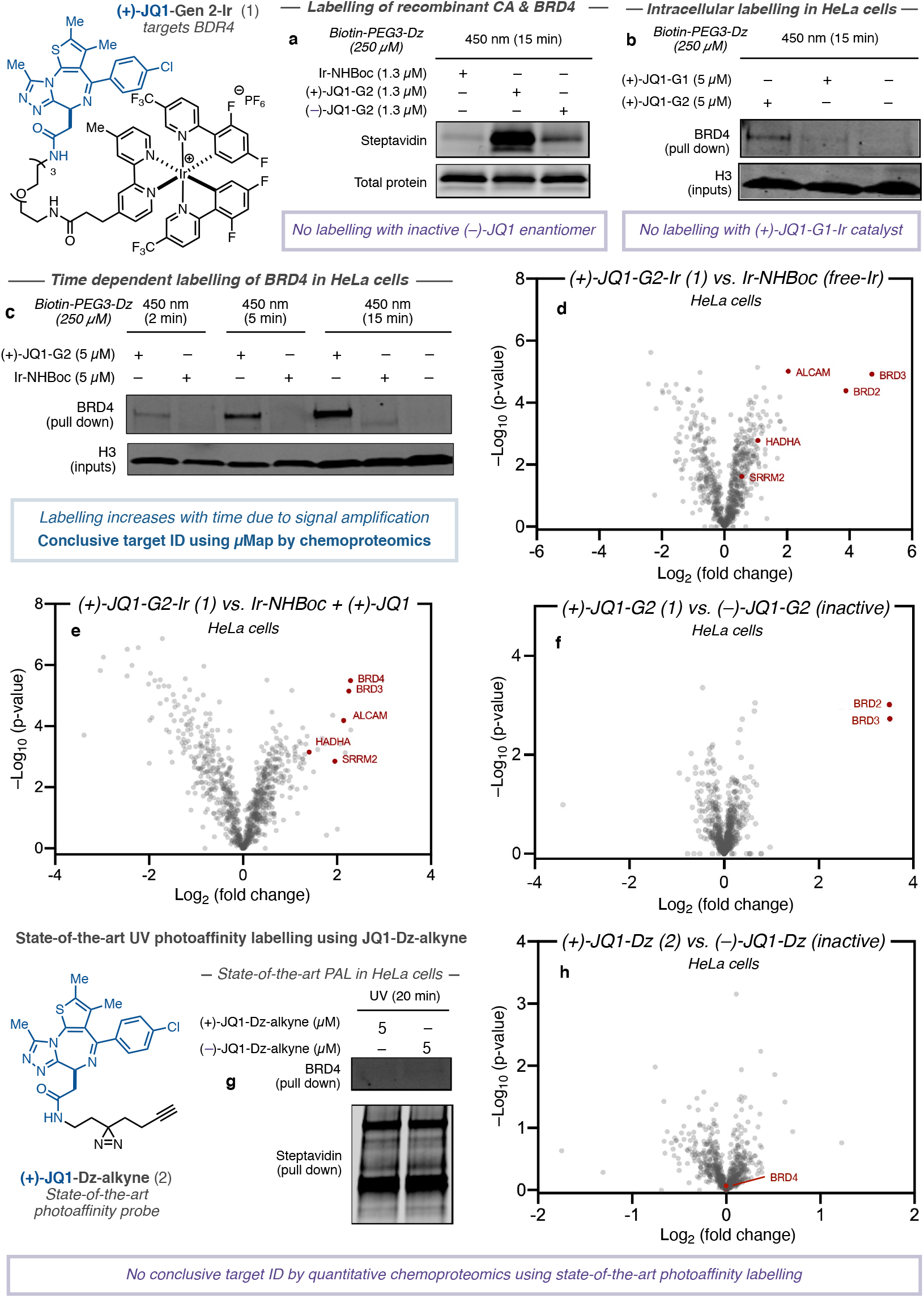
Development of photo-catalytic target ID platform for in-teractome mapping of (+)-JQ1 in HeLa cells. Structure of JQ1-based photocatalyst conjugate and state-of-the-art PAL probe. **a**, Labelling of recombinant BRD4 protein vs. spectator protein carbonic anhydrase using free iridium-, (+)-JQ1-, and (–)-JQ1-probe by immunostaining with streptavidin. **b**, Comparing permeability of G1 and G2-based (+)-JQ1 probes following irradiation in HeLa cells, streptavidin bead enrichment, and immunostaining with anti-BRD4. **c**, BRD4 labelling increases over time (2 min, 5 min, and 15 min irradiation) through photocatalytic signal amplification using (+)-JQ1-G2 probe in HeLa cells following streptavidin bead enrichment and immunostaining with anti-BRD4. **d**, TMT-based quantitative chemoproteomic analysis of JQ1-labelling in HeLa cells comparing intracellular labelling by (+)-JQ1-G2 catalysts and unconjugated iridium catalyst (control) reveals BRD proteins as highly enriched in addition to known JQ1 off-targets. **e**, TMT-based quantitative chemoproteomic analysis of JQ1-labelling in HeLa cells comparing intracellular labelling by (+)-JQ1-G2 catalysts and unconjugated iridium catalyst + (+)-JQ1 (control). **f**, TMT-based quantitative chemo-proteomic analysis of JQ1-labelling in HeLa cells comparing intracellular labelling between active (+)-JQ1-G2 and inactive (–)-JQ1-G2 cata-lysts reveals only (+)-isomer labels BRD proteins. **g**, State-of-the-art PAL employing active (+)-JQ1- and inactive (–)-JQ1-Dz-alkyne probes reveals no selective enrichment of BRD4 by western blotting despite broad biotinylation visible by immunostaining with streptavidin. **h**, TMT-based quantitative chemoproteomic analysis in HeLa cells comparing state-of-the-art PAL employing active (+)-JQ1- and inactive (–)-JQ1-Dz-alkyne probes reveals no enrichment of BRD proteins.

Based on these results, we sought to apply this method to live cells. We treated HeLa cells with 5 µM (+)-JQ1-Gen 2 (**1**) for 3h before the addition of 250 µM Dz-PEG3-Biotin and subsequent 15 min irradiation (450 nm). Following lysis and streptavidin-bead enrichment, western blot analysis with anti-BRD4 showed a clear labelling of the target protein compared to DMSO control (Figure S2). In line with previous findings, the corresponding (+)-JQ1-Gen 1 catalyst, while demonstrating similar *in vitro* labelling capability, showed no such enrichment of the target protein in cells (Scheme 2b). Consistent with our hypothesis, the intensity of labelling was found to be linearly related to irradiation time, demonstrating the photocatalytic signal amplification and temporal control offered by the µMap platform (Scheme 2c). This was also observable by confocal microscopy, wherein the degree of biotinylation imparted by increased significantly over time (Figure S3). Encouraged by our western blot validation data, we moved to TMT-based quantitative chemoproteomics in order to more completely assess the interactome of (+)-JQ1. To our delight, by comparing the labelling by (+)-JQ1-Gen 2 (**1**) vs. unconjugated (free) photocatalyst in HeLa cells, we observed several BRD proteins as the most enriched, although the precise identity of which however remains difficult to ascertain due to structural homology (Scheme 2d). We also identified two previously annotated (+)-JQ1 off-targets, HADHA^30^ and SRRM2^31^. ALCAM (CD166), a transmembrane glycoprotein, was also identified as being significantly enriched, but currently has no reported interaction with (+)-JQ1. CD166 exerts a pro-carcinogenic role via the inhibition of transcription factors along the FOXO/AKT axis and is considered a novel therapeutic target for liver cancer^32^. Interestingly, BET inhibition by (+)-JQ1 has been shown to upregulate expression of FOXO1, although the mechanism remains unclear^33^. In order to evaluate whether protein enrichment was as a direct result of labelling or up-regulation by virtue of the presence of (+)-JQ1, we repeated the experiment with an equivalent of (+)-JQ1 in the free iridium control (Scheme 2e). Upon chemoproteomic analysis we found that CD166 was similarly enriched, indicating that it may be a putative off-target binder of (+)-JQ1, although further biological validation is required. We further compared the interactomes of the enantiomers of JQ1-G2 and found the active (+) enantiomer, (**1**), delivered BRD2/3/4 as top hits, and while CD166 was detected it was not enriched, indicating that binding may not be affected by the stereogenic center (Scheme 2e). In contrast to these data, the same analysis using classical UV-based PAL employing (+)-JQ-Dz-alkyne (**2**)^34^, in our hands, did not lead to enrichment of BRD proteins by western blot (Scheme 2g) or chemoproteomic analysis (Scheme 2h).

The dual Src/Abl tryrosine kinase inhibitor dasatinib displays significant antileukemic effects against various imatinib-resistant mutants^35^. However, despite well-documented BCR/ABL inhibition, its precise downstream cellular MOA remains to be fully understood. While the dasatinib interactome has been previously characterized^16^, most methods have been performed with recombinant protein or in cell-lysate; live cell data is typically restricted to kinase-based assays that measure downstream phosphorylation or residence at engineered kinase constructs, which can be challenging to deconvolute and fail to identify non-kinase based off targets^36–39^.

As previous studies have demonstrated difficulties in maintaining potency and cell-permeability using dasatinib-derived probes^16^, we started by synthesizing three truncated (des<u>H</u>ydroxy<u>E</u>thyl<u>P</u>iperazinyl)-dasatinib iridium conjugates using our cell-permeable Ir-G2 catalyst with varying PEG linker lengths (n = 3–5) (**3**) (Scheme 3, top). Gratifyingly, upon subjection of the desHEP-dasatinib-G2 conjugates (**3**) (5 µM) to our standard µMap protocol, all of the conjugates revealed enrichment of p38 (MAP kinase) by western blot analysis compared to off-compete (4X dasatinib) controls in THP1 cells (Figure S4). As the corresponding PEG5-G2 conjugate showed the greatest enrichment (3.5X enrichment vs. off-compete and 9.5X enrichment vs. free-Ir) (Scheme 3a), we undertook label free proteomic analysis of these reactions revealing significant enrichment of p38α (Figure S5), which has been shown to play a critical role in its antileukemic properties^40^, as well as several other established kinase interactors including Src and Lyn (Scheme 3b)^41^. Furthermore, we identified multidrug resistance transporter ABCC1 amongst the most enriched proteins – an important off target; understanding the interaction between drug molecules and efflux transporters is an important consideration in many drug discovery efforts^42^. Lysosomal sequestration of dasatinib^43^, due to its lipophilic and weakly basic properties, was evident by the presence of cathepsin S (CTSS) amongst the most enriched proteins. Encouraged by these initial results, we turned our attention to the underexplored full dasatinib-PEG3-G2 catalyst (**4**), which retains the 2-hydroxyethylpiperazine tail. Importantly, we found a similar kinase inhibition profile against p38, in addition to Abl, by evaluation of downstream phosphorylation in Ph+ K562 cells, compared to the parent drug, again highlighting the compatibility of the iridium photocatalyst towards maintaining biological function and cell permeability (Scheme 3c) (Figure S6). Gratifyingly, subjection of our µMap labelling to TMT-based chemoproteomics revealed extensive enrichment of p38α as well as Myt1 and CSK kinases, both well-established binders of dasatinib (Scheme 3d)^39^. Moreover, known kinase off target ferrochelatase (FECH)^44^ was also significantly enriched, alongside large amino acid transporter (LAT3)^39^. Similarly, lysosomal protein cathepsin D (CTSD) was amongst the most enriched proteins. Notably, in our hands, state-of-the-art photoaffinity labelling, employing dasatinib-diazirine-alkyne (**5**), revealed only trace enrichment of CSK and the kinases BTK and MAPK1 (BTK was found to be similarly enriched by µMap) (Scheme 3e).

**Scheme 3.**
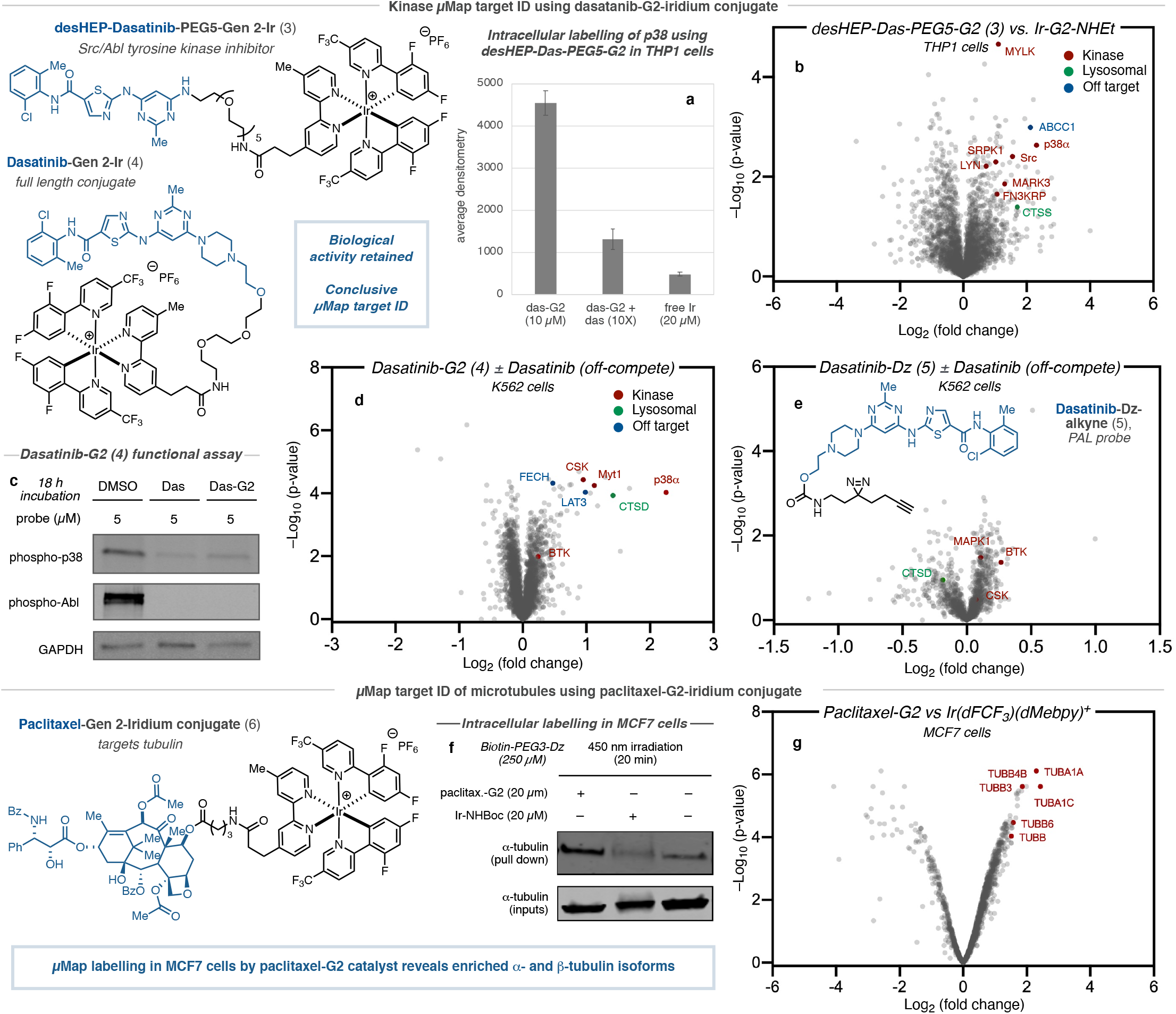
Intracellular photocatalytic target ID and interactome mapping of dasatinib and paclitaxel. **a**, Enrichment of p38 by western blot for labelling using desHEP-dasatinib-PEG5-G2 labelling in THP1 cells. **b**, Label free proteomic analysis in THP1 cells comparing intracellular labelling by desHEP-dasatinib-PEG5-G2 catalyst vs. Ir-G2-NHEt reveals enrichment of several kinases (red), as well as lysosomal proteins (green) and off-targets (blue). **c**, Kinase activity assays reveals dasatinib-G2 retains inhibition activity against Abl and p38, as well as general tyrosine phosphorylation, in K562 cells. **d**, TMT-based quantitative chemoproteomic analysis in K562 cells comparing intracellular labelling by dasatinib-G2 catalyst vs. dasatinib-G2 + dasatinib (off-compete control) reveals enrichment of several kinases (red), as well as lysosomal proteins (green) and established off-targets (blue). **e**, TMT-based quantitative chemoproteomic analysis in K562 cells comparing intracellular labelling by dasatinib-Dz-alkyne (PAL probe) vs. off-compete control does not reveal enrichment of kinases suitable for conclusive target ID. **f**, Initial western blot studies for paclitaxel-G2 labelling in MCF7 cells following irradiation and streptavidin bead enrichment reveals significant enrichment of a-tubulin by immunostaining compared to unconjugated iridium and DMSO controls. **g**, TMT-based quantitative chemoproteomic analysis in MCF7 cells comparing intracellular labelling by paclitaxel-G2 catalyst and unconjugated iridium catalyst (control) reveals enrichment of several tubulin isoforms.

The anti-cancer properties of the natural product paclitaxel (Taxol) have been proposed to be derived from binding to microtubules, leading to stabilization and mitotic arrest; however, the full extent of its mechanism remains unclear^45^. Based on its widespread use and intriguing mechanism, we prepared the corresponding paclitaxel-Gen 2-iridium conjugate (**6**) (Scheme 3, bottom) and assessed its cellular activity. Through a series of cell proliferation assays we found that our paclitaxel-G2 conjugate displayed similar anti-proliferative properties as the native compound, suggesting that the pendent Ir-catalyst did not disrupt the native function of paclitaxel (Figure S7). Encouraged by this, we proceeded to study the efficiency of labelling in the breast cancer cell line MCF7. Following our standard µMap protocol with 20 µM paclitaxel-G2 conjugate (**6**) for 3h, western blot analysis with anti-α-tubulin showed clear labelling of the target protein compared both the free iridium and DMSO controls (Scheme 3f). Subjection of our µMap labelling to TMT-based chemoproteomics revealed extensive labelling of tubulin isotypes αIa, βIII, βIVb, and αIc (Scheme 3g), which is in good agreement with previous photoaffinity labelling studies on extracted tubulin^46^.

Having established the efficacy of µMap target ID for intracellular proteins, we turned our attention to the cell surface. The exceedingly low abundance, lack of exposed residues, and aggregation-prone hydrophobic domains oftentimes confounds the detection and manipulation of membrane proteins, rendering target ID unfeasible^19,47,48^. These challenges are exacerbated when combined with the high background labelling, poor sensitivity, and low cross-linking yields systemic in PAL campaigns. We therefore felt that our µMap target ID platform was ideally placed to tackle these challenges by virtue of our catalytic signal amplification. We chose the adenosine receptor A2a (ADORA2A) as an exemplar membrane target. This GPCR has become an important target for immunotherapy^49^, but critically, has never been identified through live cell chemoproteomics^50,51^. Using a reported ligand for ADORA2A that binds from the extracellular face, SCH58261^52^, we prepared both a tethered diazirine-conjugate, SCH58261-Dz (**7**), and an Ir-conjugate based on the more hydrophilic G1 catalyst, SCH58261-G1 (**8**); the low cell permeability affording a higher effective concentration of photocatalyst probe on the cell surface (Scheme 4a). Photocatalytic labelling applied to A2a-expressing HEK293T cells, followed by western blot visualization, revealed a stark difference in labelling between the SCH58261-G1 (**8**) and the corresponding off-compete controls (Scheme 4b). Tandem mass tag (TMT)-based chemoproteomic analysis of these reactions confirmed our initial result, with our photocatalytic-labelling method using SCH58261-G1 (**8**) showing a 10-fold enrichment for ADORA2A with respect to off-competing with the parent SCH58261 ligand, and >20-fold enrichment versus free-Ir photocatalyst (Figure S8a). In contrast, PAL using SCH58261-Dz (**7**), showed poor enrichment of ADORA2A by quantitative chemoproteomics (Scheme 4c), in line with western blot data (Figure S8b). Based on the degree of enrichment in A2a-expressing HEK293T cells, we were keen to ascertain how the µMap target ID platform performed at native levels of membrane protein concentration, wherein classical PAL remains extremely challenging. Remarkably, photocatalytic labelling using SCH58261-G1 (**8**) in PC-12 cells, which have previously been validated to natively express A_2a_^53^, revealed similarly high levels of enrichment for the target protein ADORA2A – highlighting the signal amplification conferred by the µMap platform (Scheme 4d). Finally, we further validated this labelling technology by identifying the long chain fatty acid receptor GPR40, an important anti-diabetic therapeutic target, using the small molecule probe MK-8666, further expanding the repertoire of µMap membrane target ID (Figure S9–S12)^54^. In conclusion, we describe a general platform for photocatalytic target ID that utilizes cell-penetrating iridium conjugated-small molecules, which can bind protein targets, to locally activate proximal diazirines via Dexter energy transfer. The catalytic signal amplification conferred by µMap target ID has allowed for the identification of multiple protein targets and off targets across multiple drug classes and cellular compartments where established PAL have not been successful. As such, we envision that µMap target ID will find immediate use in providing a deeper biological understanding of efficacy target networks, quickly revealing off-target pharmacology, and ultimately driving pharmacotherapy forward against novel targets within drug discovery programmes in both academic and industrial settings.

**Scheme 4.**
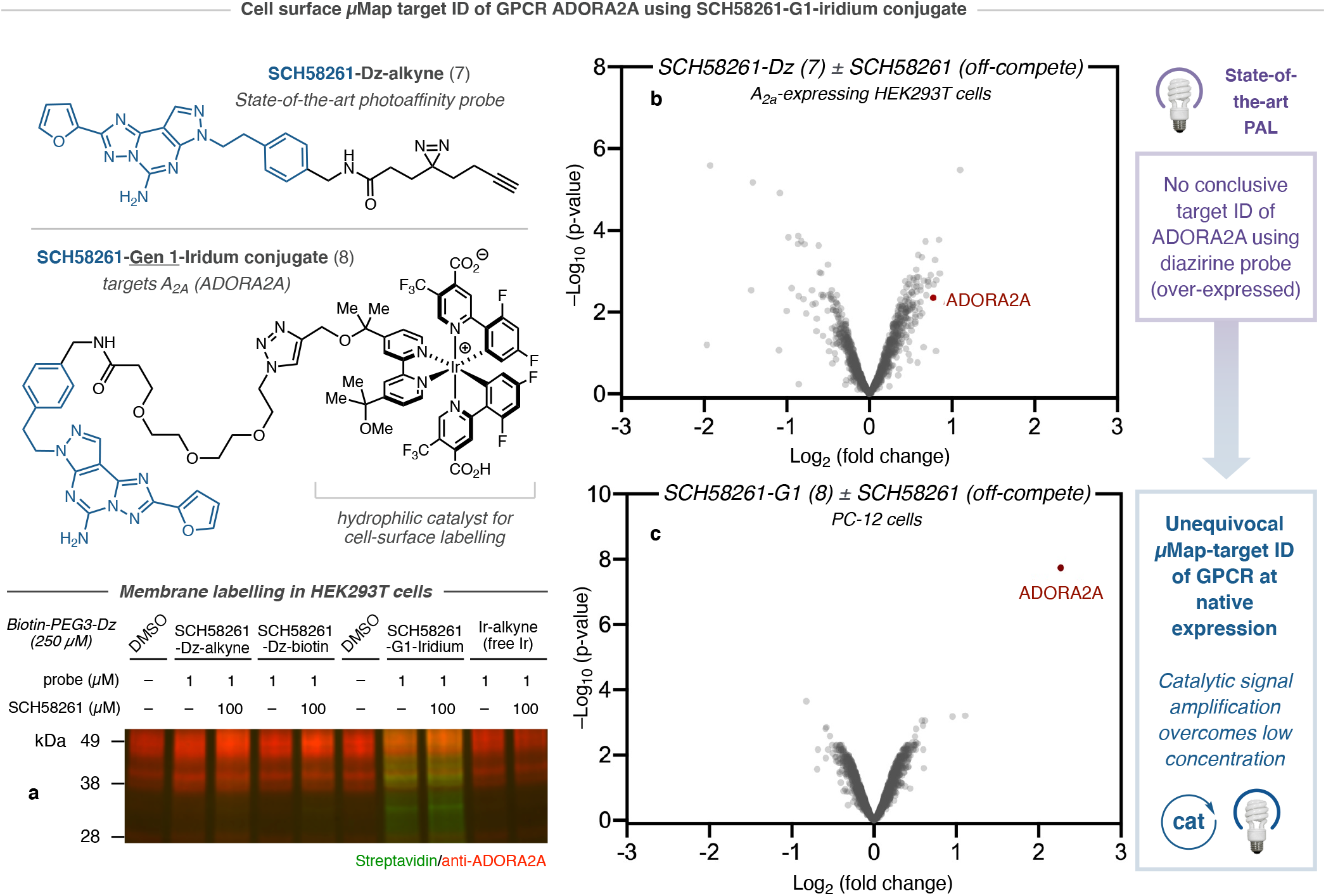
Extracellular photocatalytic target ID of GPCR ADORA2A using SCH58261 at native expression. **a**, Structure of SCH58261-based Dz-alkyne probe for PAL labelling and iridium conjugate based on hydrophilic G1 photocatalyst. **b**, Initial western blot studies for SCH58261-G1 labelling in A_2a_-expressing HEK293T cells following irradiation and streptavidin bead enrichment reveals significant biotinylation by immunostaining compared to unconjugated iridium and DMSO controls, in addition to PAL labelling. **c**, TMT-based quantitative chemoproteomic analysis in A_2a_-expressing HEK293T cells comparing extracellular labelling by SCH58261-Dz-alkyne vs. SCH58261-Dz-alkyne + SCH58261 (off-compete control) reveals inconclusive target ID of ADORA2A. **d**, TMT-based quantitative chemoproteomic analysis in PC-12 cells comparing extracellular labelling by SCH58261-G1 catalyst vs. SCH58261-G1 + SCH58261 (off-compete control) reveals significant enrichment of ADORA2A.

## Supporting information

Proteomics data

## Acknowledgments

Research reported in this publication was supported by the NIH National Institute of General Medical Sciences (R01-GM103558-03) and gifts from Merck & Co., Inc., Kenilworth, New Jersey, USA. ADT would like to thank the European Union’s Horizon 2020 research and innovation programme under Marie Sklodowska-Curie Grant Agreement No.891458. JBG acknowledges the NIH for a postdoctoral fellowship (F32-GM133133-01). JVO acknowledges the National Science Foundation Graduate Research Fellowship Programme under Grant No. (DGE-1656466). Any opinions, findings, and conclusions or recommendations expressed in this material are those of the authors and do not necessarily reflect the views of the National Science Foundation. The authors thank Saw Kyin and Henry H. Shwe at the Princeton Proteomics Facility. The authors thank Brande Thomas-Fowlkes and Xiaoping Zhang (MRL, Merck & Co., Inc, Kenilworth, NJ, USA), for providing GPR40-HEK cells and running IP-1 assay, respectively. HEK-hA2aR cell line was received as a gift from Jeremy Presland (MRL, Merck & Co., Inc, Boston, MA, USA). We acknowledge the use of Princeton’s Imaging and Analysis Center, which is partially supported by the Princeton Center for Complex Materials, a National Science Foundation/Materials Research Science and Engineering Centers programme (DMR-1420541). We also acknowledge V. G. Vendavasi and the use of Princeton’s Biophysics Core Facility. We thank Antony Burton for assistance in performing confocal microscopy. We thank T. W. Muir and members of the Muir Laboratory for their advice and analytical support.

## Author contributions

ADT, CPS, FPR-R, DLP, DWCM conceived the work. FPR-R, DLP, CPS, ADT, BXL, BD, JBG, JVO, AGS, OOF, RCO designed and executed the experiments. PT, TR-R, KAR provided insight into experimental design. ADT, CPS, FPR-R, DLP, DWCM prepared this manuscript.

