## Supporting Information for "Small molecule photocatalysis enables drug target identification via energy transfer"

Aaron D. Trowbridge[1]*, Ciaran P. Seath[1]*, Frances P. Rodriguez-Rivera[2*†], Beryl X. Li[1], Barbara E. Dul[3], Adam G. Schwaid[4], Jacob B. Geri[1], James V. Oakley[1], Olugbeminiyi O. Fadeyi[5], Rob C. Oslund[5], Keun Ah Ryu[5], Cory White[5], Tamara Reyes-Robles[5], Paul Tawa[6], Dann L. Parker, Jr.[2†], David W. C. MacMillan[1†]

[1]Merck Center for Catalysis at Princeton University, Princeton, NJ 08544, USA.
[2]Discovery Chemistry, Merck & Co., Inc, Kenilworth NJ 07033, USA
[3]Department of Chemistry, Princeton University, Princeton, NJ 08544, USA
[4]Discovery Chemistry, Merck & Co., Inc., Boston, MA 02115, USA.
[5]Merck Exploratory Science Center, Merck & Co., Inc., Cambridge, MA 02141, USA.
[6]Pharmacology, Merck & Co., Inc., Kenilworth, NJ 07033, USA.

*These authors contributed equally to this work.

#### Supporting Figures

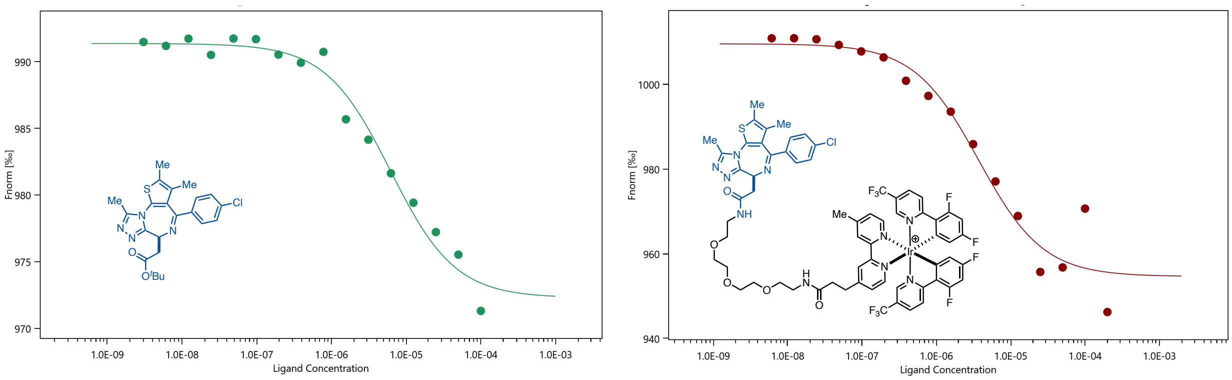

**Figure S1: Binding assay of (His)$_6$-tagged BRD4-AF633 conjugate vs. JQ1 and JQ1-PEG3-Ir via MST**

MST was subsequently performed on a Monolith NT.115 instrument with three replicates. (+)-JQ1 showed a binding $K_D$ = 6 μM and (+)-JQ1-PEG3-G2-Ir (1) showed a binding $K_D$ = 4 μM. The corresponding Ir-PEG3-NHBoc showed >85 μM binding. The lower binding of (+)-JQ1 compared to reported values is due to the use of a truncated form of the recombinant protein.

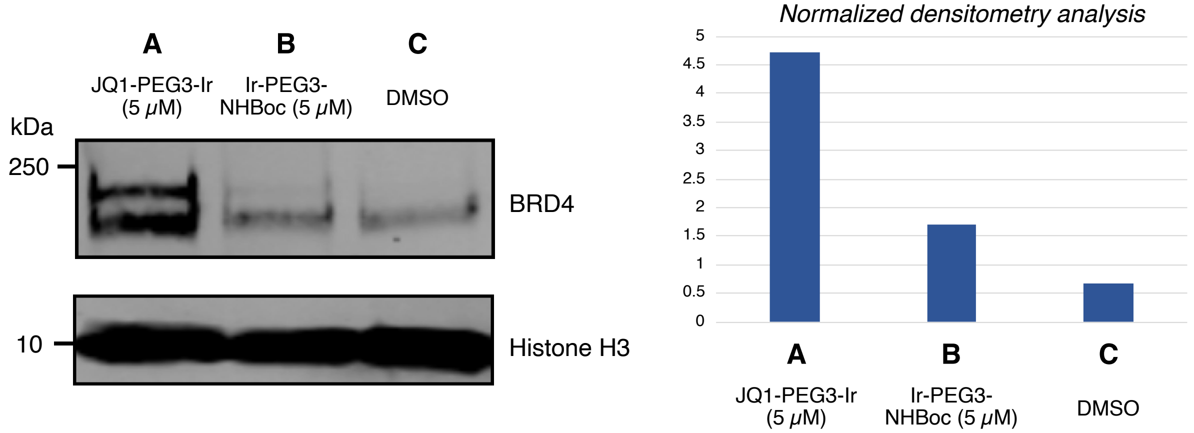

**Figure S2: Intracellular labeling in HeLa cells of BRD4 using JQ-1-PEG3-Ir conjugate compared to Ir-PEG3-NHBoc and DMSO controls.**

HeLa cells were incubated with 5 mM probe or DMSO control for 3 h before addition of Diazirine-PEG3-biotin (250 μM) and 15 min irradiation with 450 nm light. Cells were washed, lysed, and subjected to streptavidin enrichment before western blotting. BRD4 is clearly enriched in the directed experiment compared to the controls.

*Irradiation time = 2 mins*

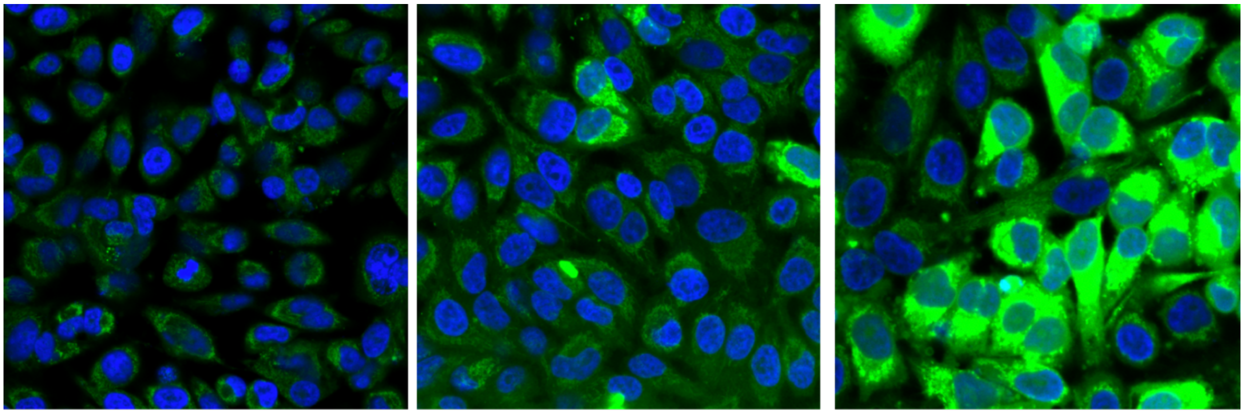

DMSO control

JQ1-PEG3-Ir
*(directed)*

Ir-PEG3-NHBoc
*(free-Ir)*

*Irradiation time = 10 mins*

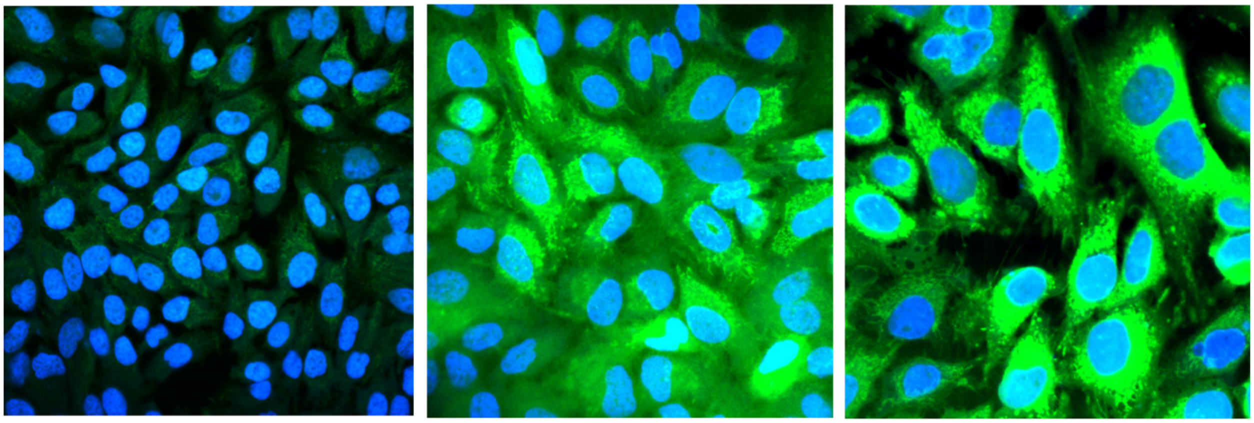

DMSO control

JQ1-PEG3-Ir
*(directed)*

Ir-PEG3-NHBoc
*(free-Ir)*

**Figure S3: Confocal microscopy of HeLa cells following probe incubation and irradiation.** Images show that labeling increases over time (streptavidin stain in green with Hoechst nuclear stain in blue). Labeling is spread evenly throughout the cell, displaying the cell permeability of the probe.

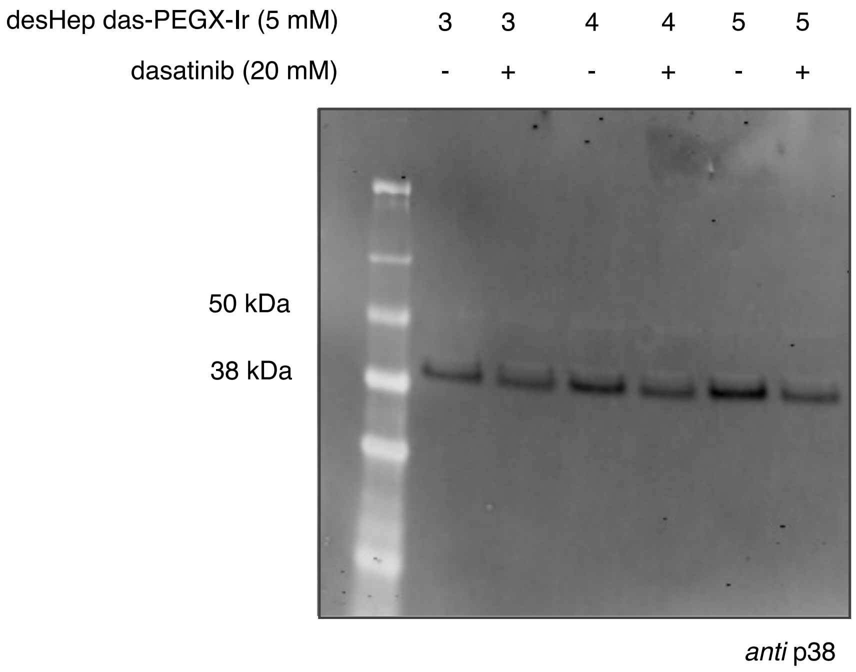

**Figure S4: Labeling experiment comparing desHep-PEG3 (3a), PEG4 (3b), and PEG5 (3c) analogs**

THP1 cells were treated with a solution of diazirine-PEG3-biotin (250 µM) in DPBS which contained either desHep-PEG3 dasatinib G2-iridium conjugate (**3a**) with or without 20 µM dasatinib, or 5 µM desHep-PEG4 dasatinib G2-iridium conjugate (**3b**) with or without 20 µM dasatinib or 5 µM desHep-PEG5 dasatinib G2-iridium conjugate (**3c**) with or without dasatinib (20 µM). Following incubation, irradiation with 450 nm light and centrifugation/washing and lysis steps described above, the samples were subjected to gel electrophoresis and WB also as described above. Selective labeling of MAPK14 (p38) with three varying linker desHep dasatinib irdium conjugates revealed by streptavidin pulldown followed by WB anti-MAPK14 (abcam ab31828) (1:1,000 dilution, 10 mL) for 1 hour then by incubation with LiCOR IRDYE-800CW goat-anti-mouse (926-32210) (1:15,000 dilution, 10 mL) for 1 hour.

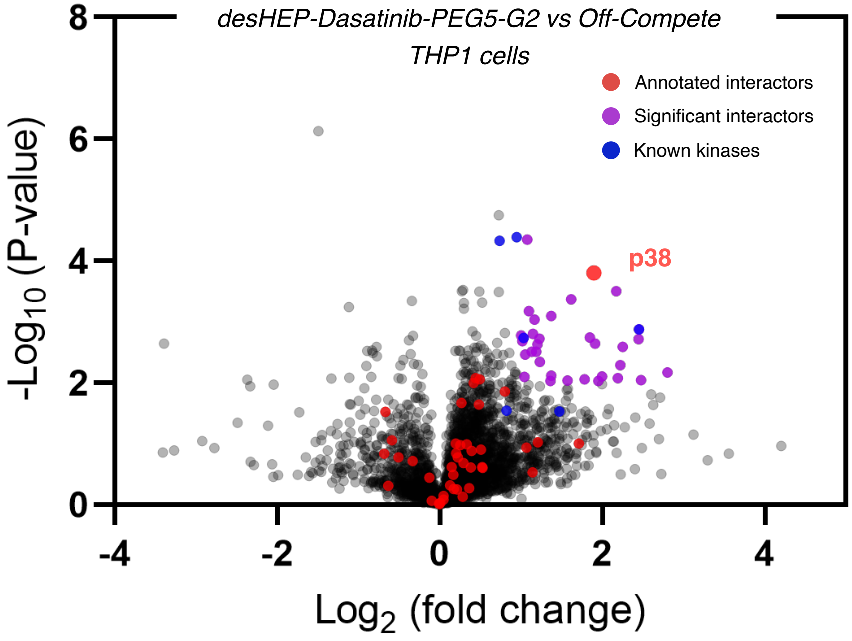

**Figure S5b: Label free proteomics experiment: desHEP-dasatinib-Ir vs off-compete.** desHEP-dasatinib-Ir (3) labelling identifies p38 as target by chemoproteomics when comparing ligand-targeted Ir versus ligand-targeted Ir with excess ligand in THP1 cells.

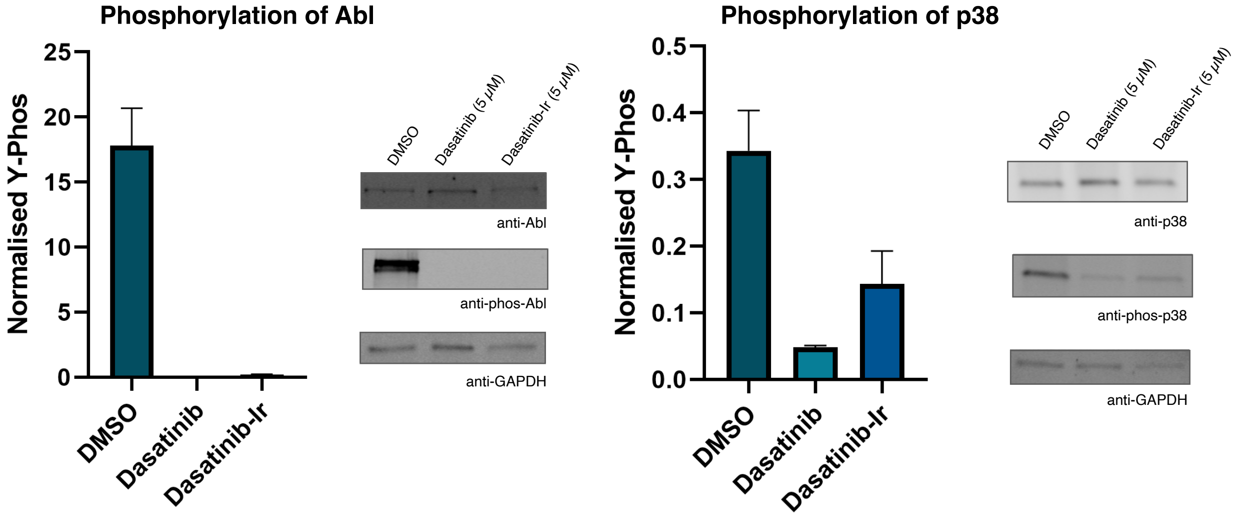

**Figure S6: Analysis of cellular activity in K562 cells with dasatinib vs Dasatinib-PEG3-Ir and DMSO controls.**

A 24 well plate was seeded with 500k K562 cells per well in 2 mL of IMDM. To each well was added either Dasatinib (5 µM), Dasatinib-Gen2-Ir (4) (5 µM), or DMSO. The cells were incubated for 24 hours, pelleted, and washed with PBS. Cell pellets were lysed (20 mM Tris, 1 mM β-glycerophosphate, 1 mM $Na_3VO_4$, 150 mM NaCl, 1 mM EDTA, 1 mM EGTA, 1% NP-40, 1% Na deoxycholate, 2.5 mM Na pyrophosphate, 0.1% SDS) and 10 µg of lysate was analysed by western blot, staining for p38 (Cell signalling: 9212), phos-p38 (Thr180/Tyr182) (Cell Signalling: 9211), abl (Cell Signalling: 2862S), phos-Abl (Cell Signalling: 2861S), and GAPDH (SCBT; 47724). Blotting shows that the addition of the Ir-catalyst has only a minor effect on dephosphorylation.

*Averaged absorbance values over 72 h (n=3)*

| A450 | 0h | 24h | 48h | 72h |
|---|---|---|---|---|
| Paclitaxel (2 μM) | 0.67 | 0.73 | 0.69 | 0.62 |
| Paclitaxel-G2 (6) (2 μM) | 0.76 | 0.77 | 0.76 | 0.54 |
| $Ir(dFCF_3)(dMebpy)^+$ (14) (2 μM) | 0.78 | 0.64 | 0.56 | 0.35 |
| DMSO | 0.68 | 0.97 | 1.60 | 1.44 |
| Control | 0.71 | 0.88 | 1.14 | 1.08 |

*Standard Deviation*

| SD | 0h | 24h | 48h | 72h |
|---|---|---|---|---|
| Paclitaxel (2 μM) | 0.07 | 0.14 | 0.10 | 0.10 |
| Paclitaxel-G2 (6) (2 μM) | 0.02 | 0.11 | 0.14 | 0.01 |
| $Ir(dFCF_3)(dMebpy)^+$ (14) (2 μM) | 0.01 | 0.10 | 0.05 | 0.01 |
| DMSO | 0.10 | 0.02 | 0.27 | 0.28 |
| Control | 0.04 | 0.08 | 0.03 | 0.04 |

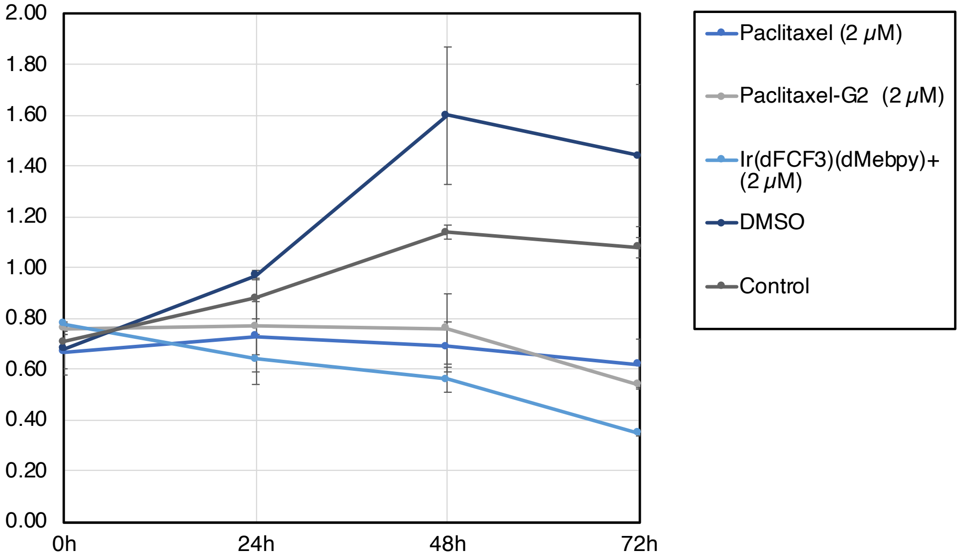

**Figure S7a: Cell viability assay with Taxol-Ir conjugate vs controls**

MCF7 Cells were grown to a confluency of about 80%, trypsinized, resuspended in fresh media and counted using a hemocytometer. Cells were diluted to 30,000 cells/ml and 100 μl was pipetted into a 96 well plate. Compounds were resuspended in DMSO at a concentration of 400 μm and diluted to 40 μm with water. Compounds were added to a final concentration of 2 μM, including controls for DMSO, to a final concentration of less than 1% DMSO. Cell viability was measured using EZ Quant Cell Quantifying Kit (ALSTEM) diluted 1:3 in PBS according to manufacturers. Briefly, at each time point, 20 µL of EZ Quant reagent was added (diluted 3-fold in PBS) and 2 h later the absorbance at 450 nm was recorded. Each viability experiment was performed in triplicate.

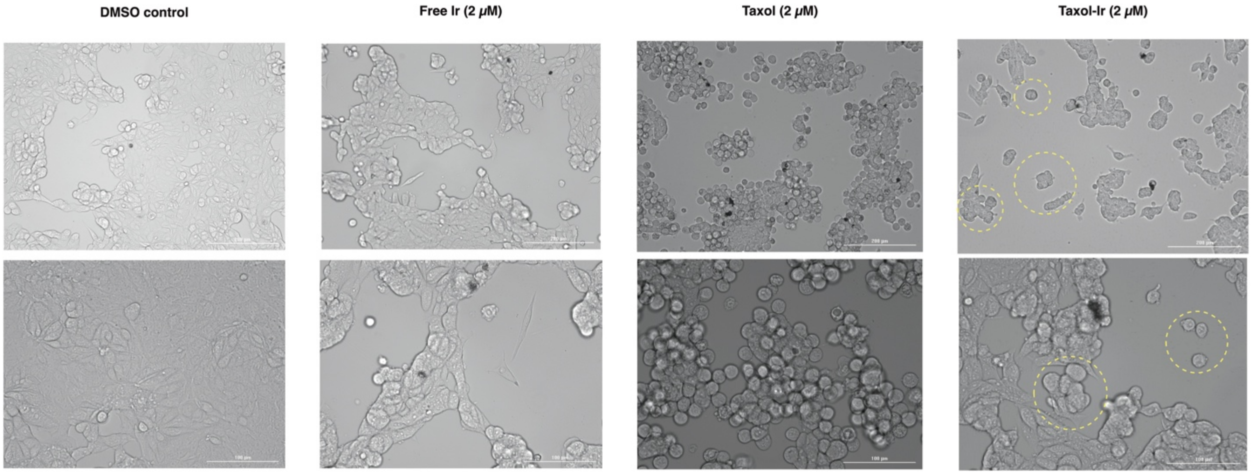

**Figure S7b: MCF7 morphology is consistent between Taxol and Taxol-Ir treatment.** Brightfield images taken after 16 h incubation with Ir(dFCF$_3$)(dMebpy)$^+$ (2 µM), Taxol-Ir (6, 2 µM), or taxol (2 µM). The MCF7 cell morphology changed when treated with both paclitaxel and paclitaxel-G2 (6). A pronounced shrinking or the membrane and rounding of the cells was observed in both cases, although viability was not impacted, the cells did not proliferate. While the free-Ir photocatalyst also resulted in loss of viability, cell death was more rapid and continuous, similar morphological changes were also not observed. Yellow circles show cells that clearly show the taxol treated phenotype in the taxol-Ir wells.

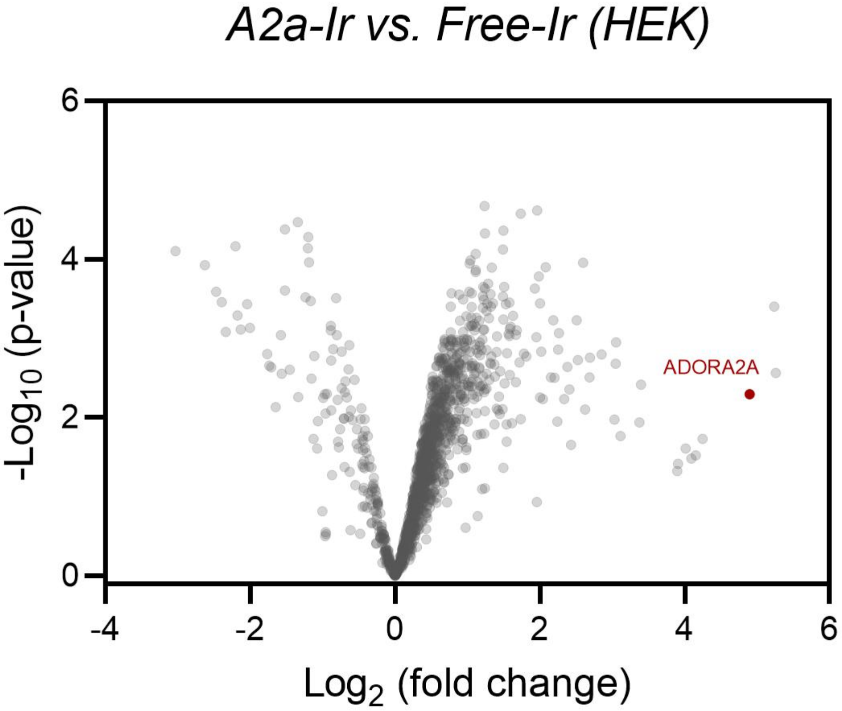

**Figure S8a: TMT-based chemoproteomic analysis of SCH58261-G1-Ir (8) labelling in A2a-expressing HEK293T cells vs. Free Iridium control (12)**

SCH58261-G1-Ir (8) $A_2$aR-Ir labelling identifies ADORA2A as target by chemoproteomics when comparing ligand-targeted Ir versus free Ir photocatalyst (**12**). HEK-hA2aR cells were treated with SCH58261-G1-Ir conjugate (8) or free photocatalyst for 30 min, washed and irradiated for 10 min at 450 nm after treatment with Diaz-PEG3-Biotin (**9**). Cell lysates were prepared and processed for chemoproteomic experiments in triplicate.

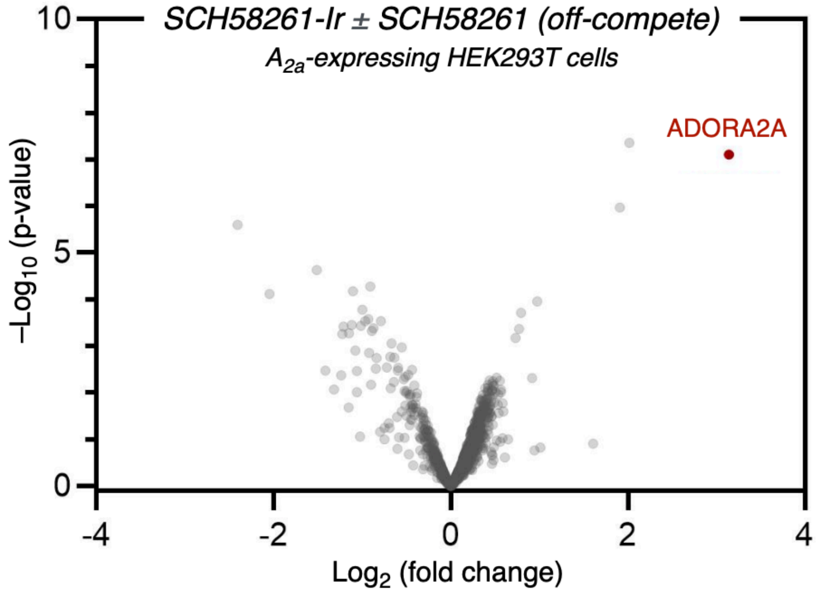

**Figure S8b: TMT-based chemoproteomic analysis of SCH58261-G1-Ir (8) labelling in A2a-expressing HEK293T cells**

The labeling reaction was performed according to the general procedures. TMT-based chemoproteomics identified ADORA2A as most enriched protein.

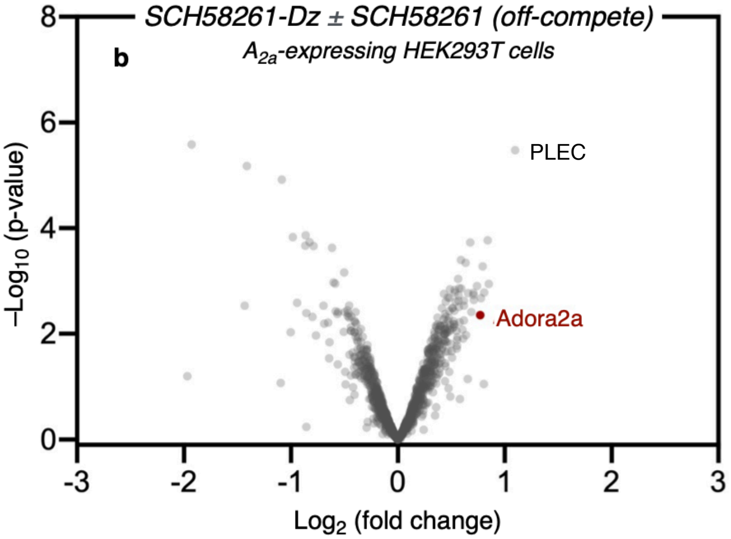

**Figure S9: PAL chemoproteomic target ID of Adora2a using SCH58261-Dz-alkyne (7) in A2a-expressing HEK293T cells**

The labeling reaction was performed according to the general procedures. TMT-based chemoproteomics identified PLEC as the target.

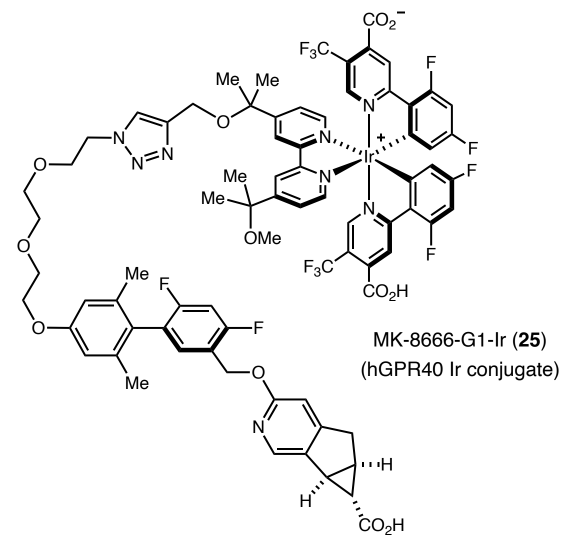

**Figure S10: Structure of MK8666-G1-Ir probe that targets GPR40**

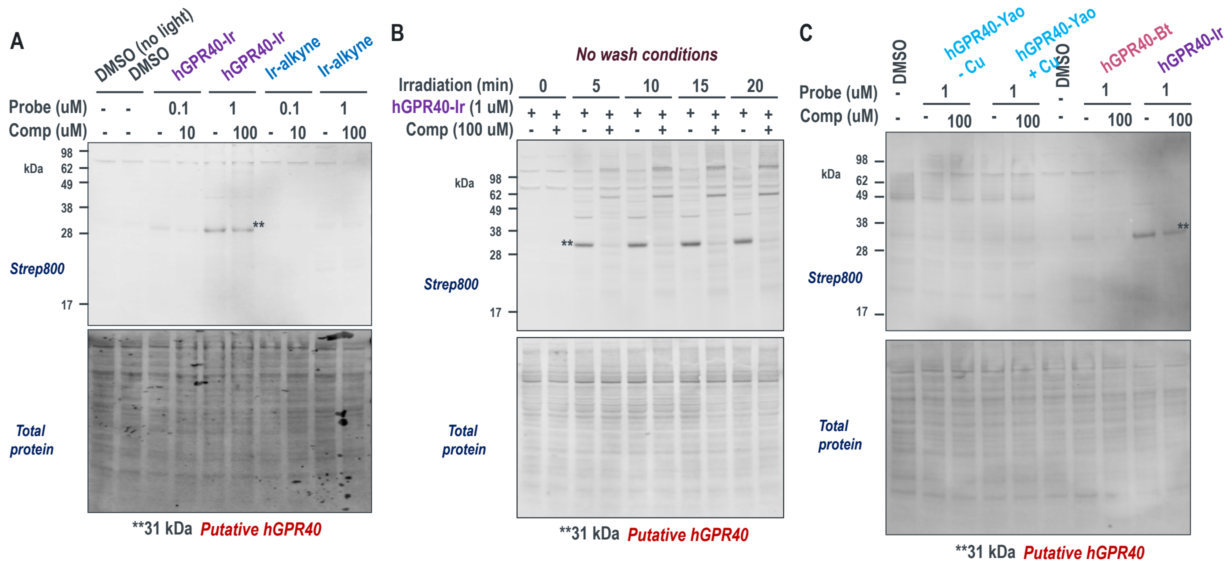

**Figure S11: WB analysis for MK8666-G1-Ir µMap labelling in HEK-hGPR40 cells**

Western blotting data for µMap experiments with MK8666-G1-Ir targeting hGPR40 in HEK293T cells stably expressing hGPR40. A) Streptavidin enrichment and staining shows enrichment of a single band that corresponds to the MW of hGPR40. The intensity of this band is diminished in the presence of off competing ligand. B) The intensity of the labeled band increases with increasing irradiation time and is ablated in the presence of excess MK8666. C) Putative hGPR40 is not observed when performing classic PAL methods.

*Streptavidin staining and enrichment of 31 kDa band** assigned as putative hGPR40*

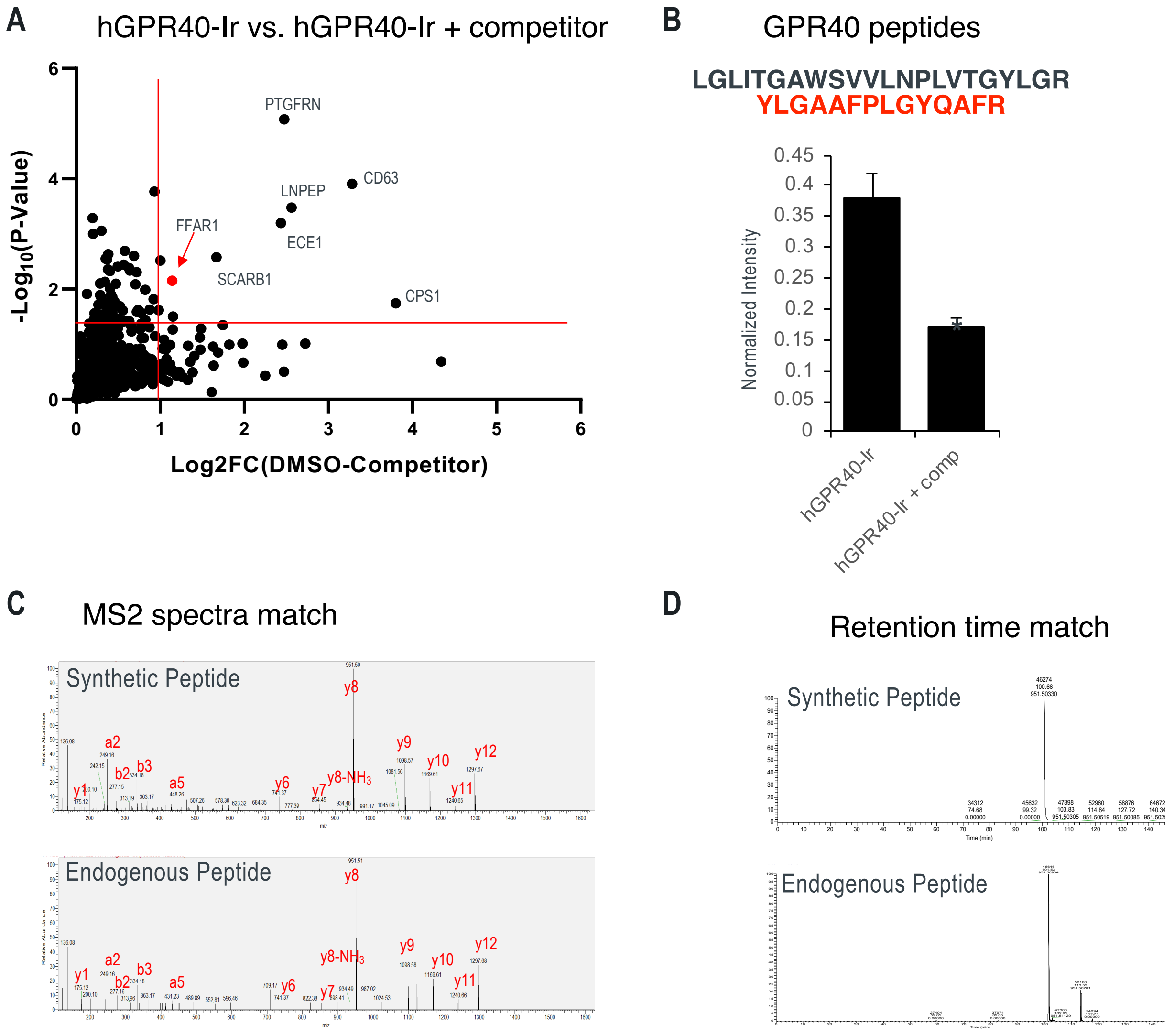

**Figure S12: MK-8666-G1-Ir labeling identifies FFAR1 (GPR40) by quantitative proteomics** (A) FFAR1 is enriched by chemoproteomics after MK8666-G1-Ir labeling. Labeling experiments were performed in triplicate and processed for proteomics. (B) hGPR40 peptides detected by LC-MS and normalized intensities for the YLGAAFPLGYQAFR peptide. (student's t-test, $p \leq 0.01$, error bars are SEM). Validation of synthetic hGPR40 peptide confirms correct identification by matching MS2 spectra (C) and retention time (D).

### General Considerations

#### Synthetic methods

Organic solvents were purified according to the method of Grubbs[1]. Water was purified using a Millipore Milli-Q Integral Water Purification System. Organic solutions were concentrated under reduced pressure on a Büchi rotary evaporator using a water bath. $^{1}$H NMR spectra were recorded on a Bruker UltraShield Plus Avance III 500 MHz unless otherwise noted and are internally referenced to residual solvent signals. Data for $^{1}$H NMR are reported as follows: chemical shift (δ ppm), multiplicity (s = singlet, d = doublet, t = triplet, q = quartet, p = quintet, m = multiplet, dd = doublet of doublets, dt = doublet of triplets…etc, br = broad), coupling constant (Hz) and integration. Irradiation of samples was performed in either a PennOC Photoreactor for small-molecule sensitization experiments (PennOC, Pennsburg, PA, Model M1) or a Biophotoreactor (Efficiency Aggregators, Richland, Tx, Fisher, NC1558343 BPR200). Spectral analysis of light sources was performed using a UPRtek MK350 handheld spectrometer. Chromatographic purification was carried out using a Biotage Isolera Prime flash chromatography system with SilaSep flash cartridges (60 mesh) and UV detection. $^{13}$C NMR spectra were recorded on a Bruker UltraShield Plus Avance III 500 MHz (125 MHz) and data are reported relative to the solvent employed. High resolution mass spectra and intact protein mass spectra were obtained from the Princeton University Mass Spectral Facility and Princeton Proteomics & Mass Spectrometry Core.

### General Materials

All buffers and synthetic starting materials were used as received from commercial sources. Ascorbic acid (BP321-500) and ethanol (BP2818100) were purchased from Fisher Scientific (Pittsburgh, PA). Bovine serum albumin (BSA) (A7906), Eppendorf Protein LoBind tubes (Z666505), Coppe(II) sulfate pentahydrate (7758-99-8), and (+)-sodium L-ascorbate (134-03-2) were purchased from Millipore Sigma (St. Louis, MO). Sodium azide (14314) was purchased from Alfa Aesar (Haverhill, MA). Mem-PER™ Plus membrane fractionation kit (89842), RIPA Buffer (89900), 1X DPBS (14190144), Pierce BCA Protein Assay Kit (23227), and iBright Prestained Protein ladder (LC5615) were purchased from Thermo Scientific (Rockford, IL). TBST (IBB-

581X) was purchased from Boston BioProducts (Ashland, MA). 5M Sodium chloride (S24600-500.0) was purchased from Research Products International (Mt. Prospect, S19 IL). 12% Criterion TGX precast gels (5671044) and 4x Laemmli sample buffer (161-0747) were purchased from Bio-Rad (Hercules, CA). 20% SDS solution (351-066-721) was purchased from Quality Biological (Gaithersburg, MD). Biotin-PEG3-diazirine, $[Ir(dCO_2HdFCF_3ppy)_2(MeCN)_2]OTf$, 2,2'-([2,2'-bipyridine]-4,4'-diyl)bis(propan-2-ol), and 2-(4'-(2-methoxypropan-2-yl)-[2,2'-bipyridin]-4-yl)propan-2-ol were synthesized as described previously[2]. $Ir(dFCF_3)(dMebppy)PF_6$ was prepared analogously to Ir-G2 catalyst using dMebppy (Aldrich). (+)-JQ1 was obtained from ApexBio (A1910). (+)-JQ1-$CO_2H$ was provided as a generous gift from Merck. (–)-JQ1 was obtained from Sigma Aldrich (SML1525). 2-(3-but-3-yn-1-yl)-3H-diazirin-3-yl)ethan-1-amine was obtained from Astatech (S10087). Biotin-PEG3-NHS was obtained from Broadpharm (BP21509). Boc-N-Amido-PEG3-amine (BP20583) and Amino-PEG3-t-Bu-ester (BP20697) were obtained from Broadpharm. DBCO-amine was obtained from Sigma Aldrich (761540). Paclitaxel was obtained from Astatech (N88686). 4-[3-(trifluoromethyl)-3H-diazirin-3-yl]benzylamine hydrochloride was obtained from TCI (T3448). Dasatinib was purchased from Astatech (62242). The A2a core (2-(furan-2-yl)-7H-pyrazolo[4,3-e][1,2,4]triazolo[1,5-c]pyrimidin-5-amine) was purchased from JW&Y Pharmlab, Co. Biotin-PEG3-azide (cat. 630721) and BTTAA (cat. 906328) were purchased from Sigma Aldrich. Yao linker building blocks were purchased from Enamine. SCH-58261 (cat. 2270) was purchased from Tocris Bioscience. EZQuant Cell Quantifying Kit was obtained from Alstem (Richmond, CA). Poly-L-Lysine Solution was obtained from Sigma-Aldrich (St. Louis, MO). Paraformaldehyde (20% solution) was obtained 5 from electron microscopy sciences (Hatfield, PA). 35 mm glass-bottom dishes were obtained from MatTek (Ashland, MA). Alexa Fluor 633 NHS ester was obtained from Thermo Fisher Scientific (A20005). Streptavidin-Alexa Fluor 488 was obtained from BioLegend (San Diego, CA). Standard Tissue Culture Dishes were obtained from Thermo Fisher Scientific (Waltham, MA). DPBS (Gibco, #14190250), DMEM high glucose (Gibco, #31053036), DMEM high glucose – no phenol red (Gibco, #31053028), RPMI 1640 (Gibco, #11875093), RPMI 1640 – no phenol red (Gibco, #11835030), Foetal Bovine Serum Gibco (#10437-028), L-glutamine (Gibco, #25030164), Penicillin-Streptomycin (Gibco, #15070063), Trypsin-EDTA (Gibco, #25300054), Trypsin protease MS (Pierce, #PI90057), and RIPA buffer (Thermo, #89900) were obtained from Thermo Fisher Scientific. PMSF (Sigma Aldrich, #78830) and cOmplete EDTA free protease inhibitor (Roche, #11873580001) were obtained from Sigma Aldrich. Streptavidin Magnetic Beads were obtained from New England Biolabs (NEB, #S1420S) or Thermo Fisher Scientific (Pierce, #88816) and stated in the text. EMEM (Sigma, cat. M6199-500) was obtained from Sigma Aldrich, Trifluoroacetic acid (Optima grade), Acetonitrile (Optima grade), Water (Optima grade), Acetic acid (Optima grade) were obtained from Thermo Fisher Scientific. Triethylammonium bicarbonate (1M Sigma Aldrich, #90360), 50% Hydroxylamine solution (Sigma Aldrich, #438227), Ammonium hydrogen carbonate (LiChropur, Merck, #5438350), and Iodoacetamide (Sigma Aldrich, #I1149) were obtained from Sigma Aldrich. TMT6plex and TMT10plex kits (Thermo), Urea (Pierce, Sequanal, #29700), and DTT (Thermo, #R0862) were obtained from Thermo Fischer Scientific.

#### Cell lines

HeLa cells (CCL2), MCF7 cells (HTB22), PC-12 cells (CRL1721), K562 (CCl243TM), and HEK293 (CRL321) were obtained from American Type Culture Collection (ATCC). HEK293 cells stably expressing TOM20-Halotag fusion protein was a gift from the Ploss Lab at Princeton University.

All cell lines were cultured in the recommended media supplemented with 10% Fetal Bovine Serum (Gibco, #10437-028), 1% L-glutamine (Gibco, #25030164) if required, and 1% Penicillin-Streptomycin (Gibco, #15070063) at 37 $^{o}$C and 5% $CO_2$ atmosphere in 10 cm dishes or T175 flasks according to standard practice unless otherwise stated.

#### Antibodies

*Primary antibodies:*
anti-BRD4 (A-7, Santa Cruz Biotech)
anti-histone H3 (polyclonal Invitrogen PA5-16183)
anti-α-tubulin (AB18251, Abcam)
anti-TAMRA (MA1-041, Thermo Fischer)
Anti-p38 (Cell signalling: 9212)
Anti-phos-p38 (Thr180/Tyr182) (Cell Signalling: 9211)

Anti-Abl (Cell Signalling: 2862S)
Anti-phos-Abl (Cell Signalling: 2861S)
Anti-P-Tyrosine (Cell Signalling: 9411S)
Anti-GAPDH (SCBT; 47724).
Goat-anti-Mouse 700, Goat-anti-Rabbit 700, Goat-anti-Mouse 800, Goat-anti-Mouse 700, Streptavidin 700, and Streptavidin 800 were obtained from Li-COR
Anti-$A_{2a}$ (Santa Cruz Biotech, 32261)

### General Structure Key – Iridium Catalysts and Diazirines

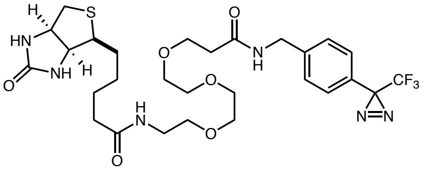

**diazirine-PEG3-biotin (9)**
*suitable for direct streptavidin enrichment*

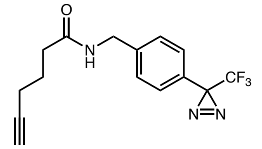

**diazirine-alkyne (10)**
*suitable for downstream CuAAC conjugation*

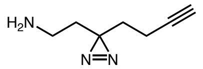

**diazirine-alkyne-amine (11)**
*conjugation handle for PAL labelling*

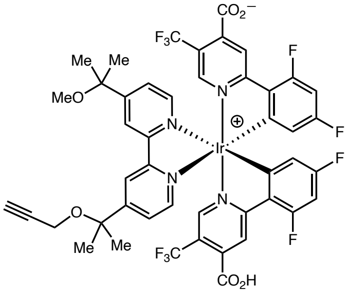

**Gen 1 Iridium photocatalyst (12)**
$[Ir(dFCF_3CO_2Hppy)_2(O\text{-}propargylbpy)]PF_6$

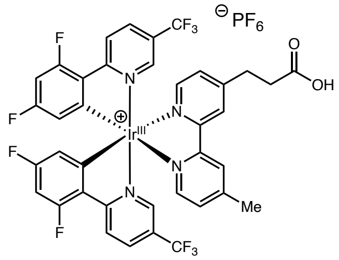

**Gen 2 Iridium photocatalyst (13)**
$[Ir(dFCF_3ppy)_2(dMebpyAcOH)]PF_6$

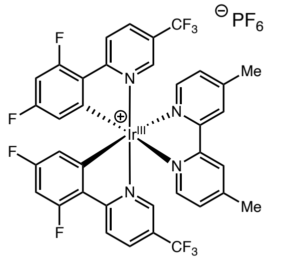

**Free Iridium catalyst (14)**
$[Ir(dFCF_3ppy)_2(dMebpy)]PF_6$

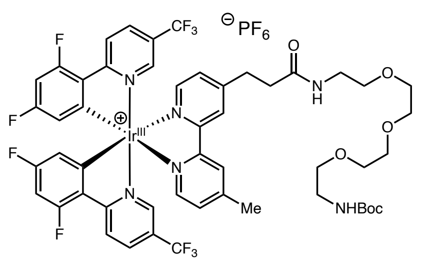

**Free Iridium catalyst (15)**
Ir-G2-PEG3-NHBoc

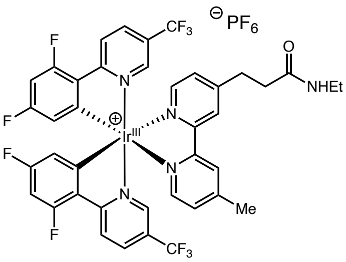

**Free Iridium catalyst (16)**
Ir-Gen2-NHEt

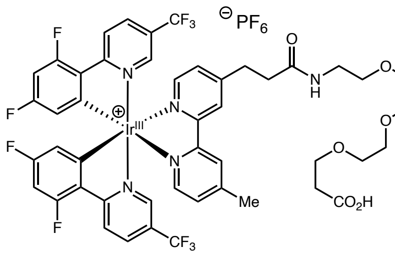

**Ir-G2-PEG3-$CO_2H$ (17)**
*conjugation handle*

#### Note on the use of free-iridium catalysts:

While multiple 'free iridium' photocatalysts have been prepared there is no specific requirement for the use of one over another. Catalyst 14 is very lipophilic and readily passes through the cell membrane leading to extensive intracellular labelling. Accordingly, it is recommended that a lower amount of the catalyst should be used for the corresponding controls compared to the drug-conjugate. Catalysts 15 and 16 better represent the solubility and cell permeability of iridium drug conjugates and can be used interchangeably with marginal differences.

### Synthesis of Iridium catalysts and Diazirines

**Dz-PEG3-biotin Synthesis (9)**

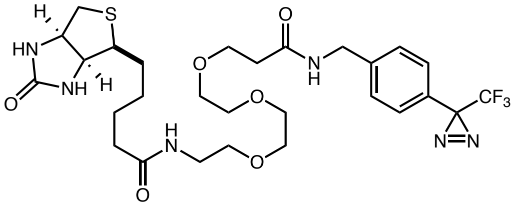

Prepared according to previously described procedure[2].

Stock solutions of the diazirine (100 mM) in DMSO can be kept frozen –20 °C to –80 °C in the dark for up to 6 months without decomposition. Deterioration of the reagent has been observed at 5 °C after several weeks. For long periods of storage it is recommended to keep as the solid in the dark at –80 °C under nitrogen. *Important: The corresponding diazo compound, formed through decomposition, leads to lower selectivity during the labelling reaction!*

While the compound is soluble in water and water-based buffers and media (DPBS, DMEM, RPMI etc.), upon addition of a concentrated stock solution a slight cloudiness can sometimes be observed. In this case, dissolution can be achieved through gentle pipetting or agitation. Alternatively, a stock solution of the diazirine can be freshly prepared in media and laid down immediately prior to irradiation.

**_N_-(4-(3-(trifluoromethyl)-3_H_-diazirin-3-yl)benzyl)hex-5-ynamide (Dz-alkyne) (10)**

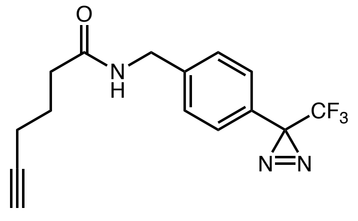

To a solution of 5-hexynoic acid (61 µL, 0.55 mmol) in anhydrous $CH_2Cl_2$ (1 mL) was added HOBt (74 mg, 0.55 mmol), EDCI (107 mg, 0.70 mmol), $Et_3N$ (103 µL, 0.73 mmol), and 4-[3-(trifluoromethyl)-3H-diazirin-3-yl]benzylamine hydrochloride (137 mg, 0.55 mmol). The reaction mixture was stirred overnight under $N_2$. The reaction mixture was diluted with EtOAc and the organic layer washed with saturated aqueous $NaHCO_3$, 0.1 M HCl, brine, and dried over $Na_2SO_4$. The solvent was removed *in vacuo*, and the crude material purified by silica column chromatography (gradient elution: 0 to 100% EtOAc/P.E.) to afford Dz-alkyne as a white solid (137 mg, 80%), which was stored at –20 ºC in the dark (no decomposition observed after 6 months). $^1$H NMR (500 MHz, DMSO) δ 8.44 (t, *J* = 6.0 Hz, 1H), 7.40 – 7.34 (m, 2H), 7.23 (d, *J* = 8.0 Hz, 2H), 4.29 (d, *J* = 6.0 Hz, 2H), 2.79 (t, *J* = 2.6 Hz, 1H), 2.25 (dd, *J* = 8.4, 6.6 Hz, 2H), 2.17 (td, *J* = 7.1, 2.7 Hz, 2H), 1.70 (p, *J* = 7.2 Hz, 2H). $^{13}$C NMR (126 MHz, DMSO) δ 172.09, 142.79, 128.50, 126.89 (d, *J* = 1.5 Hz), 126.38, 122.39 (q, *J* = 274.7 Hz), 84.45, 71.96, 42.03, 34.49, 28.48 (q, *J* = 39.9 Hz), 24.65, 17.85. $^{19}$F NMR (282 MHz, DMSO) δ -64.85.
m/z HRMS found $[M+H]^+$ = 310.12620, $[C_{15}H_{15}F_3N_3O]^+$ requires 310.11617.

*It is recommended that stock solutions in DMSO are kept in the dark at –20 $^o$C for periods >2 months. Decomposition has been overserved at 5 $^o$C after several weeks*

#### Gen 1 Iridium photocatalyst (12)

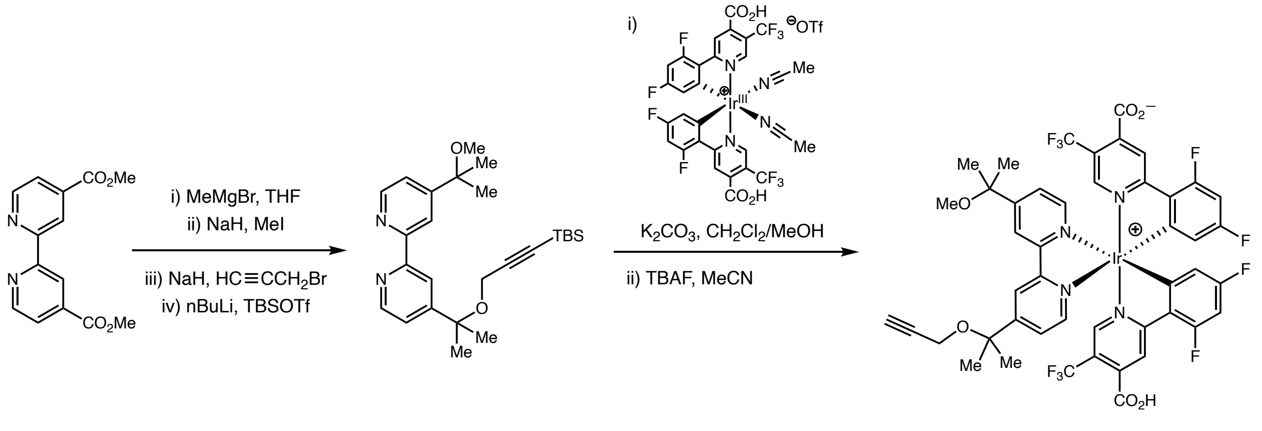

*2,2'-([2,2'-bipyridine]-4,4'-diyl)bis(propan-2-ol)*

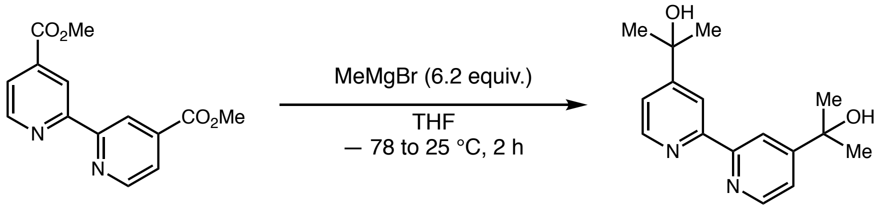

Bipyridine dicarboxylic acid dimethyl ester (11.98 g, 44 mmol) was dissolved in 500 mL anhydrous THF under nitrogen in a 1 L round bottom flask, then cooled to -78 °C. Methylmagnesium bromide (90.3 mL, 271 mmol) was rapidly added to the flask with vigorous stirring (5 cm stir bar). After continuing to stir the solution for 2 hours at room temperature, the suspension was quenched with saturated aq. NH4Cl (ca. 100 mL), evaporated to dryness, extracted into ethyl acetate (500 mL from 500 mL H2O), washed with brine (ca. 100 mL), dried over MgSO4, the solvent evaporated, and the residual solid recrystallized from boiling ethyl acetate / hexanes (75 mL EtOAc, 400 mL hexanes) to afford an off-white crystalline solid (9.0 g, 75%), which possessed spectral properties matching the reported values[2].

*2-(4'-(2-methoxypropan-2-yl)-[2,2'-bipyridin]-4-yl)propan-2-ol*

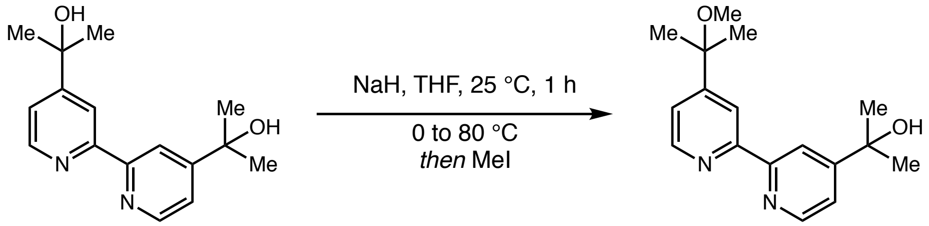

2,2'-([2,2'-bipyridine]-4,4'-diyl)bis(propan-2-ol) (4.62 g, 17 mmol) was dissolved in THF (17 mL) in a septum-equipped 40 mL vial. NaH (60% mineral oil dispersion, 350.4 mg NaH basis, 15.3 mmol) was carefully added with stirring, then the mixture stirred for 1 hour at 25 °C while vented to a bubbler through a needle puncturing the septum. Methyl iodide (0.84 mL, 13.6 mmol) was then added, the septum replaced with a cap, and the reaction mixture heated to 80 °C for 16 hours. The mixture was then quenched with saturated aq. NH4Cl (10 mL), diluted with water (100 mL), the product extracted from the mixture with DCM (4 x 50 mL), the organic phase dried with MgSO4, and solvent evaporated. The residue was purified by silica chromatography (40% ethyl acetate to 100% ethyl acetate / hexanes over 9 CV, 330 g SiO2, 180 mL/min) to afford a tan crystalline solid (1.74 g, 35%). Spectral properties matched reported values[2].

*4-(2-methoxypropan-2-yl)-4'-(2-(prop-2-yn-1-yloxy)propan-2-yl)-2,2'-bipyridine*

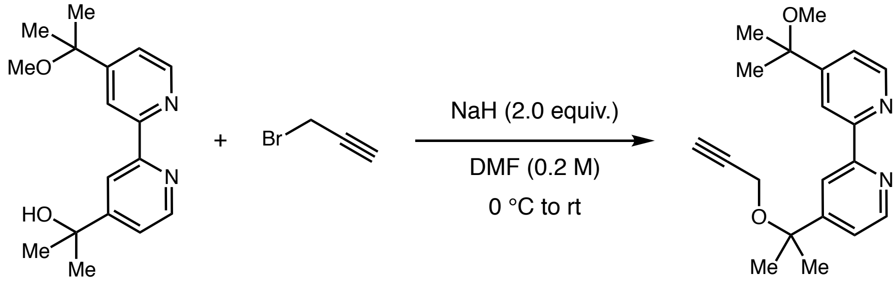

2-(4'-(2-methoxypropan-2-yl)-[2,2'-bipyridin]-4-yl)propan-2-ol (1.14 g, 4.0 mmol) was dissolved in 20 mL DMF, and NaH (137.4 mg, 6.0 mmol) was added. The mixture was stirred for 1 hour, then propargyl bromide (5.94 g, 5.0 mmol) was added and the mixture heated to 80 °C overnight. The reaction mixture was evaporated under vacuum, then purified by column chromatography (40% EtAc to 100 EtAc over 9 CV, 120 g $SiO_2$, 60 mL/min) to afford the product as a yellow oil

(0.456 g, 38%). $^{1}$H-NMR (500 MHz, $CDCl_3$): δ 8.69 (dd, 1H, J = 5.2, 0.8 Hz), 8.66 (dd, 1H, J = 5.1, 0.8 Hz), 8.39 (ddd, 2H, J = 3.0, 1.9, 0.8 Hz), 7.46 (dd, 1H, J = 5.1, 1.9 Hz), 7.40 (dd, 1H, J = 5.1, 1.8 Hz), 3.93 (d, 2H, J = 2.5 Hz), 3.15 (s, 3H), 2.41 (t, 1H, J = 2.4 Hz), 1.65 (s, 6H), 1.59 (s, 6H). $^{13}$C-NMR (126 MHz, $CDCl_3$: δ 154.21, 153.94, 153.86, 153.06, 147.07, 146.86, 118.55, 118.29, 115.91, 115.74, 78.13, 75.39, 73.98, 71.21, 49.44, 48.46, 25.28, 24.86. HRMS (ESI-TOF): m/z calcd. for $C_{20}H_{25}N_2O_2$ ([M+H]$^+$) 325.1916, found 325.18704.

*4-(2-((3-(tert-butyldimethylsilyl)prop-2-yn-1-yl)oxy)propan-2-yl)-4'-(2-methoxypropan-2-yl)-2,2'-bipyridine*

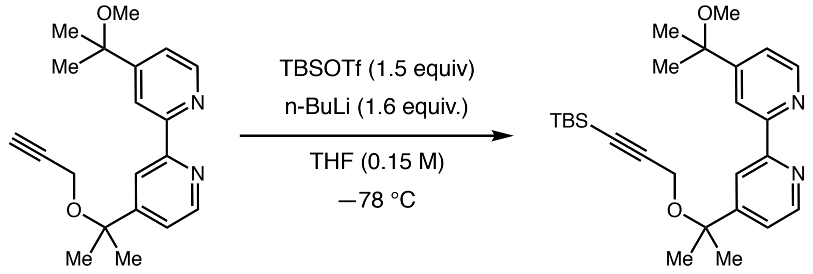

To a 40 mL vial equipped with a magnetic stir bar was added 4-(2-methoxypropan-2-yl)-4'-(2-(prop-2-yn-1-yloxy)propan-2-yl)-2,2'-bipyridine (746 mg, 2.3 mmol) and anhydrous THF (15 mL). The vial was cool to –78 °C under $N_2$, followed by addition of *n*-BuLi (1.47 mL, 3.65 mmol, 2.5 M in hexanes) in a dropwise fashion. The mixture was stirred at –78 °C for 2 hours before addition of TBSOTf (0.79 mL, 3.45 mmol). The vial was allowed to warm to room temperature and stir for 1 additional hour. The reaction mixture was then concentrated and purified via column chromatography (hexanes/EtOAc 20 to 60%) to give the product as a colorless oil (866 mg, 86% yield). $^{1}$H-NMR (500 MHz, $CDCl_3$): δ 8.68 (dd, 1H, J = 5.2, 0.8), 8.67 (dd, 1H, J = 5.2, 0.5 Hz), 8.38 (s, 2H), 7.47 (dd, 1H, J = 5.1, 1.9 Hz), 7.41 (dd, 1H, J = 5.1, 1.8 Hz), 3.96 (s, 2H), 3.15 (s, 3H), 1.64 (s, 6H), 1.59 (s, 6H), 0.91 (s, 9H), 0.09 (s, 6H). $^{13}$C-NMR (126 MHz, $CDCl_3$: δ 156.62, 156.48, 156.45, 156.07, 149.49, 149.36, 121.06, 121.00, 118.49, 118.38, 103.30, 88.65, 77.95, 76.54, 52.88, 51.01, 27.93, 27.41, 26.07, 25.95, 16.54, -4.71. HRMS (ESI-TOF): m/z calcd. for $C_{26}H_{39}N_2O_2Si$ ([M+H]$^+$) 439.2781, found 439.27540.

*TBS-protected Gen 1 Ir-photocatalyst*

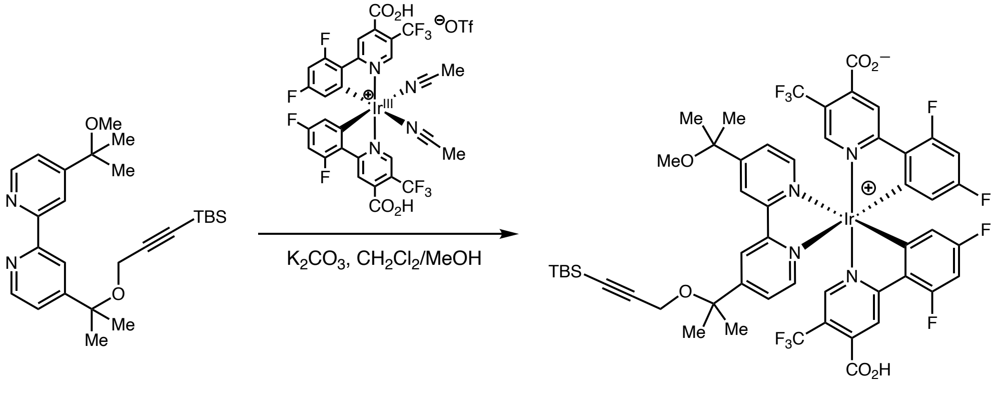

**TBS-protected Gen 1 Iridium photocatalyst**

To a 250 mL RB flask was added 4-(2-((3-(tert-butyldimethylsilyl)prop-2-yn-1-yl)oxy)propan-2-yl)-4'-(2-methoxypropan-2-yl)-2,2'-bipyridine (1.43 g, 1.39 mmol), potassium carbonate (384 mg, 2.78 mmol), [Ir(dF($CO_2H$)$CF_3$ppy)$_2$MeCN$_2$]OTf (762 mg, 1.74 mmol, *prepared according to literature procedure*) and DCM/MeOH (70 mL, 3:1 v/v). This mixture was stirred under 30 °C water bath overnight, filtered through a plug of Celite, concentrated, and purified by column chromatography ($CH_2Cl_2$/MeOH 20% to 80%) to give the product as a bright yellow powder (1.43 g, 1.03 mmol, 74% yield). $^1$H-NMR (500 MHz, MeOH-d$_4$): δ 8.80 (dd, 2H, J = 7.9, 1.8 Hz), 8.35 (s, 2H), 8.06 (dd, 2H, J = 5.8, 4.5 Hz), 7.84 (dd, 1H, J = 5.8, 1.8 Hz), 7.78 (dd, 1H, J = 5.9, 1.8 Hz), 7.47 (s, 1H), 7.44 (s, 1H), 6.75 (t, 2H, J = 10.8 Hz), 5.81 (td, 2H, J = 8.2, 2.3 Hz), 4.10 (d, 2H, J = 2.9 Hz), 3.21 (s, 3H), 1.68 (s, 3H), 1.68 (s, 3H), 1.62 (s, 3H), 1.61 (s, 3H), 0.86 (s, 9H), 0.00 (s, 6H). $^{13}$C-NMR (126 MHz, MeOH-d$_4$): δ 169.38, 167.57, 167.51, 165.90, 165.80, 163.84, 163.74, 163.65, 163.54, 161.90, 161.56, 161.46, 156.08, 155.94, 154.88, 150.79, 150.75, 145.57, 126.48, 126.22, 126.13, 125.21, 123.04, 122.32, 122.26, 121.03, 120.87, 120.75, 118.70, 113.89, 113.73, 113.65, 103.21, 99.38, 99.17, 98.95, 88.45, 77.74, 76.49, 52.26, 49.90, 26.33, 26.25, 25.76, 25.70, 25.03, 15.86, 6.03. $^{19}$F-NMR (376 MHz, $CDCl_3$): δ -62.01 (s, 6F), -80.11 (s, 3F), -104.75 (m, 2F), -108.36 (m, 2F). HRMS (ESI-TOF): m/z calcd. for $IrC_{52}H_{48}F_{10}N_4O_6Si$ ([M+H]$^+$) 1235.2807, found 1235.28010.

*Gen 1 Iridium photocatalyst (12)*

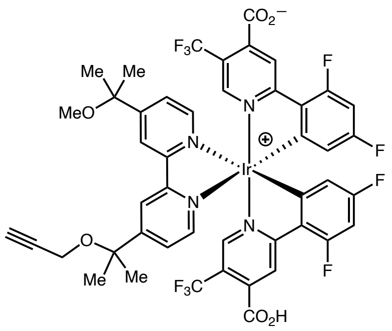

TBS-protected Gen 1 Ir photocatalyst (1.21 g, 0.8 mmol) was suspended in 80 mL acetonitrile and sonicated for one minute. To this mixture was added TBAF (1.6 mL, 1.0 M in THF) at room temperature. The reaction mixture was stirred until homogeneous and until full conversion of the starting material was overserved by mass spectrometry. The reaction mixture was then concentrated and purified successively with normal phase column chromatography (DCM/MeOH, 20-80%, reverse phase column chromatography ($H_2O$/MeOH 0-100%), then again normal phase column chromatography to afford the product **12** as a bright yellow solid (823 mg, 73% yield). $^1$H-NMR (500 MHz, $CDCl_3$): δ 8.88 (1H, d, J = 1.8 Hz), 8.83 (1H, d, J = 1.8 Hz), 8.38 (2H, s), 8.10 (2H, dd, J = 5.8, 2.0 Hz), 7.82 (2H, d, J = 5.9 Hz), 7.54 (1H, s), 7.50 (1H, s), 6.81 (2H, ddd, J = 12.0, 9.1, 2.3 Hz), 5.87 (2H, dt, J = 8.2, 2.6), 4.12 (2H, d, J = 2.4 Hz), 3.25 (3H, s), 2.86 (1H, t, J = 2.4 Hz), 1.70 (6H, s), 1.65 (6H, s). $^{13}$C-NMR (126 MHz, $CDCl_3$): δ 169.41, 167.51, 167.45, 165.88, 165.78, 163.82, 163.71, 163.63, 163.52, 161.85, 161.54, 161.44, 161.30, 155.97, 155.92, 154.83, 154.78, 151.16, 150.82, 150.79, 145.58, 126.48, 126.20, 126.10, 125.22, 123.04, 122.25, 122.18, 121.19, 121.00, 120.94, 120.87, 120.83, 120.69, 120.67, 120.41, 118.70, 113.85, 113.70, 99.36, 99.14, 98.93, 80.04, 77.86, 76.50, 74.31, 51.71, 49.91, 26.41, 26.08, 25.77, 25.69. $^{19}$F-NMR (376 MHz, $CDCl_3$): δ -60.69 (3F, s), -60.72 (3F, s), -103.43 (2F, ddd, J = 20.3, 11.8, 8.7 Hz), -107.02 (2F, tt, J = 12.4, 3.8 Hz). HRMS (ESI-TOF): m/z calcd. for $IrC_{46}H_{34}F_{10}N_4O_6$ ($[M]^+$) 1121.1948, found 1121.19531.

*Storage: Gen 1-Ir can be stored as solid in the freezer (–20 °C) without noted decomposition (> 6 months). The photocatalyst should appear luminescent (green-yellow), if the catalyst or its solutions appear yellow-orange the catalyst should be repurified by column chromatography.*

#### Gen 2 Iridium photocatalyst (13)

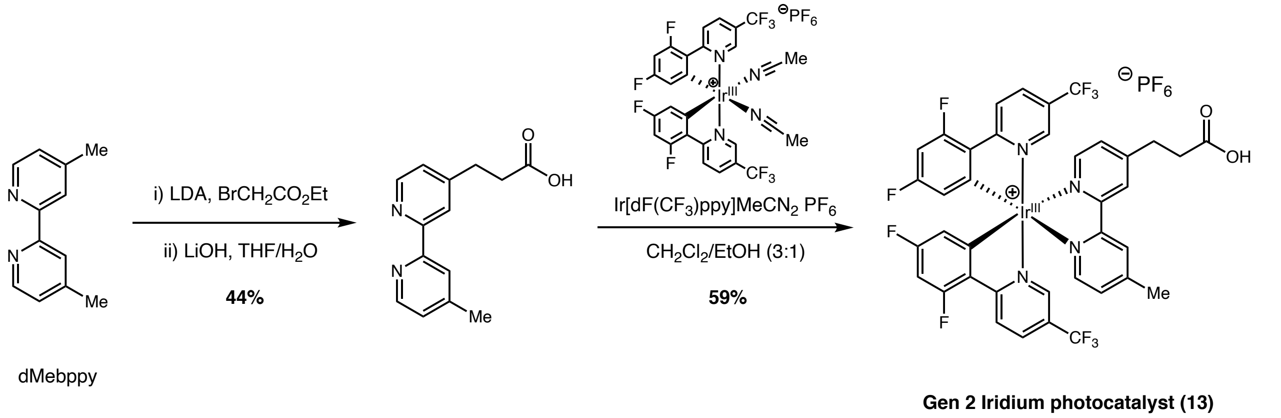

*3-(4'-Methyl-[2,2'-bipyridin]-4-yl)propanoic acid*

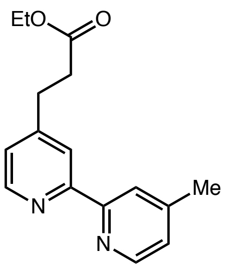

Prepared according to modified procedure[3]. 4,4'-Dimethyl-2,2'-bipyridyl (2.5 g, 13.5 mmol) was dissolved in dry THF (20 mL) under a nitrogen atmosphere in a flame-dried flask. The solution was cooled to –78 °C, and a solution of lithium diisopropylamide (14.8 mmol, 1.1 equiv) was added. The reaction mixture was allowed to warm to room temperature for 1.5 hours. This solution was cannulated into a solution of ethyl 2-bromoacetate (2.3 ml, 20 mmol) in dry THF (15 ml) at –78 °C under $N_2$. The reaction mixture was allowed to reach room temperature slowly overnight and quenched by addition of sat. sodium bicarbonate solution. Work-up using ethyl acetate followed by drying over $Na_2SO_4$ and concentration under reduced pressure provided the crude product. The crude residue was purified by column chromatography (Silica gel; DCM:MeOH:$NH_4OH$ 95:5:0.5) to provide the desired product in 69% yield.

$^1$H NMR (500 MHz, DMSO-$d_6$) δ 8.53 (dd, $J$ = 13.1, 5.0 Hz, 2H), 8.33 – 8.17 (m, 2H), 7.52 – 7.06 (m, 2H), 4.03 (q, $J$ = 7.1 Hz, 2H), 2.95 (t, $J$ = 7.4 Hz, 2H), 2.71 (t, $J$ = 7.4 Hz, 2H), 2.39 (s,

3H), 1.13 (t, $J$ = 7.1 Hz, 3H). $^{13}$C NMR (126 MHz, DMSO-d$_6$) δ 171.9, 155.3, 155.1, 150.6, 149.1, 148.9, 147.9, 125.0, 124.1, 121.3, 120.5, 60.0, 33.7, 29.7, 20.7, 14.6.

*5-(4'-Methyl-[2,2]bipyridinyl-4-yl)-pent-4-enoic acid*

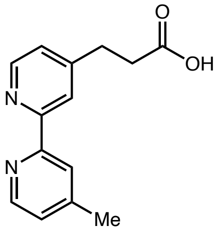

The bipyridinyl ethyl ester was taken up in 1:1 THF:$H_2O$ before the addition of LiOH (2 equiv.). The reaction mixture was stirred at room temperature for 16 h (completion by TLC) before being quenched through the addition of $NH_4Cl$ (until pH 5-6). The mixture the extracted with EtOAc, dried over $Na_2SO_4$ and concentrated under reduced pressure to provide the desired product as an off-white powder (63% yield).

$^1$H NMR (500 MHz, DMSO-d$_6$) δ 8.55 (dd, $J$ = 10.6, 5.0 Hz, 2H), 8.24 (d, $J$ = 15.4 Hz, 2H), 7.35 – 7.26 (m, 2H), 2.94 (t, $J$ = 7.4 Hz, 2H), 2.65 (t, $J$ = 7.5 Hz, 2H), 2.41 (s, 3H). $^{13}$C NMR (126 MHz, DMSO-d$_6$) δ 173.9, 155.7, 155.6, 151.5, 148.4, 125.4, 124.5, 121.7, 120.8, 34.3, 30.2, 21.2.

**Gen 2 Iridium catalyst (13)**

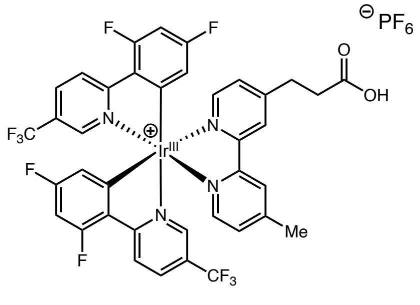

To a round bottomed flask charged with 3-(4'-methyl-[2,2'-bipyridin]-4-yl)propanoic acid (238 mg, 0.98 mmol) and Ir[dF($CF_3$)ppy]Me$CN_2$ $PF_6$ (750 mg, 0.80 mmol) was added DCM/EtOH (3:1, 10 mL) and the reaction mixture was stirred at 30 °C for 16 hours in the dark under $N_2$. The resulting solution was concentrated under reduced pressure onto silica gel and the crude product was purified by flash column chromatography (silica gel, 0-10% MeOH/DCM) to provide the desired Ir-G2 catalyst as a yellow solid (516 mg, 59%).

$^{1}$H NMR (500 MHz, $d_6$-DMSO): δ 12.31 (br, 1H), 8.84 (br, 2H), 8.46 (m, 4H), 7.83 (dd, 2H, $J$ = 10.2, 5.7 Hz), 7.67 (s, 1H), 7.65 (dd, 1H, $J$ = 5.8, 1.7 Hz), 7.59 (m, 1H), 7.50 (s, 1H), 7.08 (ddd, 2H, $J$ = 12.2, 9.3, 2.3 Hz), 5.78 (td, 2H, $J$ = 8.1, 2.4 Hz), 3.07 (t, 2H, $J$ = 7.5 Hz), 2.75 (t, 2H, $J$ = 7.5 Hz), 2.58 (s, 3H). $^{13}$C NMR (126 MHz, $d_6$-DMSO): δ 173.58, 167.24, 165.34 (d, $J$ = 13.0 Hz), 163.27 (dd, $J$ = 13.0, 6.0 Hz), 161.16 (d, $J$ = 13.4 Hz), 156.01, 155.80 (t, $J$ = 6.4 Hz), 155.61, 155.42, 153.06, 145.74 (d, $J$ = 54.4 Hz), 138.04, 130.13, 129.40, 126.91, 126.44, 125.56, 125.01 (dd, $J$ = 36.2, 2.7 Hz), 124.15 (d, $J$ = 20.4 Hz), 122.38 (dd, $J$ = 272.2, 6.8 Hz), 114.53 (t, $J$ = 15.7 Hz), 100.07 (t, $J$ = 27.0 Hz), 33.5, 30.2, 21.4. $^{19}$F NMR (376 MHz, $d_6$-DMSO): δ -61.4 (s, 3F), -61.6 (s, 3F), -70.1 (d, 6F, $J$ = 711.3 Hz), -103.3 (ddt, 2F, $J$ = 26.6, 12.1, 8.9 Hz), -106.8 (t, 2F, $J$ = 12.3 Hz). HRMS (ESI-TOF) m/z calcd. for $[M]^+$ = 951.1369, found 951.1332

*Storage: Gen 2-Ir can be stored as solid in the fridge (5 °C) without noted decomposition (> 6 months). Stock solutions should not be prepared. The photocatalyst should appear luminescent (green-yellow), if the catalyst or its solutions appear yellow-orange the catalyst should be repurified by column chromatography.*

**[Ir(dFCF$_3$ppy)$_2$(dMebpy)]PF$_6$ (14)**

Prepared according to previous reported route[4].

**Iridium-G2-PEG3-NHBoc (15)**

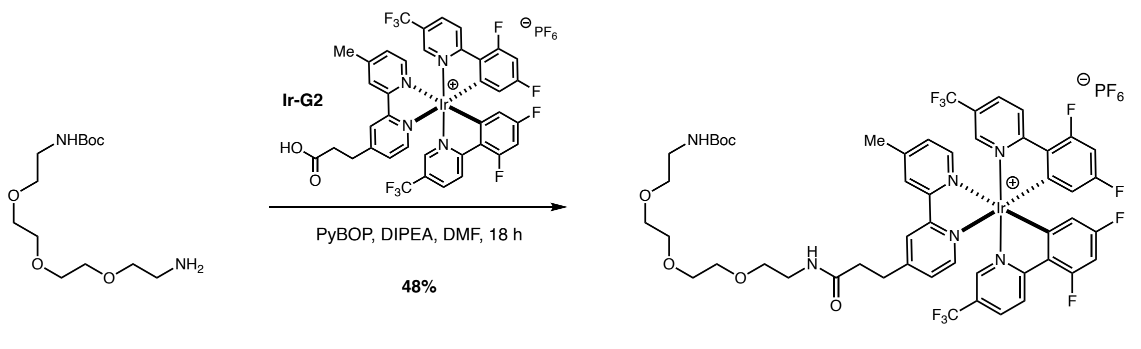

To a stirred solution of Ir-G2 (13) (50 mg, 46 μmol), and PyBOP (36 mg, 69 μmol) in anhydrous DMF (1 mL) under $N_2$ in the dark was added DIPEA (24 μL, 138 μmol). The resulting mixture was stirred at room temperature for 10 minutes and a solution of NHBoc-PEG3-$NH_2$ (40 mg, 69 μmol) in anhydrous DMF (1 mL) was added dropwise. The reaction was stirred overnight, diluted with EtOAc, and quenched by the addition of saturated aqueous $NaHCO_3$. The aqueous phase was removed and the organic layer washed with additional saturated aqueous $NaHCO_3$, 5% aqueous citric acid, brine, and dried over $Na_2SO_4$. The solvent was removed *in vacuo*, and the crude material purified by C8 reverse phase preparative HPLC (gradient elution: 30 to 100% MeCN/$H_2O$ (0.1% formic acid)) to afford NHBoc-PEG3-iridium as a yellow solid (30 mg, 48%).

$^1$H NMR (500 MHz, $CDCl_3$) δ: 8.76 (s, 1H), 8.73 (s, 1H), 8.48 (qd, *J* = 8.9, 2.1 Hz, 2H), 8.05 (app. t, *J* = 10.0 Hz, 2H), 7.74 (dd, *J* = 10.8, 5.6 Hz, 2H), 7.59 (s, 1H), 7.52 – 7.46 (m, 2H), 7.34 (d, *J* = 5.3 Hz, 1H), 6.98 (br. s, 1H), 6.65 (app. t, *J* = 10.6 Hz, 2H), 5.65 – 5.60 (m, 2H), 5.17 (br. s, 1H), 3.68 – 3.58 (m, 8H), 3.56 – 3.49 (m, 4H), 3.40 – 3.35 (m, 2H), 3.33 – 3.26 (m, 2H), 3.24 – 3.17 (m, 2H), 2.85 – 2.73 (m, 2H), 2.68 (s, 3H), 1.42 (s, 9H). $^{13}$C NMR (125 MHz, $CDCl_3$) δ: 171.6, 168.1 (dd, *J* = 14.6, 7.7 Hz), 166.1 (d, *J* = 12.2 Hz), 164.0 (d, *J* = 13.2 Hz), 163.8 (d, *J* = 13.2 Hz), 161.7 (d, *J* = 14.0 Hz), 157.5, 156.4 – 156.3 (m), 155.4, 155.2, 154.9 (d, *J* = 7.5 Hz), 154.9, 154.7 (d, *J* = 7.5 Hz), 149.7, 149.2, 144.9 (dd, *J* = 10.2, 4.4 Hz), 136.7, 129.8 (d, *J* = 22.6 Hz), 127.4, 126.6 – 126.3 (m), 126.2 (d, *J* = 3.4 Hz), 126.1, 126.0 – 125.9 (m), 123.9 (t, *J* = 21.4 Hz), 122.7 (d, *J* = 9.3 Hz), 114.2 (ddd, *J* = 18.7, 7.9, 2.3 Hz), 100.2 (td, *J* = 25.3, 10.5 Hz), 79.2, 70.5, 70.5, 70.4, 70.3, 70.2, 69.7, 40.4, 39.3, 35.4, 31.3, 29.8, 28.5, 21.6. $^{19}$F NMR (376 MHz, $CDCl_3$) δ: –62.8 (d, *J* = 35.9 Hz), –71.1, –73.0, –101.4 (dq, *J* = 38.5, 11.8 Hz), –105.8 (dt, *J* = 32.7, 12.5 Hz). m/z HRMS found $[M]^+$ = 1225.33992 (100), 1223.33115 (78), 1226.33635 (76), 1224.32887 (59),

1227.33202 (39), $[C_{51}H_{50}F_{10}IrN_6O_6]^+$ requires 1225.32559 (100), 1223.32326 (60), 1226.32894 (55), 1224.32661 (33), 1227.33230 (15). HPLC (Vydac 218TP C18 HPLC, gradient: 0 – 90% MeCN/$H_2O$ (0.1% TFA) 10 minutes, 5 minutes 90% MeCN (0.1% TFA), 1 mL/min, 260 nm): $\tau_r$ = 12.2 min

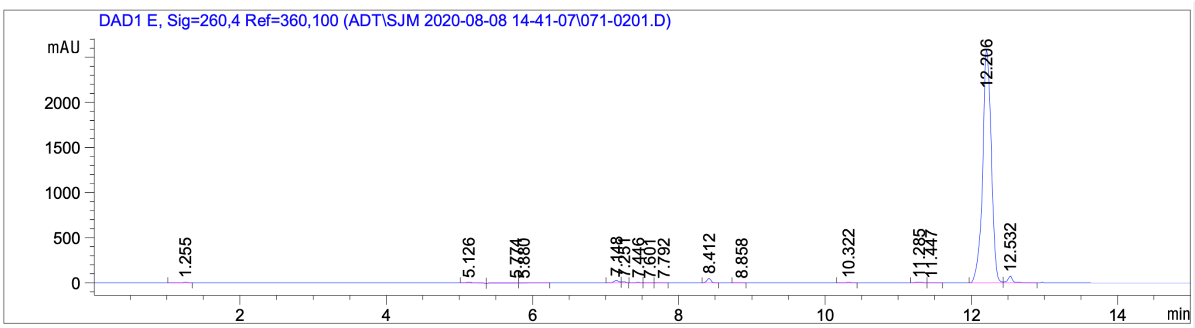

**Gen 2-Iridium NHEt (16)**

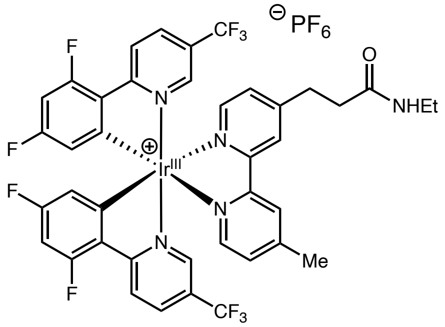

Ir-G2 (13) (53 mg, 0.05 mmol) and BOP (50 mg, 0.11 mmol) were combined in a reaction vial and then DMF (0.7 mL), ethylamine (2 M solution in THF, 0.13 mL, 0.26 mmol) and diisopropylethylamine (0.2 mL, 0.11 mL) were added. The reaction was stirred at room temperature for 2.5 hours then diluted to 2 mL with DMSO. This mixture was then filtered 0.45 µm Acrodisc then purified by preparative LCMS (43-73% acetonitrile/water (0.16%TFA) over 12 minutes, 25 mL/min, Waters CSH C18 5 µm 100 x 19 mm). The resulting product fraction was then lyophilized to give the Ir-G2-NHEt (16) (29 mg, 0.03 mmol, 56% yield) as a yellow solid. $^1$H NMR (500 MHz, Methanol-$d_4$) δ 8.72 – 8.65 (m, 2H), 8.58 (d, $J$ = 9.0 Hz, 2H), 8.33 (dt, $J$ = 8.8, 3.0 Hz, 2H), 7.94 (dd, $J$ = 16.3, 5.7 Hz, 2H), 7.73 (d, $J$ = 17.7 Hz, 2H), 7.57 (ddd, $J$ = 11.7, 5.7, 1.2 Hz, 2H), 6.82 (ddd, $J$ = 11.9, 9.2, 2.0 Hz, 2H), 5.79 (td, $J$ = 8.0, 2.3 Hz, 2H), 3.23 – 3.10 (m, 4H), 2.70 – 2.63 (m, 5H), 1.01 (t, $J$ = 7.3 Hz, 3H). $^{13}$C NMR (126 MHz, MeOD) δ 171.97, 167.80, 165.96, 165.86, 163.90, 163.79, 163.68, 161.69, 161.59, 156.36, 155.45, 155.20, 155.04, 153.78,

150.21, 149.93, 145.11, 129.60, 128.86, 125.92, 125.76, 125.48, 125.31, 125.20, 123.78, 123.61, 123.04, 120.88, 113.82, 113.73, 99.41, 99.19, 98.98, 34.91, 33.79, 30.61, 20.05, 13.32. $^{19}$F NMR (471 MHz, MeOD) δ -64.47, -64.49, -104.57 (ddt, $J$ = 17.5, 12.2, 8.8 Hz), -108.11 (t, $J$ = 12.4 Hz). HRMS (ESI-TOF) m/z calcd. for $[M]^+$ = 976.1818, found 976.1815

**Ir-G2-PEG3-$CO_2$H (17)**

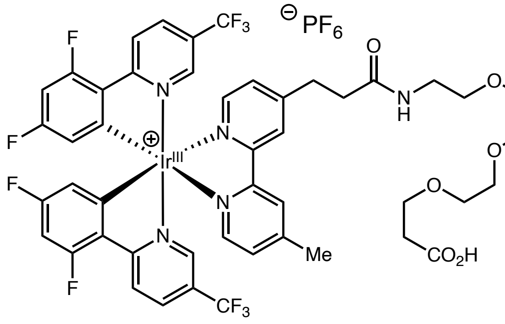

**Ir-G2-PEG3-$CO_2$H (17)**
*conjugation handle*

To a stirred solution of Ir-G2 (13) (83 mg, 76 μmol), and PyBOP (59 mg, 113 μmol) in anhydrous DMF (1.5 mL) under $N_2$ in the dark was added DIPEA (29 μL, 224 μmol). The resulting mixture was stirred at room temperature for 10 minutes and a solution of $NH_2$-PEG3-$CO_2{}^t$Bu (21 mg, 76 μmol) in anhydrous DMF (1.5 mL) was added dropwise. The reaction was stirred overnight, diluted with EtOAc, and quenched by the addition of saturated aqueous $NaHCO_3$. The aqueous phase was removed and the organic layer washed with additional saturated aqueous $NaHCO_3$, 5% aqueous citric acid, brine, and dried over $Na_2SO_4$. The solvent was removed *in vacuo*, and the crude material purified by C8 reverse phase preparative HPLC (gradient elution: 30 to 100% MeCN/$H_2O$ (0.1% formic acid)) to afford Ir-G2-PEG3-$CO_2{}^t$Bu as a yellow solid (65 mg, 63%). HRMS (ESI-TOF) m/z calcd. for $[M]^+$ = 1210.31524, found 1210.31612.

*Ir-G2-PEG3-$CO_2{}^t$Bu can be stored as solid in the fridge (5 °C) without noted decomposition (> 6 months). Deprotection is performed immediately prior to use.*

Ir-G2-PEG3-$CO_2{}^t$Bu (47 mg) was dissolved in $CH_2Cl_2$ (5 mL) and cooled to 0 °C under $N_2$. TFA (2 mL) was added dropwise and the solution warmed to r.t. overnight in the dark. The solvent was removed *in vacuo* and sample further dried under hi-vac. The resulting Ir-G2-PEG3-$CO_2H$ (17) was used directly without purification.

$^1$H NMR (500 MHz, MeOD) δ 8.70 (d, $J$ = 4.1 Hz, 2H), 8.56 (dt, $J$ = 7.5, 3.4 Hz, 2H), 8.30 (t, $J$ = 8.2 Hz, 2H), 7.94 (d, $J$ = 5.7 Hz, 1H), 7.90 (d, $J$ = 5.6 Hz, 1H), 7.74 (s, 1H), 7.70 (s, 1H), 7.58 (d, $J$ = 5.7 Hz, 1H), 7.53 (d, $J$ = 5.7 Hz, 1H), 6.78 (ddt, $J$ = 12.0, 9.1, 2.7 Hz, 2H), 5.76 (ddd, $J$ = 10.5, 8.0, 2.2 Hz, 2H), 3.67 (t, $J$ = 6.2 Hz, 2H), 3.61 – 3.55 (m, 4H), 3.52 (d, $J$ = 4.5 Hz, 2H), 3.43 (dp, $J$ = 9.5, 4.7 Hz, 2H), 3.20 (t, $J$ = 7.3 Hz, 2H), 2.70 (t, $J$ = 7.3 Hz, 2H), 2.64 (s, 3H), 2.47 (t, $J$ = 6.3 Hz, 2H). $^{13}$C NMR (126 MHz, MeOD) δ 173.8, 172.4, 167.8 (d, $J$ = 6.8 Hz), 165.9 (d, $J$ = 12.7 Hz), 163.9 (d, $J$ = 12.7 Hz), 163.7 (d, $J$ = 12.9 Hz), 161.6 (d, $J$ = 13.1 Hz), 156.3, 155.4, 155.3, 155.1 (t, $J$ = 7.7 Hz), 153.8, 150.3, 149.9, 145.2 (dd, $J$ = 11.9, 5.0 Hz), 136.9 (t, $J$ = 3.6 Hz), 129.6, 128.8, 126.5 (d, $J$ = 4.1 Hz), 126.0, 125.8, 125.5, 125.4, 125.2, 123.8 (d, $J$ = 3.8 Hz), 123.6 (d, $J$ = 3.8 Hz), 121.97 (d, $J$ = 271.8 Hz), 115.30 – 111.94 (m), 99.20 (t, $J$ = 27.1 Hz), 70.16, 70.03, 69.92, 69.7, 69.0, 66.4, 38.9, 34.7, 34.4, 30.6, 20.1. $^{19}$F NMR (376 MHz, MeOD) δ -64.39 (d, $J$ = 28.1 Hz), -76.93, -104.48, -108.02 (d, $J$ = 7.1 Hz). HRMS (ESI-TOF) m/z calcd. for $[M]^+$ = 1154.25264, found 1154.25445.

##### Gen 1-Iridium PEG4-$C_6H_{12}$Cl (18)

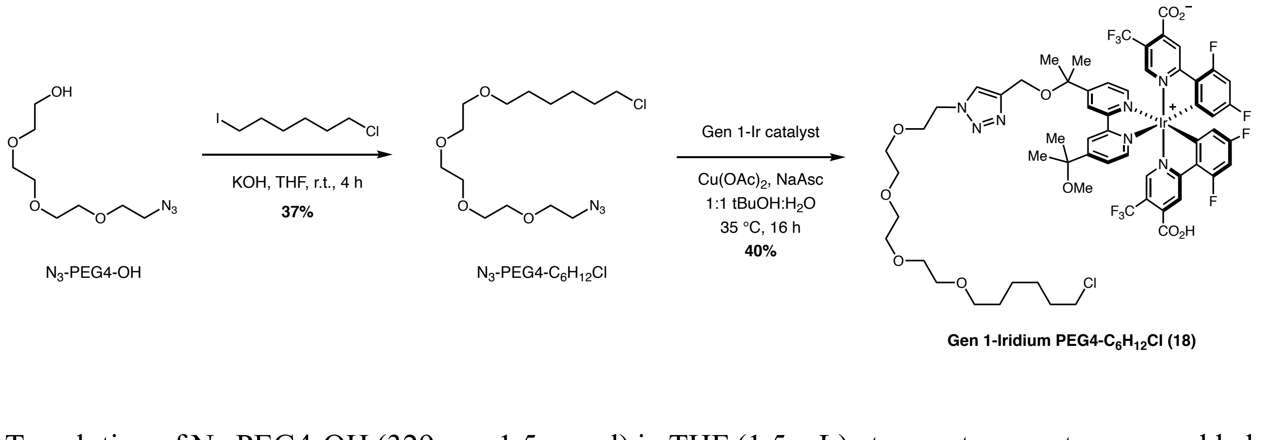

Gen 1-Iridium PEG4-$C_6H_{12}$Cl (18)

To solution of $N_3$-PEG4-OH (329 mg, 1.5 mmol) in THF (1.5 mL) at room temperature was added KOH (85%, 84 mg, 1.5 mmol). The solution was sonicated and stirred vigorously for 1 hour before the addition of 6-chloro-1-iodohexane (0.23 mL, 1.5 mmol) in THF (1.5 mL) dropwise at room temperature. The mixture was vigorously stirred for an additional 4 hours in the dark and quenched by the addition of brine (30 mL). The mixture was extracted with EtOAc (3 x 20 mL), dried over $Na_2SO_4$, and the solvent removed *in vacuo*. Purification by silica column chromatography (gradient elution: 0 to 50% EtOAc/hexane) gave the product $N_3$-PEG4-$C_6H_{12}Cl$ as a pale-yellow oil (194 mg, 37%). $^1H$ NMR (500 MHz, $CDCl_3$) δ: 3.69 – 3.64 (m, 12H), 3.58 (dd, $J$ = 5.3, 3.3 Hz, 2H), 3.53 (t, $J$ = 6.6 Hz, 2H), 3.46 (t, $J$ = 6.6 Hz, 2H), 3.39 (t, $J$ = 5.1 Hz, 2H), 1.78 (quin, $J$ = 7.3 Hz, 2H), 1.60 (quin, $J$ = 7.7 Hz, 2H), 1.45 (quin, $J$ = 7.7 Hz, 2H), 1.37 (quin, $J$ = 7.3 Hz, 2H). $^{13}C$ NMR (125 MHz, $CDCl_3$) δ: 71.3, 70.8, 70.7 (multiple peaks), 70.2, 70.1, 50.7, 45.1, 32.6, 29.5, 26.8, 25.5. m/z HRMS found $[M+H]^+$ = 338.18141, $[C_{14}H_{29}ClN_3O_4]^+$ requires 338.18411.

An 8 mL vial was charged with a magnetic stirrer bar, $N_3$-PEG4-$C_6H_{12}Cl$ (4.1 mg), Ir-alkyne (13.7 mg), $CuSO_4 \cdot 5H_2O$ (6.2 mg), sodium ascorbate (24.7 mg), and 1:1 *t*-butanol:$H_2O$ (2.5 mL). The mixture was stirred overnight at 37 °C in the dark for 18 hours. The solvent was subsequently removed *in vacuo* and purified by C8 reverse phase preparative HPLC (gradient elution: 30 to 100% MeCN/$H_2O$ (0.1% formic acid)) to afford Ir-G2-PEG4-$C_6H_{12}Cl$ (18) as a yellow solid (7 mg, 40%).

**Ir-PEG4-$C_6H_{12}$Cl (19)**

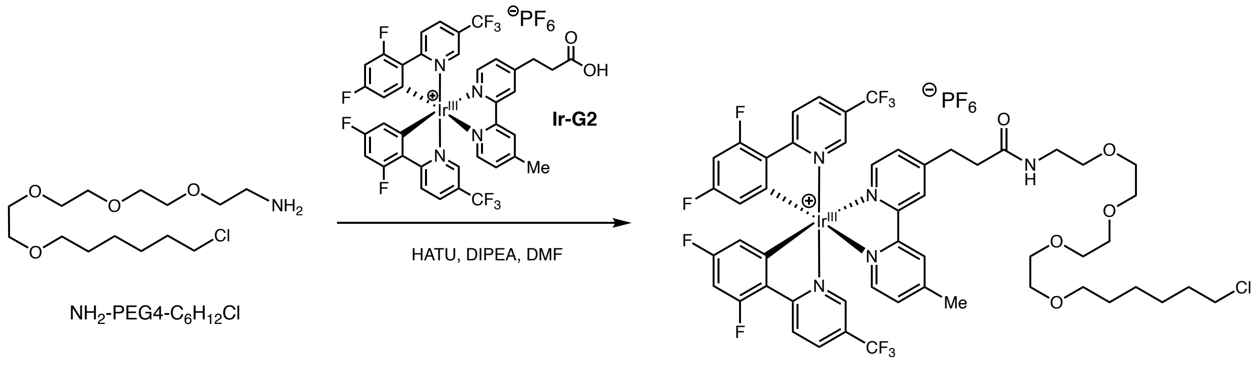

To a solution of Ir-G2 (13) (333 mg, 304 µmol) in DMF (1 mL) was added DIPEA (106 µL, 511 µmol) and HATU (237 mg, 456 µmol). The resulting suspended solution was stirred for at least 15 min under air. Meanwhile, to a stirring solution of $NH_2$-PEG4-$C_6H_{12}Cl$[5] (155 mg, 365 µmol) in DMF (2 mL) at room temperature was added DIPEA (106 µL, 511 µmol); this solution was stirred for at least 5 min. The Ir/HATU solution was then added to the amine solution, and the mixture stirred for 4 hours until completion. The solvent was subsequently removed *in vacuo* and purified by C8 reverse phase preparative HPLC (gradient elution: 30 to 100% MeCN/$H_2O$ (0.1% formic acid)) to afford Ir-G2-PEG4-$C_6H_{12}Cl$ (19) as a bright yellow solid (378 mg, 49%). $^1H$ NMR (500 MHz, Acetone-d6) δ: 8.82 (s, 2H), 8.64 (ddd, *J* = 8.3, 5.2, 2.5 Hz, 2H), 8.42 (ddd, *J* = 8.5, 5.8, 2.2 Hz, 2H), 8.19 – 8.07 (m, 2H), 8.02 (d, *J* = 2.1 Hz, 1H), 7.97 – 7.88 (m, 1H), 7.71 – 7.62 (m, 2H), 7.27 (q, *J* = 6.8, 6.3 Hz, 1H), 6.83 (ddt, *J* = 12.6, 9.6, 3.1 Hz, 2H), 5.97 (ddd, *J* = 8.7, 6.4, 2.4 Hz, 2H), 3.58 (q, *J* = 8.8, 7.7 Hz, 12H), 3.53 (dt, *J* = 6.5, 3.4 Hz, 4H), 3.44 (dt, *J* = 13.1, 5.8 Hz, 4H), 3.32 (p, *J* = 5.6 Hz, 2H), 3.21 (t, *J* = 7.4 Hz, 2H), 2.70 (t, *J* = 7.4 Hz, 2H), 2.65 (d, *J* = 4.4 Hz, 3H), 1.76 (p, *J* = 6.8 Hz, 2H), 1.54 (p, *J* = 6.7 Hz, 2H), 1.45 – 1.32 (m, 2H). $^{19}F$ NMR (470 MHz, Acetone) δ: –63.35, –71.46, –72.96, –104.65, –107.84.

### Chloroalkane Penetration Assay (CAPA)

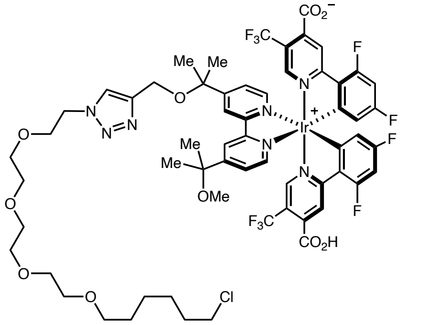

**Gen 1-Iridium PEG4-$C_6H_{12}Cl$ (18)**

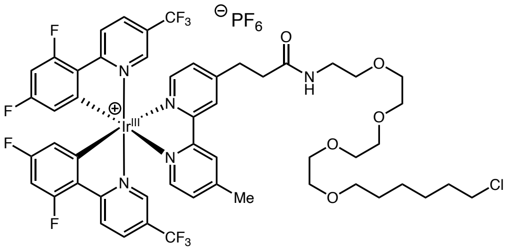

**Gen 2-Iridium PEG4-$C_6H_{12}Cl$ (19)**

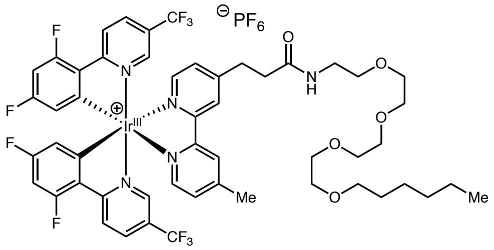

**Ir-G2-PEG4-hexyl (Ir-Me, 20)**
*no affinity for halotag*

HEK293T cells expressing TOM20-Halotag proteins were grown to 90% confluency in three 6-well cell culture dishes using DMEM. Ir-Cl Gen1 (18), Ir-Cl Gen2 (19), Ir-Me (10 mM stock solutions in DMSO), or DMSO were added to the appropriate wells to 5 or 10 µM final catalyst concentration. The cells were incubated for 1 or 2 hours at 37 °C, aspriated, then gently washed 2 x 2 mL x 10 min with fresh DMEM. TAMRA-Cl fluorescent dye (Promega, G8251) (5 mM) was diluted with OptiMEM to a 5 µM working solution. The washed cells were aspirated again, added 0.8 mL of the working dye solution, then incubated at 37 °C for 15 min.

Ir-Cl Gen 1 and Gen2 was incubated for 1 h (10 mM or 5 uM stock solution in DMSO, added to 2 mL media)

| Well | 1 | 2 | 3 | 4 | 5 | 6 |
|---|---|---|---|---|---|---|
| Compound | DMSO | Ir-Me | Ir-Cl Gen1 | | Ir-Cl Gen2 | |
| Concentration | – | 10 uM | 5 uM | 10 uM | 5 uM | 10 uM |
| Stock | 2 uL | 1 uL | 1 uL | 2 uL | 1 uL | 2 uL |

Ir-Cl Gen 1 and Gen2 was incubated for 1 or 2 h (10 uM). Three replicates of 6-well plates.

| Well | A | B | C | D | E | F |
|---|---|---|---|---|---|---|
| Compound | DMSO | Ir-Me | Ir-Cl Gen1 | | Ir-Cl Gen2 | |
| Time | 2 h | 2 h | 1 h | 2 h | 1 h | 2 h |
| Stock (10 mM) | 2 uL | 2 uL | 2 uL | 2 uL | 2 uL | 2 uL |

The cells were suspended into 1.5 mL eppendorf tubes, washed 2 x 1 mL with cold DPBS, then resuspended with 200 μL RIPA (1x cOmplete protease inhibitor). Sonicated at 35% power, 5 x 5 sec, with 30 sec rest on ice in between cycles. The lysate was centrifuged at 15,000 g for 20 min at 4 °C. The supernant was removed, concentration adjusted to 0.7 mg/mL via BCA assay. Cell lysates were processed and analyzed by western blot with anti-TAMRA and anti-GAPDH (for normalization) antibodies. TAMRA signal is inversely proportional to Ir binding to intracellularly expressing TOM20-Halotag, and therefore inversely proportional to Ir cell permeability.

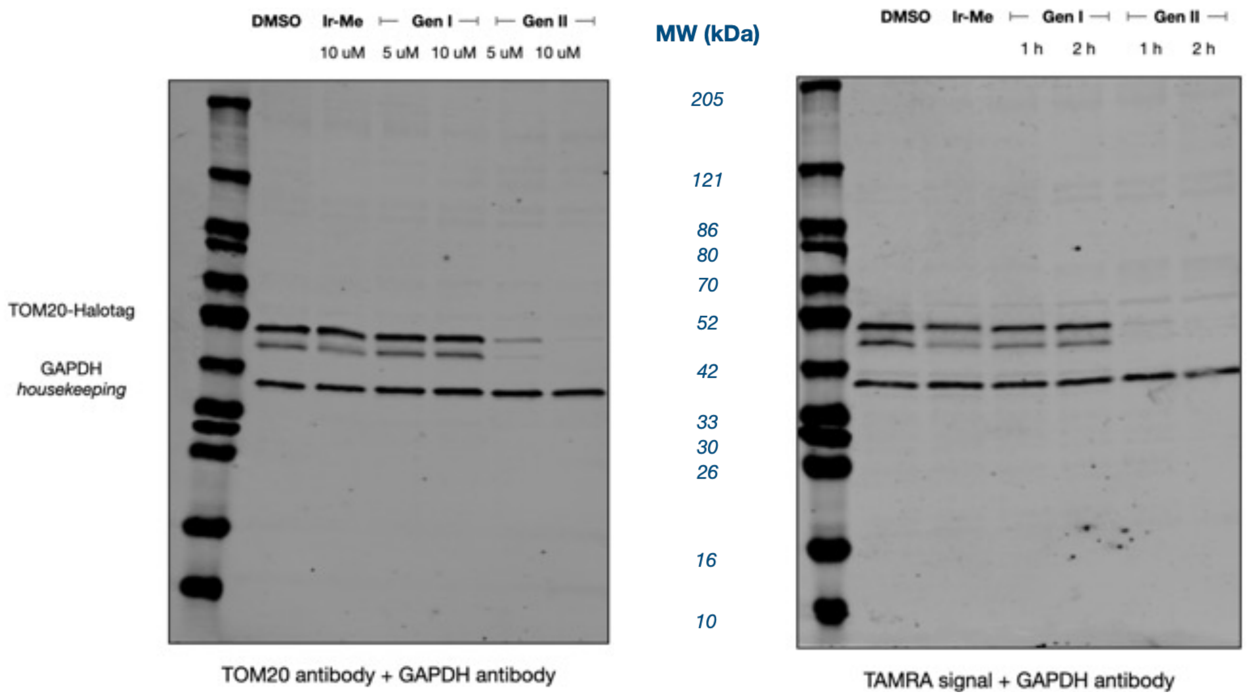

#### Preparation of G2-Ir conjugates: Tips and Tricks

Particular importance should be placed on selecting a ligation point on the small molecule that shows minimal disruption to biological activity and target binding.

We have found that a no SAR is typically required for the linker between the small molecule and Iridium catalyst. If common spacer units are employed for certain small molecules conjugates, such as PROTACs, dyes, and contrast reagents (e.g. Paclitaxel and 4-aminobutyric acid spacer has been employed previously for the synthesis of MRI contrast reagents) then it is advisable to follow such examples. Otherwise, a PEG3 spaces linker is usually sufficient.

*General Procedure:*

To a darkened 8 mL vial was added Ir-G2 (13) (32 mg, 25 μmol) and PyBOP (20 mg, 1.5 equivalents) and a magnetic stir bar. *The combination of these coupling reagents is paramount to avoid decomposition of the amine and subsequent coupling to the Ir-G2 catalyst.* The vial was closed with a cap containing a septa and placed under nitrogen gas (3 times vacuum cycling). Anhydrous DMF (2 mL) was added and the vial placed into an ice bath. To the cooled solution was added DIPEA (13 μL, 3 equivalents) and the reaction mixture stirred for an additional 30 minutes. After such time, the desired small molecule amine (25 μmol, 1 equivalent) in anhydrous DMF (1 mL) was added dropwise and the resulting mixture was warmed to room temperature with stirring overnight. The contents were diluted with EtOAc (50 mL), and quenched by the addition of saturated aqueous $NaHCO_3$ (50 mL). The organic layer washed with additional saturated aqueous $NaHCO_3$ (50 mL), 5% aqueous citric acid (50 mL), brine (50 mL), and dried over $Na_2SO_4$. The solvent was removed *in vacuo*, and the crude material purified by C8 reverse phase preparative HPLC (gradient elution: 0 to 100% MeCN/$H_2O$ (0.1% formic acid)) to afford the iridium conjugate as a yellow solid.

The photocatalyst shows diagnostic signals by $^{19}F$ NMR (376 MHz, $CDCl_3$) δ: –62.8 (d, $J$ = 35.9 Hz), –71.1, –73.0, –101.4 (dq, $J$ = 38.5, 11.8 Hz), –105.8 (dt, $J$ = 32.7, 12.5 Hz).

No decomposition of the solid Ir-G2-conjugates has been observed at – 20 °C over extended periods (> 6 months). However, particular care should be given to small molecules containing oxidatively labile functionality during synthesis, purification, and storage to exclude light and oxygen. Stock solutions of the small molecule conjugates in DMSO can be prepared (5 mM to 10 mM) and stored at – 20 °C until use. Decomposition (darkening of solution) has been observed after extended periods of storge at 5 °C (> 6 months) with certain conjugates.

| **Problem** | **Possible causes** | **Solutions** |
|---|---|---|
| Low yield of small molecule conjugate | Decomposition of iridium photocatalyst during ligation<br><br>Small molecule conjugate insoluble in EtOAc<br><br>Small molecule conjugate is too soluble in aqueous phase<br><br>Small molecule conjugate too polar for silica gel chromatography | **Use only** PyBop and DIPEA as coupling reagents<br><br>Perform coupling in dark under inert atmosphere<br><br>Extract using more solubilizing solvent (e.g. DCM, n-BuOH)<br><br>Evaporate DMF and load directly onto HPLC for purification |
| Small molecule conjugate contains no appropriate conjugation handle (e.g. amine, alcohol) | | Consider employing alternative bioconjugation handle such as azide; the corresponding Ir-G2-DBCO conjugate has been prepared and used in conjugation reactions<br>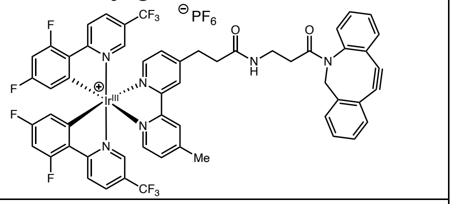 |
| Small molecule is particularly valuable/low quantity available | | Reverse order or coupling; prepare Ir-G2-PEG3-$NH_2$ or Ir-G2-PEG3-$CO_2H$ and couple to small molecule<br><br>Use Ir-G2-DBCO for Cu-free click conjugation to small molecule azide |

#### **Design of experiment for μMap photocatalytic target ID**

For new small molecule conjugates it is advisable to ascertain labelling efficiency in vitro. A typical procedure involves irradiating a mixture of recombinant target protein, a competitor protein (e.g. carbonic anhydrase, BSA etc), the small molecule conjugate and the diazirine (9). Additional controls such as off-competing with the parent ligand and a free (unconjugated) iridium catalyst should also be included. Western blot analysis and immunostaining with streptavidin shows fold enrichment of target protein vs controls (typically 3.5x to 7.5x).

Prior to proteomic analysis it is recommended to perform WB analysis checking for enrichment of the target protein by immunostaining. Experiments can be performed either in plates or Eppendorf tubes depending on quantity of reagents available and cell line. It is important to note that certain molecules may cause cells to loose adhesion after extended periods of incubation. Irradiation of plates is conducted in Efficiency Aggregators Bio-photoreactor (up to 6 plates stacked). Irradiation of tubes is conducted in either Efficiency Aggregators Bio-photoreactor or PennOptic photoreactor.

*Typical procedure for WB analysis in tubes:*

1. Remove cells (recommended 50 million per labeling experiment) from culture and pellet according to manufacturer's guidelines (e.g. 1,000xg for 4 min at 4 °C). *Note: For membrane based targets use non-enzyme based dissociation reagent.*

*2.* Resuspend cells in media (1 mL) and distribute evenly into 1.5 mL Eppendorf tubes.

3. Add the desired small molecule conjugates/controls as a solution in DMSO to a final concentration of between 1 – 10 μM (suggested off-compete 20X i.e. 20 – 200 μM). *Note: Final DMSO concentration should not exceed 0.5%.*

*For small molecule target ID the length of time and concentration of reagent during incubation is dependent upon the binding and potency of the individual molecule. The following protocol in exemplary but for individual cases may require optimization. Note Ir-catalyst is cytotoxic over extended periods of incubation and so less than 6 hours is recommended.*

*While small molecule conjugates are freely soluble in DMSO, upon addition to media, precipitation may occur. It is therefore advisable to check solubility prior to addition to cells. In the event that precipitation occurs dilute DMSO stock solution so a larger relative volume is added. Gentle pipetting and agitation can be used to aid dissolution.*

***Exemplar reaction set up***

| Tube: *Cells suspended in 1 mL media* | 1 | 2 | 3 | 4 | 5 | 6 | 7 | 8 | 9 |
|---|---|---|---|---|---|---|---|---|---|
| Ir-small molecule conjugate (5 mM) | 1 µl | 1 µl | 1 µl | – | – | – | 1 µl | 1 µl | 1 µl |
| Ir-control (5 mM) | – | – | – | 1 µl | 1 µl | 1 µl | – | – | – |
| Small molecule (20 mM) | – | – | – | – | – | – | 5 µl | 5 µl | 5 µl |

*Note: Additional DMSO is added evenly to all tubes to maintain constant volume*

4. Incubate cells on a rotisserie at 37 °C for 1 – 3 hours in the dark.

5. Centrifuge cells and remove supernatant.

6. Wash cells twice with 1 mL cold DPBS by gentle resuspension using a pipette followed by centrifugation.

7. Suspend cell pellet gently using a pipette in 1 mL of phenol red-free media containing 250 µM diazirine biotin and incubate in the dark for 15 minutes at 37 °C (DPBS can also be used at room temperature with similar results). *Note: For membrane targets incubate at 4 °C for 5 minutes.*

8. The samples were placed in the Merck Photoreactor MS2 and irradiated on 100% LED power for 3 minutes.

9. Wash cells twice with 1 mL cold DPBS by gentle resuspension using a pipette followed by centrifugation.

10. Wash cells twice with 1 mL cold DPBS by gentle resuspension using a pipette followed by centrifugation.

*While RIPA buffer is sufficient for most applications, lysis procedure should be conducted according to best practice for the target protein/class/compartment. For membrane targets it is strongly advisable to perform membrane fractionation of the samples at this stage and proceed with the streptavidin bead enrichment.*

The cells were suspended in 1 mL of cold RIPA buffer containing PMSF (1mM) and cOmplete EDTA free protease inhibitor (1x) (Roche). The lysed cells were incubated on ice for 5–10 minutes and sonicated (35%, 5 x 5s with 30s rest). The lysate was then centrifuged at 15x1000g for 15 mins at 4 °C and the supernatant collected. The concentration of the cell lysate was measured by BCA assay and adjusted accordingly to a concentration of 1 mg/mL with RIPA.

Magnetic Streptavidin beads (NEB or Pierce) were removed (9 x 250 µL) into 9 x 1.5 mL Lo-Bind tubed and washed twice with RIPA (0.5 mL) (5 minutes incubation on a rotisserie). The beads were pelleted on a magnetic rack, diluted with the samples (1 mL per tube) and incubated on a rotisserie at 4 °C overnight. The beads were pelleted on a magnetic rack, the supernatant removed. The beads were subsequently washed with 3 x 1% SDS in DPBS (1 mL), 3 x 1M NaCl in DPBS (1 mL), 3 x 10% EtOH in DPBS (1 mL). The samples were incubated with each wash for 5 minutes prior to pelleting. The beads were resuspended in RIPA buffer (0.5 mL) and transferred into 9 x 1.5 mL Lo-bind tubes.

For Western blot analysis: The beads were pelleted and the supernatant removed and the beads subsequently resuspended in freshly prepared elution buffer (30 mM biotin, 6 M urea, 2 M thiourea, 2% SDS in DPBS, pH = 11.5) (24 µL) and 4x Laemlli buffer with BME (6 µL) was added with gentle mixing. The beads were heated to 95 °C for 15 minutes, pelleted on a magnetic rack, and the supernatant was removed while hot and analysed by western blot.

| | | |
|---|---|---|
| Target protein not enriched | Protein concentration too low<br><br>Small molecule conjugate has lost binding efficiency<br><br>Labelled protein not soluble in lysis buffer | Increase number of cells<br><br>Fractionate cells prior to streptavidin enrichment<br><br>Ensure correct lysis conditions used<br><br>Check labelling efficiency of small molecule using recombinant protein *in vitro* followed by western blot analysis and streptavidin staining<br><br>Check labelling efficacy in cell lysate<br><br>Increase concentration of probe conjugate |

*Proteomics experiment:*

Both chemoproteomic and label free proteomics have been conducted for μMap target ID. It is, however, usually preferable to increase the number of cells for TMT-based proteomics experiments (adjusting the number of plates for TMT6 or TMT10 plex). We routinely conduct experiments with three replicates.

Reactions are conducted identically to WB experiments up until the final washings of the streptavidin beads. After which, the supernatant was removed and the beads washed with 3 x DPBS (0.5 mL) and 3 x $NH_4HCO_3$ (100 mM) (0.5 mL). The beads were re-suspended in 500 μL 6 M urea in DPBS and 25 μL of 200 mM DTT in 25 mM $NH_4HCO_3$ was added. The beads were incubated at 55 $^oC$ for 30 min. Subsequently, 30 μL 500 mM IAA in 25 mM $NH_4HCO_3$ was added and incubated for 30 min at room temperature in the dark. The supernatant was removed and the beads washed with 3 x 0.5 mL DPBS and 3 x 0.5 mL TEAB (50 mM). The beads were resuspended in 0.5 mL TEAB (50 mM) and transferred to a new protein LoBind tube, pelleted, and the supernatant removed. The beads were resuspended in 40 μL TEAB (50 mM) and 1.2 μL trypsin (1 mg/mL in 50 mM acetic acid) was added and the beads incubated overnight on a rotisserie at 37 $^oC$. After 16 hours, an additional 0.8 μL trypsin was added and the beads incubated for an additional 1 hour on a rotisserie at 37 $^oC$. Meanwhile, the TMT label reagents (0.8 mg) (Thermo) were equilibrated to room temperature and diluted with 41µL of anhydrous acetonitrile (Optima grade; 5 min with vortexing) and centrifuged. The beads were subsequently pelleted and the supernatant transferred to the corresponding TMT-label. The reaction was incubated for 2 hours at room temperature. The samples were quenched with 8µL of 5% hydroxylamine and incubated for 15 minutes. All of the samples were pooled in a new Protein LoBind tune and quenched with TFA (16µL, Optima). The samples were stored at –80 °C until proteomics were conducted. Samples were desalted and fractionated (high pH) and combined prior to running (3 injections).

### Labelling of the bromodomain using JQ1-Ir

#### Synthesis of conjugates

##### (+)-JQ1-PEG3-G2-iridium (1)

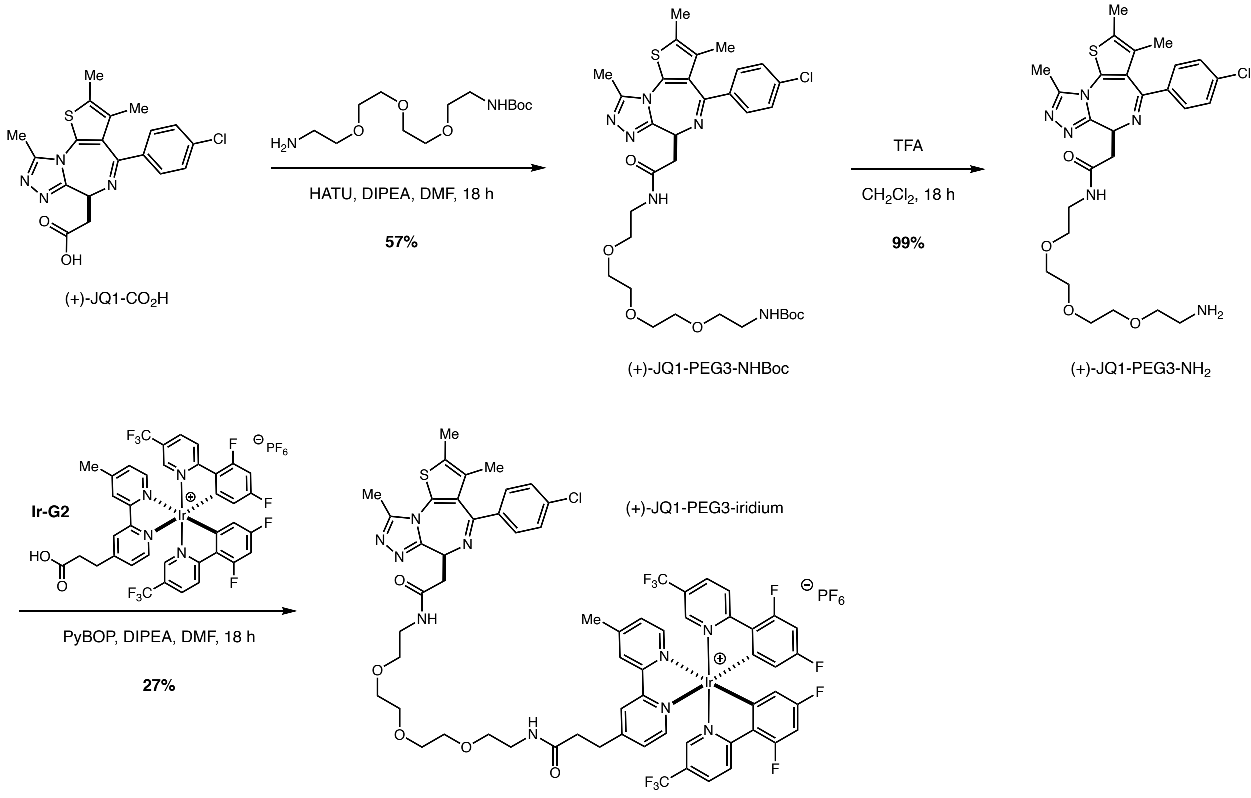

To a stirred solution of (+)-JQ1-$CO_2H$ (177 mg, 0.44 mmol) in anhydrous DMF (4.5 mL) was added HATU (176 mg, 0.46 mmol) followed by DIPEA (230 µL, 1.32 mmol). The reaction was stirred at room temperature for 10 minutes under $N_2$ and a solution of *t*-Boc-*N*-amido-PEG3-amine (143 mg, 0.49 mmol) in anhydrous DMF (0.5 mL) was added dropwise. The resulting mixture was stirred overnight, diluted with EtOAc, and quenched by the addition of saturated aqueous $NaHCO_3$. The aqueous phase was removed and the organic layer washed with additional saturated aqueous $NaHCO_3$, brine, and dried over $Na_2SO_4$. The solvent was removed *in vacuo*, and the crude material purified by silica column chromatography (gradient elution: 0 to 10% MeOH/$CH_2Cl_2$) to afford (+)-JQ1-PEG3-NHBoc as a tan solid (171 mg, 57%). $^1$H NMR (500 MHz, $CDCl_3$) δ: 7.39 (d, *J* = 8.5 Hz, 2H), 7.31 (d, *J* = 8.7 Hz, 2H), 7.20 (br. s, 1H), 5.35 (br. s, 1 H), 4.65 (t, *J* = 7.1 Hz, 1H), 3.69 – 3.46 (m, 15H), 3.36 (dd, *J* = 15.0, 6.8 Hz, 1H), 3.30 (m, 2H), 2.65 (s, 3H), 2.39 (s,

3H), 1.66 (s, 3H), 1.41 (s, 9H). $^{13}$C NMR (125 MHz, $CDCl_3$) δ: 170.7, 164.0, 156.3, 155.7, 150.0, 136.9, 136.7, 132.2, 131.0, 131.0, 130.6, 130.0, 128.8, 79.2, 70.6, 70.6, 70.4, 70.2, 70.0, 54.5, 40.4, 39.5, 39.0, 28.5, 14.5, 13.2, 11.9. m/z HRMS found $[M+H]^+$ = 675.29120, $[C_{32}H_{43}ClN_6O_6S]^+$ requires 675.27226.

To a stirred solution of (+)-JQ1-PEG3-NHBoc (146 mg, 0.22 mmol) in $CH_2Cl_2$ (2 mL) at 0 °C was added TFA (3 mL) dropwise. The reaction mixture was warmed to room temperature overnight and the solvent removed *in vacuo*. The crude mixture was basified using saturated aqueous $NaHCO_3$, extracted with $CH_2Cl_2$, and the solvent removed *in vacuo* to afford (+)-JQ1-PEG3-$NH_2$ as a tan solid (125 mg, 99%), which was used immediately without further purification m/z HRMS found $[M+H]^+$ = 575.23657, $[C_{27}H_{36}ClN_6O_4S]^+$ requires 575.22018.

To a stirred solution of (+)-JQ1-PEG3-$NH_2$ (32 mg, 56 µmol), Ir-G2 (13) (61 mg, 56 µmol), and PyBOP (45 mg, 86 µmol) in anhydrous DMF (2 mL) under $N_2$ in the dark was added DIPEA (30 µL, 172 µmol). The resulting mixture was stirred overnight, diluted with EtOAc, and quenched by the addition of saturated aqueous $NaHCO_3$. The aqueous phase was removed and the organic layer washed with additional saturated aqueous $NaHCO_3$, 5% aqueous citric acid, brine, and dried over $Na_2SO_4$. The solvent was removed *in vacuo*, and the crude material purified by silica column chromatography (gradient elution: 0 to 3% MeOH/$CH_2Cl_2$) and C8 reverse phase preparative HPLC (gradient elution: 30 to 100% MeCN/$H_2O$ (0.1% formic acid)) to afford (+)-JQ1-PEG3-G2-Ir (1) as a yellow solid (25 mg, 27%).

$^1$H NMR (500 MHz, $CDCl_3$) δ: 9.24 – 8.92 (m, 2H), 8.58 – 8.27 (m, 2H), 8.24 (s, 1H), 8.04 (dd, *J* = 12.2, 8.9 Hz, 2H), 7.79 – 7.66 (m, 2H), 7.62 (s, 1H), 7.55 (s, 1H), 7.49 (t, *J* = 5.1 Hz, 1H), 7.41 (d, *J* = 8.2 Hz, 2H), 7.31 (d, *J* = 8.2 Hz, 2H), 6.63 (dd, *J* = 12.2 8.8 Hz, 2H), 5.62 (dd, *J* = 8.0, 2.3 Hz, 2H), 4.80 (br. s, 2H), 4.66 (t, *J* = 6.9 Hz, 1H), 3.70 – 3.30 (m, 18H), 3.24 – 3.15 (m, 2H), 2.93 – 2.77 (m, 2H), 2.66 (s, 3H), 2.63 (s, 3H), 2.39 (s, 3H), 1.66 (s, 3H). $^{13}$C NMR (125 MHz, $CDCl_3$) δ: 171.9, 170.8, 167.0 (dd, *J* = 258.2, 16.8 Hz), 163.9, 162.7 (dd, *J* = 262.6, 14.2 Hz), 157.6, 155.6 (d, *J* = 9.9 Hz), 155.1 (dd, *J* = 6.9, 28.6 Hz), 154.5, 149.6, 149.1, 145.2 (d, *J* = 3.2 Hz), 136.8 (d, *J* = 2.8 Hz), 136.6, 131.1, 130.9, 130.7, 130.0, 129.9, 129.6, 128.8, 127.9, 126.6, 126.4, 126.2, 123.7 (d, *J* = 22.5 Hz), 121.7 (dd, *J* = 273.3, 8.9 Hz), 114.2 (ddd, *J* = 17.1, 10.1, 2.6 Hz), 100.1

(dt, $J$ = 27.0, 9.7 Hz), 70.7, 70.4, 70.3, 69.9, 69.8, 54.5, 39.6, 39.2, 38.9, 35.2, 32.1, 29.8, 29.5, 22.8, 21.8, 14.6, 14.3, 14.3, 13.2, 12.0. $^{19}$F NMR (376 MHz, $CDCl_3$) δ: –62.7 (d, $J$ = 3.1 Hz), –62.7 (s), –72.1 (d, $J$ = 714.3 Hz), –101.6 (dtt, $J$ = 59.3, 12.5, 8.8 Hz), –105.9 – –106.1 (m). m/z HRMS found $[M]^+$ = 1507.34161 (100), 1508.33996 (84), 1505.33212 (63), 1506.3 3296 (52), 1509.33644 (72), 1510.33378 (47) $[C_{65}H_{57}ClF_{10}IrN_{10}O_5S]^+$ requires 1507.33876 (100), 1508.34202 (70), 1505.33634 (60), 1506.33969 (42), 1509.33572 (32), 1509.34538 (24), 1510.33907 (23). HPLC (Vydac 218TP C18 HPLC, gradient: 0 – 90% MeCN/$H_2O$ (0.1% TFA) 10 minutes, 5 minutes 90% MeCN (0.1% TFA), 1 mL/min, 254 nm): $\tau_r$ = 12.5 min.

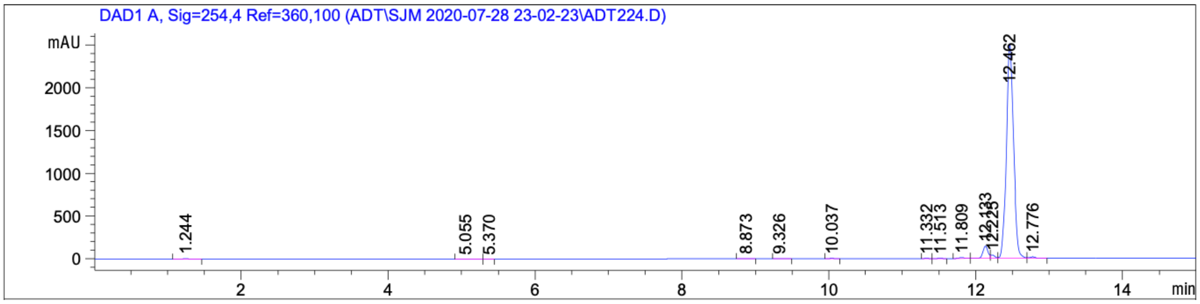

*The enantiomer was prepared analogously from (–)-JQ1-$CO_2$H and displayed identical spectral features.*

**(+)-JQ1-G1-catalyst (21)**

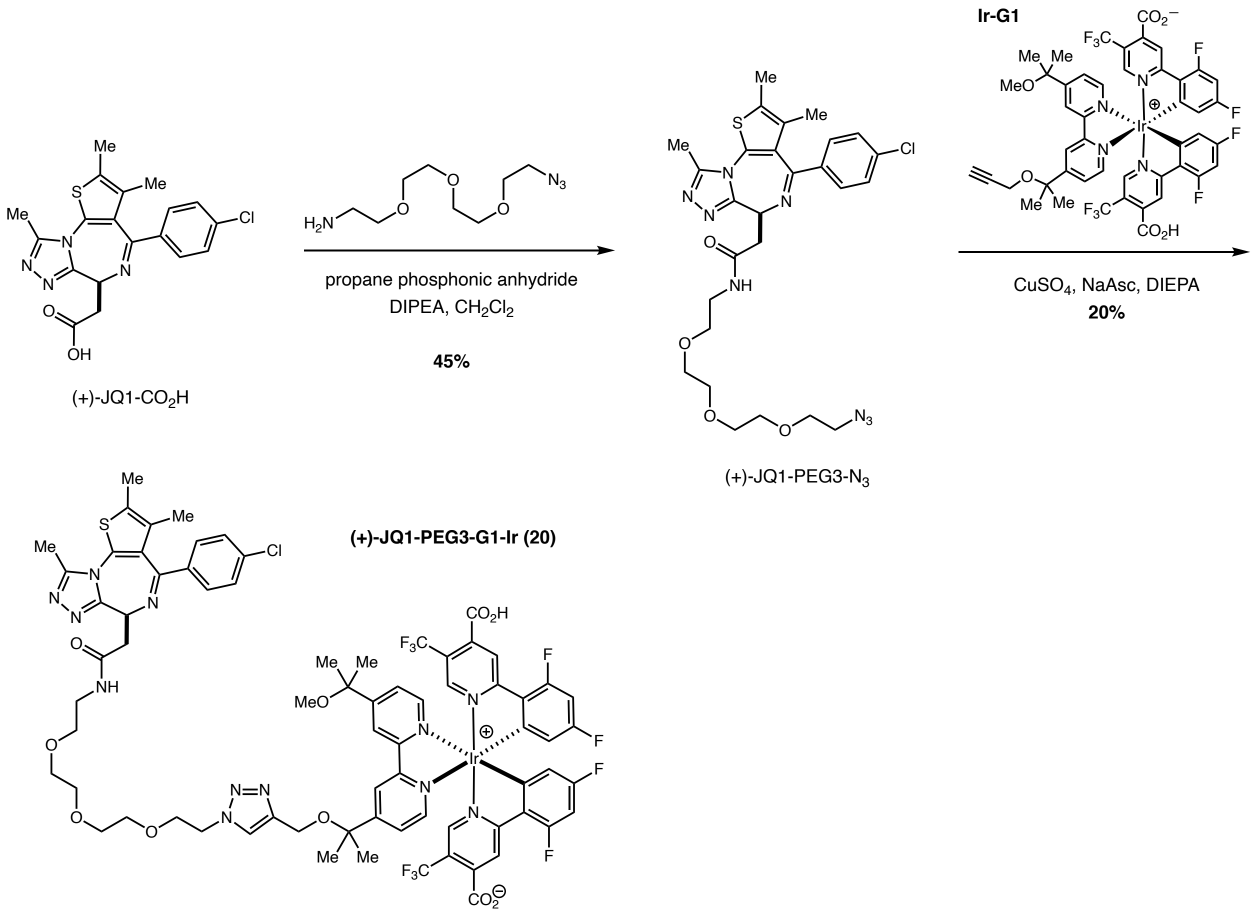

(+)-JQ1-$CO_2H$ (100 mg, 0.25 mmol), azido-PEG3-amine (60 mg, 0.27 mmol), 1-propanephosphonic anhydride (300 μL, 0.5 mmol, 50% solution in ethyl acetate) and disopropylethylamine (130 μL, 0.75 mmol) were combined in dichloromethane (0.6 mL) and stirred at room temperature for 3.5 hours. The reaction mixture was partitioned between ethyl acetate (15 mL) and water (15 mL). The aqueous layer was extracted with additional ethyl acetate and the organic layers were combined, washed with brine, dried over magnesium sulfate, filtered and then concentrated under reduced pressure. The resulting material was then purified by normal phase column chromatography (ISCO RediSep Gold 12 column, 0-100% (3:1 ethyl acetate:ethanol) in hexane to give (+)-JQ1-PEG3-azide as a colorless oil (68 mg, 45% yield). $^1$H NMR (500 MHz, $CDCl_3$) δ: 7.44 (d, 2H, *J* = 8.3 Hz), 7.36 (d, 2H, *J* = 8.4 Hz), 6.90 (bs, 1H), 4.68 (t, 1H, *J* = 7.0 Hz), 3.75 – 3.69 (m, 8H), 3.63 (m 2H), 3.55 (m, 2H), 3.45 – 3.37 (m, 2H), 2.69 (s,

3H), 2.43 (s, 3H), 1.70 (s, 3H). $^{13}$C NMR (125 MHz, $CDCl_3$) δ:170.6, 163.9, 155.7, 149.9, 136.8, 136.7, 132.2, 130.9, 130.8, 130.5, 129.9, 128.7, 70.7, 70.7, 70.7, 70.4, 70.0, 69.8, 54.4, 50.7, 39.4, 39.2, 14.4, 13.1, 11.8. m/z HRMS found $[M]^+$ = 601.2125, $[C_{27}H_{33}ClN_8O_4S]^+$ requires 601.2125.

(+)-JQ1-PEG3-azide (11 mg, 0.02 mmol) and Ir-G1 (21 mg, 0.02 mmol) and DIPEA (16 μL, 0.1 mmol) were combined in acetonitrile (0.2 mL) to give a hazy suspension. To this suspension was added a freshly prepared suspension of copper sulfate (1.4 mg, 0.005 mmol) and sodium ascorbate (3.3 mg, 0.02 mmol) in water (0.3 mL) which instantly resulted in a yellow solution. This reaction mixture was stirred at room temperature for 5 hours at which point it was diluted with 1.5 mL DMSO and purified by preparative HPLC (50-100% MeCN/water, 0.05% TFA over 10 minutes, 20 mL/min, LUNA 5 micron C18(2) 100 angstrom, 250 x 21.2 mm). The product fraction was lyophilized. Preparative HPLC (same conditions) was repeated and the product fraction was lyophilized to give JQ-1-PEG3-Ir (6 mg, 20% yield) as a yellow solid. $^1$H NMR (500 MHz, MeOH-*d*4) δ: 9.07 (s, 1H), 8.92 (s, 1H), 8.70 (s, 2H), 8.14 – 8.08 (m, 2H), 8.06 (s, 1H), 7.86 – 7.80 (m, 2H), 7.66 (d, *J* = 10.4 Hz, 2H), 7.50 – 7.43 (m, 2H), 7.40 (dd, *J* = 8.7, 3.9 Hz, 2H), 6.92 – 6.79 (m, 2H), 5.94 – 5.85 (m, 2H), 4.69 – 4.61 (m, 1H), 4.57 (q, *J* = 4.5 Hz, 2H), 4.53 – 4.43 (m, 2H), 3.89 (t, *J* = 4.8 Hz, 2H), 3.68 – 3.56 (m, 10H), 3.50 – 3.39 (m, 3H), 3.28 (dd, *J* = 14.9, 5.2 Hz, 1H), 3.24 (d, *J* = 2.5 Hz, 3H), 2.69 (d, *J* = 3.2 Hz, 3H), 2.46 (s, 3H), 1.69 (dd, *J* = 17.2, 3.9 Hz, 15H). $^{13}$C NMR (125 MHz, MeOH-*d*4) δ: 171.32, 168.40, 166.33, 164.93, 164.59, 164.17, 162.20, 161.83, 161.67, 159.62, 159.51, 159.32, 156.29, 156.16, 155.51, 155.25, 151.03, 150.77, 149.66, 146.48, 144.21, 142.69, 136.67, 136.51, 132.09, 130.71, 130.57, 130.03, 128.41, 126.38, 126.13, 124.42, 123.22, 123.05, 122.80, 122.61, 122.54, 120.37, 113.94, 99.64, 99.42, 99.21, 77.45, 76.60, 70.12, 70.10, 69.94, 69.17, 68.99, 56.80, 53.63, 49.97, 49.92, 39.15, 37.18, 26.78, 26.74, 26.42, 26.38, 25.81, 13.00, 11.53, 10.17. $^{19}$F NMR (471 MHz, MeOH-*d*4) δ: -61.73, -77.07, -103.74, -107.98. m/z calcd. for $C_{73}H_{66}ClF_{10}IrN_{12}O_{10}S$ (1719.3958 found 1719.3947 (M+H) and 860.2029 (M+2H)/2. LC retention time: 1.23 minutes using Acquity Single pole LCMS equipped with two channels (20 and 25 V). The flow rate is 0.6 ml/min on a 2.1 x 50 mm BEH 1.7 μM particle size column with gradient 5 to 100% MeCN for 1.8 min, hold for 0.2 min.

**(+)-JQ1-Dz-alkyne (2)**

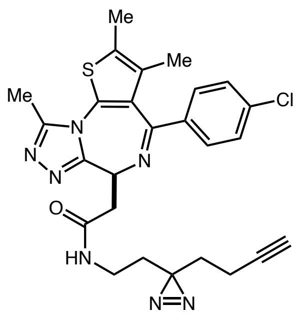

*Prepared according to reported literature procedure*[6].

(+)-JQ1-$CO_2H$ (70 mg, 0.17 mmol) was dissolved in anhydrous DMF (2 mL) followed by the addition of HOBt (32 mg, 0.24 mmol), EDCI (46 mg, 0.30 mmol), $Et_3N$ (71 $\mu$L, 0.50 mmol), and 2-(3-but-3-yn-1-yl)-3H-diazirin-3-yl)ethan-1-amine (11) (30 mg, 0.22 mmol). The reaction mixture was stirred overnight under $N_2$. The reaction mixture was diluted with EtOAc and the organic layer washed with saturated aqueous $NaHCO_3$, 0.1 M HCl, brine, and dried over $Na_2SO_4$. The solvent was removed *in vacuo*, and the crude material purified by silica column chromatography (gradient elution: 0 to 5% MeOH/$CH_2Cl_2$) to afford (+)-JQ1-Dz-alkyne (2) as a colourless film (54 mg, 59%). NMR (500 MHz, $CDCl_3$) δ: 7.39 (d, $J$ = 8.5 Hz, 2H), 7.31 (d, $J$ = 8.5 Hz, 2 H), 7.07 (t, $J$ = 5.3 Hz, 1H), 4.62 (t, $J$ = 6.9 Hz, 1 H), 3.58 (dd, $J$ = 14.4, 7.6 Hz, 1H), 3.42 – 3.33 (m, 1H), 3.15 (quint, $J$ = 5.9 Hz, 2H), 2.65 (s, 3H), 2.39 (s, 3H), 2.00 – 1.94 (m, 3H), 1.67 – 1.58 (m, 7H). $^{13}C$ NMR (125 MHz, $CDCl_3$) δ: 170.6, 164.0, 155.9, 150.0, 136.9, 136.6, 132.2, 131.0, 130.9, 130.6, 129.9, 128.8, 82.8, 69.4, 54.5, 53.5, 39.3, 34.5, 32.7, 32.1, 29.8, 26.9, 14.5, 13.3, 13.2, 11.9. m/z HRMS found $[M+H]^+$ = 520.17939, $[C_{26}H_{27}ClN_7OS]^+$ requires 520.16808.

*The enantiomer was prepared analogously from (–)-JQ1-$CO_2H$ and displayed identical spectral features.*

#### Expression of recombinant $(His)_6$-tagged BRD4

Histidine (6x)-tagged pNIC28-Bsa4 bromodomain expression constructs were obtained from Addgene (#38943). Single colonies were grown overnight at 37 ºC on LB/agar plates with 50 µg/ml kanamycin, subsequently harvested, and cultured in LB media with 50 µg/ml kanamycin (20 mL) for 16 hours. Plasmid DNA was extracted using QIAprep Spin Miniprep Kit (Qiagen, #27104) and stored at –20 ºC until use. The plasmid DNA (100 ng) was transformed into competent *E. coli* BL21(DE3) cells (NEB) (50 µL) at 0 ºC for 30 minutes. The mixture was subsequently heat-shocked at 42 ºC for 30 minutes and cooled at 0 ºC for an additional 5 minutes. The mixture was diluted with SOC outgrowth media (NEB) (950 µL), incubated at 37 ºC for 1 hour on a rotisserie, and diluted with Terrific broth containing 50 µg/ml kanamycin (9 mL). The mixture was incubated overnight at 37 ºC. Subsequent expression cultures were prepared by diluting start-up cultures 1:200 in fresh medium. Growth was allowed at 37 ºC to an optical density (OD) of about 0.7 ($OD_{600}$) before the temperature was decreased to 0 ºC. Protein expression was induced overnight at 20 ºC with 0.1 mM isopropyl-β-D-thiogalactopyranoside (IPTG). The cells were subsequently pelleted (4500xg, 45 minutes) and resuspended in lysis buffer (3 mL per g cells, 50 mM HEPES, 500 mM NaCl, 30 mM imidazole, pH = 7.5 containing lysozyme (1 mg/mL), DNAase (0.1 mg/mL), and PMSF (1 mM). The cells were lysed using a fluidizer and the lysate cleared by centrifugation (20,000xg, 45 min, 4 ºC). The supernatant was then incubated with Nickel-NTA agarose beads (Qiagen) (4 mL, 30 min, 4 ºC) and the beads washed with 5 x volume wash buffer (50 mM HEPES, 500 mM NaCl, 30 mM imidazole, pH = 7.5). The protein was eluted using elution buffer (50 mM HEPES, 500 mM NaCl, pH = 7.5) containing a gradient of imidazole (30 to 250 mM imidazole; 25 mL). The fractions were analyzed by SDS-PAGE (>90% purity) and fractions combined and dialyzed overnight (20 mM HEPES, 150 mM NaCl, pH = 7.5) at 4 ºC. The sample was subsequently purified by FPLC (Amersham Bioscience) using a Hiload 16/60 superdex 75 prep grade column, and aliquoted into 0.5 mg/mL samples and stored at –80 ºC.

#### Labelling of recombinant BRD4 using JQ1-G2

*Representative example:*

To a 0.5 mL Eppendorf tube was added DPBS (11.5 µL), $(His)_6$-BRD4 (2.5 µL, 9.7 µM, 20 mM HEPES, 150 mM NaCl), and bovine carbonic anhydrase (2.5 µL, 5.9 µM, DPBS) followed by either (+)-JQ1-G2 (1) (1.25 µL, 20 µM, 10% DMSO in DPBS), (–)-JQ1-G2 (1.25 µL, 20 µM, 10% DMSO in DPBS), or Ir-NHBoc (15) (1.25 µL, 20 µM, 10% DMSO in DPBS) or (+)-JQ1 (10x off-compete). Dz-PEG3-biotin (9) (1.25 µL, 100 µM, DPBS) was added and the sample was irradiated 15 minutes at 450 nM in the Efficiency Aggregator PhotoReactor. The sample was subsequently diluted with 4x Laemlli buffer with BME (5 µL) and heated to 95 °C for 10 minutes. The samples were cooled to room temperature and centrifuged. The samples were subsequently loaded onto a BioRad Criterion 4–20% tris-glycine gel, alongside all of the appropriate controls, and run in freshly prepared Tris running buffer (160V, 60 minutes). The gel was washed (3 x MiliQ water) and transferred via iBlot 2 to an NC membrane. Following transfer, the membranes were then immersed in REVERT total protein stain (Li-Cor, 926-11011) for 5 minutes. Excess stain was decanted, membranes washed with 6.7:30:63.3 AcOH:MeOH:$H_2O$ and imaged using a Li-Cor Odyssey CLx scanner in the 700 nm channel. The membranes were washed with water, then immersed in Odyssey Blocking Buffer (Li-Cor, 927-50000) and incubated for 1 hour. The blocking solution was then decanted, and 35 mL of fresh blocking buffer containing 70 µL of Tween 20 was added. This mixture was rocked for 5 minutes. Afterwards, 1.5 µL of IRDye 800CW streptavidin (Li-Cor, 926-32230) was added and the mixture incubated for 1 hour. The blocking

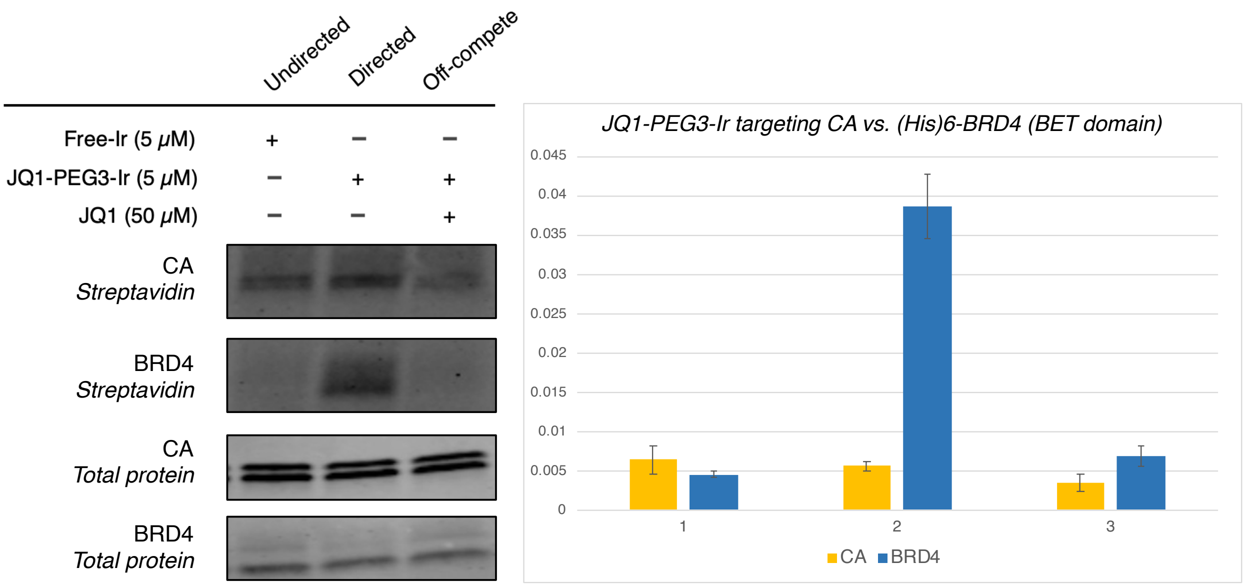

buffer was then decanted, and the membranes were washed with 1X TBST (3 x 5 min) and water before imaging via Li-Cor Odyssey CLx scanner in the 800 nm channel. Pixel densitometry was performed using Image Studio Lite V. 5.2 (Li-Cor). The streptavidin 800 channel pixel density was then divided by the total protein stain 700 channel pixel density to provide a normalized biotinylation signal for each protein band. Reactions were repeated (n=3).

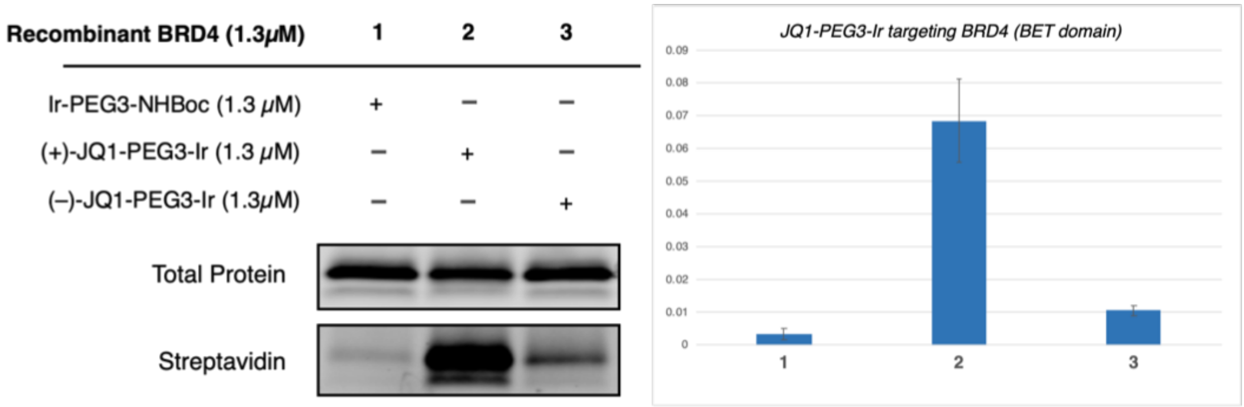

***MST study on JQ1 binding to $(His)_6$-BRD4***

*Preparation of $(His)_6$-tagged BRD4 Alexa Fluor 633 conjugate*

To a solution of $(His)_6$-tagged BRD4 (200 µL, 24 µM, 20 mM HEPES, 150 mM NaCl, pH = 7.5) was added $NaHCO_3$ (20 µL, 1 M) and Alexa Fluor 633 NHS ester (6 µL, 5 mM, DMSO). The mixture was incubated for 90 minutes at room temperature on a rotisserie in the dark and concentrated (7K MWCO Zebra Spin desalting column). The resulting solutions were analysed by NanoDrop and protein concentration determined to be between 6.4–8.9 µM with and average of 2.75–4.6 dyes per protein. The protein-dye conjugate was used immediately.

*Binding assay of $(His)_6$-tagged BRD4-AF633 conjugate vs. JQ1 and JQ1-PEG3-Ir*

A solution of (+)-JQ1 (20 mM, DMSO) /(+)-JQ1-PEG3-Ir were diluted (20 mM HEPES, 150 mM NaCl, pH = 7.5) to a final concentration of 400 mM (200 µL). A 16-fold serial dilution (10 µL) of the respective molecules was prepared and then diluted with $(His)_6$-tagged BRD4-AF633 (40 nM, 10 µL). MST was subsequently performed on a Monolith NT.115 instrument with three replicates.

(+)-JQ1 showed a binding $K_D$ = 6 µM and (+)-JQ1-PEG3-G2-Ir (1) showed a binding $K_D$ = 4 µM. The corresponding Ir-PEG3-NHBoc showed >85 µM binding. The lower binding of (+)-JQ1 compared to reported values is due to the use of a truncated form of the recombinant protein.

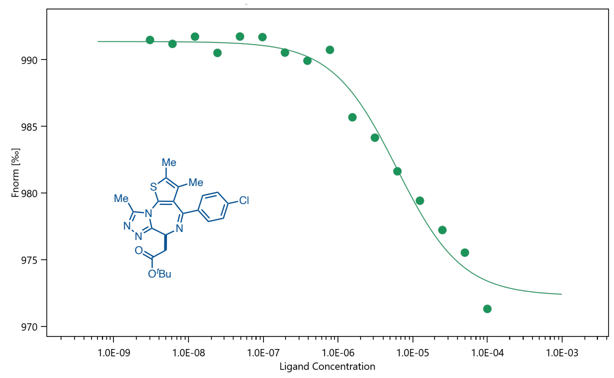

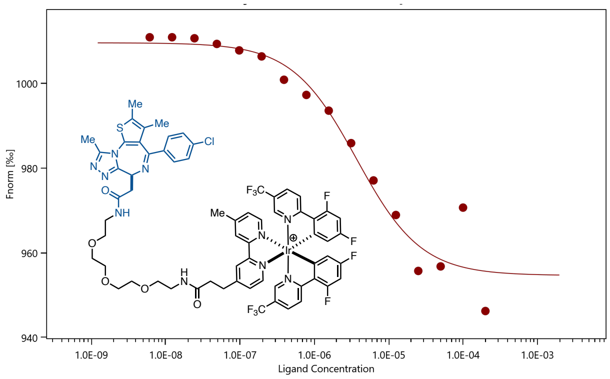

#### Intracellular labelling of BRD4 using (+)-JQ1-iridium conjugates

*Incubation and Irradiation:*

To HeLa cells in 12 x 10 cm plates at 80% confluency in DMEM with **no** phenol red (Gibco) (4 mL) was added (+)-JQ1-PEG3-G2 (1) (5 µM) (4 plates, **A**); Ir-PEG3-NHBoc (15) (5 µM) (4 plates, **B**); and DMSO (4 plates, **C**). Final DMSO concentration was always kept below 0.5%. The plates were incubated at 37 °C for 3 hours and the media removed and replaced. Diazirine-PEG3-bioitin (9) was added (250 µM) and the plates incubated at 37 °C for an additional 20 minutes. The plates were subsequently irradiated (without the lid) in the bioreactor at 450nM for 15 minutes. The media was removed and the cells washed twice with cold DPBS (4 °C). The cells were resuspended in cold DPBS (4 °C), scraped and transferred to a separate 50 mL falcon tube. The cells were pelleted (1000g for 5 minutes at 4 °C) and suspended in 1mL of cold RIPA buffer containing PMSF (1mM) and cOmplete EDTA free protease inhibitor (1x) (Roche). The lysed cells were incubated on ice for 5–10 minutes and sonicated (35%, 5 x 5s with 30s rest). The lysate was then centrifuged at 15x1000g for 15 mins at 4 °C and the supernatant collected. The concentration of the cell lysate was measured by BCA assay and adjusted accordingly to equal concentration of 1 mg/mL. A control sample was removed from each plex (15 µL) and stored at –20 °C for later analysis.

*Streptavidin pull-down:*

Magnetic Streptavidin beads (NEB) were removed (250 µL per plex) and washed twice with RIPA (0.5 mL) (5 minutes incubation on a rotisserie). The beads were pelleted on a magnetic rack, diluted with the samples (1 mL) and incubated on a rotisserie at 4 °C overnight. The beads were pelleted on a magnetic rack, the supernatant removed, and a control sample from each plex (15 µL) and stored at –20 °C for later analysis. The beads were subsequently washed with 1 x RIPA (0.5 mL), 3 x 1% SDS in DPBS (0.5 mL), 3 x 1M NaCl in DPBS (0.5 mL), 3 x 10% EtOH in DPBS and 1 x RIPA (0.5 mL). The samples were incubated with each wash for 5 minutes prior to pelleting. The beads were resuspended in RIPA buffer (300 µL) and transferred to a new 1.5 mL Lo-bind tube.

*Western Blot analysis:*

Following the final wash and transfer procedure for pull-down, the beads were pelleted on a magnetic rack and the supernatant removed. The beads were gently centrifuged to gather at the bottom of the tube and freshly prepared elution buffer (30 mM biotin, 6 M urea, 2 M thiourea, 2% SDS in DPBS, pH = 11.5) (24 µL) and 4x Laemlli buffer with BME (6 µL) was added with gentle mixing. The beads were heated to 95 °C for 15 minutes, pelleted on a magnetic rack, and the supernatant was removed while hot and beads discarded. The samples were cooled to room temperature and centrifuged. The samples (17 µL) were subsequently loaded onto a BioRad Criterion 4–20% tris-glycine gel, alongside all of the appropriate controls, and run in freshly prepared Tris running buffer (160V, 60 minutes). The gel was washed (3 x MiliQ water) and transferred via iBlot 2 to an NC membrane. The membrane was again washed (3 x MiliQ water) and blocked with Li-COR TBS Blocking Buffer for 1 hour at room temperature and then incubated with anti-BRD4 (A-7, Santa Cruz) (1:500) and anti-histone H3 (polyclonal Invitrogen PA5-16183) (1:2000) overnight in Pierce Protein-Free Blocking (1:2000) at 4 °C overnight. The membrane was washed 3 x TBST (5 mins per wash) and 5 x MiliQ water and resuspended in Pierce Protein-Free Blocking Buffer with Li-COR secondary antibodies (Goat-anti-Mouse 800) and (Goat-anti-Rabbit 700) and rocked for 1 hour at room temperature (1:12,500). The membrane was washed 3 x TBST (5 mins per wash) and 5 x MiliQ water and imaged.

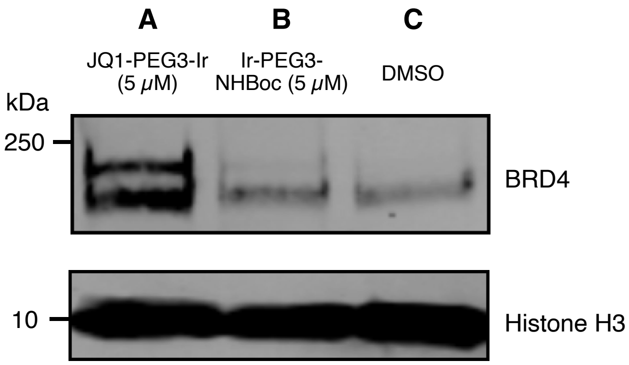

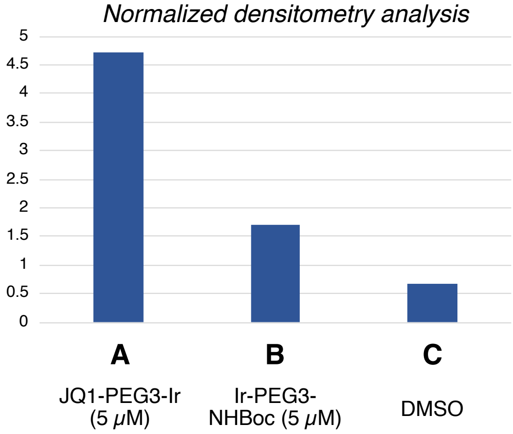

***Time dependent labelling of BRD4 using (+)-JQ1-PEG3-Ir:***

Following the in-cell labelling protocol as described above. Irradiation time was varied so as to demonstrate the degree of biotinylation over time (2, 5, and 15 mintues). The control reaction using UV light was performed using a UV-photobox wherein the plates were irradiated using 254nm light at 4 °C for 20 minutes.

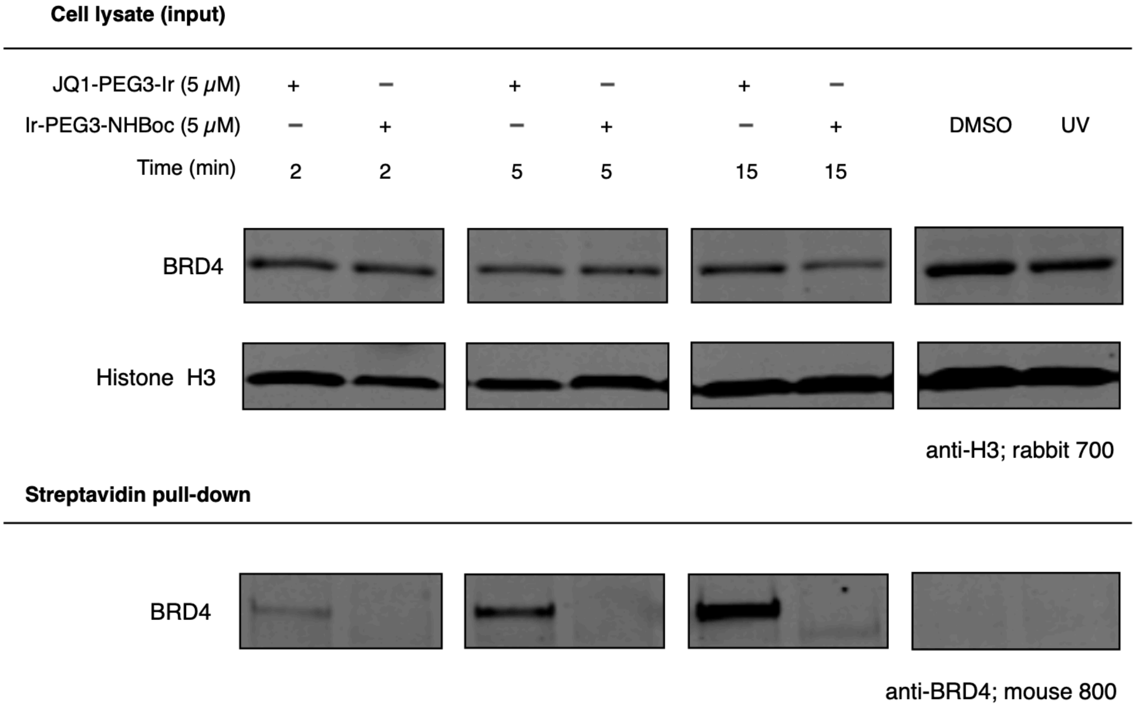

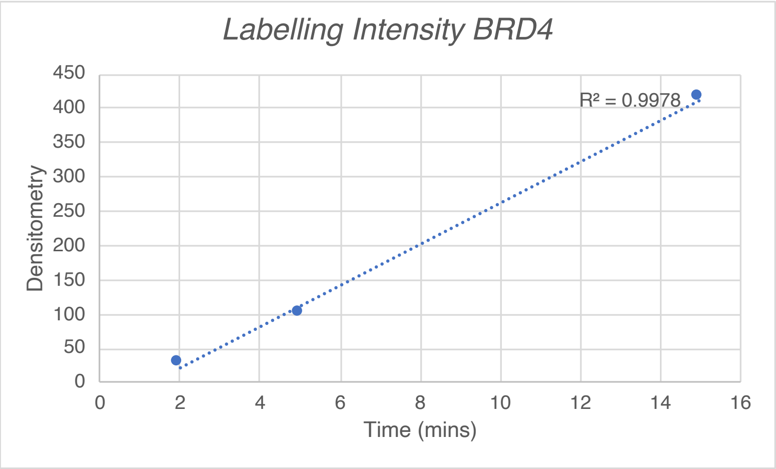

*Comparing labelling between Gen-1 and Gen-2 JQ1 iridium catalyst conjugates:*

Following the in-cell labelling protocol as described above. To HeLa cells in 12 x 10 cm plates at 80% confluency in DMEM with **no** phenol red (Gibco) (4 mL) was added JQ1-PEG3-G2-Ir (1) (5 µM) (4 plates, **A**); JQ1-PEG3-G1-Ir (21) (5 µM) (4 plates, **B**); and DMSO (4 plates, **C**). The plates were incubated at 37 °C for 3 hours and the media removed and replaced. Diazirine-PEG3-bioitin (9) was added (250 µM) and the plates incubated at 37 °C for an additional 20 minutes. The plates were subsequently irradiated (without the lid) in the bioreactor at 450nM for 20 minutes. Streptavidin enrichment and western blot performed as previously described.

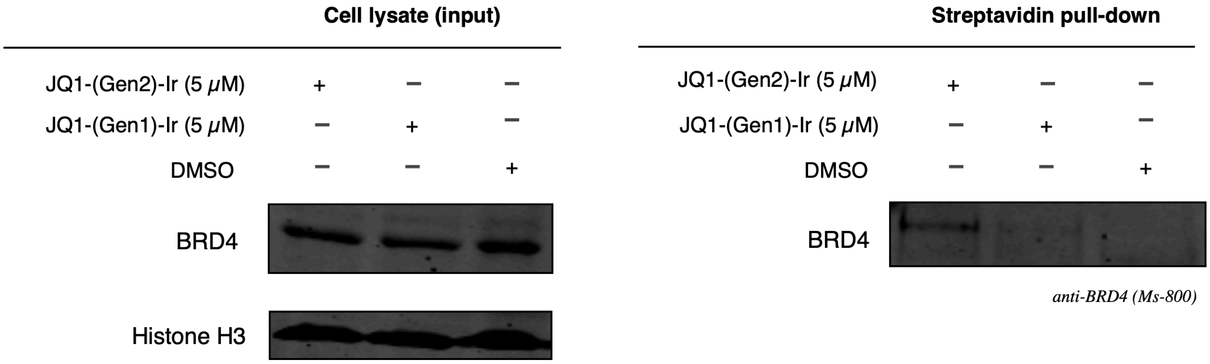

*Comparing labelling between (+)-JQ1 and (–)-JQ1 iridium catalyst conjugates (Gen-2):*

(–)-JQ1 has NO affinity for BRD-proteins and hence serves as a negative control.

Following the in-cell labelling protocol as described above. To HeLa cells in 12 x 10 cm plates at 80% confluency in DMEM with **no** phenol red (Gibco) (4 mL) was added (+)-JQ1-PEG3-G2-Ir (1) (5 µM) (4 plates, **A**); (–)-JQ1-PEG3-G2-Ir (5 µM) (4 plates, **B**); and DMSO (4 plates, **C**). The plates were incubated at 37 ºC for 3 hours and the media removed and replaced. Diazirine-PEG3-bioitin (9) was added (250 µM) and the plates incubated at 37 ºC for an additional 20 minutes. The plates were subsequently irradiated (without the lid) in the bioreactor at 450nM for 20 minutes. Streptavidin enrichment and western blot performed as previously described.

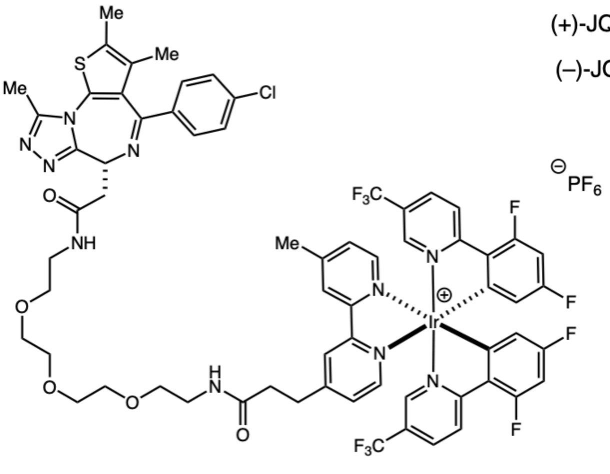

(–)-JQ1-PEG3-iridium

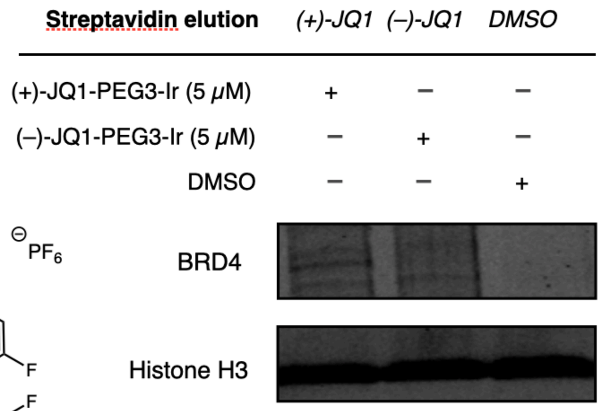

#### μMap proteomics using (+)-JQ1-G2

The procedure carried out is identical to that of the *in-cell labelling for western blot analysis*: To HeLa cells in 12 x 10 cm plates at 80% confluency in DMEM with **no** phenol red (Gibco) (4 mL) was added (+)-JQ1-PEG3-G2-Ir (1) (5 μM) (6 plates, **A**) and Ir-PEG3-NHBoc (15) (5 μM) (6 plates, **B**). The plates were incubated at 37 °C for 3 hours and the media removed and replaced. Diazirine-PEG3-bioitin (9) was added (250 μM) and the plates incubated at 37 °C for an additional 20 minutes. The plates were subsequently irradiated (without the lid) in the bioreactor at 450nM for 15 minutes. The media was removed and the cells washed twice with cold DPBS (4 °C). The cells were resuspended in cold DPBS (4 °C), scraped and transferred into separate 15 mL falcon tube (2 plates per tube; 6 tubes in total). The cells were pelleted (1000g for 5 minutes at 4 °C) and suspended in 2 mL of cold RIPA buffer containing PMSF (1mM) and cOmplete EDTA free protease inhibitor (1x) (Roche). The lysed cells were incubated on ice for 5–10 minutes and sonicated (35%, 5 x 5s with 30s rest). The lysate was then centrifuged at 15x1000g for 15 mins at 4 °C and the supernatant collected. The concentration of the cell lysate was measured by BCA assay and adjusted accordingly to a concentration of 1.5 mg/mL. Magnetic Streptavidin beads (NEB) were removed (350 μL per plex) and washed twice with RIPA (0.5 mL) (5 minutes incubation on a rotisserie). The beads were pelleted on a magnetic rack, diluted with the samples (1 mL) and incubated on a rotisserie at 4 °C overnight. The beads were pelleted on a magnetic rack, the supernatant removed, and a control sample from each plex (15 μL) and stored at –20 °C for later analysis. The beads were subsequently washed with 1 x RIPA (0.5 mL), 3 x 1% SDS in DPBS (0.5 mL), 3 x 1M NaCl in DPBS (0.5 mL), 3 x 10% EtOH in DPBS and 1 x RIPA (0.5 mL). The samples were incubated with each wash for 5 minutes prior to pelleting. The beads were resuspended in RIPA buffer (300 μL) and transferred to a new 1.5 mL Lo-bind tube.

The supernatant was removed and the beads washed with 3 x DPBS (0.5 mL) and 3 x $NH_4HCO_3$ (100 mM) (0.5 mL). The beads were re-suspended in 500 μL 6 M urea in DPBS and 25 μL of 200 mM DTT in 25 mM $NH_4HCO_3$ was added. The beads were incubated at 55 °C for 30 min. Subsequently, 30 μL 500 mM IAA in 25 mM $NH_4HCO_3$ was added and incubated for 30 min at room temperature in the dark. The supernatant was removed and the beads washed with 3 x 0.5 mL DPBS and 3 x 0.5 mL TEAB (50 mM). The beads were resuspended in 0.5 mL TEAB

(50 mM) and transferred to a new protein LoBind tube, pelleted, and the supernatant removed. The beads were resuspended in 40 µL TEAB (50 mM) and 1.2 µL trypsin (1 mg/mL in 50 mM acetic acid) was added and the beads incubated overnight on a rotisserie at 37 °C. After 16 hours, an additional 0.8 µL trypsin was added and the beads incubated for an additional 1 hour on a rotisserie at 37 °C. Meanwhile, the TMT6 plex label reagents (0.8 mg) (Thermo) were equilibrated to room temperature and diluted with 41µL of anhydrous acetonitrile (Optima grade; 5 min with vortexing) and centrifuged. The beads were subsequently pelleted and the supernatant transferred to the corresponding TMT-label. The reaction was incubated for 2 hours at room temperature. The samples were quenched with 8µL of 5% hydroxylamine and incubated for 15 minutes. All of the samples were pooled in a new Protein LoBind tune and quenched with TFA (16µL, Optima). The samples were stored at –80 °C until proteomics were conducted. Samples were desalted and fractionated prior to running.

*LC-MS/MS/MS-based proteomic analysis*

Mass spectra were obtained using an Orbitrap Fusion at Princeton Proteomics Facility and analysed using MaxQuant. TMT labeled peptides were dried down in SpeedVac, re-dissolved in 300 µl of 0.1% TFA in water and fractionated into 8 fractions using Pierce™ High pH Reversed-Phase Peptide Fractionation Kit (#84868). Fractions 1, 4, and 7 were combined as sample 1. Fractions 2 and 6 were combined as sample 2. Fractions 3, 5, and 8 were combined as sample 3. Three combined samples were dried completely in a SpeedVac and resuspended in 20 µl 5% acetonitrile/water (0.1% formic acid (pH = 3)). 2µl (~ 360ng) was injected per run using an Easy-nLC 1200 UPLC system. Samples were loaded directly onto a 45cm long 75um inner diameter nano capillary column packed with 1.9um C18-AQ resin (Dr. Maisch, Germany) mated to metal emitter in-line with an Orbitrap Fusion Lumos (Thermo Scientific, USA). Column temperature was set at 45 °C and two-hour gradient method with 300nl per minute flow. The mass spectrometer was operated in data dependent mode with synchronous precursor selection (SPS) - MS3 method[7] with 120,000 resolution of MS1 scan (positive mode, profile data type, Intensity threshold 5.0e3 and mass range of 375-1600 m/z) in the Orbitrap followed by CID fragmentation in ion trap with 35% collision energy for MS2 and HCD fragmentation in Orbitrap (50,000 resolution) with 55% collision energy for MS3. MS3 scan range was set at 100-500 with injection time of 120ms.

Dynamic exclusion list was invoked to exclude previously sequenced peptides for 60s and maximum cycle time of 2.5s was used. Peptides were isolated for fragmentation using quadrupole (0.7 m/z isolation window). Ion-trap was operated in Rapid mode.

MS/MS/MS data was searched against 2018 Uniprot human protein database containing common contaminants (forward and reverse). Samples were set to three fractions and database search criteria were applied as follows: variable modifications set to methionine oxidation and *N*-terminal acetylation and deamidation (NQ), and fixed modifications set to cysteine carbamidomethylation, with a maximum of 5 modifications per peptide. Specific tryptic digestion (trypsin/P) with a maximum of 2 missed cleavages. Peptide samples were matched between runs. The maximum peptide mass was set to 6000 Da. The label minimum ration count was set to 2 and quantified using both unique and razor peptides. FTMS MS/MS match tolerance was set to 0.05 Da, and ITMS MS/MS match tolerance was set to 0.6 Da. All other settings were left as default.

The proteinGroups.txt file was subsequently imported into Persues [Main: corrected reported intensities; the remaining entries left to default]. The rows were subsequently filtered by categorical column with '+' values with matching rows removed via a reduced matrix based upon the following criteria, 'only identified by site', 'reverse', and 'potential contaminant'. The resulting matrix was then transformed by log2($x$) and the column correlation verified to be >0.9. From the previous matrix, the rows were annotated (categorical annotation of rows) into their corresponding experiments (3 x **A**, 3 x **B**). The matrix was subsequently normalized (subtraction of columns), and the corresponding data plotted as a scatter graph (volcano plot). The FDR was determined by a 2-sample T-test (Benjamini-Hochberg).

*Due to similarities in sequence homology between BRD2, BRD3, and BRD4 – it is ambiguous as to the precise identities of the enriched bromodomain proteins.*

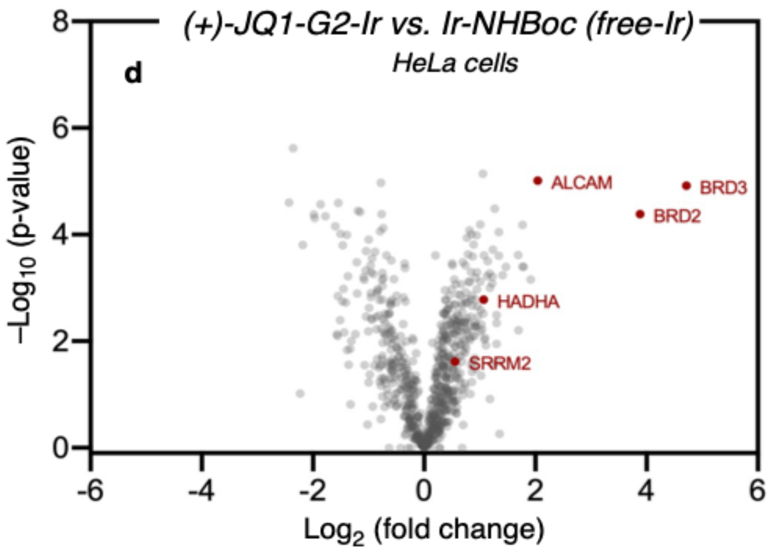

*Proteomics preparation and isobaric labelling (+)-JQ1-G2 vs Free Ir + (+)-JQ1:*

Reaction was conducted identically to previous experiment except in the samples containing free-iridium, an equimolar amount of (+)-JQ1 was added (5 µM).

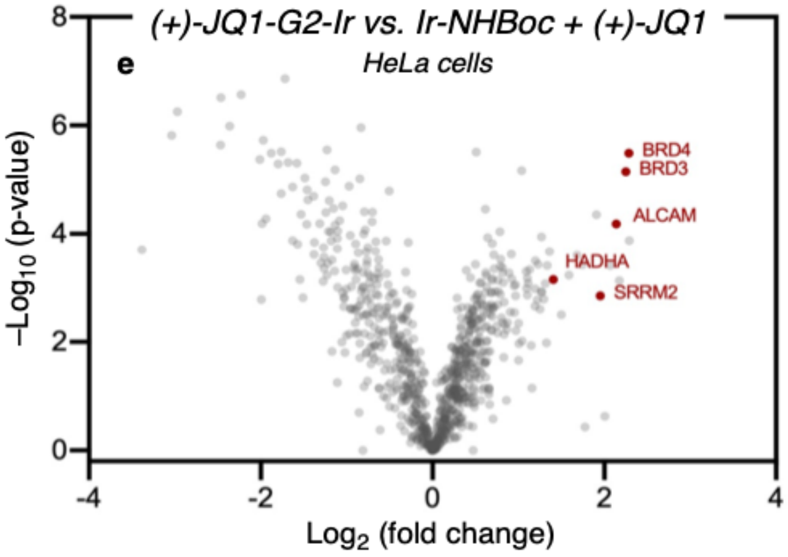

*Proteomics preparation and isobaric labelling (+)-JQ1-G2 vs (–)-JQ1-G2:*

Reaction was conducted identically to previous experiment except control lane contains *vs (–)-JQ1-G2* (5 µM).

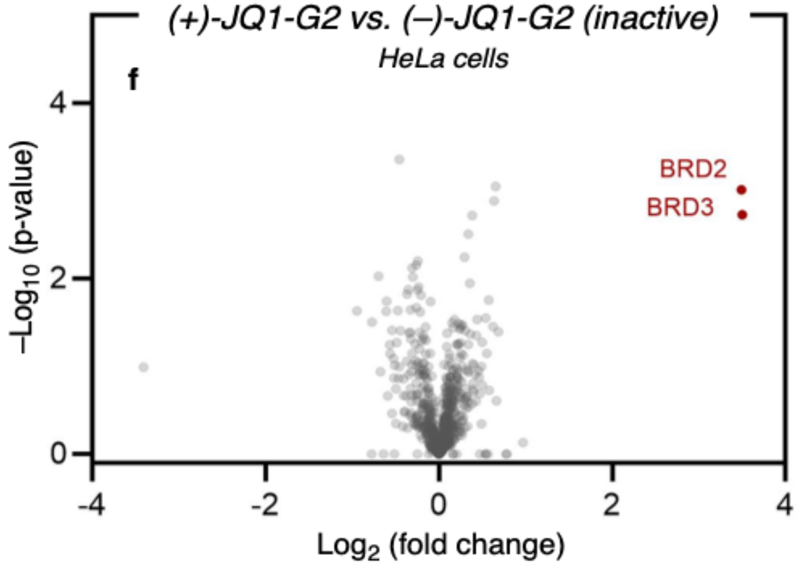

#### Confocal microscopy

HeLa cells were plated onto 35 mm glass-bottom microscopy dishes with DMEM (no phenol red) and treated with (+)-JQ1-PEG3-G2 (1) (5 μM), Ir-PEG3-NHBoc (15) (5 μM), and DMSO. The plates were incubated at 37 °C for 3 hours and the media removed and replaced. Diazirine-PEG3-bioitin (9) was added (250 μM) and the plates incubated at 37 °C for an additional 20 minutes. The plates were subsequently irradiated (without the lid) in the bioreactor at 450nM for different time periods. The media was removed, and the cells were washed with PBS. Cells were then fixed with 400 μL of 4% paraformaldehyde in PBS for 20 min at 37 C. Cells were washed 3x with PBS and permeabilized with 400 μL of 0.1% triton X-100 in PBS for 20 min at RT. Cells were washed with PBS and blocked with 400 μL of 2% BSA in PBS for 20 min at RT. Cells were washed 3x with PBS and incubated with 400 μL of Streptavidin-Alexa Fluor 488 diluted 1:500 and Hoechst diluted 1:10,000 in PBS. Confocal microscopy was performed at 40x magnification using a Nikon A1/HD25 microscope (Nikon Instruments, Inc., Melville, NY). Images shown are representative of the multiple cross-sectional images taken during each session.

*Irradiation time = 2 mins*

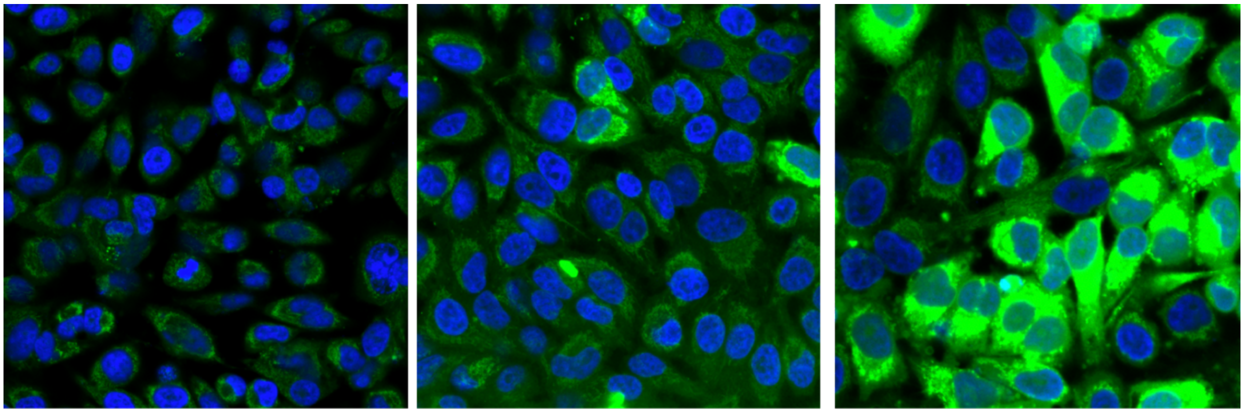

DMSO control | JQ1-PEG3-Ir *(directed)* | Ir-PEG3-NHBoc *(free-Ir)*

*Irradiation time = 10 mins*

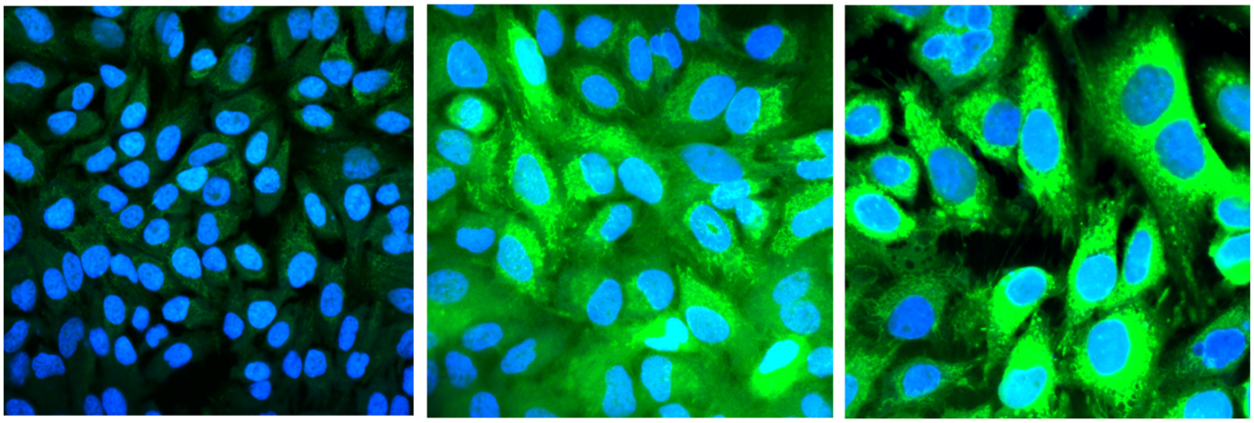

DMSO control | JQ1-PEG3-Ir *(directed)* | Ir-PEG3-NHBoc *(free-Ir)*

#### PAL using JQ1-Dz-alkyne

To HeLa cells in 8 x 10 cm plates at 80% confluency in DMEM with **<u>no</u>** phenol red (Gibco) (4 mL) was added (+)-JQ1-Dz-alkyne (2) (5 µM) (4 plates, **A**), and (–)-JQ1-Dz-alkyne (5 µM) (4 plates, **B**). The plates were incubated at 37 °C for 3 hours and the media removed and replaced. The plates were subsequently irradiated at 4 °C for 20 minutes in a UV reactor (254 nm). The media was removed and the cells washed twice with cold DPBS (4 °C). The cells were resuspended in cold DPBS (4 °C), scraped and transferred into separate 15 mL falcon tube (2 plates per tube; 6

tubes in total). The cells were pelleted (1000g for 5 minutes at 4 °C) and suspended in 1 mL cold lysis buffer (20 mM HEPES, 10 mM KCl, 1.5 mM $MgCl_2$, 0.5% NP40) containing PMSF (1mM) and cOmplete EDTA free protease inhibitor (1x) (Roche). The lysed cells were incubated on ice for 5–10 minutes and sonicated (35%, 5 x 5s with 30s rest). The lysate was then centrifuged at 15x1000g for 20 mins at 4 °C and the supernatant collected. The concentration of the cell lysate was measured by BCA assay and adjusted accordingly to a concentration of 1.5 mg/mL.

*CuAAC reaction:*

*Click-cocktail stock solution:* In a 0.5mL Lo-bind tube, 6.2 µL 500mM $CuSO_4$ was added to 62 µL 100 mM THPTA and vortexed. Subsequently, 15.5 µL 5 mM biotin-PEG7-azide was added, followed by 15.5 µL of freshly prepared 1M sodium ascorbate (*Important: addition of reagents in that order*).

To the cell-lysate (1 mL) in a 1.5mL Lo-bind tube was added 32 µL of the *click-cocktail*. The resulting solution was vortexed and incubated on a rotisserie at room temperature for 1 hour and quenched by the addition of 5 µL 250 mM $Na_4$EDTA. The mixture was cooled to 0 °C, transferred to a 15mL tube, and diluted with 4.2 mL ice-cold acetone. The samples were precipitated at –20 °C overnight, centrifuged at 4.5x1000g for 20 mins at 4 °C, and the supernatant removed. The pellet was fully resuspended in ice-cold methanol (1 mL) by sonication (2 s at 20%) and incubated at –20 °C for 30 minutes. After such time, the mixture was centrifuged at 4.5x1000g for 20 mins at 4 °C and the supernatant removed. The procedure was repeated. The pellet was allowed to air dry for 20 mins at room temperature and redissolved in 300 µL 1% SDS (1h at room temperature) and heated for 5 mins at 95 °C. The samples were cooled and diluted with 900 µL RIPA buffer. 250 µL of streptavidin magnetic beads (Thermo Fisher, cat. 88817) were added to Protein LoBind microcentrifuge tubes (Eppendorf, cat. 022431081) and washed 2x with 1 mL RIPA Buffer (Thermo Fisher, cat. 89900). Approximately 1.0 mg of cell lysate was added to the pre-washed streptavidin magnetic beads and incubated for 3 hours at room temperature. A magnetic rack was used to pellet the beads and remove the lysate supernatant. The beads were sequentially washed 3x with each of the following: 1 mL of 1% SDS, 1 mL of 1M NaCl, and 1 mL of 10% EtOH, all prepared in 1x DPBS and incubating for 5 min in between washes. A final wash was done with 1 mL RIPA Buffer. The beads were then resuspended in 30 µL of 4x Laemmli sample buffer (Boston

BioProducts, cat. BP-110R) containing 20 mM DTT and 25 mM biotin. Beads were heated for 10 min at 95 °C and were then placed on the magnetic rack. The supernatant was transferred to a new Protein LoBind microcentrifuge tube and stored at -80 °C. Western blot analysis was performed as previously described.

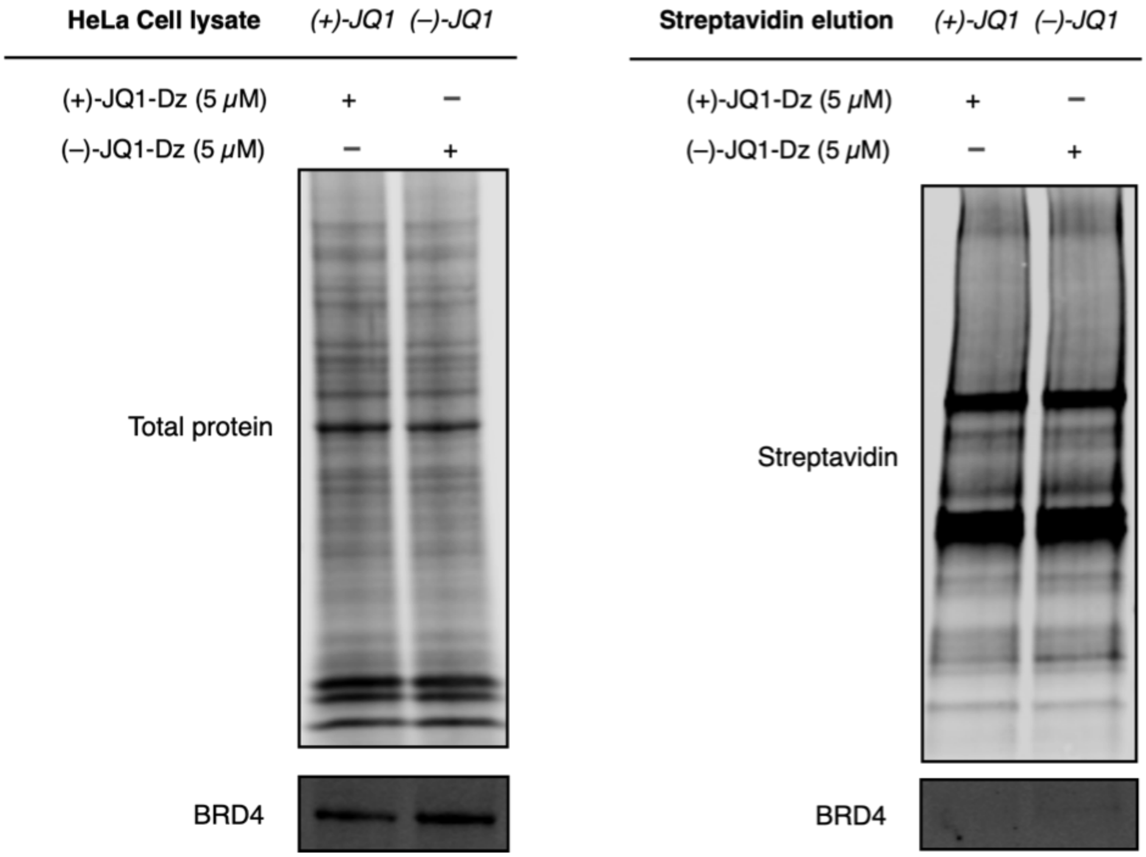

#### PAL proteomics using JQ1-Dz-alkyne

Following previously described procedure for JQ1-Dz-alkyne labelling and workflow for TMT proteomics after streptavidin bead washing.

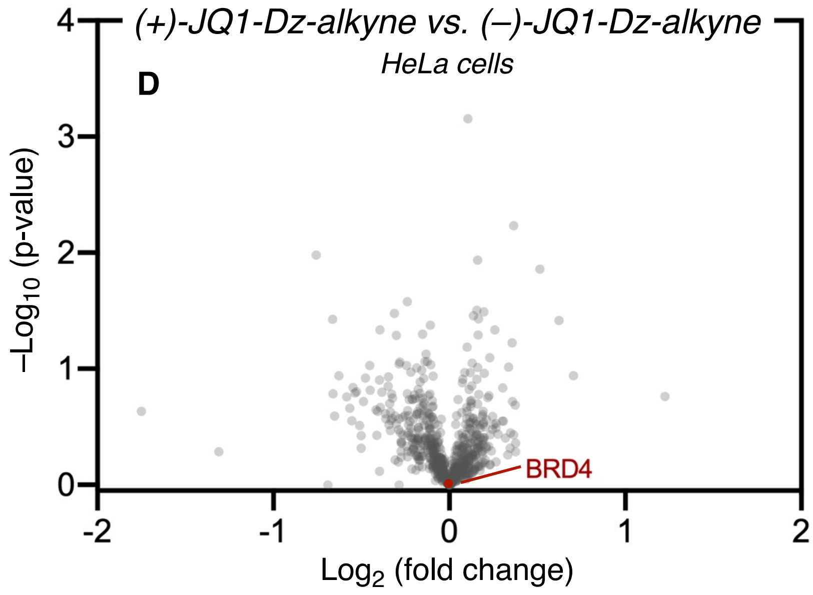

In addition to this workflow, we also conducted TMT-based proteomics in HeLa cells comparing both enantiomers of the JQ1-Dz-alkyne conjugate, based upon the workflow as described in Yao and coworkers[6]. Similarly, no enrichment of BRD proteins was observed.

***See attached data files for full lists of proteins for JQ1-based proteomics***

### Intracellular labelling of kinases using Dasatinib-Iridium

#### Synthetic Procedures

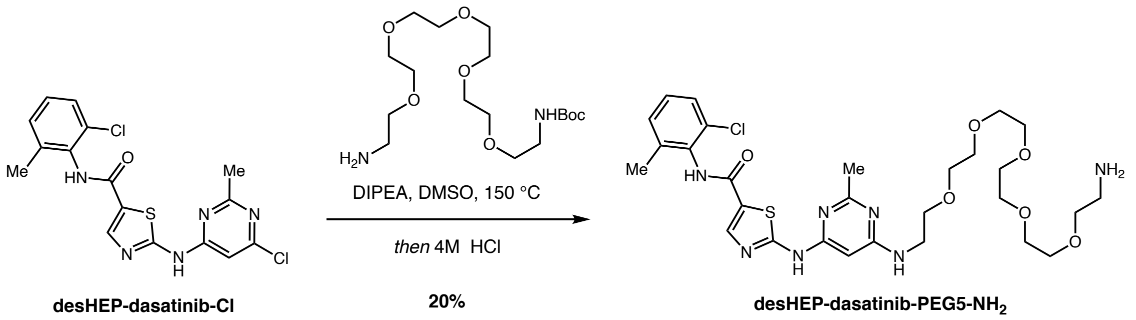

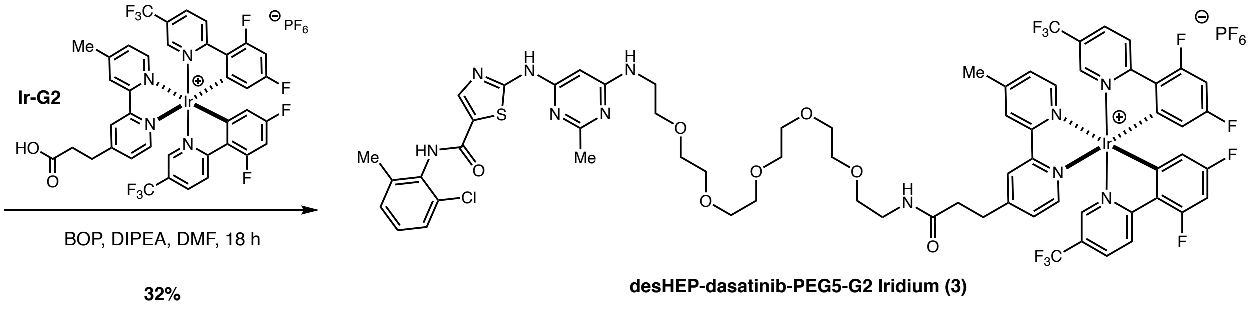

*desHEP-dasatinib-PEG5-NH₂*

desHEP-dasatinib-Cl (90 mg, 0.23 mmol) and BocNH-PEG5-$NH_2$ (87 mg, 0.23 mmol) were combined in a sealable reaction tube and DMSO (1 mL) and diisopropylethylamine (0.12 mL, 069 mmol) were added. The reaction vial was sealed and the mixture heated at 150 °C for four hours, then allowed to cool to room temperature. The reaction mixture was then diluted to 4 mL with DMSO, filtered through a 0.45 μm Acrodisc then purified in portions by preparative LCMS. The product fractions were combined and concentrated by Genevac. The resulting residue is a mixture of the desired product and the Boc-protected intermediate. A portion of this residue (60 mg) was treated with 4 N HCl in dioxane (1 mL) and stirred at room temperature for one hour. It was then dissolved in methanol (10 mL) and concentrated by rotovap and this step was repeated once more. The resulting material was dissolved in DMSO (1.8 mL), filtered 0.45 μm Acrodisc then purified in portions by preparative LCMS (10-40% acetonitrile/water (0.16%TFA) over 12 minutes, 25 mL/min, Waters CSH C18 5 μm 100 x 19 mm) The resulting product fraction was lyophilized to give desHEP-dasatinib-PEG5-$NH_2$ as white solid (37 mg, 0.06 mmol, 20% yield).

$^1$H NMR (500 MHz, Methanol-$d_4$) δ 8.24 (s, 1H), 7.39 (dd, $J$ = 7.4, 1.8 Hz, 1H), 7.32 – 7.23 (m, 2H), 6.35 (s, 1H), 3.77 – 3.60 (m, 22H), 3.16 – 3.10 (m, 2H), 2.66 (s, 3H), 2.34 (s, 3H). $^{13}$C NMR (126 MHz, Methanol-$d_4$) δ 138.88, 132.74, 132.68, 128.82, 128.35, 126.99, 70.07, 70.02, 69.94, 69.90, 69.89, 69.68, 69.00, 66.45, 41.64, 39.20, 17.27. LCMS: m/z calcd. for $C_{28}H_{40}ClN_7O_6S$ 637.2449 found 638.5

*desHEP-dasatinib-PEG5-G2 Iridium*

Ir-G2 (48 mg, 0.05 mmol) and BOP (33 mg, 0.08 mmol) were dissolved in DMF (0.23 mL) and diisopropylethylamine (0.03 mL, 0.15 mmol) was added. The resulting solution was added to desHEP-dasatinib-PEG5-$NH_2$ (32 mg, 0.05 mmol) and the resulting mixture diluted with DMF (0.23 mL). The reaction mixture as stirred at room temperature for one hour then addition BOP (15 mg, 0.04 mmol) was added before stirring the reaction overnight. The reaction mixture was then diluted with DMSO to 1.8 mL filtered 0.45 μm Acrodisc then purified in portions by preparative LCMS (36-66% acetonitrile/water (0.16%TFA) over 12 minutes, 25 mL/min, Waters CSH C18 5 μm 100 x 19 mm). The resulting product fraction was then lyophilized to give the desHep dasatinib-PEG5-G2 iridium conjugate (3) (25 mg, 0.016 mmol, 32% yield) as a yellow solid.

$^1$H NMR (500 MHz, $CD_3OD$) δ 8.71 (s, 2H), 8.58 (dd, $J$ = 8.8, 2.4 Hz, 2H), 8.35 – 8.29 (m, 2H), 8.24 (s, 1H), 7.96 (d, $J$ = 5.7 Hz, 1H), 7.92 (d, $J$ = 5.7 Hz, 1H), 7.75 (s, 1H), 7.70 (s, 1H), 7.60 – 7.53 (m, 2H), 7.37 – 7.31 (m, 1H), 7.30 – 7.21 (m, 2H), 6.87 – 6.78 (m, 2H), 6.41 (s, 1H), 5.79 (td, $J$ = 8.4, 2.3 Hz, 2H), 3.74 (t, $J$ = 4.9 Hz, 2H), 3.70 – 3.58 (m, 17 H) , 3.56 – 3.53 (m, 2H), 3.51 – 3.40 (m, 2H), 3.30 (td, $J$ = 5.5, 1.9 Hz, 2H), 3.19 (t, $J$ = 7.3 Hz, 2H), 2.69 (t, $J$ = 7.4 Hz, 2H), 2.65 (s, 6H), 2.33 (s, 3H). $^{13}$C NMR (126 MHz, $CD_3OD$) δ 172.35, 167.84, 167.78, 165.97, 163.90, 163.80, 161.57, 161.45, 156.36, 155.43, 155.21, 155.08, 155.02, 154.94, 153.78, 150.25, 149.93, 145.17, 138.88, 136.94, 132.73, 132.69, 129.61, 128.82, 128.78, 128.33, 126.96, 126.42, 125.94, 125.35, 123.79, 123.62, 123.04, 120.88, 113.81, 113.74, 99.42, 99.21, 98.99, 70.15, 70.11, 70.07, 70.05, 70.03, 69.73, 69.00, 41.89, 38.90, 34.67, 30.54, 20.49, 20.09, 17.29. $^{19}$F NMR (471 MHz, $CD_3OD$) δ -64.36, -64.43, -104.45, -108.03. HRMS (ESI-TOF): m/z calcd. for $C_{66}H_{62}ClF_{10}IrN_{11}O_7S$ 1570.3707 found 1570.3647

#### Other desHEP-dasatinib PEG-G2 Iridium conjugates prepared (3a–c)

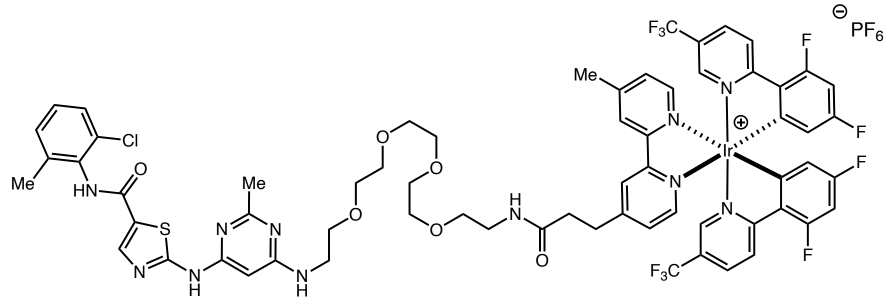

**desHEP-dasatinib-PEG4-G2 Iridium (3a)**

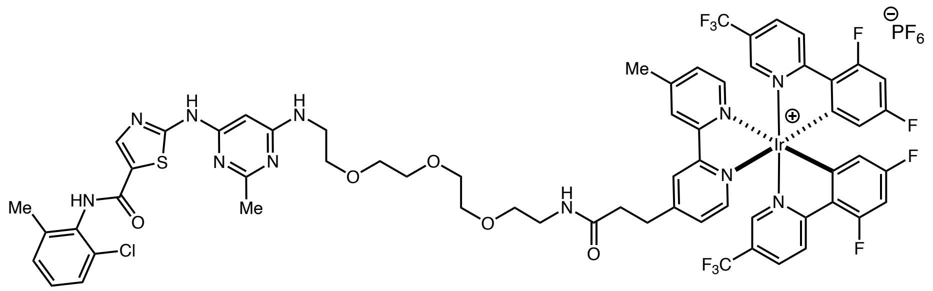

**desHEP-dasatinib-PEG3-G2 Iridium (3b)**

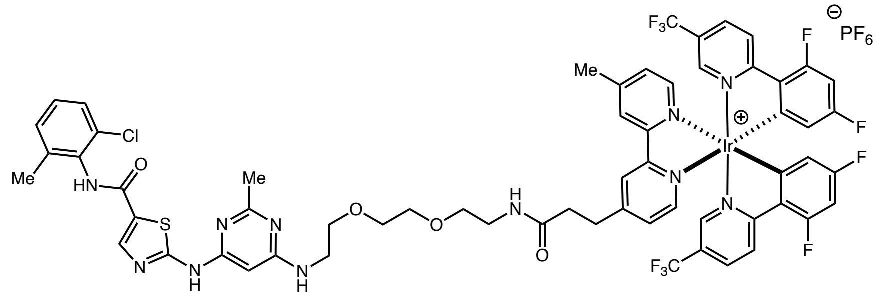

**desHEP-dasatinib-PEG2-G2 Iridium (3c)**

*All conjugates were prepared analogously using the corresponding BocNH-PEG**X**-$NH_2$ for the first step.*

**Dasatinib-PEG3-iridium (4)**

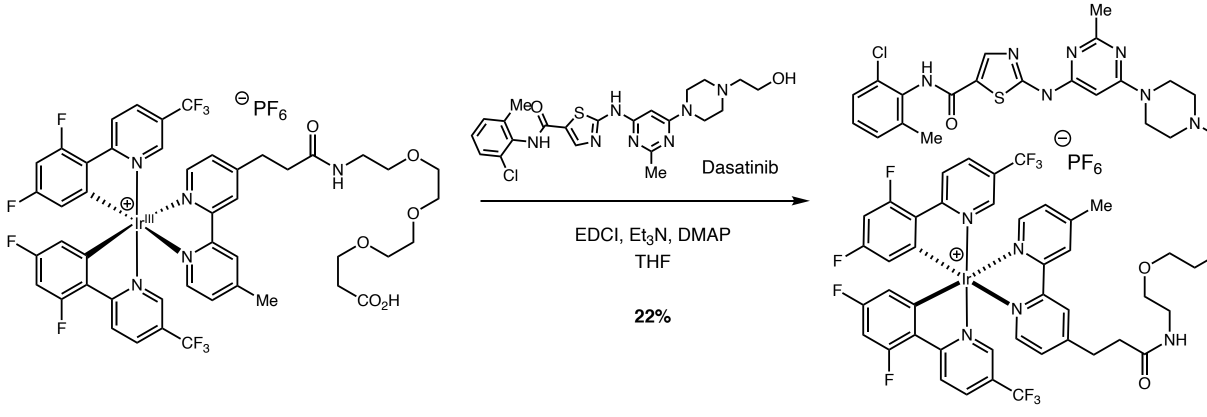

Ir-G2-PEG3-CO$_2$H (17) Dasatinib-PEG3-G2-Iridium (4)

To a stirred solution of Ir-G2-PEG3-CO$_2$H (17) (26 mg, 0.02 mmol) in DMF (0.5 mL) was cooled to 0 °C before the addition of EDCI (3.7 mg, 0.024 mmol) and Et$_3$N (5.6 μL, 0.04 mmol) under N$_2$. The reaction mixture was stirred for 1 h before the addition of dasatinib (19.5 mg, 0.04 mmol) and DMAP (3.7 mg, 0.03 mmol). The reaction mixture was slowly warmed to room temperature and left to stir for 24 h. Upon completion, the reaction mixture was diluted with EtOAc, and quenched by the addition of saturated aqueous NaHCO$_3$. The aqueous phase was removed, and the organic layer washed with additional saturated aqueous NaHCO$_3$, brine, and dried over Na$_2$SO$_4$. The solvent was removed *in vacuo*, and the crude material purified by silica column chromatography (gradient elution: 0 to 2% MeOH/CH$_2$Cl$_2$) followed by C8 reverse phase chromatography (gradient elution: 30 to 100% MeCN/H$_2$O (0.1% formic acid)) to give Dasatinib-PEG3-G2-Iridium (4) as a yellow solid (7 mg, 22%).

$^1$H NMR (500 MHz, CDCl$_3$) δ 8.94 (s, 2H), 8.46 (ddd, *J* = 16.3, 8.8, 3.2 Hz, 3H), 8.25 (t, *J* = 5.5 Hz, 1H), 8.03 (ddd, *J* = 13.5, 8.8, 2.1 Hz, 2H), 7.98 – 7.79 (m, 2H), 7.73 (dd, *J* = 11.3, 5.6 Hz, 2H), 7.61 (d, *J* = 2.0 Hz, 1H), 7.54 (d, *J* = 2.0 Hz, 1H), 7.49 (dd, *J* = 5.7, 1.6 Hz, 1H), 7.30 (dd, *J* = 5.7, 1.5 Hz, 1H), 7.21 – 7.05 (m, 2H), 6.64 (ddt, *J* = 11.5, 8.7, 2.5 Hz, 2H), 6.17 (s, 1H), 5.62 (dt, *J* = 7.9, 2.5 Hz, 2H), 4.27 (t, *J* = 5.3 Hz, 2H), 3.75 (t, *J* = 6.1 Hz, 2H), 3.67 (t, *J* = 4.9 Hz, 3H), 3.59 (d, *J* = 8.6 Hz, 8H), 3.50 (t, *J* = 5.6 Hz, 2H), 3.34 (q, *J* = 5.6 Hz, 2H), 3.21 (dt, *J* = 8.3, 4.0 Hz, 2H), 2.84 (qt, *J* = 14.8, 7.2 Hz, 2H), 2.73 (t, *J* = 5.3 Hz, 2H), 2.60 (q, *J* = 5.7 Hz, 8H), 2.48 (s,

3H), 2.32 (s, 3H). $^{13}$C NMR (126 MHz, $CDCl_3$) δ 172.0, 171.8, 168.1 (dd, $J$ = 16.7, 6.6 Hz), 166.9, 166.0 (d, $J$ = 12.6 Hz), 165.9, 164.0 (d, $J$ = 14.2 Hz), 163.7 (d, $J$ = 10.5 Hz), 163.0, 161.6 (d, $J$ = 13.0 Hz), 157.5, 156.7, 155.6, 155.5, 155.1 (d, $J$ = 6.9 Hz), 154.8 (d, $J$ = 6.9 Hz), 154.3, 149.7, 149.2, 145.2, 143.1 (d, $J$ = 509.5 Hz), 138.7, 136.6, 132.7, 129.9, 129.5, 129.4, 128.0, 127.7, 127.2, 126.5, 126.4, 126.1, 123.8 (dd, $J$ = 25.0, 21.0 Hz), 121.7 (dd, $J$ = 272.8, 8.4 Hz), 114.2 (dd, $J$ = 18.5, 8.2 Hz), 100.1 (td, $J$ = 26.8, 13.9 Hz), 83.6, 70.5, 70.5, 70.3, 69.7, 66.8, 61.9, 56.6, 52.8, 44.0, 39.3, 35.4, 35.3, 31.3, 25.5, 21.7, 19.2. $^{19}$F NMR (282 MHz, $CDCl_3$) δ -62.68 (dd, $J$ = 19.5, 3.9 Hz), -101.44 (dd, $J$ = 51.7, 12.6 Hz), -105.84 (dd, $J$ = 39.1, 12.6 Hz).

m/z HRMS found $[M+H]^+$ = 1624.4023, $[C_{69}H_{66}ClF_{10}IrN_{12}O_7S]^+$ requires 1624.4046. HPLC (Vydac 218TP C18 HPLC, gradient: 0 – 90% MeCN/$H_2O$ (0.1% TFA) 10 minutes, 5 minutes 90% MeCN (0.1% TFA), 1 mL/min, 254 nm): $\tau_r$ = 11.0 min.

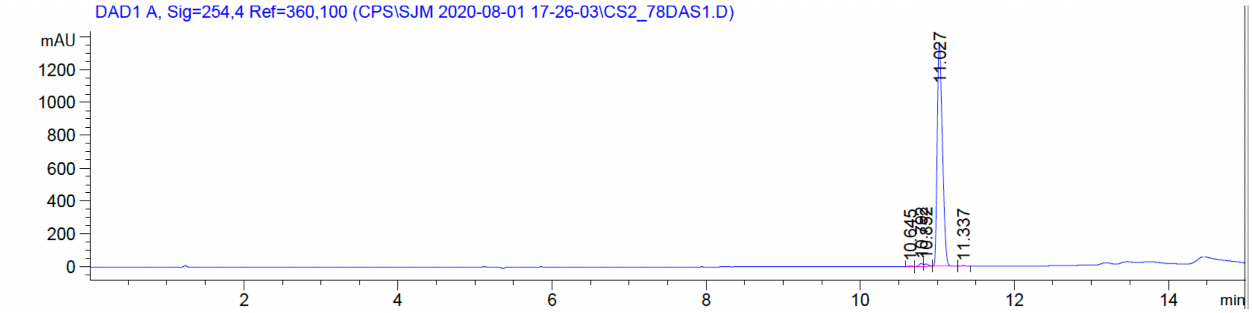

**Dasatinib-Dz-alkyne (5)**

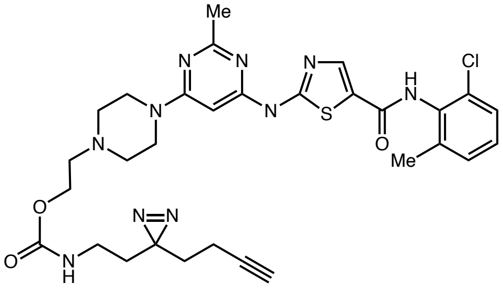

To a solution of dasatinib (200 mg, 0.4 mmol) in anhydrous THF (100 mL) was added 4-nitrophenyl chloroformate (90 mg, 0.44 mmol, 1.1 equiv.) and DIPEA (200 μL, 1.1 mmol). The reaction was stirred at room temperature under $N_2$ for 6 hours before an additional portion of 4-nitrophenyl chloroformate (90 mg, 0.44 mmol) was added. The reaction mixture was stirred

overnight. 2-(3-but-3-yn-1-yl)-3H-diazirin-3-yl)ethan-1-amine (67 mg, 0.50 mmol) and DIPEA (200 $\mu$L, 1.1 mmol) were added to the solution and stirring continued for an additional 24 hours. The reaction mixture was concentrated *in vacuo* and residue purified by reverse phase (C8) preparative HPLC (0 to 100% MeCN/$H_2O$ (0.1% FA)) to give the product (5) as a pale-yellow glass (103 mg, 40%). $^1$H NMR (500 MHz, DMSO-$d_6$) δ: 11.48 (br. s, 1H), 9.89 (s, 1H), 8.23, (s, 1H), 8.15 (s, 1H), 7.41 (d, $J$ = 7.7 Hz, 1H), 7.31 – 7.23 (m, 2H), 7.21 (t, $J$ = 5.0 Hz, 1H), 6.05 (s, 1H), 4.11 – 4.06 (m, 2H), 2.90 – 2.82 (m, 3H), 2.58 – 2.53 (m, 2H), 2.41 (br. s, 3H), 2.24 (br. s, 3H), 1.99 (dt, $J$ = 7.7, 2.7 Hz, 2H), 1.59 (t, $J$ = 7.7 Hz, 2H), 1.52 (t, $J$ = 7.1 Hz, 2H). $^{13}$C NMR (125 MHz, DMSO-$d_6$) δ: 165.6, 163.7, 163.0, 162.8, 160.4, 157.4, 156.5, 141.3, 139.3, 134.0, 132.9, 129.5, 128.7, 127.5, 126.2, 83.6, 83.1, 72.3, 61.5, 57.1, 52.9, 44.0, 43.4, 35.8, 32.7, 31.8, 27.6, 26.1, 18.8, 13.1. *Peaks in both $^1$H and $^1$3C NMR present under residual solvent.*
m/z HRMS found $[M+H]^+$ = 651.24181, $[C_{30}H_{36}ClN_{10}O_3S]^+$ requires 651.23756.

**Cell Culture:**
THP1 cells (ATCC, cat. TIB-202) were grown in RPMI (Lifetech cat. 11875-093) with added 10% FBS (Lifetech cat. 10082-147), NEAA (Lifetech 11140050), 1X penicillin-streptomycin-glutamine (Lifetech cat. 10378-016) and 40 μM betamercaptoethanol (Sigma M3184) in T175 flasks at 37 °C under 5% $CO_2$.

#### Labelling of recombinant p38 and Abl using Dasatinib-PEG3-Ir

*Representative example:*
To a 1.5 mL Eppendorf tube was added a DPBS solution containing Abl or p38 (0.5 μM), bovine carbonic anhydrase (0.5 μM), either Dasatinib-G2-Ir (4) (0.4 μM) or Ir-NHBoc (15) (0.4 μM), either Dasatinib (5 μM, 10x off-compete) or DMSO vehicle, and Dz-PEG3-biotin (9) (100 μM). The sample was irradiated 15 minutes at 450 nM in the Efficiency Aggregator PhotoReactor. The sample was subsequently diluted with 4x Laemlli buffer with BME (5 μL) and heated to 95 °C for 10 minutes. The samples were cooled to room temperature and centrifuged. The samples were subsequently loaded onto a BioRad Criterion 10% tris-glycine gel, alongside all of the appropriate controls, and run in freshly prepared Tris running buffer (150V, 60 minutes). The gel was washed (3 x MiliQ water) and transferred to an NC membrane. Following transfer, the membranes were

then immersed in REVERT total protein stain (Li-Cor, 926-11011) for 5 minutes. Excess stain was decanted, membranes washed with 6.7:30:63.3 AcOH:MeOH:$H_2O$ and imaged using a Li-Cor Odyssey CLx scanner in the 700 nm channel. The membranes were washed with water, then immersed in Odyssey Blocking Buffer (Li-Cor, 927-50000) and incubated for 1 hour. The blocking solution was then decanted, and 35 mL of fresh blocking buffer containing 70 µL of Tween 20 was added. This mixture was rocked for 5 minutes. Afterwards, 1.5 µL of IRDye 800CW streptavidin (Li-Cor, 926-32230) was added and the mixture incubated for 1 hour. The blocking buffer was then decanted, and the membranes were washed with 1X TBST (3 x 5 min) and water before imaging via Li-Cor Odyssey CLx scanner in the 800 nm channel. Pixel densitometry was performed using Image Studio Lite V. 5.2 (Li-Cor). The streptavidin 800 channel pixel density was then divided by the total protein stain 700 channel pixel density to provide a normalized biotinylation signal for each protein band.

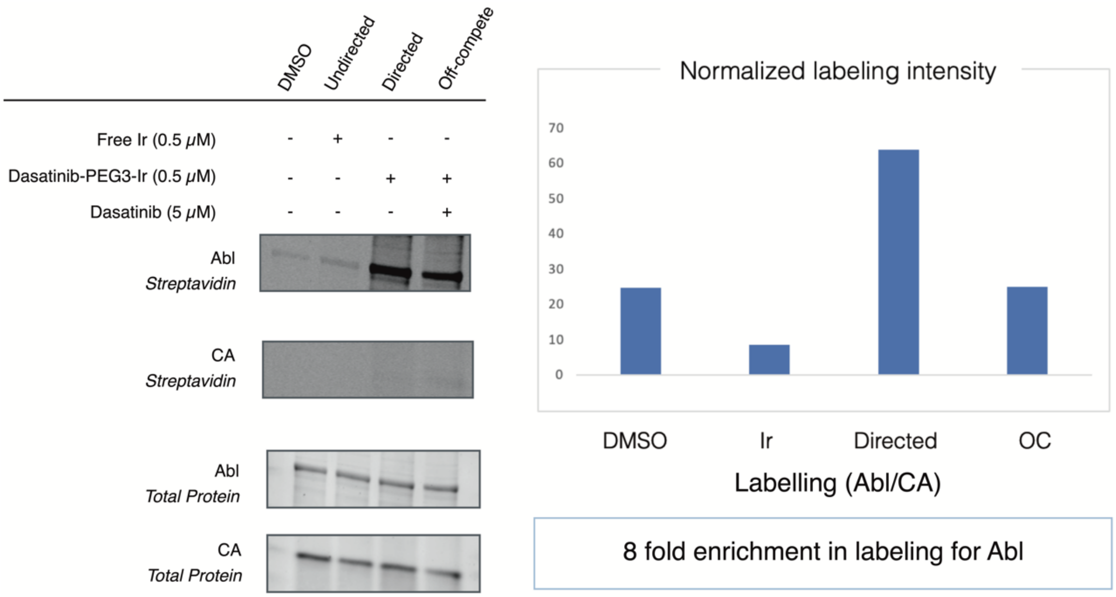

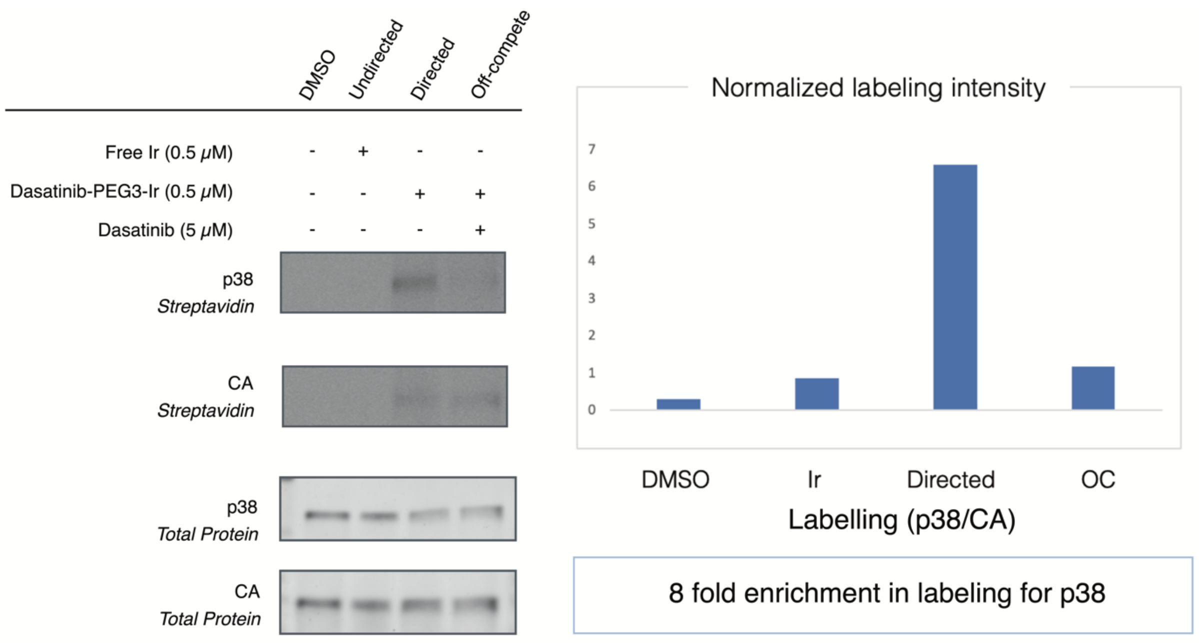

#### Kinase inhibition assays in K562 cells

##### Dephosphorylation assay

A 24 well plate was seeded with 500k K562 cells per well in 2 mL of IMDM. To each well was added either Dasatinib (5 µM), Dasatinib-Gen2-Ir (5) (5 µM), or DMSO. The cells were incubated for 24 hours, pelleted, and washed with PBS. Cell pellets were lysed (20 mM Tris, 1 mM β-glycerophosphate, 1 mM $Na_3VO_4$, 150 mM NaCl, 1 mM EDTA, 1 mM EGTA, 1% NP-40, 1% Na deoxycholate, 2.5 mM Na pyrophosphate, 0.1% SDS) and 10 µg of lysate was analysed by western blot, staining for p38 (Cell signalling: 9212), phos-p38 (Thr180/Tyr182) (Cell Signalling: 9211), abl (Cell Signalling: 2862S), phos-Abl (Cell Signalling: 2861S), P-Tyrosine (Cell Signalling: 9411S) and GAPDH (SCBT; 47724).

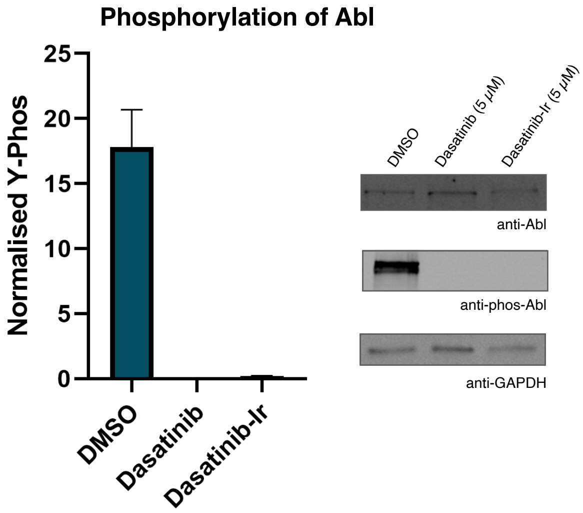

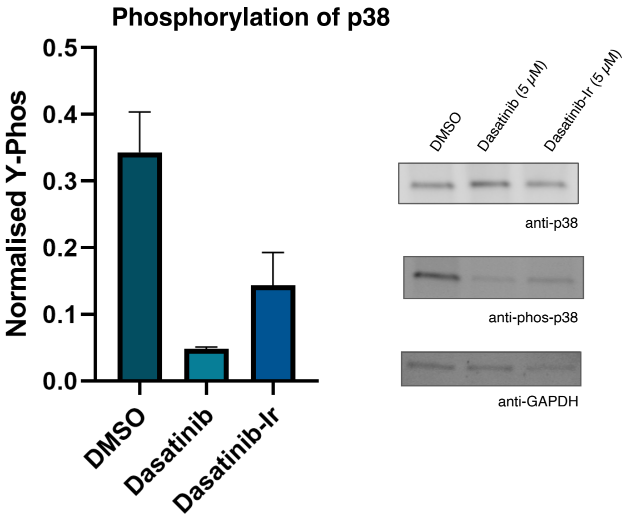

#### Dasatinib μMap target ID experiments

**Labeling experiment protocol 1 (THP1):**

THP1 cells in DPBS (approx. 9 million cells per replicate) were treated with Dz-PEG3-biotin (9) (250 μM) and contained either Ir-Gen2-NHBoc (16) (10 μM) or desHep-dasatinib-PEG5-G2 iridium conjugate (3) (10 μM), or (4) (10 μM) plus dasatinib (100 μM) in 1.5 mL eppendorf tubes. Each tube was incubated with end-over-end mixing at room temperature for one hour. Tubes were then subjected to 450 nm light for 30 minutes at room temperature. Cells were then washed twice by centrifugation at 5000xg for 2 minutes at 4 °C followed by removal of supernatant and resuspension in DPBS. After a third centrifugation, supernatant was removed and the cells were treated with RIPA (Thermofisher cat. 89900) buffer containing 1X protease inhibitors (Promega G6521) (1 mL). These samples were then sonicated at 60% power for 3-5 seconds then subjected to the pulldown protocol below.

**Pulldown:**

250 μL of streptavidin magnetic beads (Thermo Fisher, cat. 88817) were added to Protein LoBind microcentrifuge tubes (Eppendorf, cat. 022431081) and washed 2x with 1 mL RIPA Buffer (Thermo Fisher, cat. 89900) (using a magnetic rack to pellet the beads for each wash). Approximately 3 mg of cell lysate for each sample was added to the pre-washed streptavidin magnetic beads and incubated for 3 hours at room temperature with end-over-end mixing. A magnetic rack was used to pellet the beads and remove the lysate supernatant. The beads were

washed with 3x with RIPA buffer with 5 minute incubations with end-over-end mixing. The beads were then resuspended in 45 µL of 2.5:1 (4x NuPage LDS buffer (Thermofisher NP0007): NuPage sample reducing agent (Thermo Fisher NP0004)) which contains 25 mM Biotin. Beads were heated for 10 min at 70 °C and were then placed on the magnetic rack. The supernatant was transferred to a new Protein LoBind microcentrifuge tube. Approximately 10–15 µL was removed for gel and WB, and the remainder of each sample stored at -80 °C.

**Gel and WB:**

Gel electrophoresis was performed using the XCell SureLock Mini-Cell Electrophoresis System (Thermo Fisher, cat. EI0001). Samples were loaded onto a 12 % bistris NuPage gel (under reducing conditions) and run at 200 V for 35 min to resolve proteins using 1X MES SDS buffer (Invitrogen, cat. NP0002) and molecular weight ladder SeeBlue Plus2 prestained protein standard (Invitrogen, cat. LC5925). Proteins were blotted onto a PVDF membrane (Invitrogen, cat. IB24001) using iBlot2 gel transfer device (Thermo Fisher, cat. IB21001,IB23001) with settings as 23V for 7 min. The membrane was blocked with 10 mL of 3% BSA (Sigma, cat. A3983-100G) TBST (0.1% tween 20 in 1X TBS, Thermo Scientific, cat. 28358) buffer overnight at room temperature, followed by incubation with anti-MAPK14 (abcam ab31828) (1:1,000 dilution, 10 mL) for 1 hour then by incubation with LiCOR IRDYE-800CW goat-anti-mouse (926-32210) (1:15,000 dilution, 10 mL) for 1 hour. Membranes were subsequently washed 3 times with TBST (5 minutes per

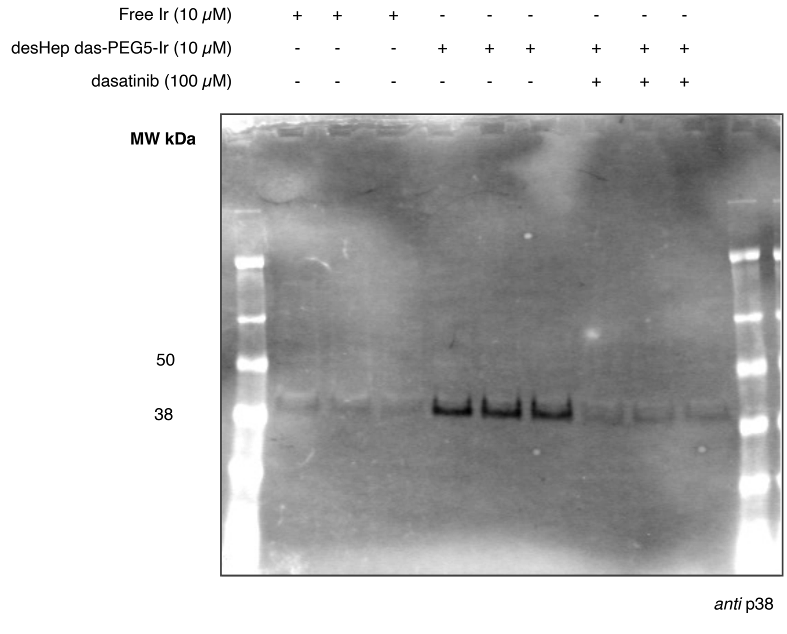

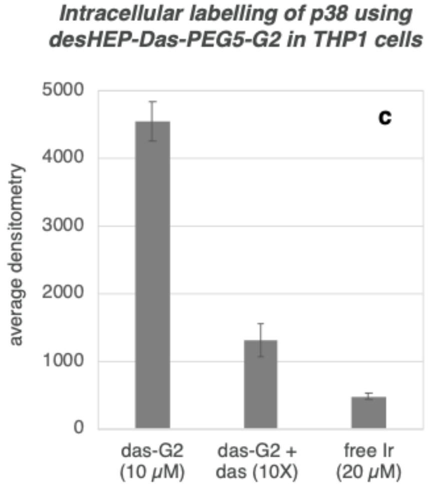

wash), then imaged LI-COR Odyssey using automatic settings. Labelling intensities were normalized by protein concentration and plotted.

#### Dasatinib-PEG5-G2 label free proteomics

**Sample Preparation for Mass Spectrometry:**

Samples were reduced and alkylated with 10mM TCEP and 20mM iodoacetamide for 30 minutes at 65 degrees. Samples were prepared for mass spectrometry using the SP3 method as previously described[8]. Subsequent to elution from SP3 beads, samples were desalted using C18 columns.

**LC-MS/MS Data Acquisition:**

Samples were resuspended in 95% water and 5% acetonitrile analyzed with 0.1% formic acid and analyzed on a QE-HF mass spectrometer coupled to a Dionex 3000 LC system. The LC system used a C18 analytical column with a 2 hour gradient from 3% to 40% acetonitrile. The mass spectrometer was operated in data dependent mode with a top 20 method.

**Data Analysis:**

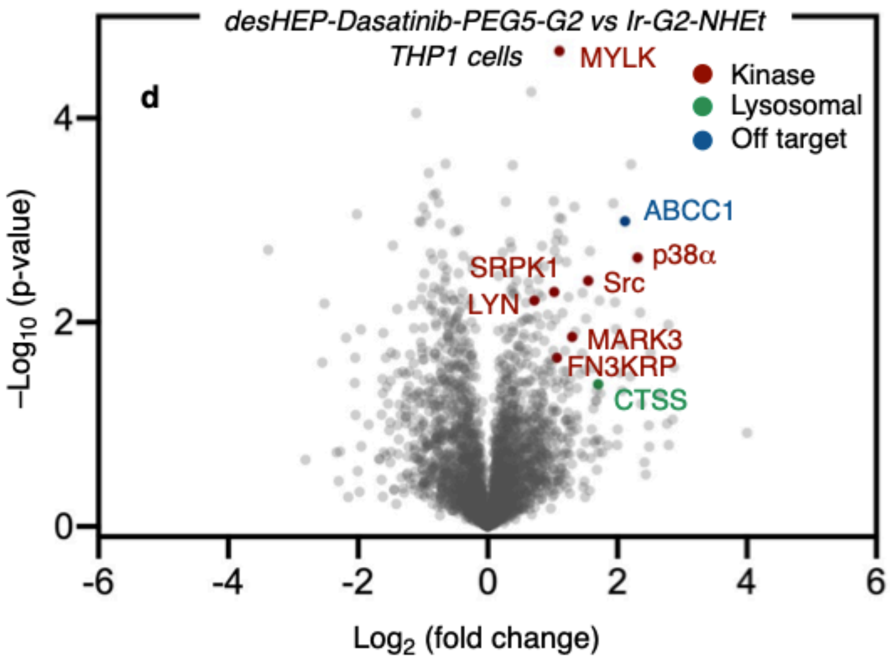

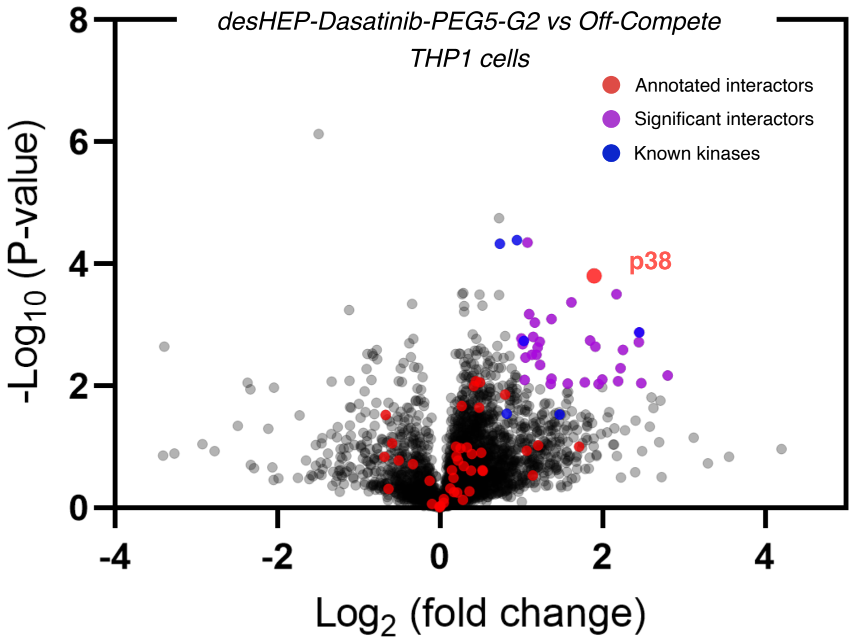

*Please see associated datafiles for full list of interactors.*

Data was searched with Maxquant version 1.6.17 using the align between runs feature. Cysteine carbamidomethylation was selected as a fixed modification. Methionine oxidation and N-terminal protein acetylation were set as variable modifications. Data was analyzed in Perseus 1.6.14. Data was filtered to remove contaminants and reverse database hits. Proteins with at least one unique peptide with a peptide FDR of 1% were accepted. Protein intensities were Log2 transformed, median normalized, and missing values were imputed.

**Labeling experiment comparing desHep-PEG3 (3a), PEG4 (3b), and PEG5 (3c) analogs:**
Using protocol as above, THP1 cells were treated with a solution of diazirine-PEG3-biotin (9) (250 µM) in DPBS which contained either desHep-PEG3 dasatinib G2-iridium conjugate (**3a**) with or without 20 µM dasatinib, or 5 µM desHep-PEG4 dasatinib G2-iridium conjugate (**3b**) with or without 20 µM dasatinib or 5 µM desHep-PEG5 dasatinib G2-iridium conjugate (**3c**) with or without dasatinib (20 µM). Following incubation, irradiation with 450 nm light and centrifugation/washing and lysis steps described above, the samples were subjected to gel electrophoresis and WB also as described above.

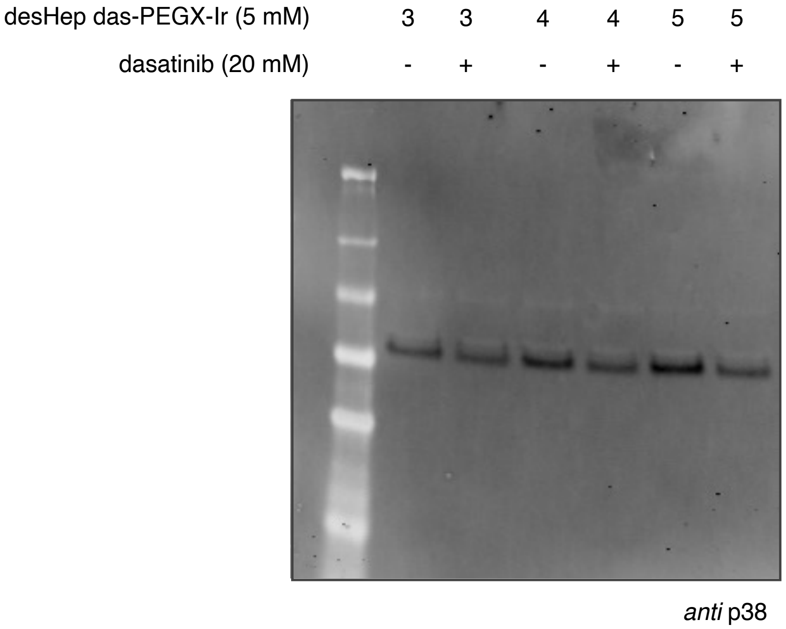

#### µMap labeling using Dasatinib-G2-Ir

200M K562 (grown in IMDM) cells were removed from T75 flasks and divided into 9 portions in IMDM (T25 flasks, 4 mL each). To three flasks (**B**, off compete) was added 50 µM dasatinib and the cells were incubated for 30 min at 37 °C. At this point, Dasatinib-G2-Ir (4) (5 µM) was added to 6 flasks (**A**, directed and **B,** off-compete), and Ir-dF($CF_3$)(dMebpy)$PF_6$ (14) (5 µM) was added to 3 flasks (**C**, free Ir) and the samples were incubated at 37 °C for a further 3 hours. At this stage, the cells were removed, collected, and resuspended in HEPES buffered saline (1 mL/plex) and transferred to 1.5 mL Eppendorf tubes. Diazirine-PEG3-biotin (9) was added to each tube (250 µM) and the samples were incubated at 23 °C for an additional 20 minutes. The tubes were subsequently irradiated in the MPR at 450 nM for 3 minutes at 100% intensity. The cells were then washed twice with cold DPBS (4 °C) and resuspended in 1 mL of cold kinase lysis buffer (20 mM Tris, 1 mM β-glycerophosphate, 1 mM $Na_3VO_4$, 150 mM NaCl, 1 mM EDTA, 1 mM EGTA, 1% NP-40, 1% Na deoxycholate, 2.5 mM Na pyrophosphate, 0.1% SDS) containing HALT protease and phosphatase inhibitor. The lysed cells were incubated on ice for 15 minutes and sonicated (35%, 5 x 5s with 30s rest). The lysate was then centrifuged at 15x1000g for 20 mins at 4 °C and the supernatant collected. The concentration of the cell lysate was measured by BCA assay (typically 2 mg/mL) and volumes adjusted accordingly to equal concentration for streptavidin enrichment. A sample was removed and stored at –20 °C for future analysis.

Streptavidin enrichment and proteomics preparation were carried out as described above.

**TMT-based chemoproteomics analysis of Dasatinib-Dz-alkyne labelling in K562 cells**

Following previously described procedure for Dasatinib-Dz-alkyne (5) labelling and workflow for CuAAC reaction and TMT proteomics after streptavidin bead washing.

#### Proteomics results

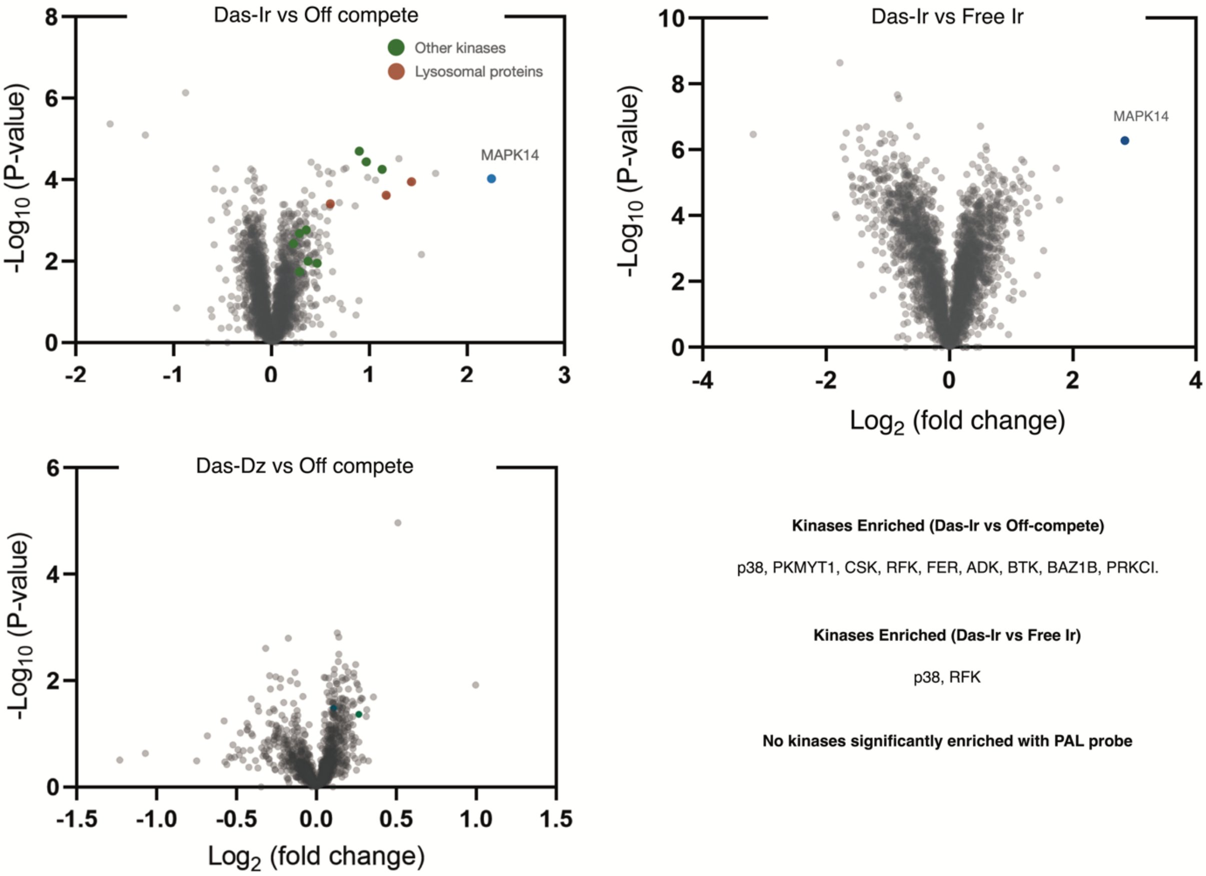

***See attached data files for full lists of proteins***

### Labelling of microtubules using Paclitaxel-Ir

#### Synthesis of conjugates

##### Paclitaxel-G2 (6)

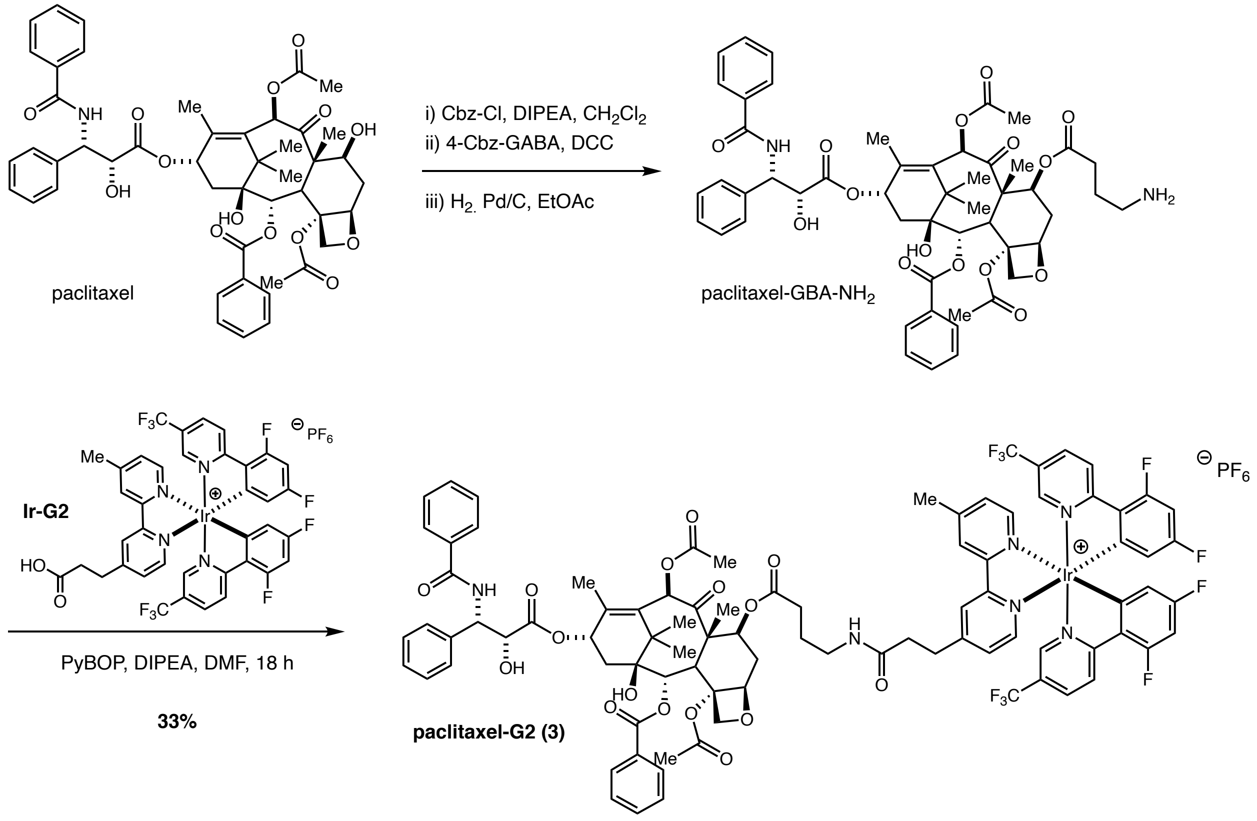

Following a previously reported procedure[9]. To a solution of Paclitaxel (312 mg, 0.37 mmol) in $CH_2Cl_2$ (8 mL) was added DIPEA (193 µL, 1.1 mmol) at room temperature. Cbz-Cl (156 µL, 1.1 mmol) was added and the mixture stirred for 3h under $N_2$. The solvent was removed *in vacuo* and the mixture purified by column chromatography (silica: 1:1 EtOAc/P.E.) to afford Cbz-Paclitaxel as a white solid (330 mg, 90%). Spectral data was consistent with literature reports.

Cbz-Paclitaxel (250 mg, 0.25 mmol), DMAP (30 mg, 1 equiv.), and 4-Cbz-aminobutyric acid (225 mg, 4.4 equiv.) was dissolved in $CH_2Cl_2$ (10 mL) and DCC (210 mg, 1 mmol) was added portion wise. The resulting mixture was stirred at room temperature under $N_2$ for 16 hours and filtered over a pad of celite. The solvent was removed *in vacuo* and the residue purified by column chromatography (silica: 1:1 EtOAc/P.E.) to afford the product as a white solid (quant. yield). Spectral data was consistent with literature reports[9].

To a solution of Cbz-Paclitaxel-GBA-Cbz (100 mg, 83 μmol) in EtOAc (2 mL) was added Pd/C (10%, 50 mg) and placed under an atmosphere of hydrogen (balloon). The mixture was stirred vigorously overnight, filtered over a pad of celite, and the solvent removed *in vacuo*. The resulting Paclitaxel-GBA-$NH_2$ was used without further purification (72 mg, 92%). Spectral data was consistent with literature reports[9].

To a stirred solution of Ir-G2 (13) (75 mg, 69 μmol), and PyBOP (55 mg, 105 μmol) in anhydrous DMF (1 mL) under $N_2$ in the dark was added DIPEA (30 μL, 172 μmol). The resulting mixture was stirred at room temperature for 10 minutes and a solution of Paclitaxel-GBA-$NH_2$ (66 mg, 70 μmol) in anhydrous DMF (1 mL) was added dropwise. The reaction was stirred overnight, diluted with EtOAc, and quenched by the addition of saturated aqueous $NaHCO_3$. The aqueous phase was removed and the organic layer washed with additional saturated aqueous $NaHCO_3$, 5% aqueous citric acid, brine, and dried over $Na_2SO_4$. The solvent was removed *in vacuo*, and the crude material purified by silica column chromatography (gradient elution: 0 to 3% MeOH/$CH_2Cl_2$) and C8 reverse phase preparative HPLC (gradient elution: 30 to 100% MeCN/$H_2O$ (0.1% formic acid)) to afford Paclitaxel-G2-Ir (6) as a yellow solid (47 mg, 33%).

$^1$H NMR (500 MHz, $CDCl_3$) δ: 8.77 (d, *J* = 7.3 Hz, 1H), 8.75 (s, 1H), 8.77 – 8.65 (m, 1H), 8.48 (t, *J* = 10.5 Hz, 2H), 8.14 – 7.99 (m, 4H), 7.92 – 7.77 (m, 2H), 7.82 (d, *J* = 7.3 Hz, 2H), 7.74 (t, *J* = 7.3 Hz, 2H), 7.66 – 7.28 (m, 13H), 7.04 – 6.94 (m, 1H), 6.64 (t, *J* = 9.4 Hz, 2H), 6.16 (s, 1H), 6.10 (t, *J* = 8.4 Hz, 1H), 5.79 – 5.68 (m, 1H), 5.67 – 5.57 (m, 3H). 5.55 – 5.45 (m, 1H), 5.29 (s, 1H), 4.90 (d, *J* = 9.6 Hz, 1H), 4.84 (d, *J* = 3.6 Hz, 1H), 4.27 (d, *J* = 8.9 Hz, 1H), 4.15 (d, *J* = 7.9 Hz, 1H), 3.87 (d, *J* = 7.9 Hz, 1H), 3.16 (app. s, 4H), 2.95 – 2.58 (m, 7H), 2.58 – 2.50 (m, 1H), 2.35 (app. s, 3H), 2.26 – 2.09 (m, 5H), 1.86 – 1.63 (m, 7H), 1.25 (app. s, 3H), 1.16 (s, 3H), 1.13 (s, 3H). $^{13}$C NMR (125 MHz, $CDCl_3$) δ: 202.03, 172.7, 172.5, 171.5 (d, *J* = 3.2 Hz), 170.5, 169.6, 169.5, 168.2 – 168.0 (m), 167.3, 167.0, 165.0 (dd, *J* = 262.5, 13.0 Hz), 262.7 (dd, *J* = 263.7, 13.0 Hz), 157.7, 153.4 – 155.2 (m), 155.1 – 155.0 (m), 154.8 – 154.6 (m), 149.7, 149.3, 145.1 – 144.8 (m), 140.8 (d, *J* = 2.6 Hz), 138.7 (d, *J* = 2.0 Hz), 136.8 – 136.6 (m), 134.1 (d, *J* = 2.1 Hz), 133.9, 132.8,

131.8, 130.3, 130.1 (d, $J$ = 6.1 Hz), 129.8, 129.3, 128.9, 128.8, 128.7, 128.1, 127.4, 126.4, 126.2, 123.9 (t, $J$ = 21.3 Hz), 122.7 (d, $J$ = 9.1 Hz), 120.6 (d, $J$ = 9.1 Hz), 114.2 (dd, $J$ = 16.5, 6.7 Hz), 100.1 (td, $J$ = 27.0, 9.8 Hz), 84.1, 81.0, 78.6, 76.5, 75.4, 74.5, 73.5, 71.6, 71.5, 71.5, 56.2, 55.9, 55.8, 53.6, 47.1, 43.3, 38.8, 35.5, 35.4, 35.3, 33.4, 31.2, 29.8, 26.5, 26.4, 23.8, 23.8, 22.7, 21.6, 21.0, 20.9, 14.6, 11.0. $^{19}$F NMR (376 MHz, $CDCl_3$) δ: –62.7 (d, $J$ = 5.6 Hz), –62.8 (d, $J$ = 5.0 Hz), –71.0, –72.9, –101.3 – –101.5 (m), –105.7 – –105.9 (m). m/z HRMS found $[M]^+$ = 1871. 51783 (100), 1872.51899 (89), 1869.51134 (55), 1870.51373 (55), 1873.51932 (52), 1874.52130 (22), $[C_{89}H_{80}F_{10}IrN_6O_{16}]^+$ requires 1871.50949 (100), 1872.51284 (96), 1869.50715 (60), 1870.51051 (57), 1873.51620 (46), 1874.51955 (14). HPLC (Vydac 218TP C18 HPLC, gradient: 0 – 90% MeCN/$H_2O$ (0.1% TFA) 10 minutes, 5 minutes 90% MeCN (0.1% TFA), 1 mL/min, 254 nm): $\tau_r$ = 13.3 min.

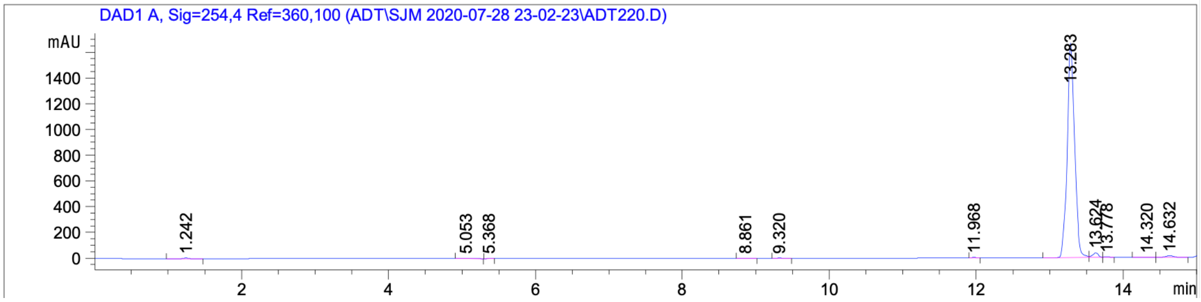

#### Cell viability assay

MCF7 Cells were grown to a confluency of about 80%, trypsinized, resuspended in fresh media and counted using a hemocytometer. Cells were diluted to 30,000 cells/ml and 100 µl was pipetted into a 96 well plate. Compounds were resuspended in DMSO at a concentration of 400 µm and diluted to 40 µm with water. Compounds were added to a final concentration of 2 µM, including controls for DMSO, to a final concentration of less than 1% DMSO. Cell viability was measured using EZ Quant Cell Quantifying Kit (ALSTEM) diluted 1:3 in PBS according to manufacturers. Briefly, at each time point , 20 µL of EZ Quant reagent was added (diluted 3-fold in PBS) and 2 h later the absorbance at 450 nm was recorded. Each viability experiment was performed in triplicate.

*Averaged absorbance values over 72 h (n=3)*

| A450 | 0h | 24h | 48h | 72h |
|---|---|---|---|---|
| Paclitaxel (2 μM) | 0.67 | 0.73 | 0.69 | 0.62 |
| Paclitaxel-G2 (6) (2 μM) | 0.76 | 0.77 | 0.76 | 0.54 |
| Ir(dFCF$_3$)(dMebpy)$^+$ (14) (2 μM) | 0.78 | 0.64 | 0.56 | 0.35 |
| DMSO | 0.68 | 0.97 | 1.60 | 1.44 |
| Control | 0.71 | 0.88 | 1.14 | 1.08 |

*Standard Deviation*

| SD | 0h | 24h | 48h | 72h |
|---|---|---|---|---|
| Paclitaxel (2 μM) | 0.07 | 0.14 | 0.10 | 0.10 |
| Paclitaxel-G2 (6) (2 μM) | 0.02 | 0.11 | 0.14 | 0.01 |
| Ir(dFCF$_3$)(dMebpy)$^+$ (14) (2 μM) | 0.01 | 0.10 | 0.05 | 0.01 |
| DMSO | 0.10 | 0.02 | 0.27 | 0.28 |
| Control | 0.04 | 0.08 | 0.03 | 0.04 |

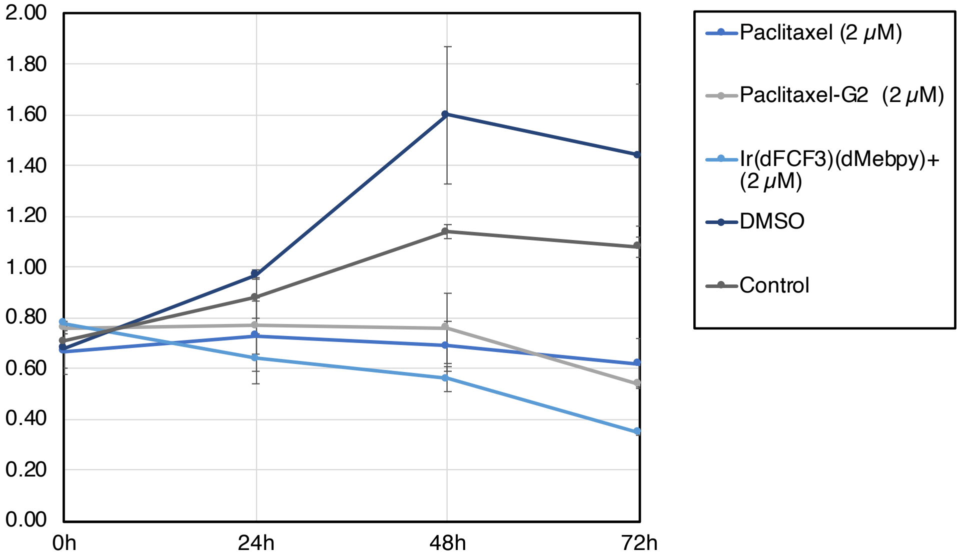

Notably, the MCF7 cell morphology changed when treated with both Paclitaxel and Paclitaxel-G2 (6). A pronounced shrinking or the membrane and rounding of the cells was observed in both cases, although viability was not impacted, the cells did not proliferate. While the free-Ir photocatalyst also resulted in loss of viability, cell death was more rapid and continuous, similar morphological changes were also not observed.

#### Paclitaxel-G2 labelling of recombinant tubulin

*Representative procedure:*

To a 0.5 mL Eppendorf tube was added DPBS (2.3 µL), β-tubulin (1.2 µL, 2 µM, DPBS) and BSA (1.0 µL, 5.9 µM, DPBS) followed by either Paclitaxel-G2 (6) (1.5 µL, 3.3 µM), free iridium (14) (2.5 µL, 1.9 µM), or Paclitaxel (1.5 µL, 3.3 µM). Dz-PEG3-biotin (9) (2 µL, 250 µM, DPBS) was added and the sample was irradiated 15 minutes at 450 nM in the Efficiency Aggregator PhotoReactor. The sample was subsequently diluted with 4x Laemlli buffer with BME (5 µL) and heated to 95 $^{o}$C for 10 minutes. The samples were cooled to room temperature and centrifuged. The samples were subsequently loaded onto a BioRad Criterion 4–20% tris-glycine gel, alongside all of the appropriate controls, and run in freshly prepared Tris running buffer (160V, 60 minutes). The gel was washed (3 x MiliQ water) and transferred via iBlot 2 to an NC membrane. Following transfer, the membranes were then immersed in REVERT total protein stain (Li-Cor, 926-11011) for 5 minutes. Excess stain was decanted, membranes washed with 6.7:30:63.3 AcOH:MeOH:$H_2O$ and imaged using a Li-Cor Odyssey CLx scanner in the 700 nm channel. The membranes were washed with water, then immersed in Odyssey Blocking Buffer (Li-Cor, 927-50000) and incubated for 1 hour. The blocking solution was then decanted, and 35 mL of fresh blocking buffer containing

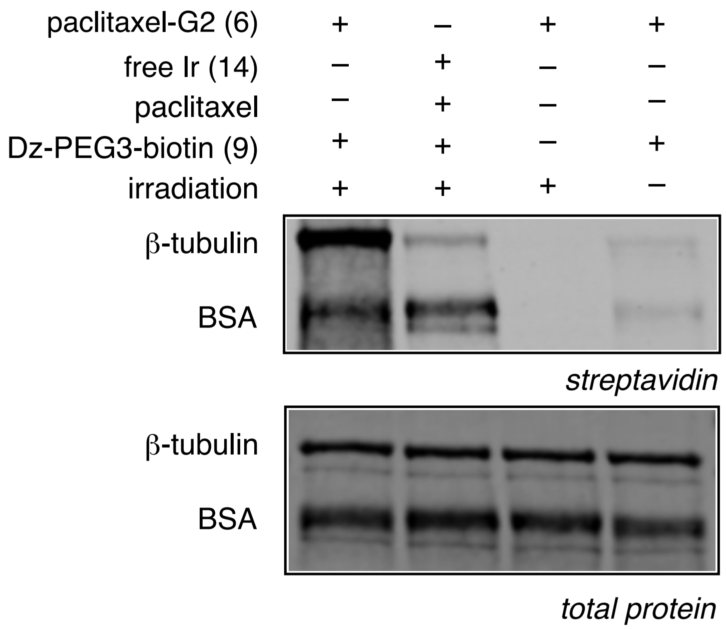

70 μL of Tween 20 was added. This mixture was rocked for 5 minutes. Afterwards, 1.5 μL of IRDye 800CW streptavidin (Li-Cor, 926-32230) was added and the mixture incubated for 1 hour. The blocking buffer was then decanted, and the membranes were washed with 1X TBST (3 x 5 min) and water before imaging via Li-Cor Odyssey CLx scanner in the 800 nm channel. Pixel densitometry was performed using Image Studio Lite V. 5.2 (Li-Cor).

#### μMap labelling of $\alpha$-tubulin using Paclitaxel-Ir

To MCF-7 cells in 15 clear 10 cm plate at 80% confluency in RPMI 1640 with **no** phenol red (Gibco) (4 mL) was added Paclitaxel-G2 (6) (20 μM) (5 plates, **A**), Ir-dF($CF_3$)(dMebpy)$PF_6$ (14) (2 μM) (5 plates, **B**), and DMSO control (5 plates, **C**). The plates were incubated at 37 °C for 3 hours and the media removed and replaced. Diazirine-PEG3-bioitin (9) was added (250 μM) and the plates incubated at 37 °C for an additional 20 minutes. The plates were subsequently irradiated (without the lid) in the bioreactor at 450nM for 15 minutes. The media was removed and the cells washed twice with cold DPBS (4 °C). The cells were resuspended in cold DPBS (4 °C), scraped and transferred to a separate 50 mL falcon tube. The cells were pelleted (1000g for 5 minutes at 4 °C) and suspended in 1mL of cold RIPA buffer containing PMSF (1mM) and cOmplete EDTA free protease inhibitor (1x) (Roche). The lysed cells were incubated on ice for 5–10 minutes and sonicated (35%, 5 x 5s with 30s rest). The lysate was then centrifuged at 15x1000g for 15 mins at 4 °C and the supernatant collected. The concentration of the cell lysate was measured by BCA assay and adjusted accordingly to equal concentration of 1 mg/mL. A control sample was removed from each plex (15 μL) and stored at –20 °C for later analysis.

*Streptavidin pull-down:*
Magnetic Streptavidin beads (NEB) were removed (250 μL per plex) and washed twice with RIPA (0.5 mL) (5 minutes incubation on a rotisserie). The beads were pelleted on a magnetic rack, diluted with the samples (1 mL) and incubated on a rotisserie at 4 °C overnight. The beads were pelleted on a magnetic rack, the supernatant removed, and a control sample from each plex (15 μL) and stored at –20 °C for later analysis. The beads were subsequently washed with 1 x RIPA (0.5 mL), 3 x 1% SDS in DPBS (0.5 mL), 3 x 1M NaCl in DPBS (0.5 mL), 3 x 10% EtOH in DPBS and 1 x RIPA (0.5 mL). The samples were incubated with each wash for 5 minutes prior to

pelleting. The beads were resuspended in RIPA buffer (300 µL) and transferred to a new 1.5 mL Lo-bind tube.

*Western Blot analysis:*

Following the final wash and transfer procedure for pull-down, the beads were pelleted on a magnetic rack and the supernatant removed. The beads were gently centrifuged to gather at the bottom of the tube and freshly prepared elution buffer (30 mM biotin, 6 M urea, 2 M thiourea, 2% SDS in DPBS, pH = 11.5) (24 µL) and 4x Laemlli buffer with BME (6 µL) was added with gentle mixing. The beads were heated to 95 °C for 15 minutes, pelleted on a magnetic rack, and the supernatant was removed while hot and beads discarded. The samples were cooled to room temperature and centrifuged. The samples (17 µL) were subsequently loaded onto a BioRad Criterion 4–20% tris-glycine gel, alongside all of the appropriate controls, and run in freshly prepared Tris running buffer (160V, 60 minutes). The gel was washed (3 x MiliQ water) and transferred via iBlot 2 to an NC membrane. The membrane was again washed (3 x MiliQ water) and blocked with Li-COR TBS Blocking Buffer for 1 hour at room temperature and then incubated with anti-α-tubulin (AB18251, Abcam) (1:1000) overnight in Pierce Protein-Free Blocking (1:2000) at 4 °C overnight. The membrane was washed 3 x TBST (5 mins per wash) and 5 x MiliQ water and resuspended in Pierce Protein-Free Blocking Buffer with Li-COR secondary antibody (Goat-anti-Rabbit 700) and rocked for 1 hour at room temperature (1:12,500). The membrane was washed 3 x TBST (5 mins per wash) and 5 x MiliQ water and imaged.

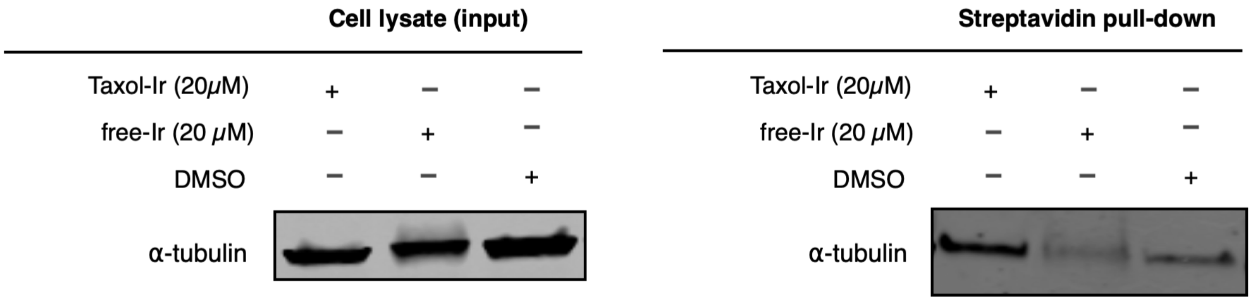

#### TMT-based chemoproteomic analysis

To MCF-7 cells in 10 clear 10 cm plate at 80% confluency in RPMI 1640 with **no** phenol red (Gibco) (4 mL) was added Paclitaxel-G2 (6) (20 μM) (5 plates, **A**) and Ir(dFCF$_3$ppy)$_2$(dMebpy)PF$_6$ (14) (2 μM) (5 plates, **B**). The plates were incubated at 37 °C for 3 hours and the media removed and replaced. Diazirine-alkyne (10) was added (250 μM) and the plates incubated at 37 °C for an additional 20 minutes. The plates were subsequently irradiated (without the lid) in the bioreactor at 450nM for 20 minutes. The plate was subsequently irradiated (without the lid) in the Merck bioreactor at 450nM for 15 minutes. The media was subsequently removed and the cells gently washed with cold DPBS (2 x 5 mL), the cells scraped (in 5 mL cold DPBS), combined, and pelleted (1000g for 5 minutes at 4 °C). The supernatant was removed and the cells suspended in 1mL of cold lysis buffer (1% SDS in 10mM HEPES, 150mM NaCl, 1.3mM $MgCl_2$) containing PMSF (1mM) and cOmplete EDTA free protease inhibitor (Roche). The lysed cells were incubated on ice and sonicated (35%, 4 x 5s with 30s rest). The lysate was then centrifuged at 15x1000g for 15 mins at 4 °C and the supernatant collected. The concentration of the cell lysate was measured by BCA assay (typically 3 mg/mL).

*CuAAC reaction:*

*Click-cocktail stock solution:* In a 0.5mL Lo-bind tube, 6.2 μL 500mM $CuSO_4$ was added to 62 μL 100 mM THPTA and vortexed. Subsequently, 15.5 μL 5 mM biotin-PEG7-azide (broadpharm) was added, followed by 15.5 μL of freshly prepared 1M sodium ascorbate (*Important: addition of reagents in that order*).

To the cell-lysate (1 mL) in a 1.5mL Lo-bind tube was added 32 μL of the *click-cocktail*. The resulting solution was vortexed and incubated on a rotisserie at room temperature for 1 hour and quenched by the addition of 5 μL 250 mM $Na_4$EDTA. The mixture was cooled to 0 °C, transferred to a 15mL tube, and diluted with 4.2 mL ice-cold acetone. The samples were precipitated at –20 °C overnight (3 hours was also found to be satisfactory), centrifuged at 4.5x1000g for 20 mins at 4 °C, and the supernatant removed. The pellet was fully resuspended in ice-cold methanol (1 mL) by sonication (2 s at 20%) and incubated at –20 °C for 30 minutes. After such time, the mixture was centrifuged at 4.5x1000g for 20 mins at 4 °C and the supernatant removed. The procedure was repeated. The pellet was allowed to air dry for 20 mins at room temperature and redissolved in 300 µL 1% SDS (1h at room temperature) and heated for 5 mins at 95 °C. The samples were cooled and diluted with 900 µL RIPA buffer. 250 µL of streptavidin magnetic beads (Thermo Fisher, cat. 88817) were added to Protein LoBind microcentrifuge tubes (Eppendorf, cat. 022431081) and washed 2x with 1 mL RIPA Buffer (Thermo Fisher, cat. 89900). Approximately 1.0 mg of cell lysate was added to the pre-washed streptavidin magnetic beads and incubated for 3 hours at room temperature. A magnetic rack was used to pellet the beads and remove the lysate supernatant. The beads were sequentially washed 3x with each of the following: 1 mL of 1% SDS, 1 mL of 1M NaCl, and 1 mL of 10% EtOH, all prepared in 1x DPBS and incubating for 5 min in between washes. A final wash was done with 1 mL RIPA Buffer. The beads were then resuspended in 30 µL of 4x Laemmli sample buffer (Boston BioProducts, cat. BP-110R) containing 20 mM DTT and 25 mM biotin. Beads were heated for 10 min at 95 °C and were then placed on the magnetic rack. The supernatant was transferred to a new Protein LoBind microcentrifuge tube and stored at -80 °C. Quantitative proteomic sample preparation and analysis was performed by IQ Proteomics (Cambridge, MA).

For LC-MS analysis at IQ Proteomics, mass spectra were acquired on an Orbitrap Fusion Lumos coupled to an EASY nanoLC-1000 (or nanoLC-1200) (Thermo Fisher) liquid chromatography system. Approximately 2 µg of peptides were loaded on a 75 µm capillary column packed in-house with Sepax GP-C18 resin (1.8 µm, 150 Å, Sepax) to a final length of 35 cm. Peptides were separated using a 110-minute linear gradient from 8% to 28% acetonitrile in 0.1% formic acid. The mass spectrometer was operated in a data dependent mode. The scan sequence began with FTMS1 spectra (resolution = 120,000; mass range of 350-1400 m/z; max injection time of 50 ms; AGC target of $1 \cdot 10^6$; dynamic exclusion for 60 seconds with a +/- 10 ppm window). The ten most intense precursor ions were selected for MS2 analysis via collisional-induced dissociation (CID) in the ion trap (normalized collision energy (NCE) = 35; max injection time = 100 ms; isolation window of 0.7 Da; AGC target of $1.5 \cdot 10^4$). Following MS2 acquisition, a synchronous-precursor-selection (SPS) MS3 method was enabled to select eight MS2 product ions for high energy collisional-induced dissociation (HCD) with analysis in the Orbitrap (NCE = 55; resolution =

50,000; max injection time = 86 ms; AGC target of $1.4 \cdot 10^5$; isolation window at 1.2 Da for +2 m/z, 1.0 Da for +3 m/z or 0.8 Da for +4 to +6 m/z). All mass spectra were converted to mzXML using a modified version of ReAdW.exe. MS/MS spectra were searched against a concatenated 2018 human Uniprot protein database containing common contaminants (forward + reverse sequences) using the SEQUEST algorithm[10]. Database search criteria are as follows: fully tryptic with two missed cleavages; a precursor mass tolerance of 50 ppm and a fragment ion tolerance of 1 Da; oxidation of methionine (15.9949 Da) was set as differential modifications. Static modifications were carboxyamidomethylation of cysteines (57.0214) and TMT on lysines and N-termini of peptides (229.1629). Peptide-spectrum matches were filtered using linear discriminant analysis[11] and adjusted to a 1% peptide false discovery rate (FDR)[12].

All bioinformatic analysis of LC-MS/MS data was performed in the R statistical computing environment [R Development Core Team, R (R foundation for statistical computing Vienna, Austria, 2011)]. Peptide level abundance data was used to identify the number of peptides corresponding to a protein in the experiment. Any protein with a single peptide quantification was removed to reduce the possibility that outliers would affect downstream proximal calls. Peptide level abundance data was then normalized to the summed total abundance for each sample separately. These totals were then averaged, and each normalized protein abundance value was multiplied by this average to rescale abundance data. Peptide level data was then merged to protein level data by taking the median of all peptides corresponding to a protein. Proteins were then filtered to remove any known contaminants identified from the database search and proteins which are known antibody contaminants (e.g. having IGK, IGK, or IGH present in the gene symbol and Immunoglobulin present in the Uniprot description). Data were then filtered to remove PRNP, a protein which is a known false positive consistently detected across almost all experiments. Protein abundances were $\log_2$ transformed and subjected to linear modeling analysis with Limma[13]. Limma utilizes an empirical Bayes approach that allows for a realistic distribution of biological variance with small sample sizes per group. This program further utilizes the full dataset to shrink the observed sample variances towards a pooled estimate. This borrowing of variance information across proteins allows for a more accurate estimate of true variance, and improved power to detect real differences between groups. For each protein, abundance data was fit to a linear model with the experimental group as the input variable using the lmFit function. The $\log_2$FC values were estimated and p-values calculated for significance. P-values were then corrected for multiple comparisons using the false discovery rate (FDR) method by Benjamini and Hochberg[14]. Volcano plots were generated in R with the ggplot2 library[15]. $Log_2FC$ and p-value estimates from Limma were subset to those reaching a specified $log_2FC$ cutoff. Proteins were colored based on whether they fell above or below the $log_2$-fold cutoff threshold and were statistically significant (FDR corrected p-value of $< 0.05$).

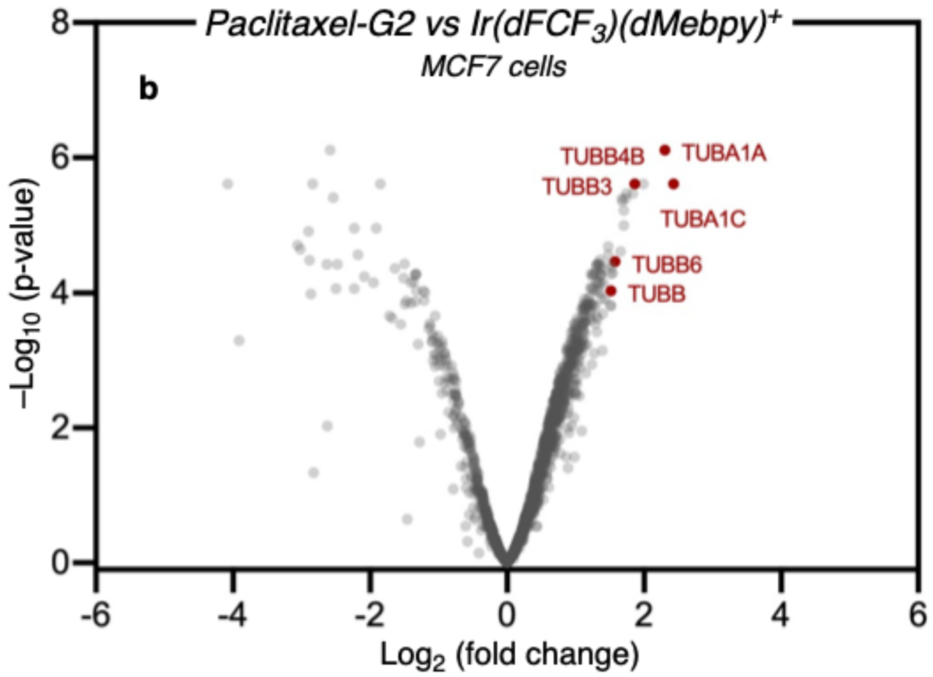

***See attached data files for full lists of proteins***

### Extracellular labelling of GPCR $A_{2a}$ using SCH58261-Ir

#### SCH58261 probes targeting Adora2a Structure Key

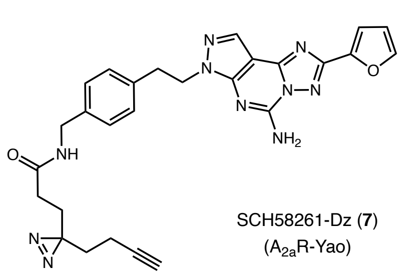

SCH58261-Dz (**7**)
($A_{2a}$R-Yao)

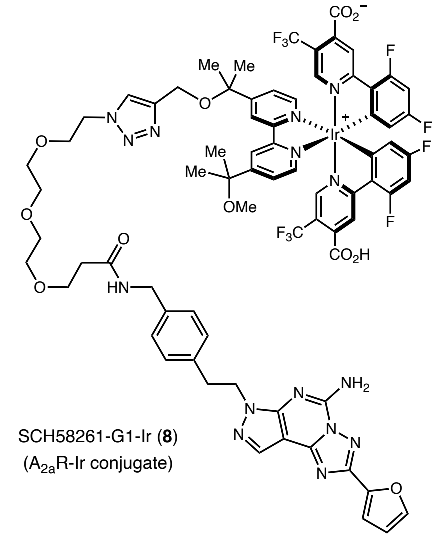

SCH58261-G1-Ir (**8**)
($A_{2a}$R-Ir conjugate)

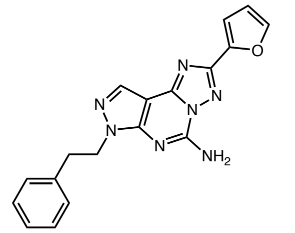

SCH58261-ligand (**22**)
($A_{2a}$R ligand)
***competitor***

SCH58261-Dz-biotin (**23**)
($A_{2a}$R-Yao-biotin)

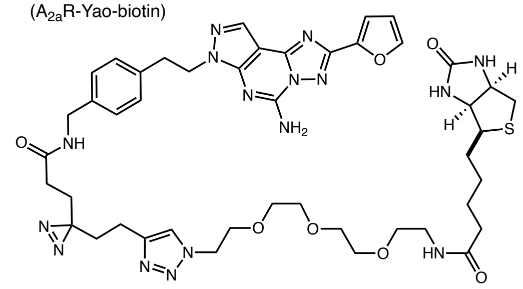

SCH58261-Dz-TAMRA (**24**)
($A_{2a}$R-Yao-TAMRA)

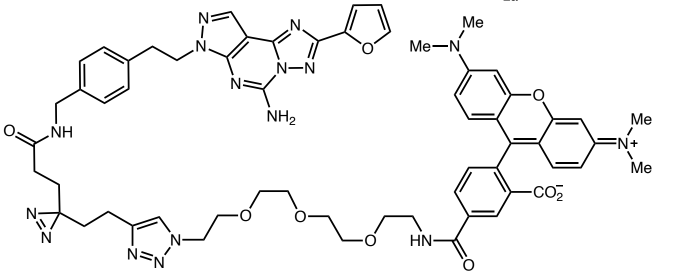

#### Synthesis of conjugates

##### 3-(3-(but-3-yn-1-yl)-3H-diazirin-3-yl)-N-(4-(2-hydroxyethyl)benzyl)propenamide

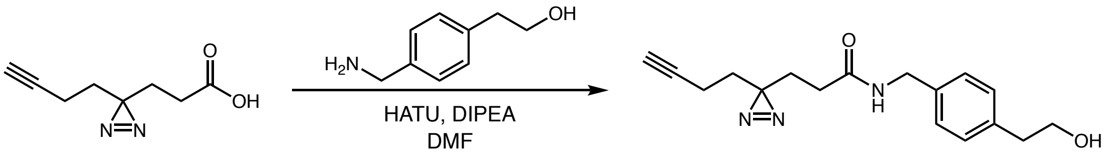

2-(4-(Aminomethyl)phenyl)ethan-1-ol (34 mg, 0.225 mmol) was dissolved in DMF (2 mL) and 3-(3-(but-3-yn-1-yl)-3H-diazirin-3-yl)propanoic acid (38.6 mg, 0.232 mmol) and HATU (124.6 mg, 0.328 mmol) were added, followed by DIPEA (0.200 mL, 1.145 mmol). The reaction mixture was stirred overnight at room temperature. This mixture was diluted with EtOAc, and washed twice with water. The organic layer was dried over $MgSO_4$, filtered and concentrated to afford a yellow oil. The residue was purified by column chromatography on silica gel (Isolute Flash Si; 12 g prepacked column, eluting with EtOAc/hexanes (EtOAc gradient from 0 to 80%)) to give the desired product **38** (49 mg, 0.164 mmol, 70.5 % yield) as a clear oil.

$^1$H-NMR (500 MHz, $CDCl_3$): δ 7.22 (m, 4H), 5.98 (s, 1H), 4.39 (d, 2H, $J$ = 5.6 Hz), 3.84 (t, 2H, $J$ = 6.5 Hz, 2H), 2.86 (t, 2H, $J$ = 6.5 Hz), 2.01 (m, 6H), 1.87 (t, 2H, $J$ = 7.7 Hz), 1.66 (t, 2H, $J$ = 7.4 Hz). LC-MS (ESI) m/z calcd. for $C_{17}H_{21}N_3O_2$ ([M+H]$^+$) 300.1667, found 300.23

**SCH58261-Dz-alkyne (7),**

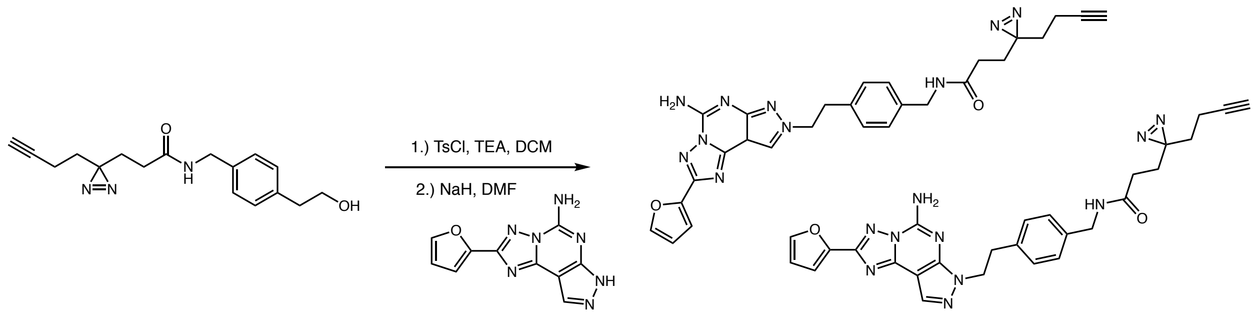

A solution of diazirine alcohol (0.049 g, 0.164 mmol) and triethylamine (0.050 mL, 0.359 mmol) in DCM (1.5 ml) was cooled to 0 °C. TsCl (0.053 g, 0.278 mmol) was added at 0 °C and stirred at this temperature for 3 h. The reaction was gradually warmed to room temperature and stirred for 20 h. The reaction mixture was diluted with DCM and washed with water (25 mL). The organic layer was dried over $Na_2SO_4$ and concentrated in vacuo to afford a clear oil, which was carried to the next step without further purification. To a solution of 2-(furan-2-yl)-7H-pyrazolo[4,3-e][1,2,4]triazolo[1,5-c]pyrimidin-5-amine (0.0378 g, 0.157 mmol) in anhydrous DMF (1 mL), NaH (12.0 mg, 0.300 mmol) was added under nitrogen. The reaction mixture was stirred for 20 min at room temperature, followed by addition of the freshly prepared tosylate from previous step (0.074 g, 0.164 mmol) dissolved in anhydrous DMF (1 mL). The reaction was stirred at room temperature for 48 h, then filtered and purified by HPLC (Gibson, Phenomenex C18, flow rate 20 mL/min, 8 min run, 20-100% MeCN in water with 0.05% TFA). The desired compounds eluted at 80% MeCN to give regioisomers N-(4-(2-(5-amino-2-(furan-2-yl)-8H-pyrazolo[4,3-e][1,2,4]triazolo[1,5-c]pyrimidin-8-yl)ethyl)benzyl)-3-(3-(but-3-yn-1-yl)-3H-diazirin-3-yl)propanamide, TFA- (14 mg, 0.022 mmol, 14.06 % yield) and N-(4-(2-(5-amino-2-(furan-2-yl)-7H-pyrazolo[4,3-e][1,2,4]triazolo[1,5-c]pyrimidin-7-yl)ethyl)benzyl)-3-(3-(but-3-yn-1-yl)-3H-diazirin-3-yl)propanamide, TFA- (40 mg, 0.063 mmol, 40.2 % yield). Both regioisomers were obtained as white solids.

**SCH58261-Dz-alkyne (7)**

$^{1}$H-NMR (500 MHz DMSO-d6): δ 8.31 (t, 1H, *J* = 5.8 Hz), 8.18 (s, 1H), 7.96 (s, 1H), 7.24 (d, 1H, *J* = 3.3 Hz), 7.15 (s, 4H), 6.75 (d, 1H, *J* = 1.7 Hz), 4.48 (t, 2H, *J* = 7.4 Hz), 4.20 (d, 2H, *J* = 5.8 Hz), 3.18 (t, 2H, *J* = 7.3 Hz), 2.84 (t, 1H, *J* = 2.4 Hz), 2.01 – 1.93 (m, 4H), 1.67 (t, 2H, *J* = 7.6 Hz), 1.57 (t, 2H, *J* = 7.4 Hz). $^{13}$C-NMR (126 MHz, DMSO): δ 171.0, 155.8, 149.1, 148.8, 146.7, 145.9, 145.5, 137.9, 137.2, 131.7, 129.0, 127.7, 112.7, 112.6, 96.1, 83.6, 72.2, 48.2, 42.2, 34.9, 31.9, 29.8, 28.7, 28.5, 13.1. HRMS (ESI-TOF): m/z calcd. for $C_{27}H_{26}N_{10}O_2$ ([M+H]$^{+}$) 523.2318, found 523.2330

**Regioisomer (7a)**

$^{1}$H-NMR (500 MHz, DMSO-d6): δ 8.48 (s, 1H), 8.31 (t, *J* = 5.7 Hz, 1H), 7.94 (s, 1H), 7.64 (s, 2H), 7.19 (d, 1H, *J* = 3.4 Hz), 7.15 (d, 5H, *J* = 4.4 Hz), 6.74 (dd, 1H, *J* = 3.3, 1.7 Hz), 4.53 (t, 2H, *J* = 7.1 Hz), 4.23 – 4.18 (m, 3H), 3.57 (t, 2H, *J* = 7.2 Hz), 3.20 (t, 2H, *J* = 7.1 Hz), 2.84 (m, 1H), 2.69 (t, 1H, *J* = 7.1 Hz), 2.02 – 1.91 (m, 6H), 1.67 (q, 3H, *J* = 7.4 Hz), 1.60 – 1.53 (m, 3H). LC-MS (ESI): m/z calcd. for $C_{27}H_{26}N_{10}O_2$ ([M+H]$^{+}$) 523.23, found 523.29

**3-(2-(2-(2-azidoethoxy)ethoxy)ethoxy)-N-(4-(2-hydroxyethyl)benzyl)propenamide**

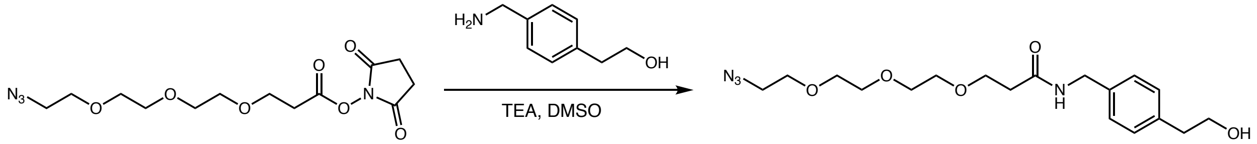

2-(4-(aminomethyl)phenyl)ethan-1-ol (34.8 mg, 0.230 mmol) was dissolved in DMSO (1.5 mL) and 2,5-dioxopyrrolidin-1-yl 3-(2-(2-(2-azidoethoxy)ethoxy)ethoxy)propanoate (87.2 mg, 0.253 mmol) was added, followed by triethylamine (0.070 mL, 0.502 mmol). The mixture was stirred at room temperature overnight, then filtered and purified by HPLC (Gibson, Phenomenex C18, flow rate 20 mL/min, 8 min run, 15-100% MeCN in water with 0.05% TFA). The desired compound eluted at 80% MeCN to give 3-(2-(2-(2-azidoethoxy)ethoxy)ethoxy)-N-(4-(2-hydroxyethyl)benzyl)propanamide (70 mg, 0.184 mmol, 80 % yield) as a white solid.

$^{1}$H-NMR (500 MHz, $CD_3OD$): δ 7.22 (q, 4H, *J* = 8.1 Hz), 4.37 (s, 2H), 3.78 – 3.72 (m, 4H), 3.66 – 3.60 (m, 10H), 3.36 (t, 2H, *J* = 7.0 Hz), 2.82 (t, 2H, *J* = 7.1 Hz), 2.51 (t, 2H, *J* = 6.1 Hz).
LC-MS (ESI-TOF): m/z calcd. for $C_{18}H_{28}N_4O_5$ ($[M+H]^+$) 381.2093, found 381.05

**SCH58261-PEG3-azide**

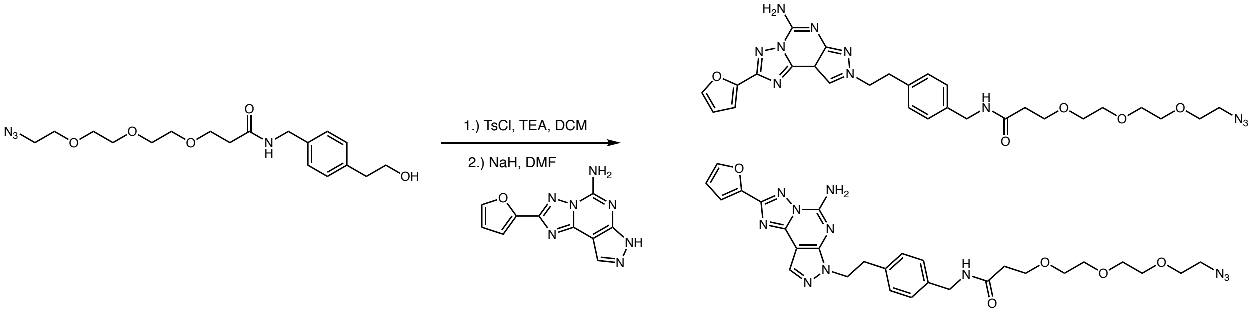

A solution of compound PEG azido alcohol (0.0396 g, 0.104 mmol) and triethylamine (0.035 mL, 0.251 mmol) in DCM (1.5 mL) and was cooled to 0 °C. Tosyl-Cl (0.0317 g, 0.166 mmol) was added at 0 °C and stirred for 1 h at this temperature. The reaction was gradually warmed to room temperature and stirred overnight, then diluted with DCM and washed with water (25 mL x 2). The organic layer was dried over $Na_2SO_4$ and concentrated in vacuo. This material was carried to the next step without further purification.

To a solution of 2-(furan-2-yl)-7H-pyrazolo[4,3-e][1,2,4]triazolo[1,5-c]pyrimidin-5-amine (26.2 mg, 109 µmol) in anhydrous DMF (1.5 mL) under nitrogen, NaH (8.7 mg, 218 µmol) was added. The reaction mixture was stirred for 20 min at room temperature, followed by addition of tosylate (53.5 mg, 0,100 µmol) dissolved in anhydrous DMF (1 mL). The reaction was stirred at room temperature for 24 h, then filtered and purified by HPLC (Gibson, Phenomenex C18, flow rate 20 mL/min, 8 min run, 20-100% MeCN in water with 0.05% TFA). The desired compounds eluted at 80% MeCN to give regioisomers N-(4-(2-(5-amino-2-(furan-2-yl)-8H-pyrazolo[4,3-e][1,2,4]triazolo[1,5-c]pyrimidin-8-yl)ethyl)benzyl)-3-(2-(2-(2-

azidoethoxy)ethoxy)ethoxy)propanamide, TFA- (3.5 mg, 4.88 µmol, 4.50 % yield) and N-(4-(2-(5-amino-2-(furan-2-yl)-7H-pyrazolo[4,3-e][1,2,4]triazolo[1,5-c]pyrimidin-7-yl)ethyl)benzyl)-3-(2-(2-(2-azidoethoxy)ethoxy)ethoxy)propanamide, TFA- (8.4 mg, 11.72 µmol, 10.79 % yield). Both regioisomers were obtained as white solids.

**Isomer**

$^{1}$H-NMR (500 MHz, $CH_3OH$-d4): δ 8.46 (s, 1H), 8.30 (t, 1H, *J* = 5.9 Hz), 7.93 (s, 1H), 7.20 – 7.09 (m, 6H), 6.72 (d, 1H, *J* = 1.7 Hz), 4.52 (t, 2H, *J* = 7.0 Hz), 4.22 (d, 2H, *J* = 4.7 Hz), 3.60 (dt, 5H, *J* = 13.3, 5.6 Hz), 3.54 – 3.44 (m, 10H), 3.39 – 3.29 (m, 5H), 3.16 (s, 10H), 2.35 (t, 2H, *J* = 6.3 Hz). LC-MS (ESI): m/z calcd. for $C_{28}H_{33}N_{11}O_5$ ($[M+H]^+$) 604.27, found 603.91

**SCH58261-PEG3-azide**

$^{1}$H-NMR (500 MHz, DMSO-d6): δ 8.30 (t, 1H, *J* = 5.9 Hz), 8.17 (s, 1H), 7.97 – 7.93 (m, 1H), 7.24 (d, 1H, *J* = 3.4 Hz), 7.14 (s, 4H), 6.75 (dd, 1H, *J* = 3.4, 1.8 Hz), 4.48 (t, 2H, *J* = 7.4 Hz), 4.22 (d, 2H, *J* = 5.9 Hz), 3.66 – 3.45 (m, 15H), 3.40 – 3.37 (m, 2H), 3.18 (t, 2H, *J* = 7.3 Hz), 2.37 (t, 2H, *J* = 6.4 Hz). $^{13}$C-NMR (126 MHz, DMSO-d6): δ 170.5, 155.8, 149.1, 148.8, 146.7, 145.9, 145.5, 138.0, 137.1, 131.7, 128.9, 128.7, 127.8, 127.6, 112.7, 112.6, 96.1, 70.2, 70.1, 70.1, 69.9, 69.7, 67.3, 50.4, 48.2, 42.1, 36.5, 34.9. HRMS (ESI-TOF): m/z calcd. for $C_{28}H_{33}N_{11}O_5$ ($[M+H]^+$) 604.2744, found 604.2762

**SCH58261-Dz-biotin (23),**

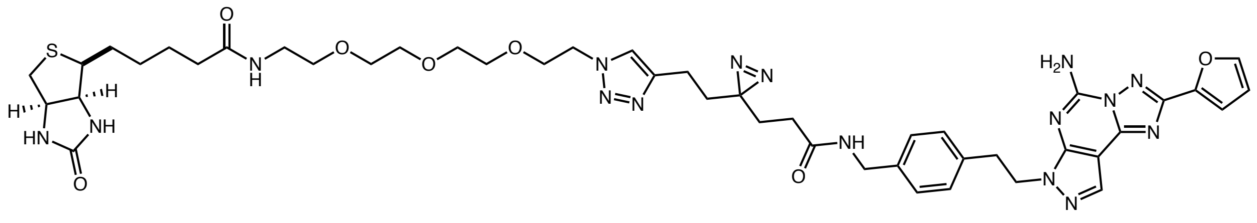

Alkyne fragment (0.012 mmol, 1 eq) and azide fragment (0.012 mmol, 1 eq) were dissolved in DMSO (0.9 mL). Copper (II) sulfate pentahydrate (0.030 mmol, 2.5 eq) was dissolved in water (0.1 mL) and added to reaction mixture. Reaction mixture was stirred for 5 min followed by addition of sodium ascorbate (0.114 mmol, 10 eq) and stirred at room temperature overnight. Reaction mixture was filtered and purified by HPLC, Gibson, Phenomenex C18, flow rate 20 mL/min, 8 min run, 30-100% ACN in water with 0.05% TFA.

$^1$H-NMR (500 MHz, DMSO-d6): δ 8.31 (t, 1H, *J* = 5.8 Hz), 8.17 (s, 1H), 7.97 – 7.94 (m, 1H), 7.85 (d, 2H, *J* = 6.9 Hz), 7.24 (d, 1H, *J* = 3.4 Hz), 7.14 (s, 4H), 6.75 (dd, 1H, *J* = 3.3, 1.7 Hz), 6.41 (d, 2H, *J* = 28.8 Hz), 4.47 (q, 4H, *J* = 6.3, 5.1 Hz), 4.31 (dd, 1H, *J* = 7.6, 5.0 Hz), 4.21 (d, 2H, *J* = 5.8 Hz), 4.13 (dd, 1H, *J* = 7.7, 4.4 Hz), 3.78 (t, 2H, *J* = 5.2 Hz), 3.52 – 3.44 (m, 8H), 3.38 (t, 2H, *J* = 5.9 Hz), 3.22 – 3.12 (m, 4H), 3.11 – 3.06 (m, 1H), 2.82 (dd, 1H, *J* = 12.4, 5.1 Hz), 2.60 – 2.55 (m, 1H), 2.44 – 2.38 (m, 2H), 2.06 (t, 2H, *J* = 7.4 Hz), 1.96 (t, 2H, *J* = 7.7 Hz), 1.73 (dd, 2H, *J* = 9.0, 7.1 Hz), 1.66 (t, 2H, *J* = 7.7 Hz), 1.63 – 1.56 (m, 1H), 1.53 – 1.41 (m, 3H), 1.38 – 1.16 (m, 3H). $^{13}$C-NMR (126 MHz, DMSO): δ 172.5, 171.0, 163.1, 158.6, 158.4, 155.8, 149.1, 148.8, 146.7, 145.9, 145.6, 145.5, 137.9, 137.2, 131.7, 129.0, 127.7, 122.8, 112.7, 112.6, 96.1, 70.1, 70.0, 70.0, 70.0, 69.6, 69.2, 61.4, 59.6, 55.8, 49.7, 48.2, 42.2, 38.8, 35.5, 34.9, 32.4, 29.8, 28.9, 28.6, 28.6, 28.5, 25.7, 20.0. HRMS (ESI-TOF): m/z calcd. for $C_{45}H_{58}N_{16}O_7S$ ([M+H] $^+$) 967.4473, found 967.4421

**SCH58261-PEG3-TAMRA (24),**

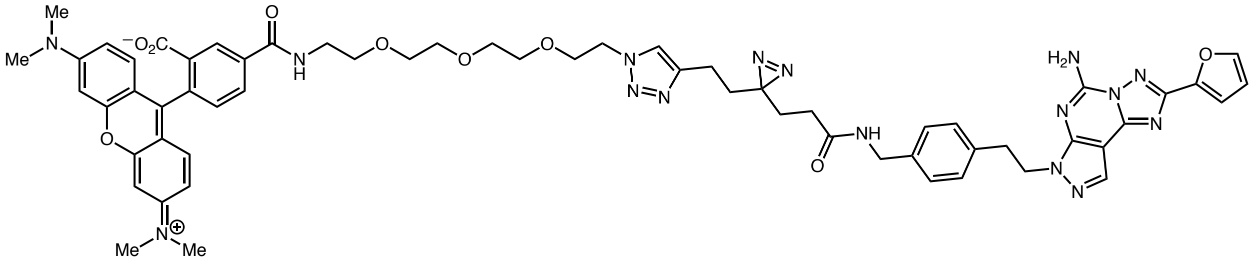

Alkyne fragment (0.012 mmol, 1 eq) and azide fragment (0.012 mmol, 1 eq) were dissolved in DMSO (0.9 mL). Copper (II) sulfate pentahydrate (0.030 mmol, 2.5 eq) was dissolved in water (0.1 mL) and added to reaction mixture. Reaction mixture was stirred for 5 min followed by addition of sodium ascorbate (0.114 mmol, 10 eq) and stirred at room temperature overnight. Reaction mixture was filtered and purified by HPLC, Gibson, Phenomenex C18, flow rate 20 mL/min, 8 min run, 30-100% ACN in water with 0.05% TFA.

$^1$H-NMR (500 MHz, $CD_3OD$): δ 8.76 (d, 1H, *J* = 1.8 Hz), 8.30 (dd, 1H, *J* = 7.9, 1.8 Hz), 8.05 (s, 1H), 7.80 (s, 1H), 7.76 (d, *J* = 1.6 Hz, 1H), 7.60 (d, *J* = 7.9 Hz, 1H), 7.20 (d, 1H, *J* = 3.4 Hz), 7.15 (d, 2H, *J* = 8.0 Hz), 7.09 (d, 2H, *J* = 8.1 Hz), 7.01 (d, 2H, *J* = 9.5 Hz), 6.90 (dd, 2H, *J* = 9.5, 2.4 Hz), 6.80 (d, 2H, *J* = 2.4 Hz), 6.66 (dd, 1H, *J* = 3.4, 1.7 Hz), 4.50 (t, 2H), 4.44 (t, 2H, *J* = 7.2 Hz), 4.26 (s, 2H), 3.87 – 3.83 (m, 2H), 3.75 (t, 2H, *J* = 5.2 Hz), 3.72 – 3.67 (m, 4H), 3.66 – 3.59 (m, 6H), 3.26 (s, 12H), 3.16 (t, 2H, *J* = 7.2 Hz), 2.45 (t, 2H), 2.00 (t, 2H, *J* = 7.6 Hz), 1.72 (dd, 2H, *J* = 8.7, 7.1 Hz), 1.67 (t, 2H, *J* = 7.6 Hz). $^{13}$C-NMR (126 MHz, $CD_3OD$): δ 172.6, 166.7, 165.9, 160.1, 159.8, 158.9, 157.3, 157.3, 155.3, 148.7, 148.6, 146.3, 145.9, 145.2, 144.7, 137.0, 136.9, 136.6, 136.1, 131.3, 131.1, 131.0, 130.6, 130.3, 129.8, 128.6, 127.4, 122.9, 114.0, 113.1, 111.9, 111.6, 95.9, 95.7, 70.2, 70.1, 69.9, 69.9, 69.1, 68.9, 49.9, 42.4, 39.9, 39.5, 34.7, 32.0, 29.5, 28.4, 27.6, 19.3. HRMS (ESI-TOF): m/z calcd. for $C_{60}H_{64}N_{16}O_9$ ($[M+H]^+$) 1152.5042; for $C_{60}H_{64}N_{16}O_9$ ($[M+H]^{+2}$) 577.2599, found 577.2620

**SCH58261-G1-Ir (8)**

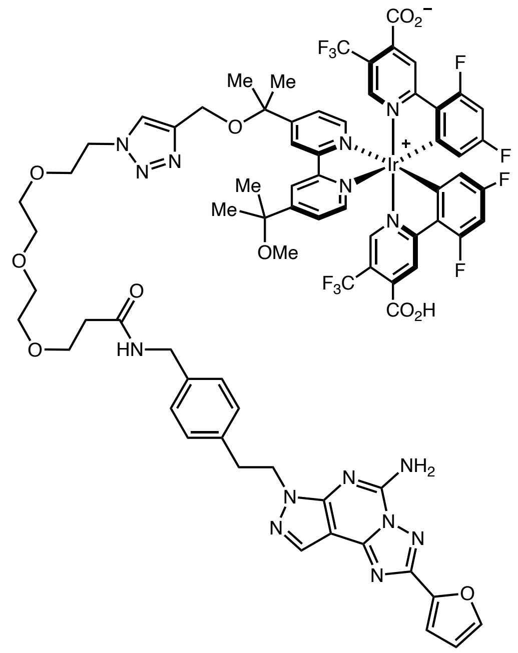

Alkyne fragment (0.012 mmol, 1 eq) and azide fragment (0.012 mmol, 1 eq) were dissolved in DMSO (0.9 mL). Copper (II) sulfate pentahydrate (0.030 mmol, 2.5 eq) was dissolved in water (0.1 mL) and added to reaction mixture. Reaction mixture was stirred for 5 min followed by addition of sodium ascorbate (0.114 mmol, 10 eq) and stirred at room temperature overnight. Reaction mixture was filtered and purified by HPLC, Gibson, Phenomenex C18, flow rate 20 mL/min, 8 min run, 30-100% ACN in water with 0.05% TFA.

$^{1}$H-NMR (500 MHz, DMSO-d6): δ 8.99 (s, 1H), 8.87 (s, 1H), 8.55 (s, 2H), 8.27 (t, 1H, J = 5.8 Hz), 8.13 (s, 1H), 8.12 (s, 1H), 8.07 (bs, 2H), 7.99 (d, 1H, J = 5.8 Hz), 7.98 (d, 1H, J = 5.8 Hz), 7.94 (d, 1H, J = 1.0 Hz), 7.90 (dd, 1H, J = 5.8, 1.4 Hz), 7.82 (dd, 1H, J = 5.8, 1.5 Hz), 7.44 (s, 1H), 7.41 (s, 1H), 7.21 (d, 1H, J = 3.4 Hz), 7.11 (m, 6H), 6.72 (d, 1H, J = 3.4, 1.8 Hz), 5.95 (d, 2H, J = 7.2 Hz), 4.50 (t, 2H, J = 5.1 Hz), 4.46 (t, 2H, J = 7.4 Hz), 4.41 (s, 2H), 4.17 (d, 2H, J = 5.9 Hz), 3.80 (t, 2H, J = 5.2 Hz), 3.57 (t, 2H, J = 6.4Hz), 3.50 (m, 2H), 3.47-3.40 (m, 6H), 3.15 (t, 2H, J = 7.3 Hz), 3.11 (s, 3H), 2.33 (t, 2H, J = 6.4 Hz), 1.65 (s, 3H), 1.63 (s, 3H), 1.61 (s, 3H), 1.59 (s, 3H)
$^{13}$C-NMR (126 MHz, DMSO-d6): δ 169.9, 167.5 (m), 164.5, 164.2 (dd, J = 262.5, 12.5 Hz), 161.9 (dd, J = 262.5, 12.5 Hz), 160.7, 160.7, 157.5, 155.8, 155.7, 155.4 (m), 155.3, 151.1, 151.0, 148.6, 148.3, 146.4 (m), 146.2, 145.4, 145.1, 143.8, 142.0, 137.5, 136.6, 131.2, 128.4, 127.2, 126.1,

124.2, 122.9, 122.8, 122.5, 122.3, 121.4 (d, J = 32.5 Hz), 121.3 (d, J = 271.2 Hz), 121.3 (d, J = 270.0 Hz), 114.6, 114.4, 112.2 (d, J = 16.6 Hz), 99.8 (t, J = 26.5 Hz), 95.7, 77.1, 76.3, 69.6, 69.5, 69.4, 68.6, 66.8, 56.9, 50.3, 49.3, 47.7, 41.6, 40.4, 36.0, 34.4, 27.6, 27.1, 26.6, 26.4

$^{19}$F-NMR (471 MHz, DMSO-d6): δ -59.09, -74.51, -102.27, -102.28, -102.29, -106.63, -106.66, -106.68. HRMS (ESI-TOF): m/z calcd. for $C_{74}H_{66}F_{10}IrN_{15}O_{11}$ ($[M+H]^{+}$) 1722.4590, found 1722.4479; for $C_{74}H_{66}F_{10}IrN_{15}O_{11}$ ($[M+H]^{+2}$) 861.7334, found 861.7294

#### Cell culture

HEK293-A2a cells were grown and expanded in EMEM (Sigma, cat. M6199-500) with 10% FBS, supplemented with P/S and geneticin (400 ug/mL). PC-12 (ATCC CRL-1721) were grown and expanded in RPMI-1640 (Gibco, cat. 11875) media with 5% FBS, 10% heat-inactivated horse serum. All cells were maintained at 37°C and 5% $CO_2$ atmosphere in T175 flasks (Nunc, cat. 159910).

#### Binding and cell functional assays for adenosine receptors

##### $A_2$ Adenosine receptor cAMP protocol

The ability of compounds to antagonize human $A_{2A}$ and $A_{2B}$ adenosine receptors was determined using a kit to measure changes in intracellular cyclic AMP levels (LANCE cAMP 384 Kit, Perkin Elmer, Cat. No. AD0264). HEK293 cells recombinantly expressing either human $A_{2A}$ (Perkin Elmer, Cat. No. ES011CV) or $A_{2B}$ (Perkin Elmer, Cat. No. ES013CF) receptors, previously frozen in Recovery Medium (Life Technologies, Cat. No. 12648-010), were thawed and diluted into stimulation buffer (HBSS (Hyclone SH 30268.01), 5 mM HEPES (Gibco 15630-106), 200 nM rolipram (Tocris, Cat. No. 0905), and 1.5 %(v/v) BSA stabilizer (kit component). The cell suspension was centrifuged at 200 x g for 10 min and then resuspended in stimulation buffer, supplemented with a 1:10,000 dilution of Alexa Fluor 647 anti-cAMP antibody, to a density of 6.0 x$10^5$ cells/mL. A Labcyte Echo 550 acoustic dispenser was used to transfer up to 25 nL of test compound dissolved in DMSO into the wells of a dry Optiplate-384 plate (Perkin Elmer, Cat. No. 6008289). All subsequent liquid additions were performed using a multichannel pipettor. Next, 5 µL of the cell suspension was added to the wells of the Optiplate-384 and incubated for 30 min at

37 °C and 5% $CO_2$ in a humidified environment. After this time 5 µL of either 300 nM or 600 nM adenosine (Sigma Cat. No. A9251) for $A_{2A}$ and $A_{2B}$ respectively was added and incubated for 30 minutes at 37 °C and 5% $CO_2$ in a humidified environment. At this time, detection mix was prepared by combining the LANCE Eu-W8044 labeled streptavidin and Biotin-cAMP in detection buffer according to the manufacturers protocol. 10 µL of the detection mix was added to each well of the Optiplate-384, which was covered with a plate seal and incubated under ambient conditions for 2 h prior to reading the plate using an Envision (Perkin Elmer, Waltham, MA) multimode plate reader. Data was normalized by defining minimal effect as stimulation in the presence of 0.25% (v/v) DMSO and maximal effect as stimulation in the presence of 1 µM ZM241385 (Cayman, Cat. No. 1036). Curve fitting of the percent effect data versus the log of compound concentration used a 4-parameter concentration response curve fitting algorithm to calculate $EC_{50}$ values. Compound concentrations tested were typically from 10,000, 3,333, 1,111, 370.4, 123.4, 41.2, 13.7, 4.6, 1.5 and 0.5 nM with 0.25% residual DMSO. Compounds were run as two biological replicates with reported values as average EC50.

**$A_{2A}$ Adenosine receptor binding protocol**

The ability of compounds to bind to the human $A_{2A}$ Adenosine receptor was determined by competitive radioligand binding using [$^3$H] SCH58261. Membranes made from HEK293 cells recombinantly expressing the human $A_{2A}$ receptors (Perkin Elmer, Cat. No. ES011CV) or $A_{2B}$ receptors were thawed and diluted in assay buffer (50 mM Tris-HCl pH = 7.4, 10 mM $MgCl_2$, 0.005% (v/v) Tween20) to a concentration of 6.67 µg/mL. To create a homogenous suspension, this preparation was slowly passed through a 27-gauge syringe needle. To individual wells of a fresh 1.2 mL 96-well polypropylene plate 2 µL of test compound or control samples were combined with 148 µL of the membrane suspension. Following a 30 min incubation at room temperature, the assay incubation was started by the addition of 50 µL of 4 nM [$^3$H] SCH58261, also prepared in assay buffer. After a further 60 min incubation at room temperature with shaking, the assay was terminated by rapid filtration through Unifilter GF/C PEI coated plates (Perkin Elmer, Cat. No. 6005277) using a Filtermate Harvester (Perkin Elmer). The filter plates were washed 3 times with approximately 1 mL of cold wash buffer (50 mM Tris-HCl pH = 7.4, 150 mM NaCl) and allowed to dry in a 55 °C oven for 3 h. After sealing the bottom of the plate, 50 µL Ultima Gold (Perkin Elmer, Cat. No. 6013329), scintillant was added to each well and the plate was sealed with a TopSeal-A PLUS clear plate seal. The amount of radioactivity remaining in each well was determined using a Topcount NXT scintillation counter (Perkin Elmer). Data was normalized by defining minimal effect as [$^3$H] SCH-58261 binding in the presence of 1% (v/v) DMSO and maximal effect as binding in the presence of 1 μM ZM241385 (Cayman, Cat. No. 1036). Curve fitting of the percent effect data versus the log of compound concentration used a 4-parameter concentration response curve fitting algorithm to calculate $EC_{50}$ values. Compound concentrations tested were typically from 10,000, 3,333, 1,111, 370.4, 123.4, 41.2, 13.7, 4.6, 1.5 and 0.5 nM with 0.25% residual DMSO.

**$A_1$ Adenosine receptor binding protocol**

The ability of compounds to bind to the human $A_1$ Adenosine receptor was determined by competitive radioligand binding using [$^3$H] Cyclopentyl-1,3-dipropylxanthine (CPA; Perkin Elmer, Cat. No. NET-974). Membranes made from CHO cells recombinantly expressing the human $A_1$ receptor (Perkin Elmer, Cat. No. ES-010-M400UA) were thawed and diluted in assay buffer (25 mM HEPES pH = 7.4, 5 mM $MgCl_2$, 1 mM $CaCl_2$, 100 mM NaCl, 0.005% (v/v) Tween20) to a concentration of 20 μg/mL. To create a homogenous suspension, this preparation was slowly passed through a 27-gauge syringe needle. To individual wells of a fresh 1.2 mL 96-well polypropylene plate, 2 μL of test compound or control samples were combined with 148 μL of the membrane suspension. Following a 30 min incubation at room temperature, the assay incubation was started by the addition of 50 μL of 4 nM [$^3$H] CPA, also prepared in assay buffer. After a further 60 min incubation at room temperature with shaking, the assay was terminated by rapid filtration through Unifilter GF/B PEI coated plates (Perkin Elmer, Cat. No. 6005277) using a Filtermate Harvester (Perkin Elmer). The filter plates were washed 3 times with approx. 1 mL of cold wash buffer (25 mM HEPES pH = 7.4, 5 mM $MgCl_2$, 1 mM $CaCl_2$, 100 mM NaCl) and the plates allowed to dry in a 55 °C oven for 3 h. After sealing the bottom of the plate 50 μL Ultima Gold (Perkin Elmer, Cat. No. 6013329), scintillant was added to each well and the plate sealed with a TopSeal-A PLUS clear plate seal. The amount of radioactivity remaining in each well was determined using a Topcount NXT scintillation counter (Perkin Elmer). Data was normalized by defining minimal effect as [$^3$H] CPA binding in the presence of 1% (v/v) DMSO and maximal effect as binding in the presence of 1 μM SLV320 (Cayman, Cat. No. 27666). Curve fitting of the percent effect data versus the log of compound concentration used a 4-parameter concentration response curve fitting algorithm to calculate $EC_{50}$ values. Compound concentrations tested were typically from 10,000, 3,333, 1,111, 370.4, 123.4, 41.2, 13.7, 4.6, 1.5 and 0.5 nM with 0.25% residual DMSO. Compounds were run as two biological replicates with reported values as average EC50.

| | Receptor binding | | Receptor function | |
|---|---|---|---|---|
| Annotation | hA2a (nM) | hA1 (nM) | hA2a (nM) | hA2b (nM) |
| SCH58261-Dz-TAMRA (24) | 93.7 | > 10,000 | 899 | > 10,000 |
| SCH58261-G1-Ir (8) | 643.4 | > 10,000 | 1,000 | > 10,000 |
| SCH58261-Dz-biotin (23) | 5.1 | > 10,000 | 7.3 | 8490 |
| SCH58261-azide | 0.8 | 1,119 | 24.3 | 7,566 |
| SCH58261-Dz (7) | 0.8 | 1.8 | 12.5 | 27.9 |
| SCH58261-ligand (22) | 3.8 | 383.7 | 7.9 | 411.3 |
| SCH58261-Dz-isomer (7a) | 4.1 | 5.7 | 160.4 | 279.2 |
| SCH58261-azide-isomer | 46.9 | 361.8 | 470.1 | 3915 |

#### General procedure for cell-membrane µMap

Growth media was removed, and cells were resuspended in enzyme-free dissociation buffer (Gibco, cat. 13150-016). Dissociated cells were consolidated and centrifuged at room temperature at 1,000 x g for 5 min. Cells were resuspended in cold (4 °C) 1X DPBS (Gibco, cat. 14190-136) and aliquoted in Eppendorf microcentrifuge tubes at 1-3 x $10^6$ cells/mL. Cells were treated with 100 µM unlabeled competitor 5 min prior to incubation with 1 µM chemical probes for 30 min at 4 °C. Prior to irradiation, PCAT samples were resuspended in 500 µL of 250 µM Diaz-PEG3-Biotin (9) in DPBS. Alkyldiazirine samples resuspended in 0.5 mL 1x DPBS were kept on ice and irradiated for 30 min at 365 nm in photocrosslinker box (UVP Crosslinker, Analytikjena,100% intensity). PCAT samples were irradiated for 10 min at 450 nm in a biophotoreactor (BPR200, Efficiency Aggregators, Fisher, cat. NC1558343). Cells were washed with cold 1X DPBS (1x; 1,000 x g for 4 min) and split into two tubes to evaluate two lysis conditions (regular vs. membrane enrichment). Eppendorf tubes were centrifuged at 10,000 x g for 4 min at 4 °C. 150 µL of lysis buffer (1% Triton-100, 150 mM NaCl, 10 mM Tris, 0.5% deoxycholic acid, 5 mM EDTA, pH = 7.4) with 1X protease inhibitor cocktail (Promega, cat. G6521, 50X in DMSO). Cells were lysed on ice for 30 min, followed by centrifuging at 16,000 x g for 30 min at 4 °C to clarify supernatant. The supernatant was separated from insoluble material to be flash frozen and stored at -80 °C until western blot analysis.

##### Membrane enrichment protocol for western blot (WB) samples

After the final wash, the cells were pelleted by centrifugation (4 min at 1,000 x g, 4 °C) and resuspended in 0.3 mL of membrane permeabilization buffer (MEM-PER Plus Membrane Fractionation Kit, Thermo Fisher Scientific, cat. 89842) containing 1x protease inhibitors (Promega, cat. G6521, 50X in DMSO) and incubated for 30 min at 4 °C. The samples were then spun at 16,000 x g for 30 min at 4 °C. The supernatant containing cytosolic proteins was removed, and the pellet was resuspended in 150 µL lysis buffer (RIPA buffer, Thermo Fisher Scientific, cat. 89900) containing 1% SDS and 1X protease inhibitors. The samples were sonicated to break up the membrane pellet (Fisher, model FB50, tip CL-18; 1x 5 s at power level 6) and then heated for

5 min at 95 °C. The protein concentration was measured by Bradford (see below) and the samples were processed for streptavidin bead enrichment.

**Copper click reaction conditions**

Lysate samples (200 μL as total volume) were treated with 2 μL biotin-azide of 10 mM stock (Broadpharm, cat. BP-20701; 100 μM final concentration), 2 μL of premixed 2:1 BTTAA:CuSO4 (equal volumes of 50 mM CuSO4 and 100 mM BTTAA stocks; final concentration as 250 μM CuSO4 and 500 μM BTTAA), and 2 μL of freshly prepared sodium ascorbate solution (250 mM stock, 2.5 mM final concentration). Reactions were allowed to proceed for 2 h at room temperature while rotating. EDTA at 5 mM final concentration was added to quench reactions. Samples were carried forward for protein quantification and western blot analysis as described below.

**Protein quantification and sample preparation for Western Blot analysis**

Protein concentration was assessed with Bradford assay using Coomassie reagent (Pierce, cat. 23200) to read sample absorbance at 595 nm with a SpectraMax M5 plate reader. For western blot analysis, samples were diluted with water, treated with 10X reducing agent (Invitrogen, cat. NP0004; 1X final concentration) and 4X NuPage loading buffer (Invitrogen, cat. NP0007; 1X final concentration). All samples incubated for 10 min at 95 °C. A short spin was performed prior to loading onto gel. Gel electrophoresis was performed using the XCell SureLock Mini-Cell Electrophoresis System (Thermo Fisher, cat. EI0001). Samples were loaded onto a 12 % bistris NuPage gel (under reducing conditions) and ran at 190 V for 50 min to resolve proteins using 1X MES SDS buffer (Invitrogen, cat. NP0002) and molecular weight ladder SeeBlue Plus2 pre-stained protein standard (Invitrogen, cat. LC5925). Proteins were blotted onto a PVDF membrane (Invitrogen, cat. IB24001) using iBlot2 gel transfer device (Thermo Fisher, cat. IB21001, IB23001) with settings as 23V for 7 min. The membrane was blocked with 10 mL of 3% BSA (Sigma, cat. A3983-100G) TBST (0.1% tween 20 in 1X TBS, Thermo Scientific, cat. 28358) buffer for 1 h at room temperature, followed by incubation with 2 μL of Streptavidin IR Dye 800 CW (LI-COR, cat. 926-32230), (1:5,000 dilution, 0.2 μg/mL) and 50 μL anti-A2aR (Santa Cruz Biotech, cat. sc-32261, clone 7F6-G5-A2, 200 μg/mL; at 1:200 or 1:1,000 dilution) for 1 h at room temperature. Next, membrane was washed 3 times with TBST buffer (5 min per wash) and incubated with 1 µL IRDye 680RD goat anti-mouse IgG antibody (LI-COR, cat. 926-68070; 1:10,000 dilution at 0.1 µg/mL) and 2 µL of Streptavidin IR Dye 800CW (LI-COR, cat. 926-32230), (1:5,000 dilution, 0.2 µg/mL). Membranes were subsequently washed 3 times with TBST buffer (5 min per wash), twice with MQ water and imaged on a LI-COR Odyssey using automatic settings.

**Preparation of cell surface enriched samples for quantitative proteomic analysis**
250 µL of streptavidin magnetic beads (Thermo Fisher, cat. 88817) were added to Protein LoBind microcentrifuge tubes (Eppendorf, cat. 022431081) and washed 2x with 1 mL RIPA Buffer (Thermo Fisher, cat. 89900). Approximately 1.0 mg of cell lysate was added to the pre-washed streptavidin magnetic beads and incubated for 3 hours at room temperature. A magnetic rack was used to pellet the beads and remove the lysate supernatant. The beads were sequentially washed 3x with each of the following: 1 mL of 1% SDS, 1 mL of 1M NaCl, and 1 mL of 10% EtOH, all prepared in 1x DPBS and incubating for 5 min in between washes. A final wash was done with 1 mL RIPA Buffer. The beads were then resuspended in 30 µL of 4x Laemmli sample buffer (Boston BioProducts, cat. BP-110R) containing 20 mM DTT and 25 mM biotin. Beads were heated for 10 min at 95 °C and were then placed on the magnetic rack. The supernatant was transferred to a new Protein LoBind microcentrifuge tube and stored at -80 °C. Quantitative proteomic sample preparation and analysis was performed by IQ Proteomics (Cambridge, MA).

**LC-MS/MS-based proteomic analysis of labeled cells**

For LC-MS analysis at IQ Proteomics, mass spectra were acquired on an Orbitrap Fusion Lumos coupled to an EASY nanoLC-1000 (or nanoLC-1200) (Thermo Fisher) liquid chromatography system. Approximately 2 µg of peptides were loaded on a 75 µm capillary column packed in-house with Sepax GP-C18 resin (1.8 µm, 150 Å, Sepax) to a final length of 35 cm. Peptides were separated using a 110-minute linear gradient from 8% to 28% acetonitrile in 0.1% formic acid. The mass spectrometer was operated in a data dependent mode. The scan sequence began with FTMS1 spectra (resolution = 120,000; mass range of 350-1400 m/z; max injection time of 50 ms; AGC target of $1 \cdot 10^6$; dynamic exclusion for 60 seconds with a +/- 10 ppm window). The ten most intense precursor ions were selected for MS2 analysis via collisional-induced dissociation (CID) in the ion trap (normalized collision energy (NCE) = 35; max injection time = 100 ms; isolation window of 0.7 Da; AGC target of $1.5 \cdot 10^4$). Following MS2 acquisition, a synchronous-precursor-selection (SPS) MS3 method was enabled to select eight MS2 product ions for high energy collisional-induced dissociation (HCD) with analysis in the Orbitrap (NCE = 55; resolution = 50,000; max injection time = 86 ms; AGC target of $1.4 \cdot 10^5$; isolation window at 1.2 Da for +2 m/z, 1.0 Da for +3 m/z or 0.8 Da for +4 to +6 m/z). All mass spectra were converted to mzXML using a modified version of ReAdW.exe. MS/MS spectra were searched against a concatenated 2018 human Uniprot protein database containing common contaminants (forward + reverse sequences) using the SEQUEST algorithm[10]. Database search criteria are as follows: fully tryptic with two missed cleavages; a precursor mass tolerance of 50 ppm and a fragment ion tolerance of 1 Da; oxidation of methionine (15.9949 Da) was set as differential modifications. Static modifications were carboxyamidomethylation of cysteines (57.0214) and TMT on lysines and N-termini of peptides (229.1629). Peptide-spectrum matches were filtered using linear discriminant analysis[11] and adjusted to a 1% peptide false discovery rate (FDR)[12].

All bioinformatic analysis of LC-MS/MS data was performed in the R statistical computing environment [R Development Core Team, R (R foundation for statistical computing Vienna, Austria, 2011)]. Peptide level abundance data was used to identify the number of peptides corresponding to a protein in the experiment. Any protein with a single peptide quantification was removed to reduce the possibility that outliers would affect downstream proximal calls. Peptide level abundance data was then normalized to the summed total abundance for each sample separately. These totals were then averaged, and each normalized protein abundance value was multiplied by this average to rescale abundance data. Peptide level data was then merged to protein level data by taking the median of all peptides corresponding to a protein. Proteins were then filtered to remove any known contaminants identified from the database search and proteins which are known antibody contaminants (e.g. having IGK, IGK, or IGH present in the gene symbol and Immunoglobulin present in the Uniprot description). Data were then filtered to remove PRNP, a protein which is a known false positive consistently detected across almost all experiments. Protein abundances were $\log_2$ transformed and subjected to linear modeling analysis with Limma[13]. Limma utilizes an empirical Bayes approach that allows for a realistic distribution of biological variance with small sample sizes per group. This program further utilizes the full dataset to shrink the observed sample variances towards a pooled estimate. This borrowing of variance information across proteins allows for a more accurate estimate of true variance, and improved power to detect real differences between groups. For each protein, abundance data was fit to a linear model with the experimental group as the input variable using the lmFit function. The $log_2FC$ values were estimated and p-values calculated for significance. P-values were then corrected for multiple comparisons using the false discovery rate (FDR) method by Benjamini and Hochberg[14]. Volcano plots were generated in R with the ggplot2 library[15]. $Log_2FC$ and p-value estimates from Limma were subset to those reaching a specified $log_2FC$ cutoff. Proteins were colored based on whether they fell above or below the $log_2$-fold cutoff threshold and were statistically significant (FDR corrected p-value of $< 0.05$).

#### Western blot analysis for cell surface target ID using SCH58261

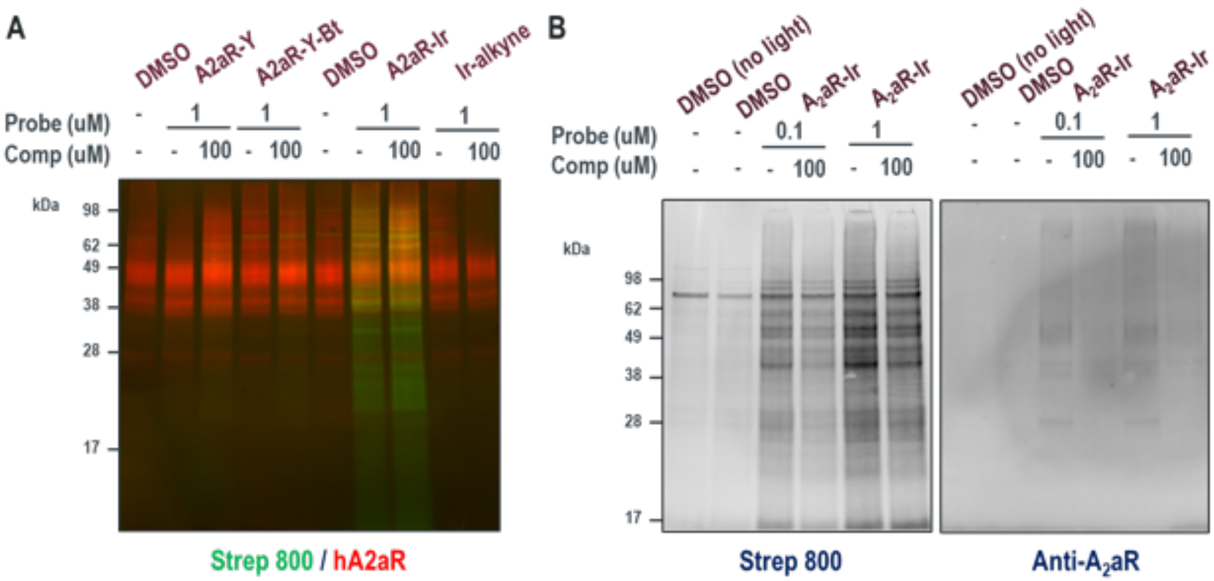

**A**) HEK-hA2aR cells were treated with A2aR conjugates or free photocatalyst in the presence or absence of competitor for 30 min, washed and irradiated for 10 min at 450 nm after treatment with Diaz-PEG3-Biotin. Cell lysates were processed and analyzed by western blot with Streptavidin-800 and anti-A2aR stains. Representative blots are shown from two independent biological replicates. **B**) Cell lysates were enriched with Streptavidin magnetic beads and analyzed by western blot with Streptavidin-800 and anti-A2aR stains.

**Western blot analysis of concentration and light-dependence for targeting of cell surface target ID of hA2aR**

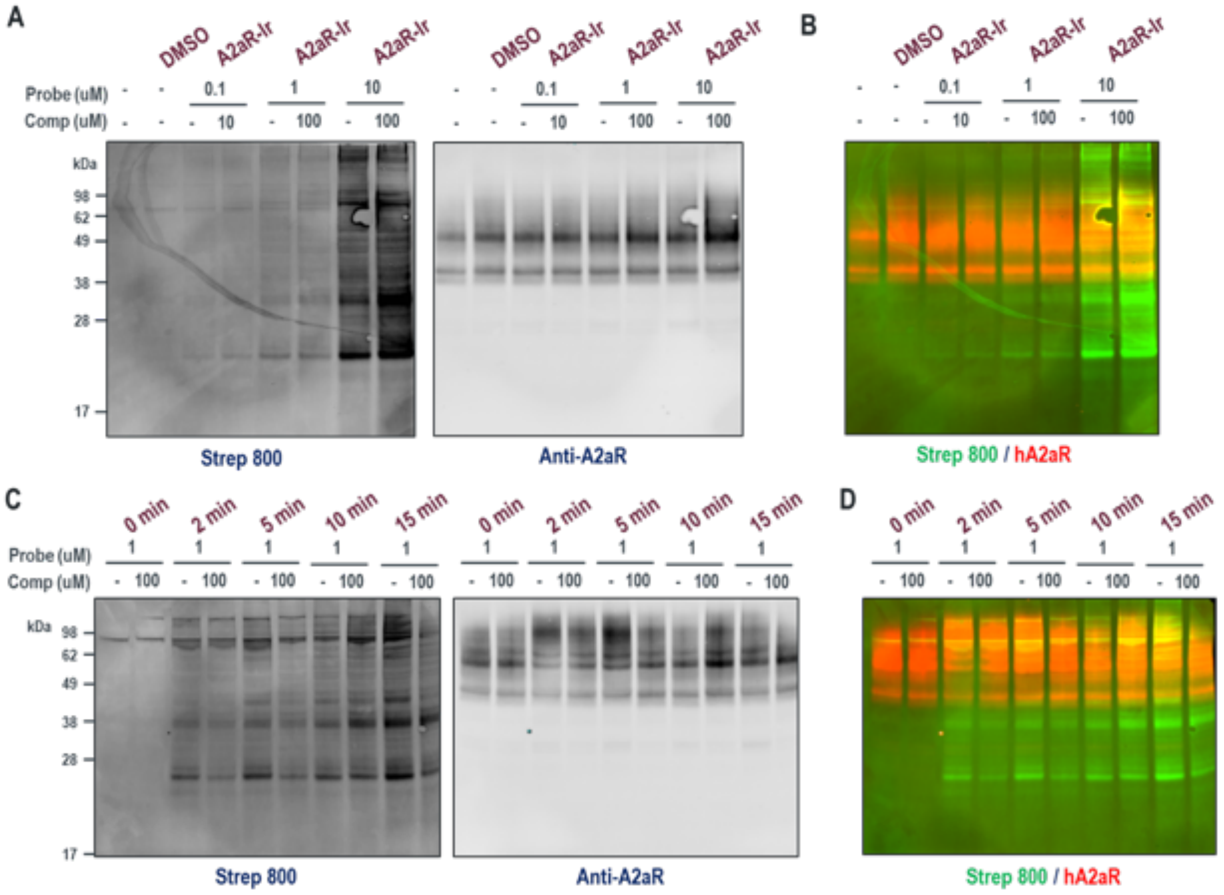

HEK-hA2aR cells were treated with SCH58261 conjugates (8) or free Ir photocatalyst (**12**) in the presence or absence of competitor for 30 min, washed and irradiated for 10 min at 450 nm after treatment with Diaz-PEG3-Biotin (9). Cell lysates were processed and analyzed by western blot with Streptavidin-800 and total protein stains. Representative blots are shown from two independent biological replicates (**A-D**). Panels (**A** and **C**) and (**B** and **D**) highlight Streptavidin 800 and anti-A2aR staining in separate or merged channels, respectively.

#### TMT proteomic analysis for SCH58261 μMap

##### PAL chemoproteomic target ID of Adora2a using SCH58261-Dz-alkyne (7) in A2a-expressing HEK293T cells

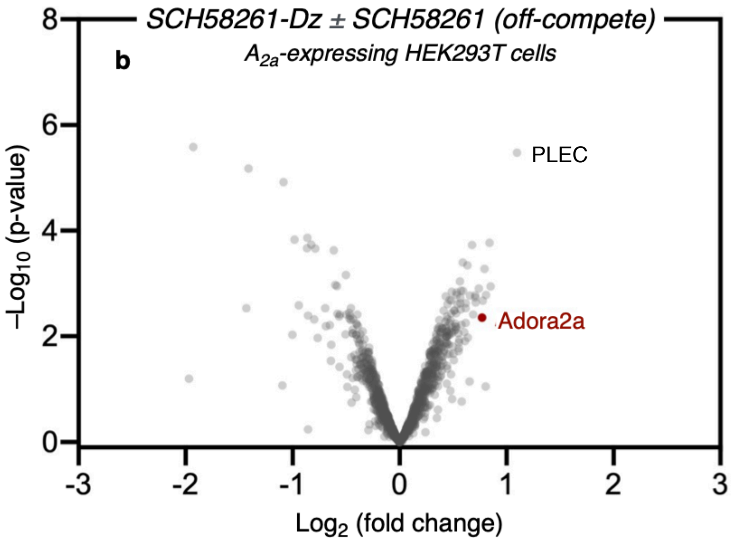

The labeling reaction was performed according to the general procedures. TMT-based chemoproteomics identified PLEC as the target.

**TMT-based chemoproteomic analysis of SCH58261-G1-Ir (8) labelling in A2a- expressing HEK293T cells vs. Free Iridium control (12)**

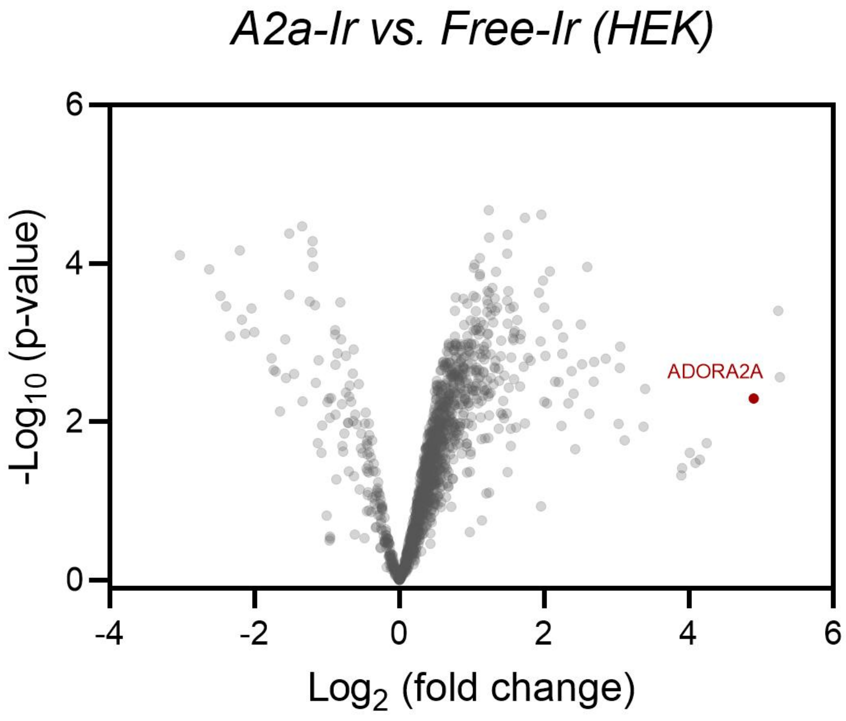

SCH58261-G1-Ir (8) $A_2$aR-Ir labelling identifies Adora2a as target by chemoproteomics when comparing ligand-targeted Ir versus free Ir photocatalyst (**12**). HEK-hA2aR cells were treated with SCH58261-G1-Ir conjugate (8) or free photocatalyst for 30 min, washed and irradiated for 10 min at 450 nm after treatment with Diaz-PEG3-Biotin (**9**). Cell lysates were prepared and processed for chemoproteomic experiments in triplicate.

**TMT-based chemoproteomic analysis of SCH58261-G1-Ir (8) labelling in A2a- expressing HEK293T cells**

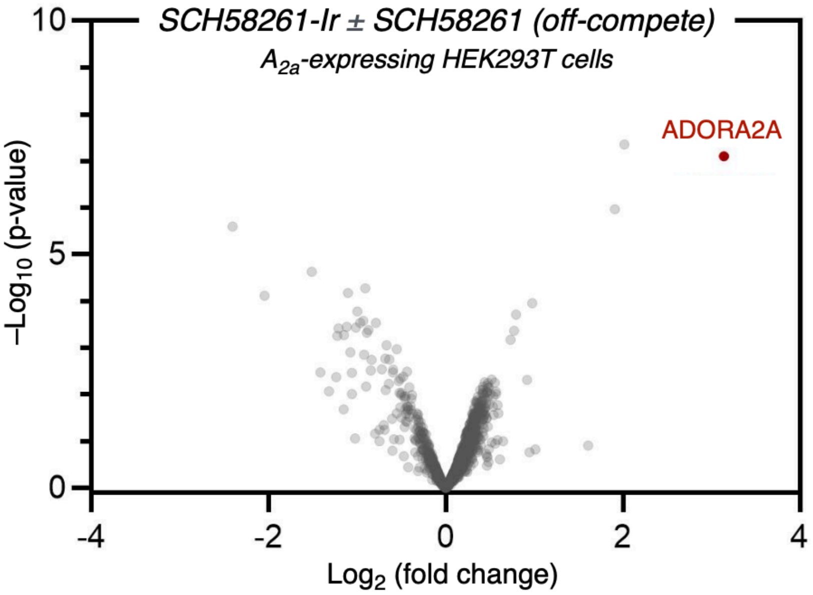

The labeling reaction was performed according to the general procedures. TMT-based chemoproteomics identified Adora2a as most enriched protein.

**TMT-based chemoproteomic analysis of SCH58261-G1-Ir (8) labelling in PC-12 cells**

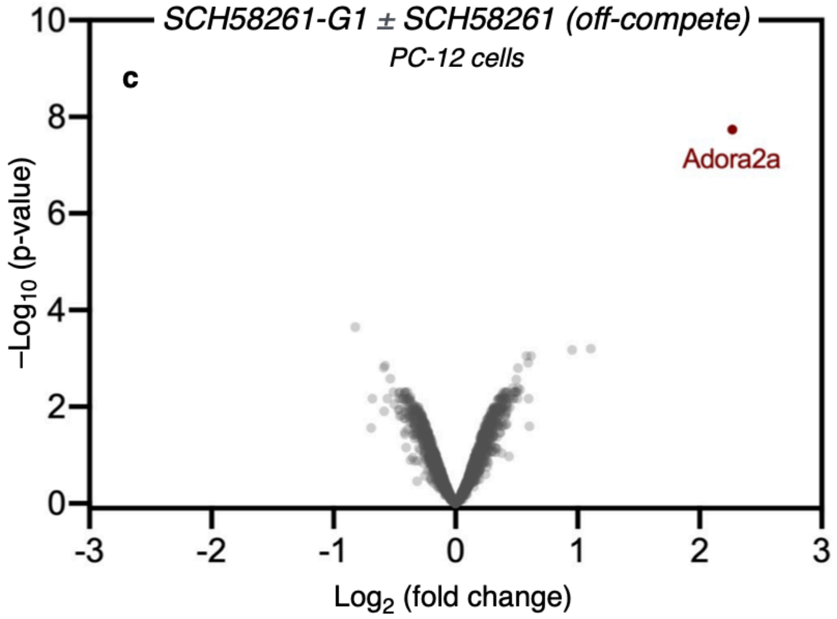

The labeling reaction was performed according to the general procedures. TMT-based chemoproteomics identified Adora2a as most enriched protein.

### GPR40 cell-based labelling, biochemical assays and proteomics

#### MK-8666 probes targeting GPR40 structure key

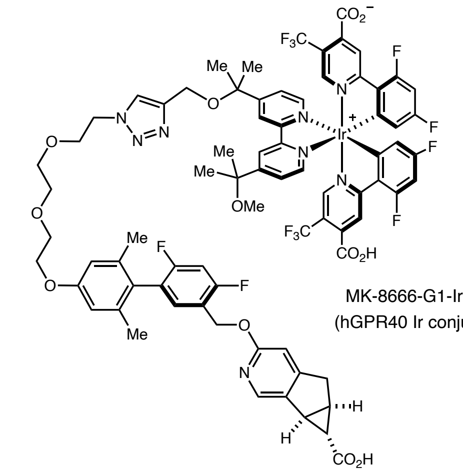

MK-8666-G1-Ir (**25**)
(hGPR40 Ir conjugate)

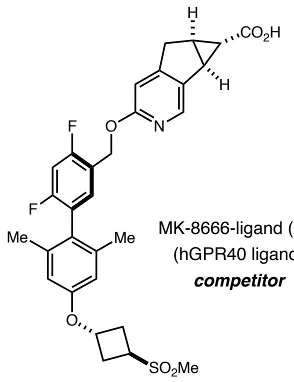

MK-8666-ligand (**26**)
(hGPR40 ligand)
***competitor***

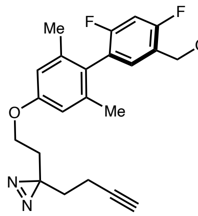

MK-8666-Dz (**27**)
(hGPR40-Yao)

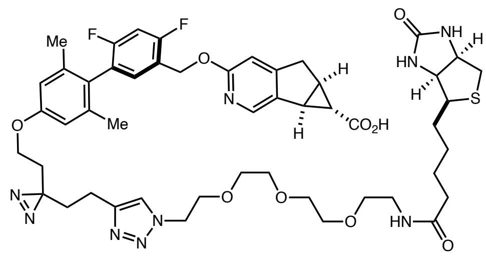

MK-8666-Dz-biotin (**28**)
(hGPR40-Yao-biotin)

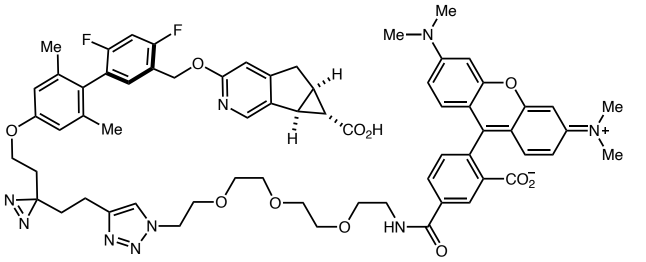

MK-8666-Dz-TAMRA (**29**)
(hGPR40-Yao-TAMRA)

#### Synthesis of conjugates

##### *Tert*-butyl (3-oxo-1-(4-(3-(trifluoromethyl)-3*H*-diazirin-3-yl)phenyl)-6,9,12-trioxa-2-azatetradecan-14-yl)carbamate (22)

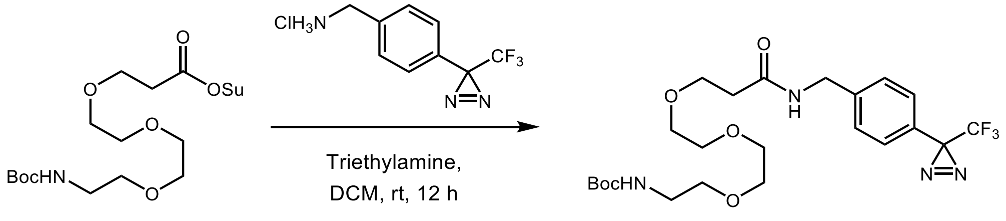

21 22

An 8 mL scintillation vial was charged with 4-[3-(trifluoromethyl)-3*H*-diazirin-3-yl]benzylamine HCl (20.0 mg, 0.079 mmol). This was dissolved in 1 ml of DCM, and triethylamine (0.40 mmol) and NHS-PEG3-NHBoc (33.1 mg, 0.079 mmol) were added. The reaction was stirred at room temperature for 12 hours. After completion, the reaction solution was concentrated under reduced pressure, the residue dissolved in DMSO, and purified by normal phase chromatography (0-50% EtOAc/Hexanes) to give the product as a white solid (36.2 mg, 88% yield).

$^{1}$H NMR (500 MHz, $d_6$-DMSO): δ 7.35 (d, 2H, $J$ = 8.1 Hz), 7.17 (d, 2H, $J$ = 8.0 Hz), 7.00 (bs, 1H), 6.00 (bs, 1H), 4.48 (d, 2H, $J$ = 5.9 Hz), 3.78 (t, 2H, $J$ = 5.7 Hz), 3.66 (dd, 2H, $J$ = 6.1, 3.2 Hz), 3.61 (dd, 2H, $J$ = 5.9, 3.3 Hz), 3.57-3.50 (m, 4H), 3.48 (t, 2H, $J$ = 5.2 Hz), 3.29 (s, 2H), 2.56 (t, 2H, $J$ = 5.7 Hz), 1.45 (s, 9H). $^{13}$C NMR (125 MHz, $d_6$-DMSO): δ 171.8, 155.9, 140.7, 127.9, 126.7, 125.4, 122.1 (q, $J$ = 272.8, 70.4, 70.3, 70.2, 70.1, 67.2, 42.7, 42.5, 40.3, 36.9, 28.4, 70.4, 70.3, 70.2, 70.1. 67.2, 42.7, 42.5, 40.3, 36.9, 28.4. $^{19}$F NMR (376 MHz, $d_4$-MeOD): δ -65.34 (s) HRMS (ESI-TOF) m/z calcd. for $C_{23}H_{34}F_3N_4O_6^+$ ([M + H]$^+$) 519.2425, found 519.2374.

#### MK-8666-PEG3-$N_3$

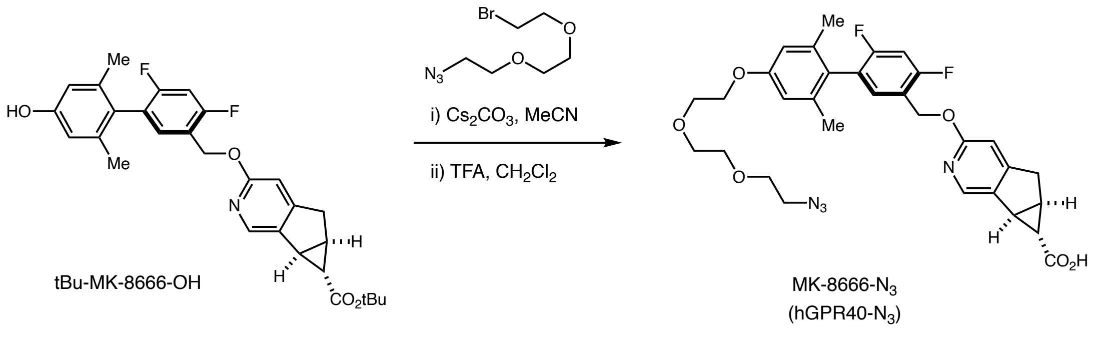

tBu-MK-8666-OH (96.2 mg, 0.195 mmol) was dissolved in MeCN (2 mL) and $Cs_2CO_3$ (325.6 mg, 0.999 mmol) was added, followed by 1-azido-2-(2-(2-bromoethoxy)ethoxy)ethane (78.5 mg, 0.330 mmol) dissolved in 0.3 mL DMF. The reaction mixture was stirred at 60 °C for 2 h, diluted with EtOAc, washed with saturated $NH_4Cl$ once and twice with brine. The organic layers were dried over $Na_2SO_4$, filtered, and concentrated to afford tert-butyl-MK-8666-PEG3-$N_3$ (127 mg, 0.195 mmol, 100 yield) as a clear oil. This intermediate (127 mg, 0.195 mmol) was dissolved in DCM (1.5 mL) and TFA (1 mL, 12.98 mmol) was added. Stirred at room temperature for 1 h, at which point LC-MS indicated complete conversion. Reaction mixture was concentrated and purified by HPLC, Gibson, Phenomenex C18, flow rate 20 mL/min, 8 min run, 30-100% MeCN in water with 0.05% TFA. Desired compound eluted at 100% MeCN to give MK-8666-$N_3$ (119 mg, 0.168 mmol, 86 % yield) as a white solid.

$^{1}$H-NMR (500 MHz, $CD_3OD$): δ 8.15 (s, 1H), 7.31 (t, J = 8.2 Hz, 1H), 7.11 (t, J = 9.6 Hz, 1H), 7.00 (s, 1H), 6.71 (s, 2H), 5.43 (s, 2H), 4.16 – 4.11 (m, 2H), 3.89 – 3.83 (m, 2H), 3.74 (dd, J = 6.0, 3.3 Hz, 2H), 3.72 – 3.65 (m, 4H), 3.37 (t, J = 4.9 Hz, 3H), 3.14 (d, J = 19.1 Hz, 1H), 3.02 – 2.96 (m, 1H), 2.50 (td, J = 6.4, 3.3 Hz, 1H), 1.96 (s, 6H), 1.23 (t, J = 2.7 Hz, 1H). $^{13}$C-NMR (126 MHz, $CD_3OD$): δ 173.9, 161.4, 161.3, 160.8, 160.7, 159.4, 159.3, 159.2, 158.8, 158.7, 158.4, 138.3, 137.8, 134.6, 133.5, 133.5, 133.4, 126.4, 124.0, 123.9, 123.9, 123.8, 119.9, 119.9, 119.9, 119.8, 113.2, 107.5, 103.8, 103.5, 103.3, 70.4, 70.2, 69.8, 69.6, 67.0, 62.5, 50.4, 34.9, 30.4, 30.0, 26.2, 19.4. $^{19}$F-NMR (471 MHz, $CD_3OD$): δ -77.31, -111.61, -111.63, -117.41, -117.43.
HRMS (ESI-TOF): m/z calcd. for $C_{31}H_{32}F_2N_4O_6$ ($[M+H]^+$) 595.2368, found 595.2374

**MK-8666-Dz-alkyne (27)**

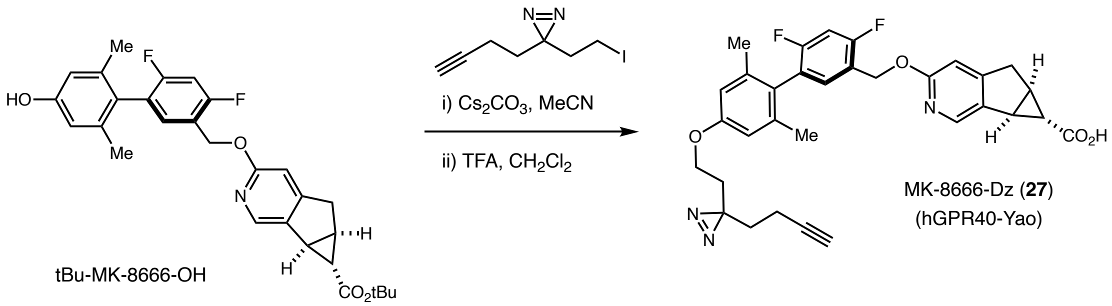

tBu-MK-8666-OH (100.5 mg, 0.203 mmol) was dissolved in MeCN (2 mL) and $Cs_2CO_3$ (332.2 mg, 1.020 mmol) was added, followed by addition of 3-(but-3-yn-1-yl)-3-(2-iodoethyl)-3*H*-diazirine (77.6 mg, 0.313 mmol) dissolved in 0.3 mL DMF. The reaction was stirred at 35 °C for overnight. After such time, the mixture was stirred at 40 °C for 2 h following the addition of another portion of alkyl iodide (77.6 mg, 0.313 mmol). The reaction mixture was diluted with EtOAc, washed with saturated $NH_4Cl$ once and twice with brine. The organic layers were dried over $Na_2SO_4$, filtered and concentrated. This residue was purified by column chromatography to on silica gel Isolute Flash Si; 12 g prepacked, eluting with EtOAc/hexanes (EtOAc gradient from 20 to 80%) to give tert-butyl tBu-MK-8666-Dz (80.2 mg, 0.131 mmol, 64.3 % yield) as a very light yellow oil. This intermediate (49.5 mg, 0.081 mmol) was dissolved in DCM (1.5 mL) and TFA (1 mL, 12.98 mmol) was added and stirred at room temperature for 1 h. LC-MS indicated complete conversion. Reaction mixture was concentrated and purified by HPLC, Gibson, Phenomenex C18, flow rate 20 mL/min, 10 min run, 35-100% ACN in water with 0.05% TFA. Desired compound eluted at 100% MeCN to give MK-8666-Dz (27) (48 mg, 0.071 mmol, 89 % yield) as a white solid.

$^1$H-NMR (500 MHz, $CD_3OD$): δ 8.16 (s, 1H), 7.33 (t, J = 8.2 Hz, 1H), 7.12 (t, J = 9.6 Hz, 1H), 7.04 (s, 1H), 6.70 (s, 2H), 5.44 (s, 2H), 3.86 (t, J = 6.0 Hz, 2H), 3.40 – 3.34 (m, 1H), 3.17 (d, J = 19.2 Hz, 1H), 3.01 (dd, J = 6.5, 1.9 Hz, 1H), 2.52 (td, J = 6.4, 3.3 Hz, 1H), 2.29 (t, J = 2.6 Hz, 1H), 2.09 (td, J = 7.5, 2.6 Hz, 2H), 1.98 (s, 6H), 1.88 (t, J = 6.0 Hz, 2H), 1.71 (t, J = 7.5 Hz, 2H), 1.25 (t, J = 2.8 Hz, 1H). $^{13}$C-NMR (126 MHz, $CD_3OD$): δ 173.9, 161.4, 161.3, 161.2, 160.9, 160.8, 159.8, 159.4, 159.3, 158.9, 158.8, 158.2, 137.9, 137.8, 134.7, 133.6, 133.5, 133.5, 126.5, 124.1,

124.0, 123.9, 123.9, 119.8, 119.7, 119.7, 113.3, 113.1, 107.5, 103.8, 103.6, 103.4, 82.3, 68.9, 62.8, 62.3, 34.9, 32.5, 32.4, 30.3, 30.0, 26.5, 26.2, 19.4, 12.5. $^{19}$F-NMR (471 MHz, $CD_3OD$): δ -77.42, -111.54, -111.56, -117.42, -117.44. HRMS (ESI-TOF): m/z calcd. for $C_{32}H_{29}F_2N_3O_4$ ([M+H] $^+$) 558.2204, found 558.2214

**MK-8666-Dz-Biotin (28)**

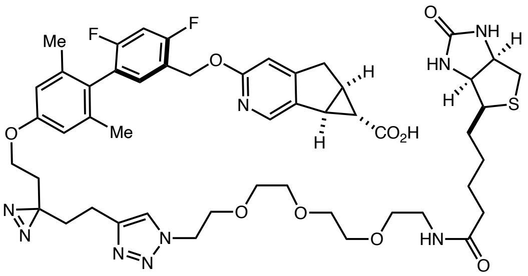

MK-8666-Dz-biotin (**28**)
(hGPR40-Yao-biotin)

Alkyne fragment (0.012 mmol, 1 eq) and azide fragment (0.012 mmol, 1 eq) were dissolved in DMSO (0.9 mL). Copper (II) sulfate pentahydrate (0.030 mmol, 2.5 eq) was dissolved in water (0.1 mL) and added to reaction mixture. Reaction mixture was stirred for 5 min followed by addition of sodium ascorbate (0.114 mmol, 10 eq) and stirred at room temperature overnight. Reaction mixture was filtered and purified by HPLC, Gibson, Phenomenex C18, flow rate 20 mL/min, 8 min run, 30-100% ACN in water with 0.05% TFA.

$^1$H-NMR (500 MHz, $CD_3OD$): δ 8.20 (s, 1H), 7.90 (s, 1H), 7.35 (t, J = 8.2 Hz, 1H), 7.16 (t, 2H), 6.71 (s, 2H), 5.48 (s, 2H), 4.54 (t, J = 5.0 Hz, 2H), 4.49 (dd, J = 7.8, 4.9 Hz, 1H), 4.30 (dd, J = 7.8, 4.5 Hz, 1H), 3.91 – 3.83 (m, 4H), 3.60 (d, J = 11.5 Hz, 8H), 3.53 (t, J = 5.5 Hz, 2H), 3.42 (dd, J = 19.4, 6.4 Hz, 1H), 3.36 (t, J = 5.5 Hz, 8H), 3.25 – 3.17 (m, 2H), 3.04 (dd, J = 6.4, 2.0 Hz, 1H), 2.93 (dd, J = 12.7, 5.0 Hz, 1H), 2.71 (d, J = 12.7 Hz, 1H), 2.68 (s, 4H), 2.61 (t, J = 7.8 Hz, 2H), 2.55 (dt, J = 6.4, 3.2 Hz, 1H), 2.21 (t, J = 7.4 Hz, 2H), 1.99 (s, 6H), 1.95 – 1.90 (m, 2H), 1.85 (t, J = 6.0 Hz, 2H), 1.67 (dddd, J = 38.9, 28.9, 14.2, 6.9 Hz, 4H), 1.44 (p, J = 7.8 Hz, 2H), 1.28 (t, J = 2.8 Hz, 1H). $^{13}$C-NMR (126 MHz, $CD_3OD$): δ 174.7, 173.8, 164.7, 160.9, 158.9, 158.2, 137.9, 137.1,

133.7, 126.5, 123.2, 113.1, 107.6, 103.9, 103.7, 103.5, 70.1, 70.1, 70.0, 69.8, 69.2, 68.9, 63.4, 62.3, 61.9, 60.2, 55.6, 50.1, 39.6, 39.0, 38.9, 35.3, 35.2, 32.6, 32.5, 30.2, 29.9, 28.4, 28.1, 26.6, 26.2, 25.4, 19.4, 19.3. $^{19}$F-NMR (471 MHz, $CD_3OD$): δ -77.54, -111.31, -111.33, -117.31, -117.33. HRMS (ESI-TOF): m/z calcd. for $C_{50}H_{61}F_2N_9O_9S$ ($[M+H]^+$) 1002.4359, found 1002.4335

**MK-8666-Dz-TAMRA (29)**

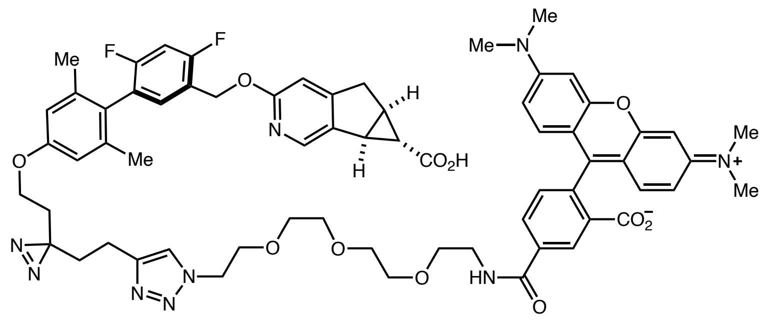

MK-8666-Dz-TAMRA (**29**)
(hGPR40-Yao-TAMRA)

$^1$H-NMR (500 MHz, $CD_3OD$): δ 8.79 (d, J = 1.6 Hz, 1H), 8.27 (dd, J = 7.9, 1.6 Hz, 1H), 8.12 (s, 1H), 7.85 (s, 1H), 7.50 (d, J = 7.9 Hz, 1H), 7.28 (t, J = 8.2 Hz, 1H), 7.12 (t, J = 8.4 Hz, 3H), 7.04 (d, J = 9.5 Hz, 2H), 6.97 (d, J = 2.2 Hz, 2H), 6.87 (s, 1H), 6.66 (s, 2H), 5.40 (s, 2H), 4.49 (t, J = 5.0 Hz, 2H), 3.85 (t, J = 5.0 Hz, 2H), 3.81 (t, J = 6.0 Hz, 2H), 3.73 (t, J = 5.3 Hz, 2H), 3.70 – 3.58 (m, 10H), 3.31 (s, 11H), 3.10 (d, J = 18.9 Hz, 1H), 2.96 (dd, J = 6.5, 2.0 Hz, 1H), 2.54 (t, J = 7.8 Hz, 2H), 2.48 (td, J = 6.4, 3.3 Hz, 1H), 1.94 (s, 6H), 1.88 (t, 2H), 1.80 (t, J = 6.0 Hz, 2H), 1.19 (t, J = 2.8 Hz, 1H). $^{13}$C-NMR (126 MHz, $CD_3OD$): δ 174.0, 166.7, 161.7, 159.2, 158.1, 157.6, 157.5, 139.0, 137.9, 136.7, 136.2, 133.3, 131.4, 130.9, 130.5, 129.9, 126.6, 123.1, 114.2, 113.3, 113.1, 107.5, 103.5, 103.3, 96.1, 70.2, 70.1, 69.9, 69.9, 69.1, 68.9, 62.3, 61.9, 49.9, 39.8, 39.5, 34.7, 32.6, 32.5, 30.4, 30.1, 26.6, 26.2, 19.4, 19.3. $^{19}$F-NMR (471 MHz, $CD_3OD$): δ -77.35, -112.20, -112.22, -117.62, -117.64. HRMS (ESI-TOF): m/z calcd. for $C_{65}H_{67}F_2N_9O_{11}$ ($[M+H]^+$) 1188.5006, found 1188.5004

**MK-8666-G1-Ir (25)**

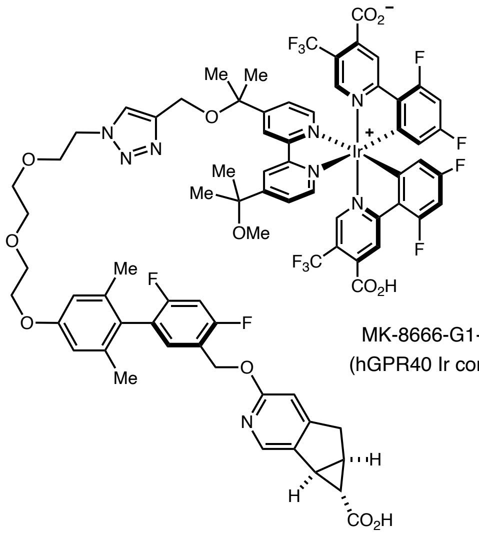

[1]H-NMR (500 MHz, $CD_3OD$): δ 9.02 (s, 1H), 8.87 (s, 1H), 8.68 (s, 2H), 8.06 (dd, J = 6.1, 3.2 Hz, 3H), 7.80 – 7.74 (m, 2H), 7.62 (d, J = 10.9 Hz, 2H), 7.23 (t, J = 8.0 Hz, 1H), 7.04 (t, J = 9.6 Hz, 1H), 6.82 (ddd, J = 11.7, 9.1, 2.0 Hz, 2H), 6.58 (s, 2H), 5.87 – 5.82 (m, 2H), 5.37 (s, 2H), 4.55 (d, J = 617.5 Hz, 2H), 4.38 (s, 2H), 4.07 – 4.03 (m, 2H), 3.87 (t, J = 4.9 Hz, 2H), 3.76 – 3.72 (m, 2H), 3.65 – 3.60 (m, 4H), 3.20 (s, 3H), 3.08 (d, J = 18.9 Hz, 1H), 2.93 (d, J = 6.5 Hz, 1H), 2.63 (s, 1H), 2.47 – 2.42 (m, 1H), 1.91 (s, 1H), 1.87 (s, 6H), 1.68 – 1.56 (m, 12H), 1.15 (s, 1H).

[13]C-NMR (126 MHz, $CD_3OD$): δ 173.9, 168.4, 162.3, 161.8, 158.9, 158.3, 156.2, 156.1, 151.0, 150.8, 146.4, 137.8, 133.4, 126.5, 126.4, 126.1, 124.6, 123.1, 122.8, 122.6, 120.4, 114.0, 113.1, 103.5, 99.4, 77.4, 76.6, 70.2, 70.0, 69.4, 68.9, 67.0, 62.3, 56.8, 50.0, 49.9, 34.8, 30.4, 30.1, 26.8, 26.2, 26.1, 25.8, 19.4. [19]F-NMR (471 MHz, $CD_3OD$): δ -61.55, -61.59, -77.42, -103.13, -103.16, -103.21, -103.23, -107.66, -107.69, -107.71, -107.74, -111.94, -111.95, -117.48, -117.50.

HRMS (ESI-TOF): m/z calcd. for $C_{31}H_{32}F_2N_4O_6$ ([M+H] $^+$) 1713.4214, found 1713.4114, for $C_{31}H_{32}F_2N_4O_6$ ([M+H] $^{+2}$) 857.2146, found 857.2159

**GPR40 competitor** (26), which is derived from MK-8666, was synthesized as previously described[16].

#### Cell culture

HEK293-A2a cells were grown and expanded in EMEM (Sigma, cat. M6199-500) with 10% FBS, supplemented with 1X P/S and 400 µg/mL geneticin. Jurkat cells were grown and expanded in McCoy5a (Gibco, cat. 16600-082) media with 10% FBS and 1X P/S. All cells were maintained at 37 $^{\circ}$C and 5% $CO_2$ atmosphere in T175 flasks (Nunc, cat. 159910).

HEK293T WT and HEK cells stably expressing hGPR40 (clone 13) were grown in DMEM (Gibco, cat. 11965-084) with 10% FBS (Gibco, cat. 26140079), supplemented with 1X NE amino acids (Gibco, cat. 1140-050), 100 units/mL penicillin and 100µg/mL streptomycin (1X dilution from 100X stock Gibco, cat. 15140-122) and geneticin (final concentration 500 µg/mL; Gibco, cat. 10131-035). HEK293-A2a cells were grown and expanded in EMEM (Sigma, cat. M6199-500) with 10% FBS, supplemented with P/S and geneticin (400 ug/mL). SK-BR-3 cells were grown and expanded in McCoy5a (Gibco, cat. 16600-082) media with 10% FBS and P/S. All cells were maintained at 37$^{o}$C and 5% $CO_2$ atmosphere in T175 flasks (Nunc, cat. 159910).

#### Functional hGPR40 cellular assay

In Vitro Inositol Phosphate Turnover (hIP1) assay was used to evaluate potency of hGPR40 probes as previously described[17,18] with deviations as described below. Stable cell lines expressing human GPR40 (hGPR40/HEK293) were cultured in DMEM supplemented with 10% Fetal Bovine Serum (FBS), 1X Non-Essential Amino Acids (NEAA), 1X Penicillin-Streptomycin (P-S) and 0.5 mg/mL Geneticin (G418). Assay ready frozen Cell stocks were thawed and grown in a subconfluent state using standard cell culture procedures. The day before the experiment, the cells were harvested with nonenzymatic cell dissociation buffer and re-suspended cells to the 3M cells/20 mL for a final concentration of 7.5K cells/well in 50 µl media. Seed 50 µL into a 384-well CulturPlate using the Combi. The seeded plates were incubated overnight at 37 C. On the day of the experiment, the growth media was removed from the assay by Blue Cat Bio BlueWasher and 10 µL of IP1 stimulation buffer (Cis Bio IP-one Tb HTRF kit) supplemented with LiCl is added to each well. Test compounds dissolved in DMSO were serially diluted in ½ log increments starting from 2 mM and 50 nL of the compound dilution was acoustically added to each well (final starting concentration 10 uM). Plates were then incubated for 60 minutes at 37°C, 5% $CO_2$ and 10 μL of detection buffer (prepared as described in the Tb kit) is added to each well. The plates are then incubated one additional hour at room temperature. After the final incubation, the plates were read in a PHERAstar (337 nm excitation, dual emission 590 and 655 nm) with the method designed for HTRF assays. For each assay, a standard curve plate in which IP1 is titrated is also included. All fluorescent readings (using 590/665 nm ratio) are back calculated to a concentration of IP1 using the IP1 standard curve and the percent activity at each concentration of test compound is determined using 0% activation (basal activity) determined in those wells that contain DMSO alone, while 100% activity is determined in wells that contained a concentration of a partial allosteric agonist know to activate GPR40. The % activity is then plotted versus the concentration of test compound and the dose response curve fitted to a standard 4-parameter non- regression model using a custom in-house developed software package. Maximal % activity and EC50 are then determined for each test compound.

**Potency (nM) of hGPR40 probes in IP1 assay**

| Probe | Potency (nM) |
|---|---|
| MK-8666-ligand competitor (26) | 0.35 ± 0.16 |
| MK-8666-azide | < 0.038 |
| MK-8666-Dz-alkyne (27) | 0.49 ± 0.05 |
| MK-8666-Dz-biotin (28) | 0.84 ± 0.04 |
| MK-8666-Dz-TAMRA (29) | 14.74 ± 2.78 |
| MK-8666-G1-Ir (25) | n.d. |
| G1-Ir catalyst (12) | > 9950 |

#### Validation of GPR40 identification by proteomics

Following the general procedure for cell-membrane labelling experiments, HEK-hGPR40 labelling experiments were performed with 30 min incubation at room temperature to increase solubility of MK-8666 chemical probes. Cells were washed with cold 1X DPBS (2x; 1,000xg for

4 min). After elution from streptavidin beads, as described by the general procedure, samples were reduced and alkylated with 10 mM TCEP and 20 mM iodoacetamide at 65 °C for 20 minutes. Samples were cleaned up using SP3 beads as described previously[8]. Samples were digested using 2 ug/sample trypsin/lys-C (Pierce A40007) overnight at 37 °C. Supernatant was collected and desalted using an AssayMAP Bravo with C18 cartridges (Agilent 5190-6532). Samples were eluted from cartridges with 70% acetonitrile, 30% water 0.1% TFA. Samples were dried down by speedvac and resuspended in 95% water, 5% acetonitrile, 0.1% formic acid and injected on a QE-HF using a 50 cm column. Samples were analyzed with either a four-hour gradient from 3% to 40% acetonitrile or a two-hour gradient from 10% to 47% acetonitrile.

Data was searched with Maxquant version 1.6.17 for tryptic peptides allowing for variable methionine oxidation, and fixed carbamidomethylation. Data was analyzed using Perseus version 1.6.5. Data was filtered for contaminants and reverse data base hits, log2 transformed, and median normalized.

To verify the YLGAAFPLGYQAFR peptide hit the peptide was synthesized by Thermo Fisher Scientific. The synthetic peptide was analyzed on a QE-HF with a two-hour gradient from 10% to 47% acetonitrile. The retention time and MS2 were identical to what was observed for the endogenous YLGAAFPLGYQAFR peptide.

### WB analysis for MK8666-G1-Ir μMap labelling in HEK-hGPR40 cells

*Streptavidin staining and enrichment of 31 kDa band** assigned as putative hGPR40*

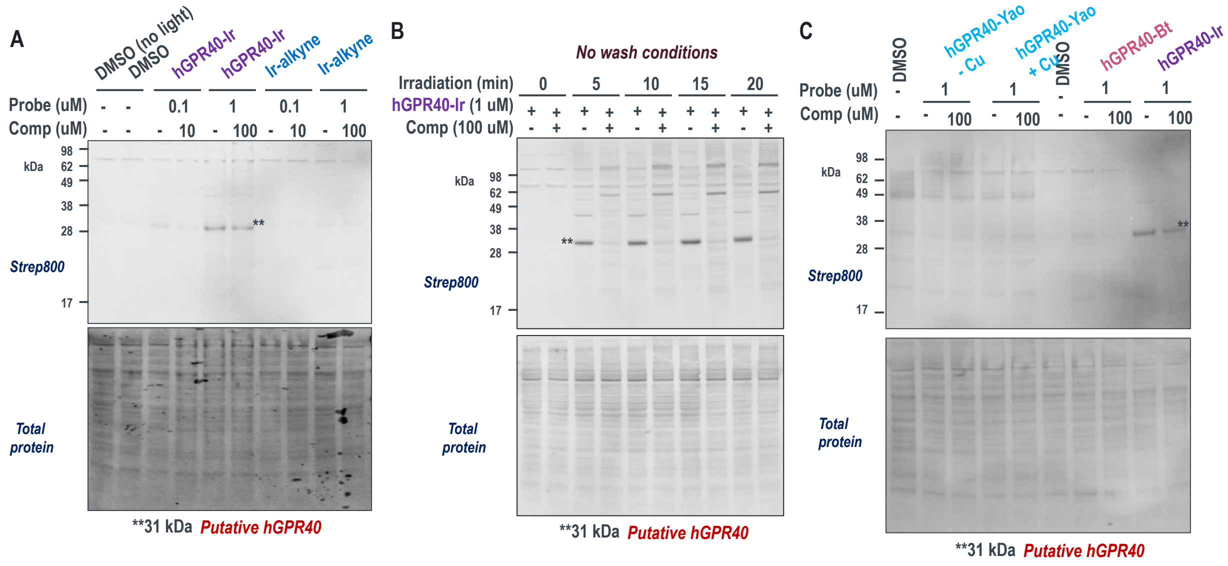

#### MK-8666-G1-Ir labeling identifies FFAR1 (GPR40) by quantitative proteomics

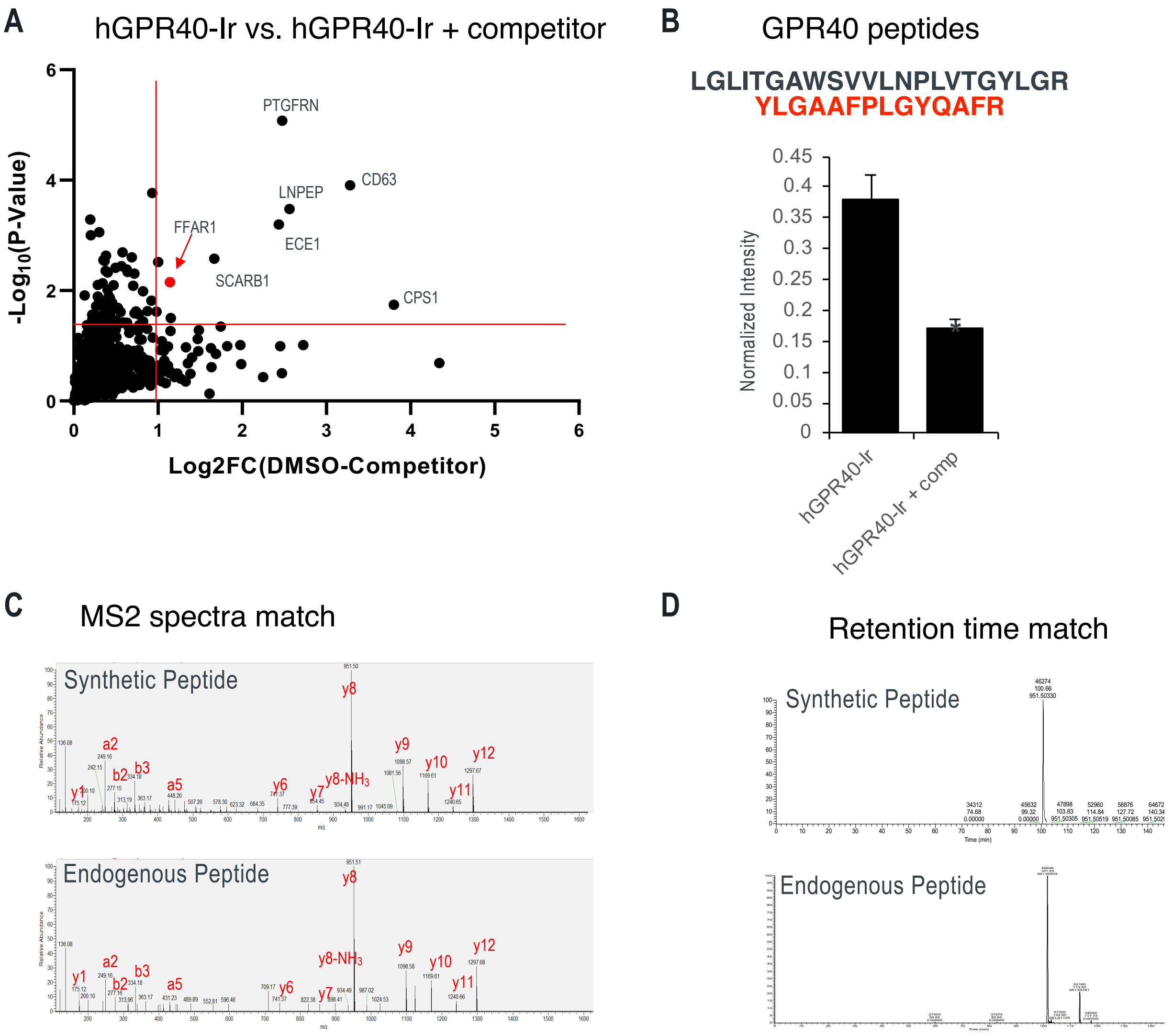

(A) FFAR1 is enriched by chemoproteomics after MK8666-G1-Ir labeling. Labeling experiments were performed in triplicate and processed for proteomics. (B) hGPR40 peptides detected by LC-MS and normalized intensities for the YLGAAFPLGYQAFR peptide. (student's t-test, $p \leq 0.01$, error bars are SEM). Validation of synthetic hGPR40 peptide confirms correct identification by matching MS2 spectra (C) and retention time (D).

(8) Hughes, C. S.; Moggridge, S.; Müller, T.; Sorensen, P. H.; Morin, G. B.; Krijgsveld, J.

Single-Pot, Solid-Phase-Enhanced Sample Preparation for Proteomics Experiments. *Nat. Protoc.* **2019**, *14* (1), 68–85. https://doi.org/10.1038/s41596-018-0082-x.

(15) Wickham, H. *Ggplot2: Elegant Graphics for Data Analysis*; Springer-Verlag New York, 2016.

(16) ANTIDIABETIC TRICYCLIC COMPOUNDS. WO2015073342, 2015.

(17) Pachanski, M. J.; Kirkland, M. E.; Kosinski, D. T.; Mane, J.; Cheewatrakoolpong, B.; Xue, J.; Szeto, D.; Forrest, G.; Miller, C.; Bunzel, M.; Plummer, C. W.; Chobanian, H. R.; Miller, M. W.; Souza, S.; Thomas-Fowlkes, B. S.; Ogawa, A. M.; Weinglass, A. B.; Di Salvo, J.; Li, X.; Feng, Y.; Tatosian, D. A.; Howard, A. D.; Colletti, S. L.; Trujillo, M. E. GPR40 Partial Agonists and AgoPAMs: Differentiating Effects on Glucose and Hormonal Secretions in the Rodent. *PLoS One* **2017**, *12* (10), e0186033.

(18) Plummer, C. W.; Clements, M. J.; Chen, H.; Rajagopalan, M.; Josien, H.; Hagmann, W. K.;

Miller, M.; Trujillo, M. E.; Kirkland, M.; Kosinski, D.; Mane, J.; Pachanski, M.; Cheewatrakoolpong, B.; Nolting, A. F.; Orr, R.; Christensen, M.; Campeau, L.-C.; Wright, M. J.; Bugianesi, R.; Souza, S.; Zhang, X.; Di Salvo, J.; Weinglass, A. B.; Tschirret-Guth, R.; Nargund, R.; Howard, A. D.; Colletti, S. L. Design and Synthesis of Novel, Selective GPR40 AgoPAMs. *ACS Med. Chem. Lett.* **2017**, *8* (2), 221–226. https://doi.org/10.1021/acsmedchemlett.6b00443.

### Spectroscopic data

#### Diazirine alkyne: [1]H NMR

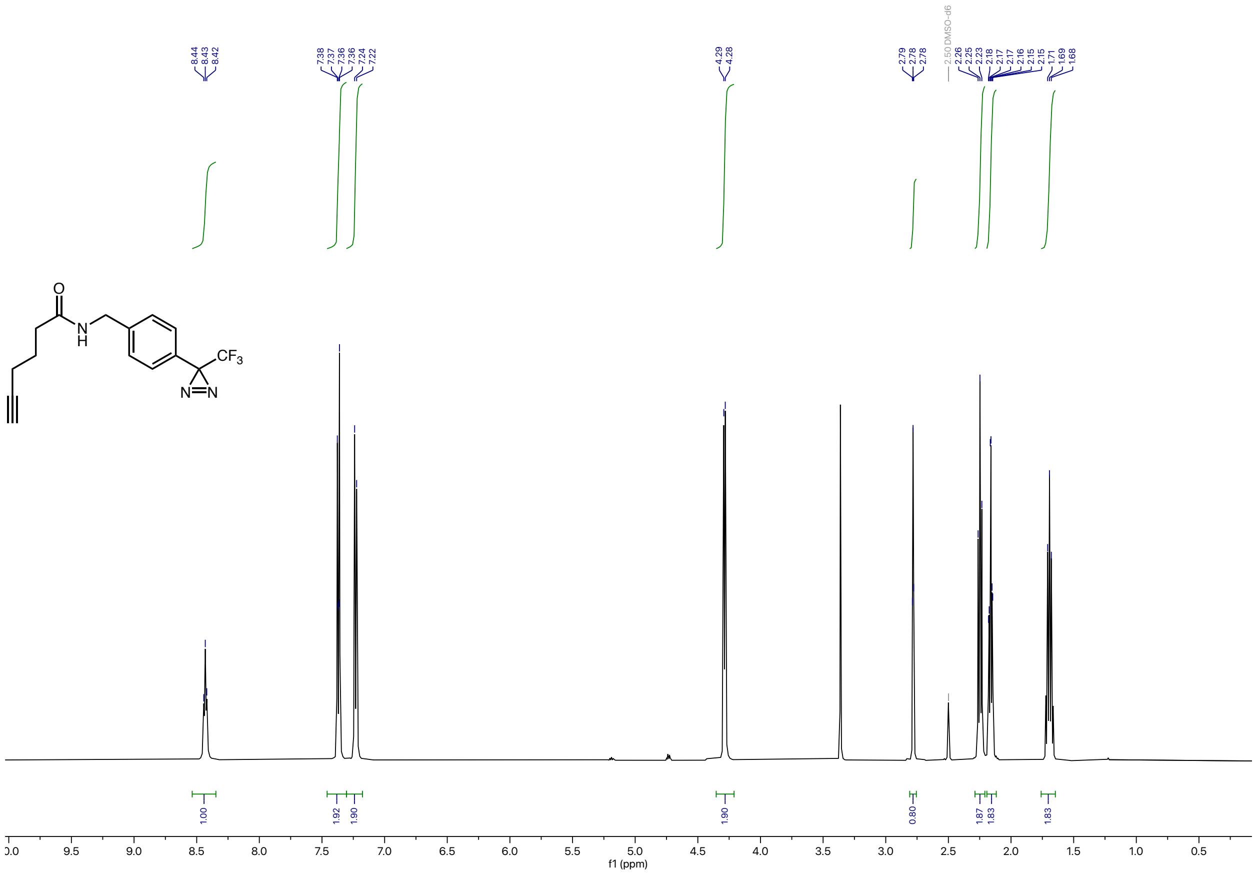

### Diazirine alkyne: $^{13}C$ NMR

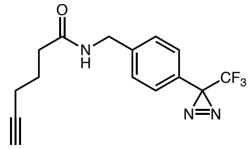

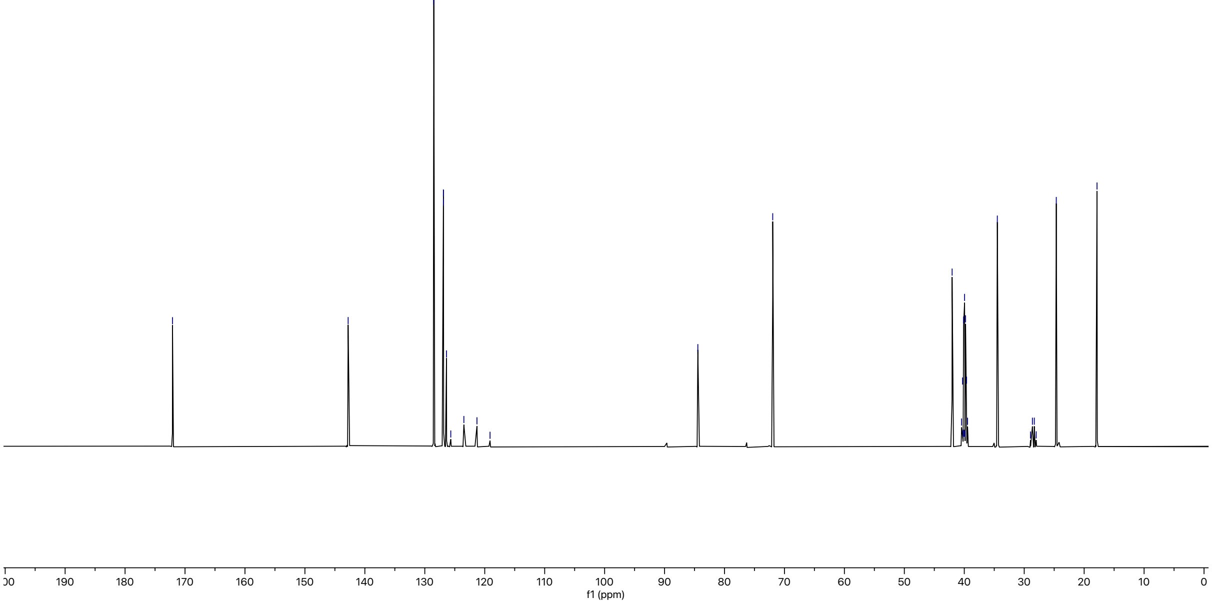

### Diazirine alkyne: $^{19}$F NMR

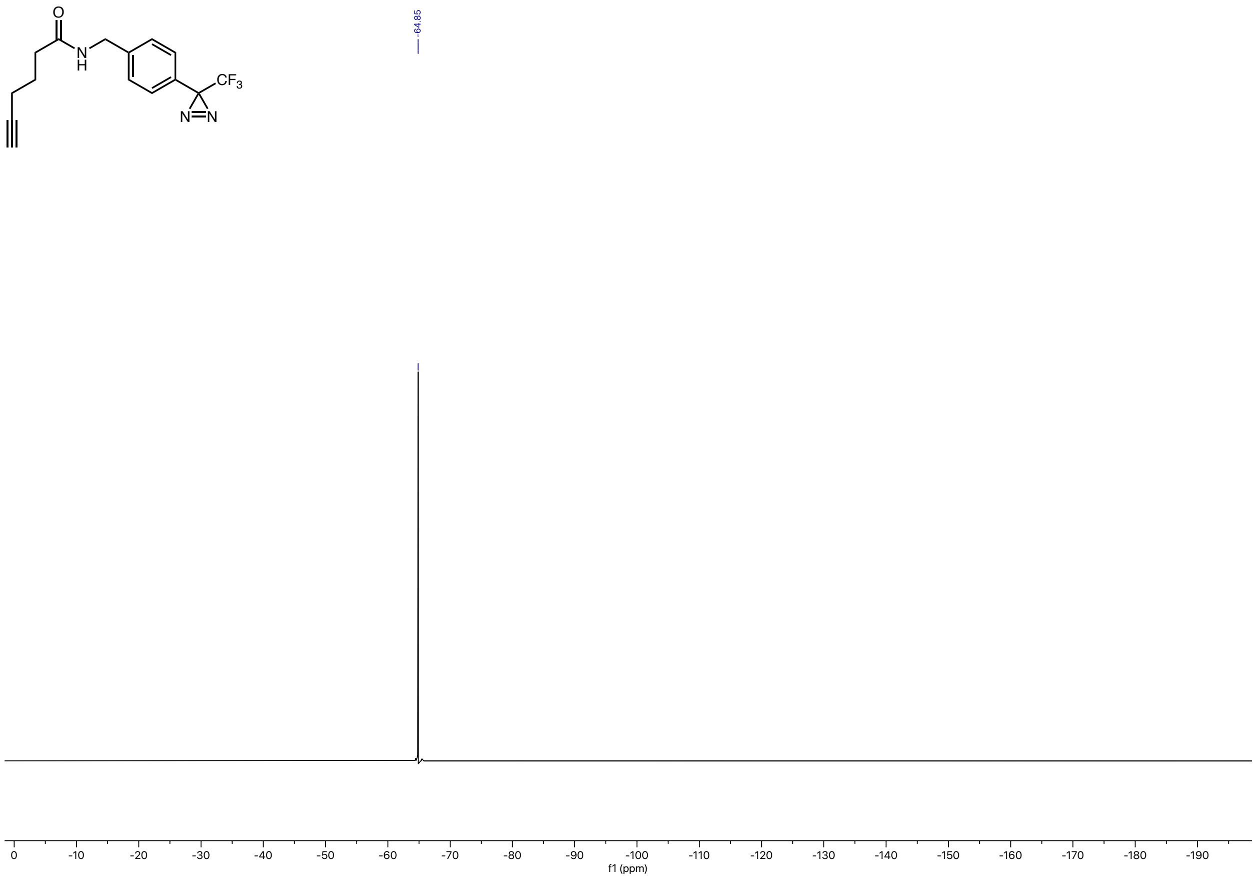

### 4-(2-methoxypropan-2-yl)-4'-(2-(prop-2-yn-1-yloxy)propan-2-yl)-2,2'-bipyridine: $^1$H NMR

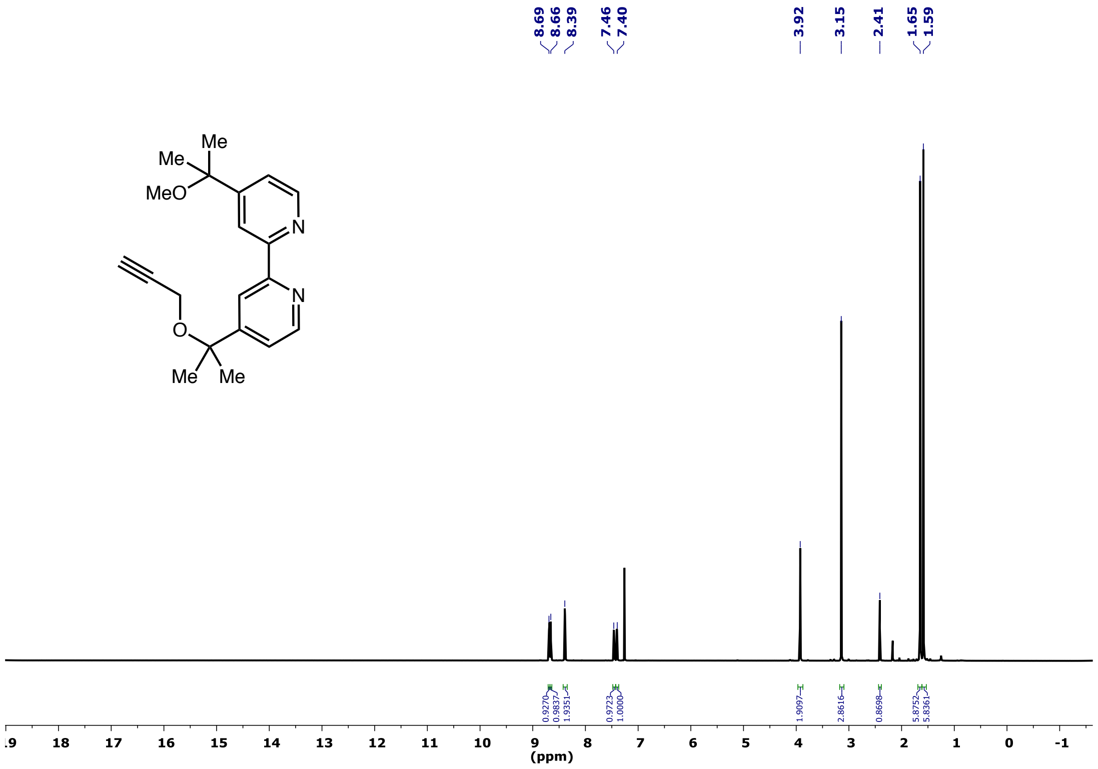

#### 4-(2-methoxypropan-2-yl)-4'-(2-(prop-2-yn-1-yloxy)propan-2-yl)-2,2'-bipyridine: $^{13}C$ NMR

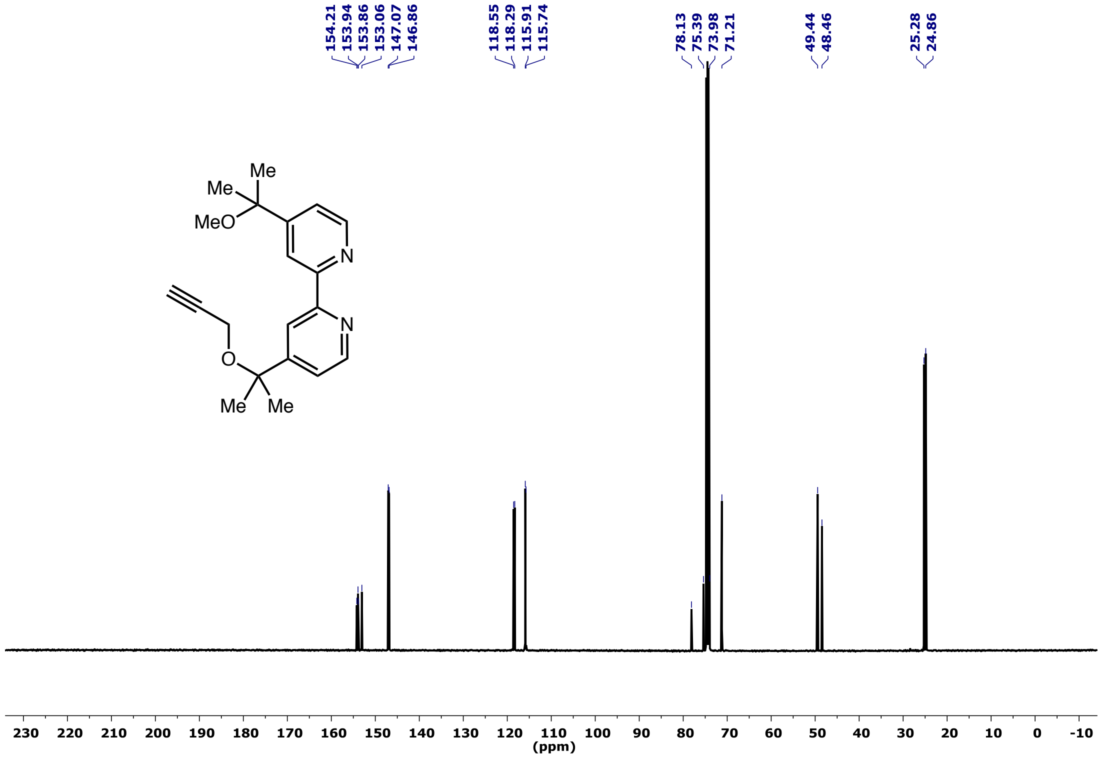

**4-(2-((3-(tert-butyldimethylsilyl)prop-2-yn-1-yl)oxy)propan-2-yl)-4'-(2-methoxypropan-2-yl)-2,2'-bipyridine: $^{1}$H NMR**

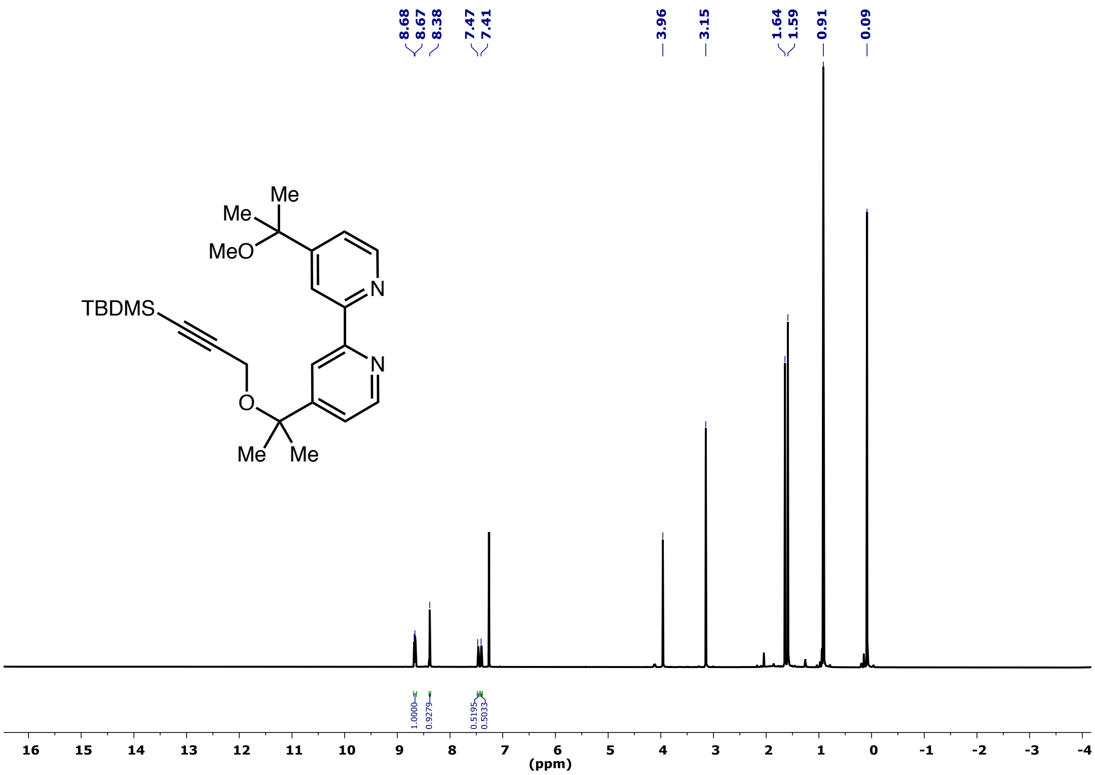

**4-(2-((3-(*tert*-butyldimethylsilyl)prop-2-yn-1-yl)oxy)propan-2-yl)-4'-(2-methoxypropan-2-yl)-2,2'-bipyridine: $^{13}$C NMR**

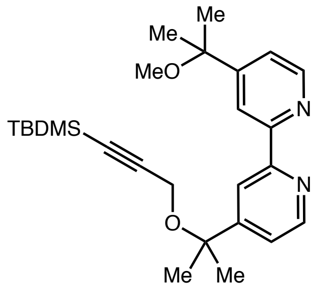

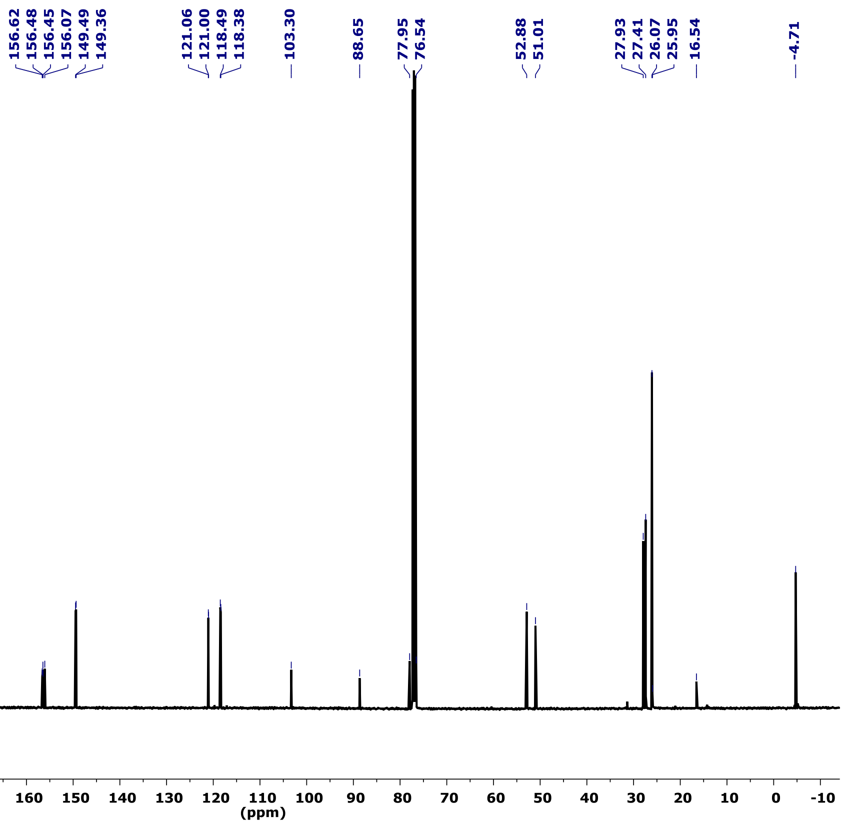

### Ir-TBS-alkyne: $^{1}H$ NMR

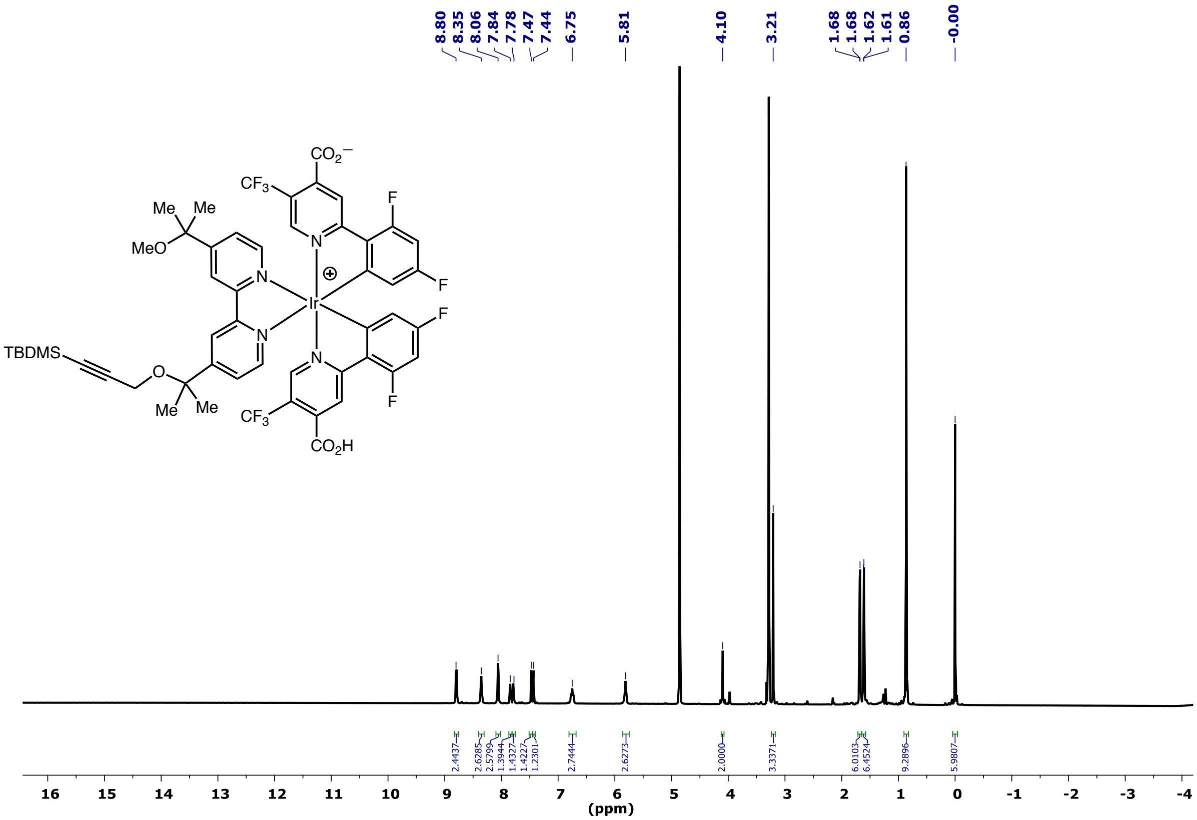

### Ir-TBS-alkyne: $^{13}C$ NMR

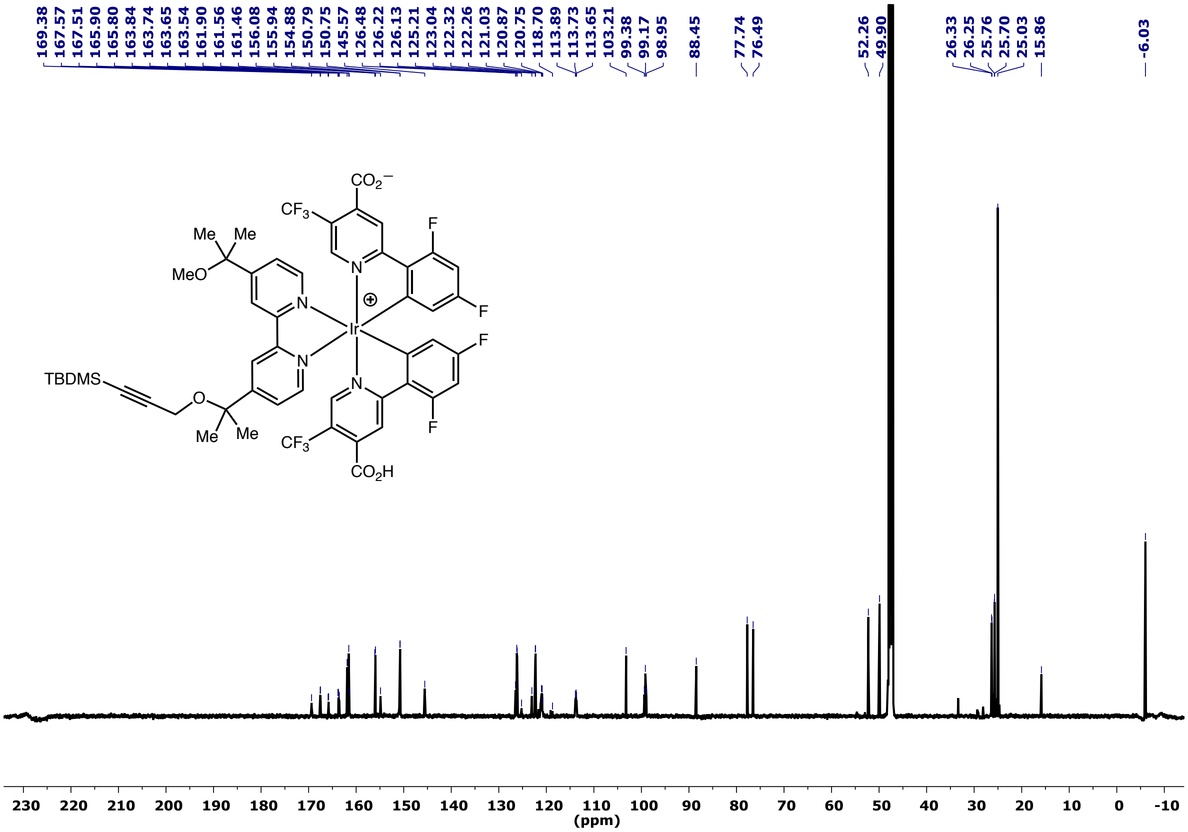

**Ir-TBS-alkyne: $^{19}$F NMR**

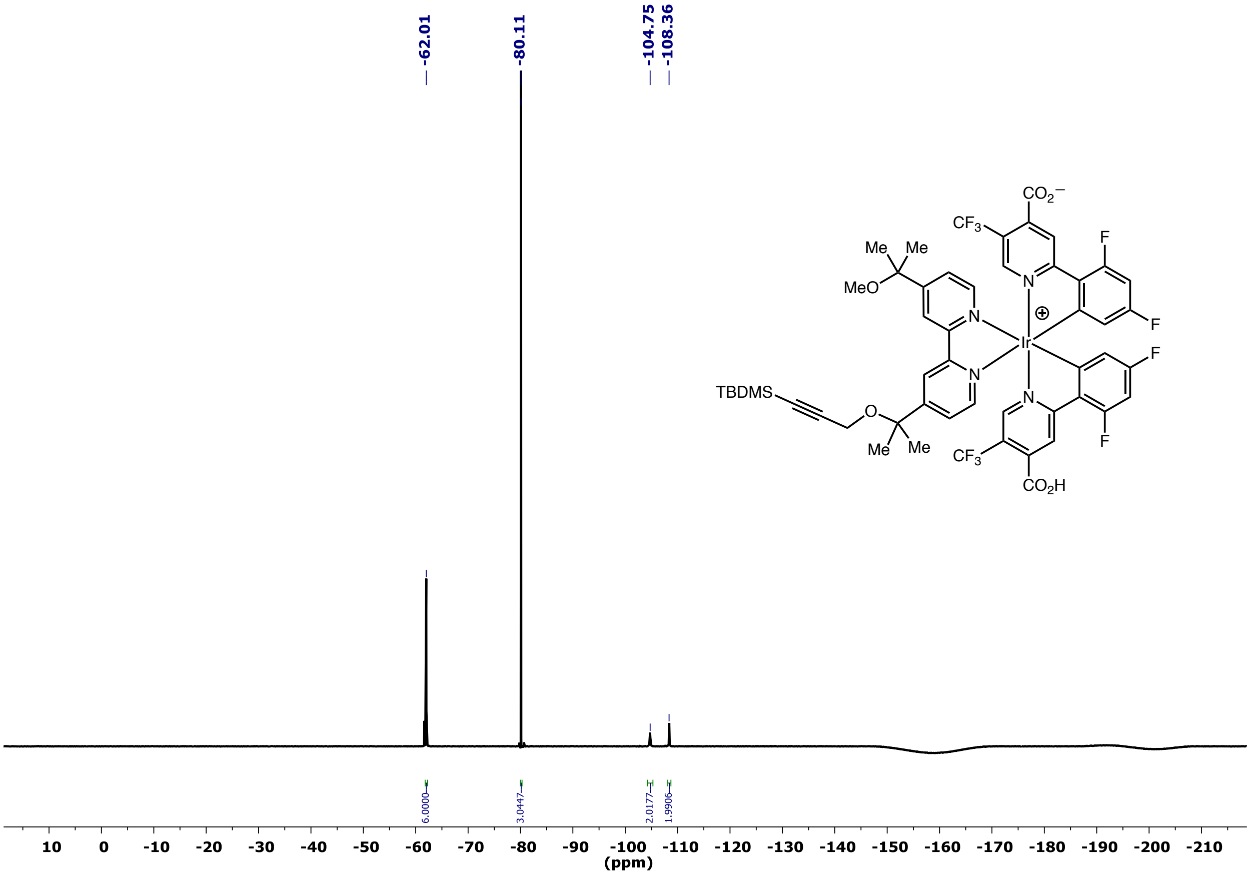

### Ir-alkyne: $^1$H NMR

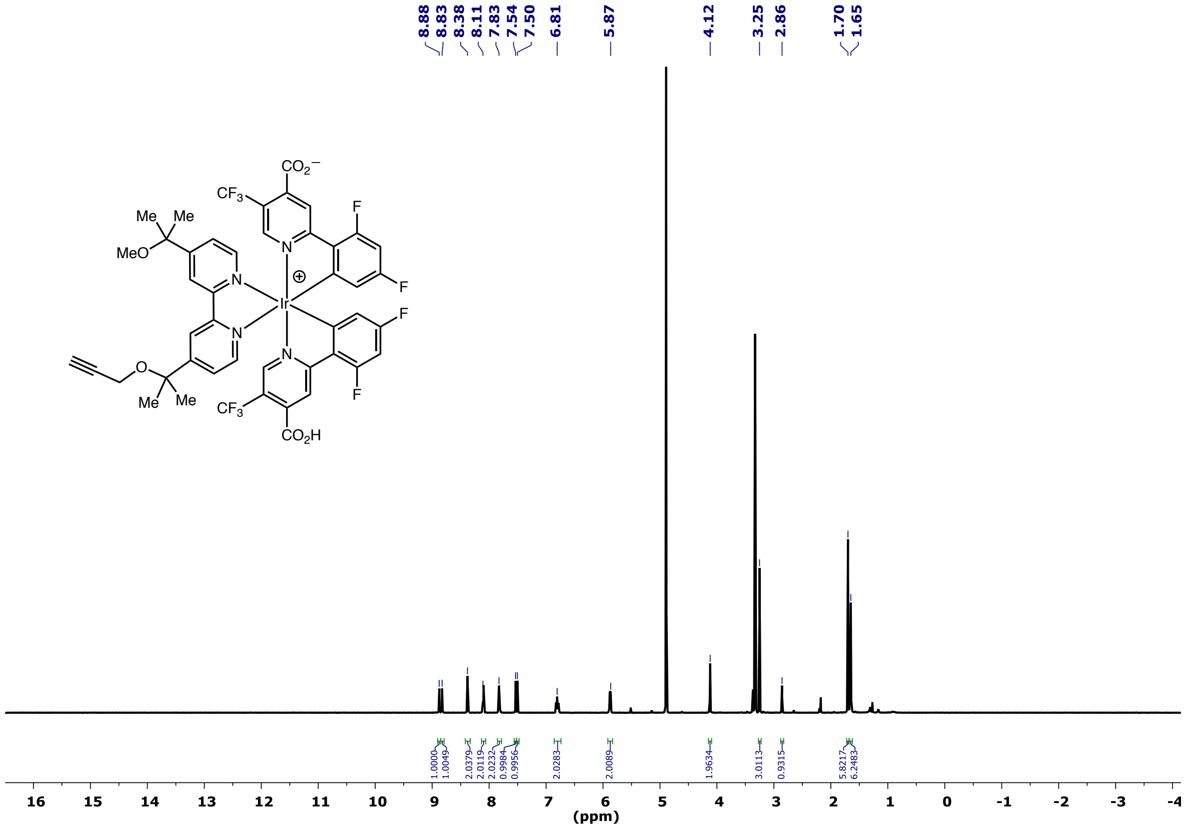

### Ir-alkyne: $^{13}C$ NMR

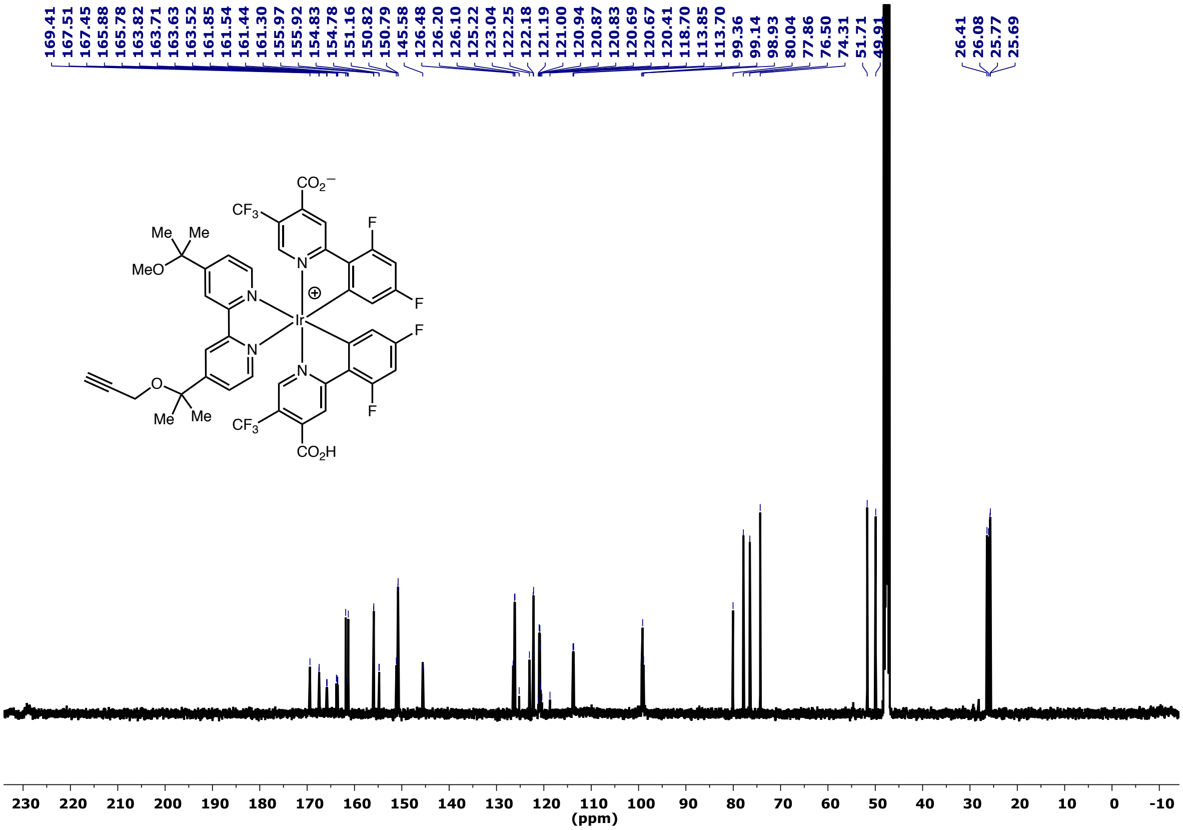

### Ir-alkyne: $^{19}$F NMR

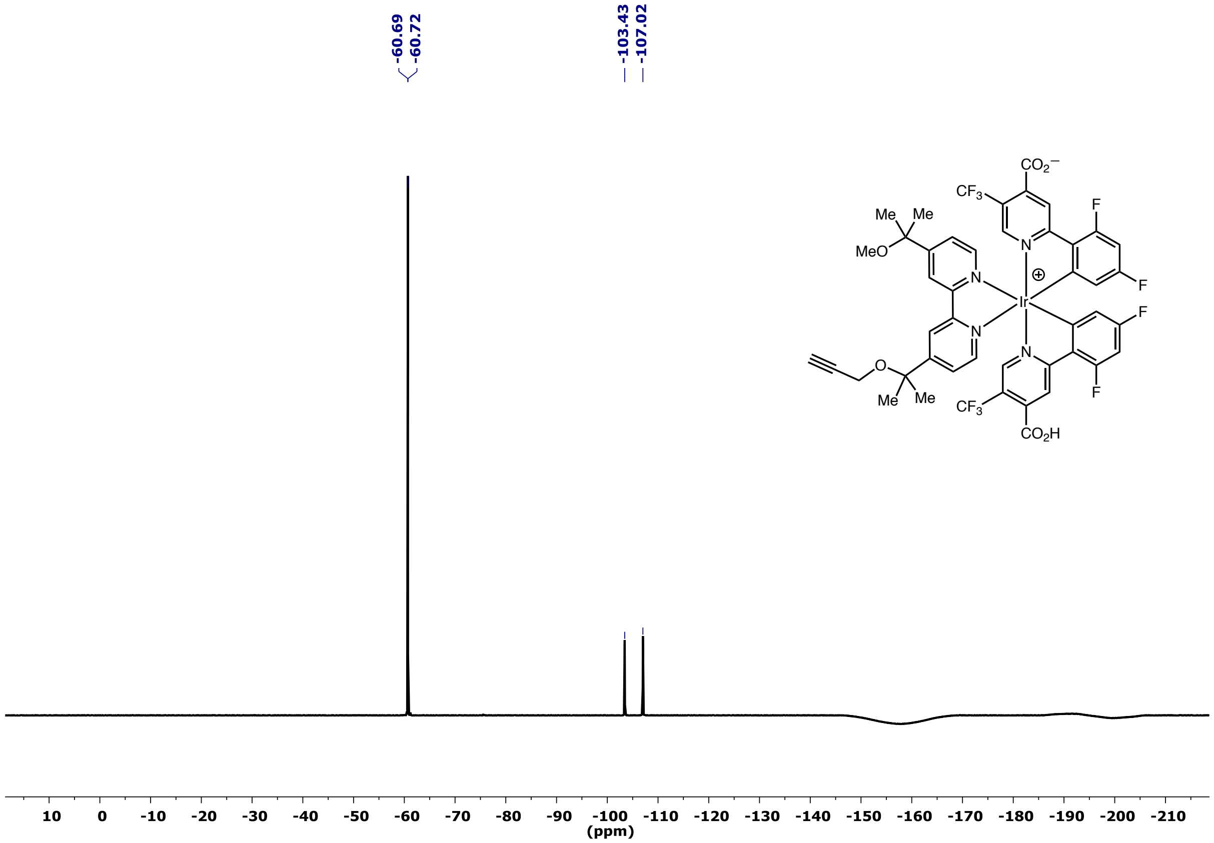

**Ir(dCO$_2$HdFCF$_3$ppy)$_2$(diolbppy): $^1$H NMR**

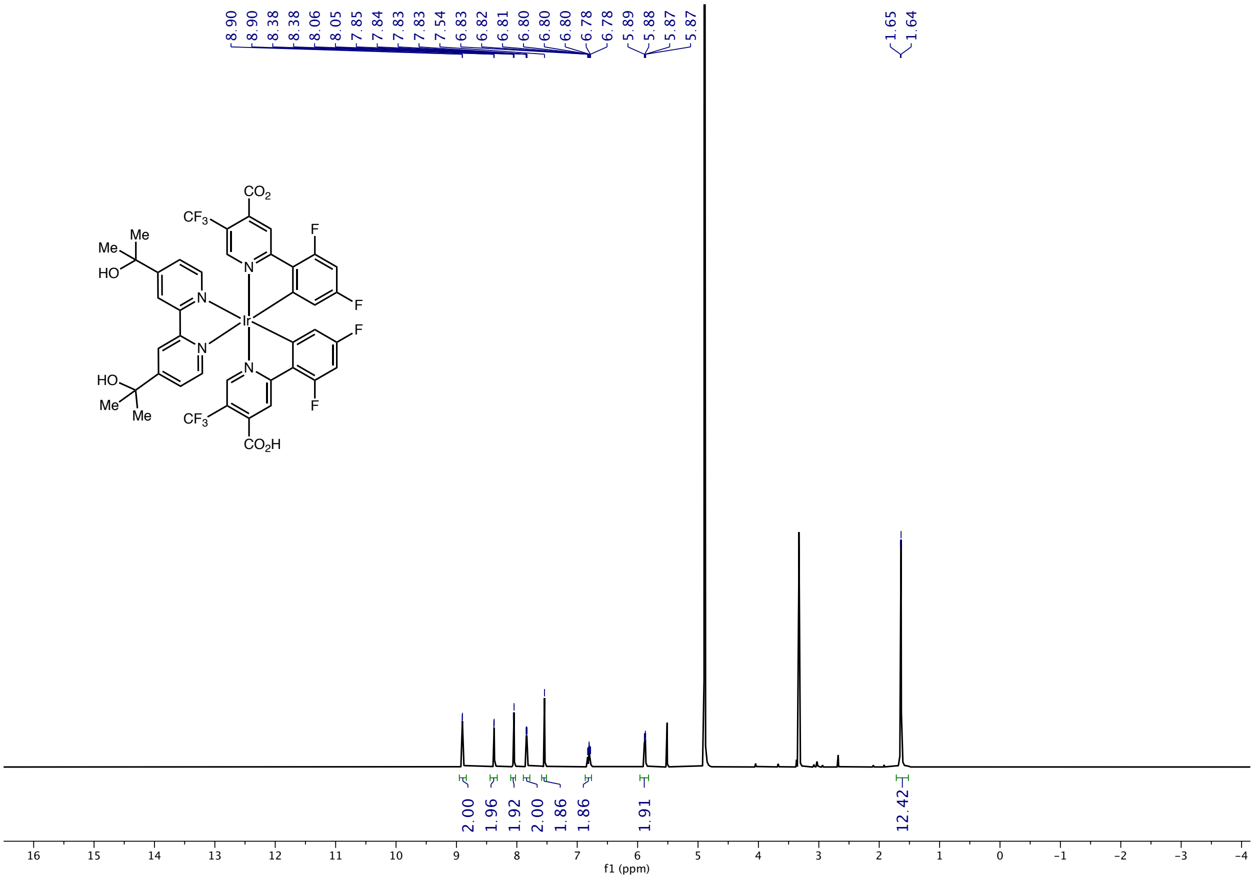

### Ir(dCO$_2$HdFCF$_3$ppy)$_2$(diolbppy): $^{13}$C NMR

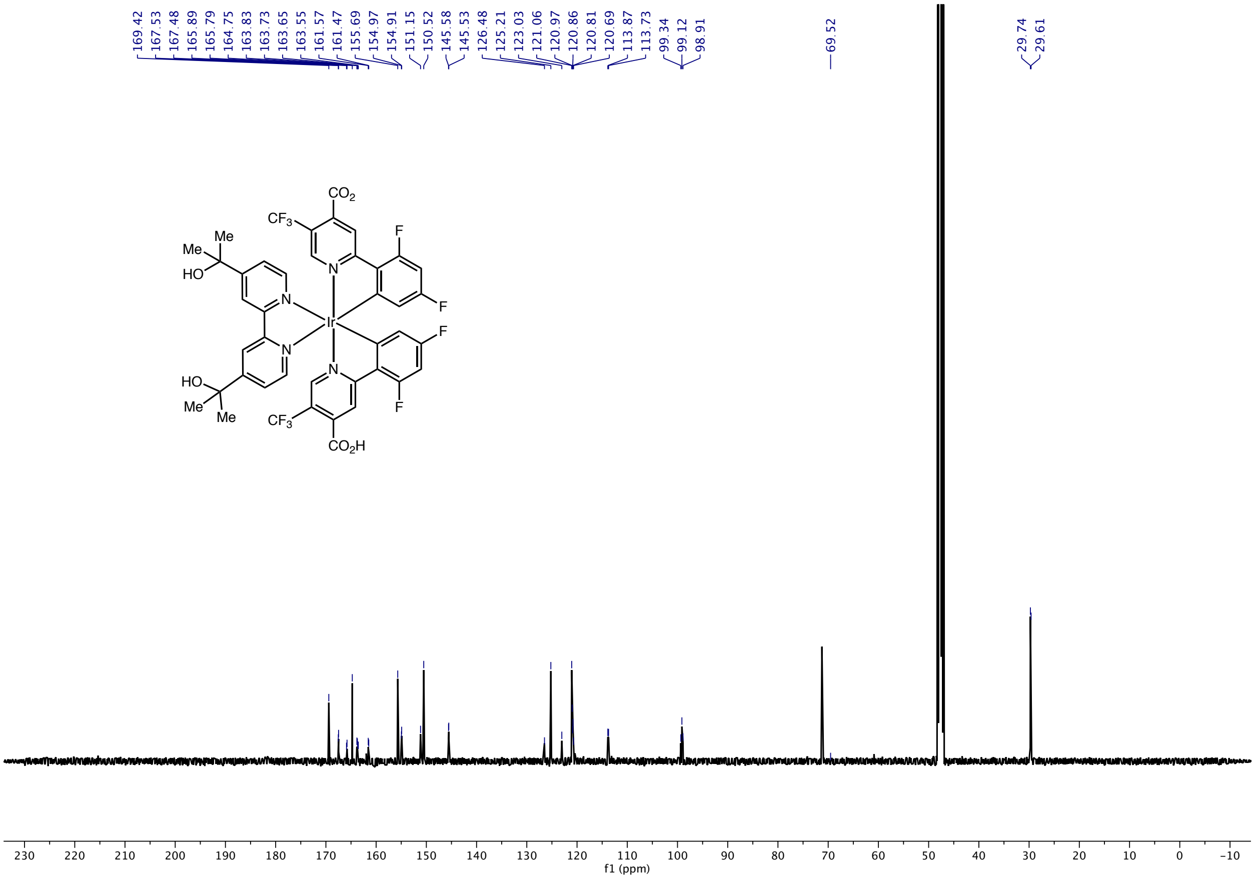

### Ir(dCO$_2$HdFCF$_3$ppy)$_2$(diolbppy): $^{19}$F NMR

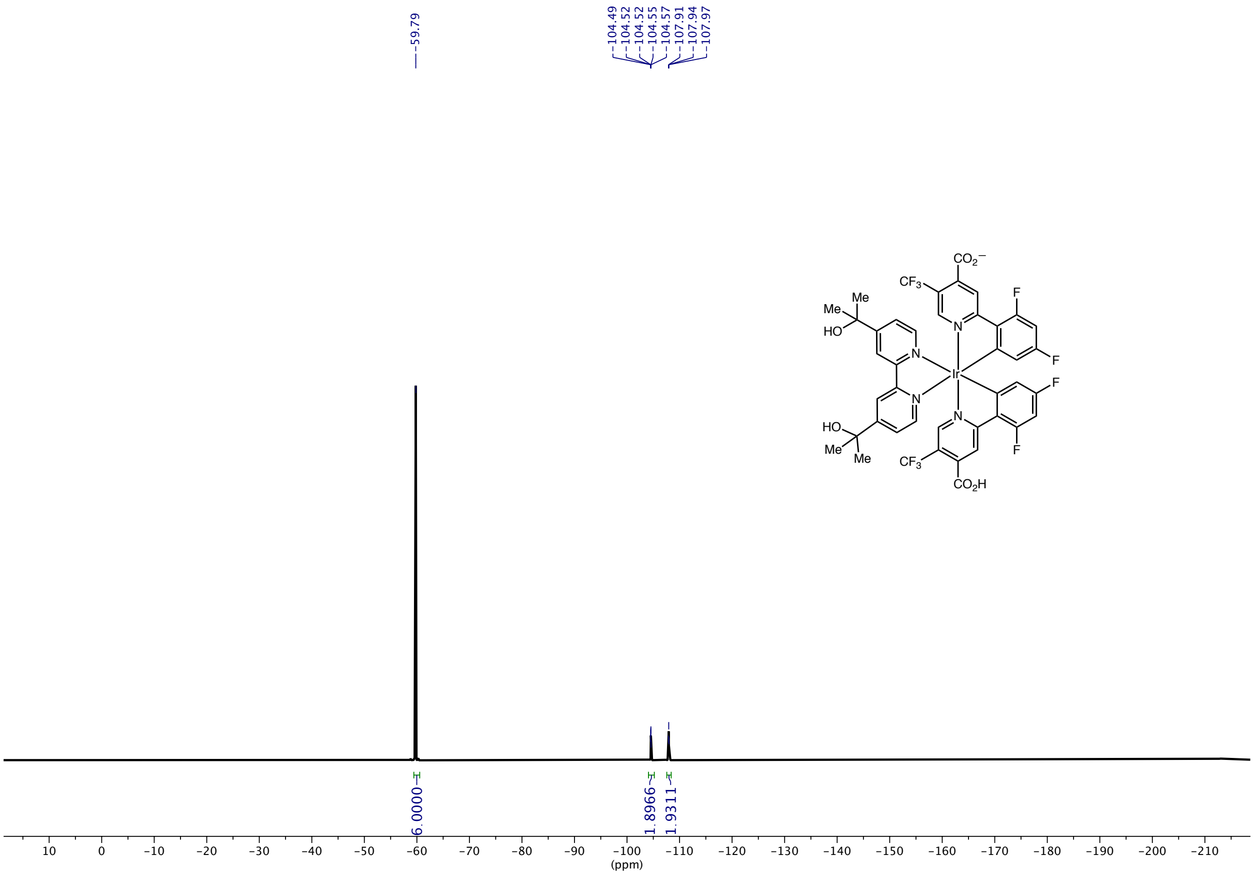

#### Ethyl 3-(4'-methyl-[2,2'-bipyridin]-4-yl)propanoate: $^{1}$H NMR

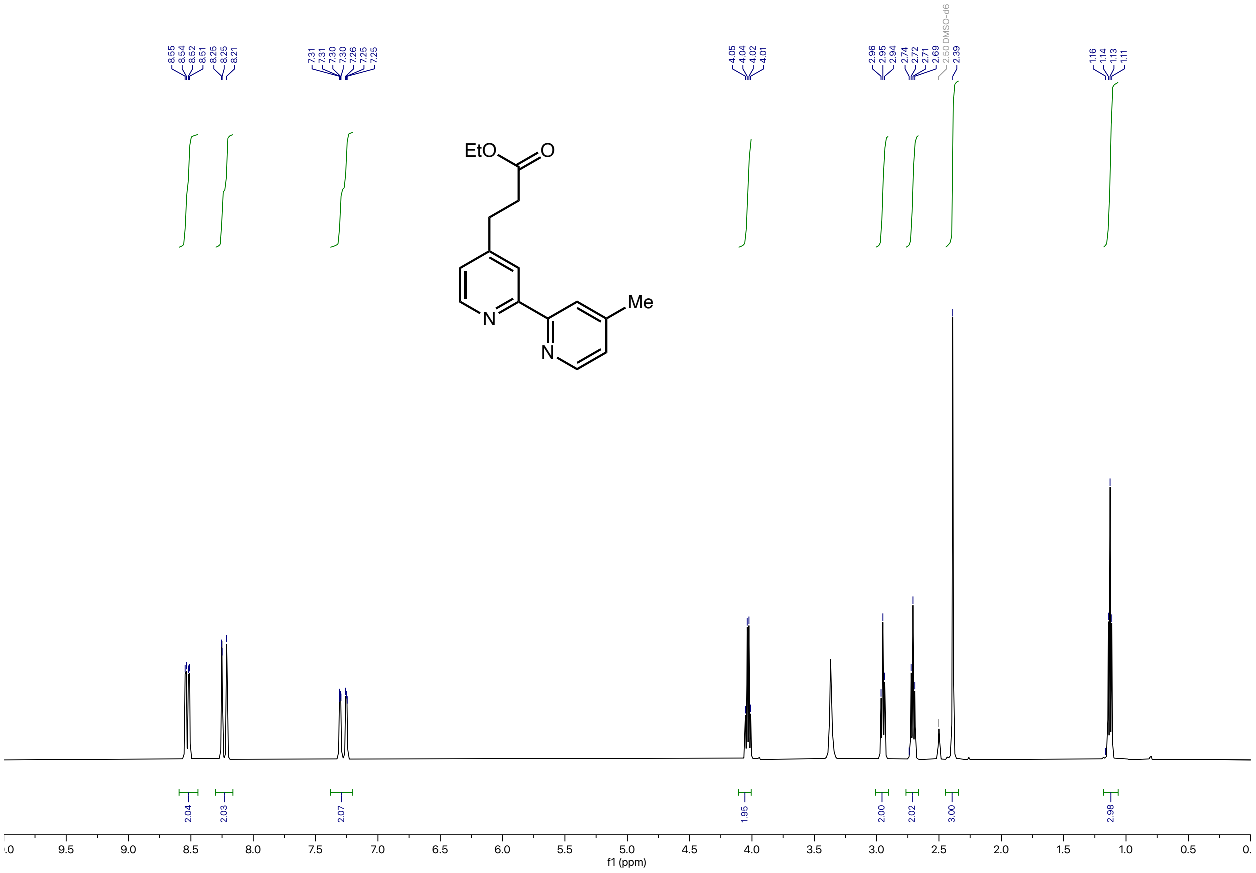

### Ethyl 3-(4'-methyl-[2,2'-bipyridin]-4-yl)propanoate: $^{13}$C NMR

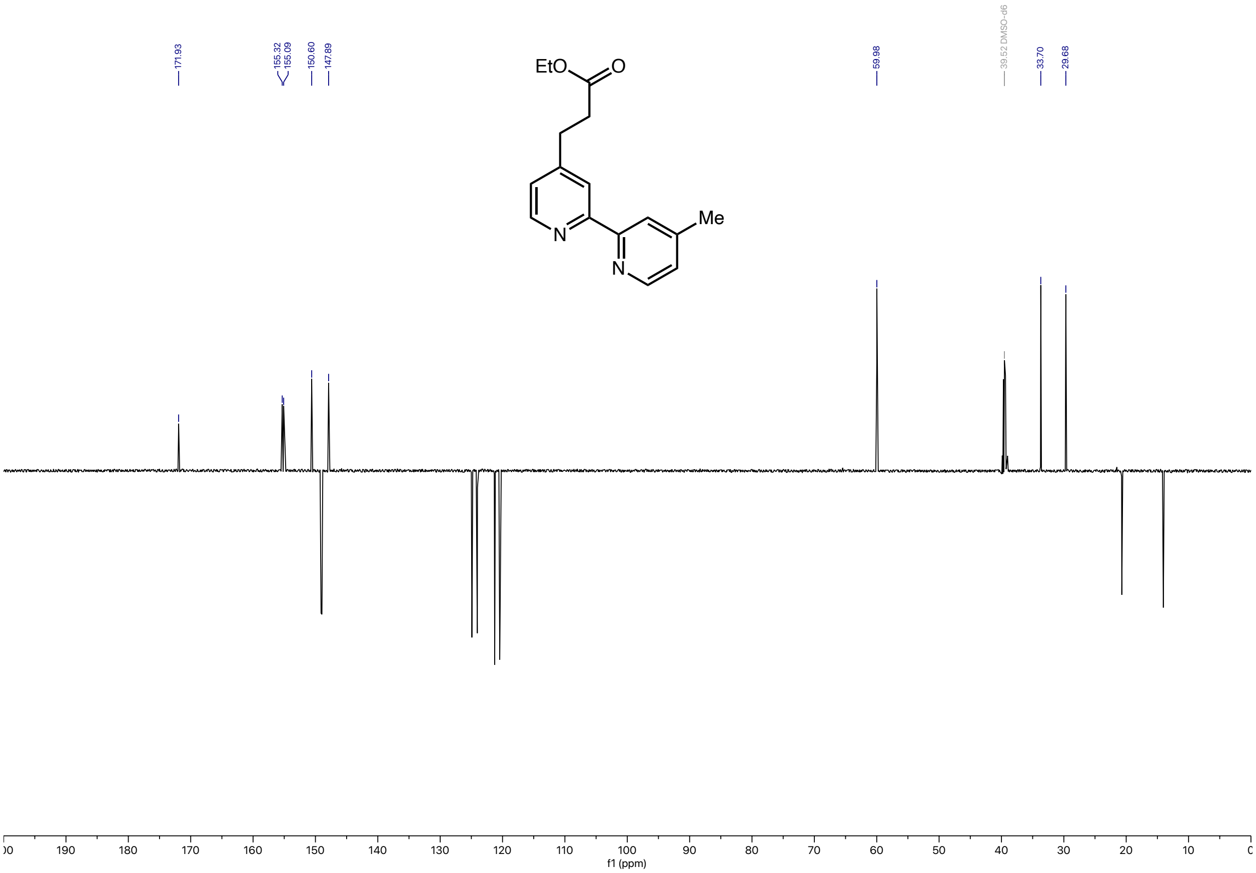

### 3-(4'-methyl-[2,2'-bipyridin]-4-yl)propanoic acid: $^{1}$H NMR

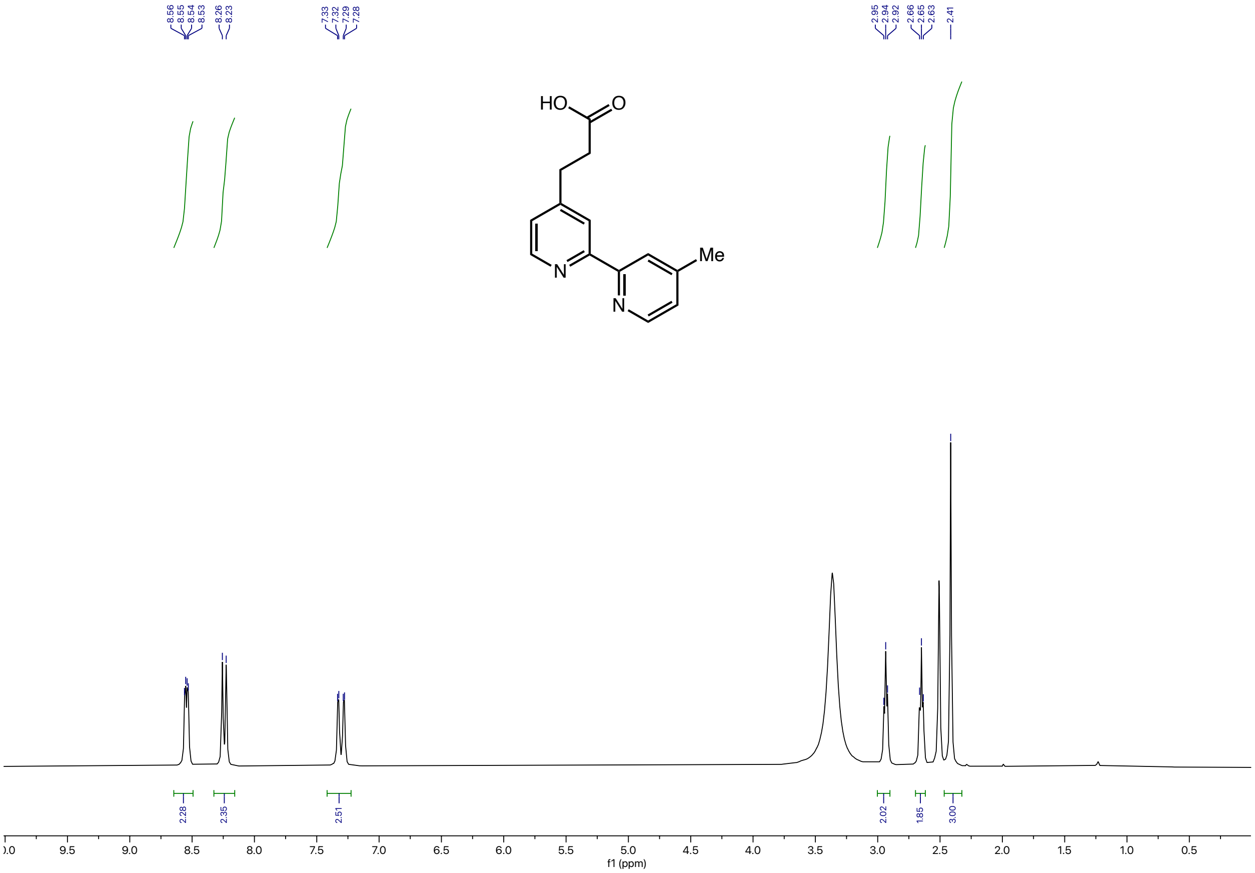

### 3-(4'-methyl-[2,2'-bipyridin]-4-yl)propanoic acid: $^{13}$C NMR

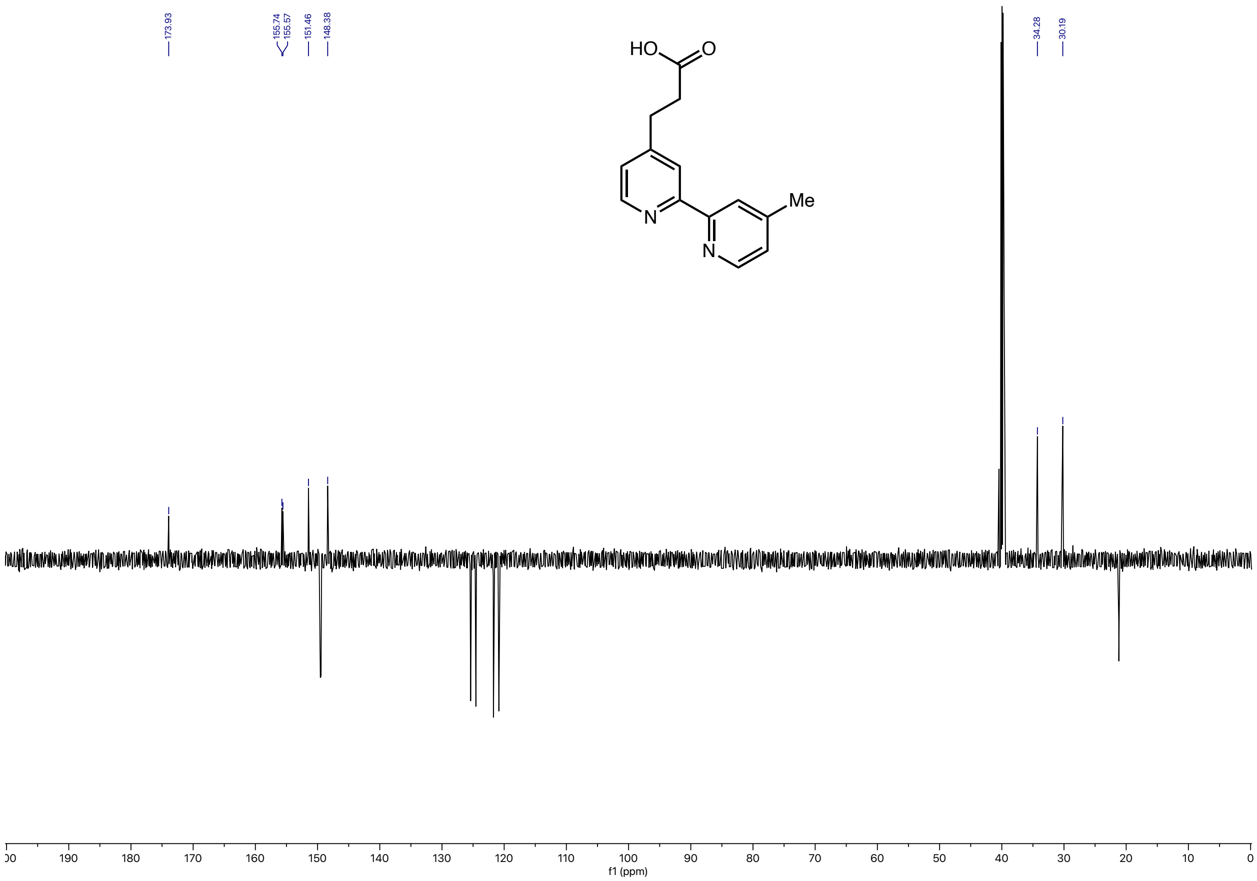

#### Ir-$CO_2$H: [1]H NMR

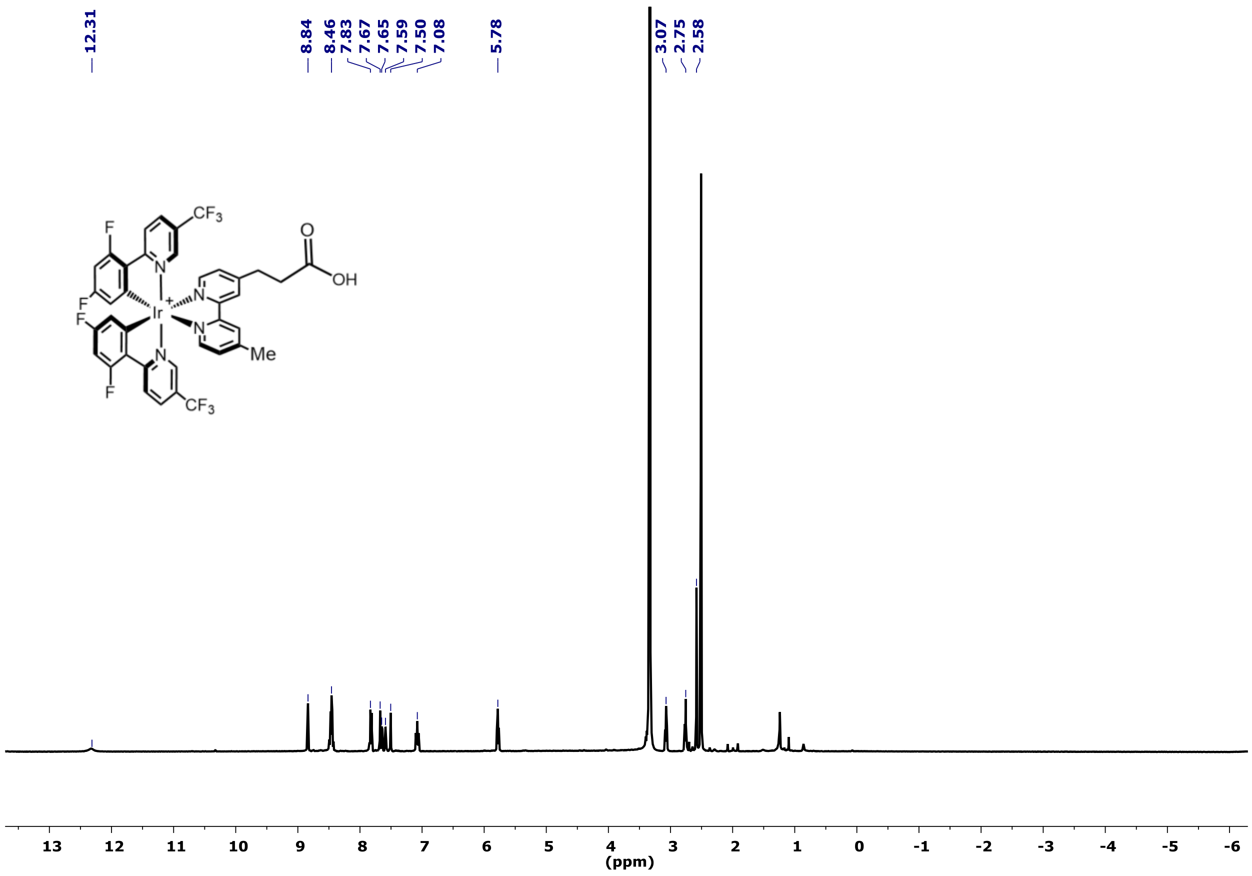

#### Ir-$CO_2$H: $^{13}$C NMR

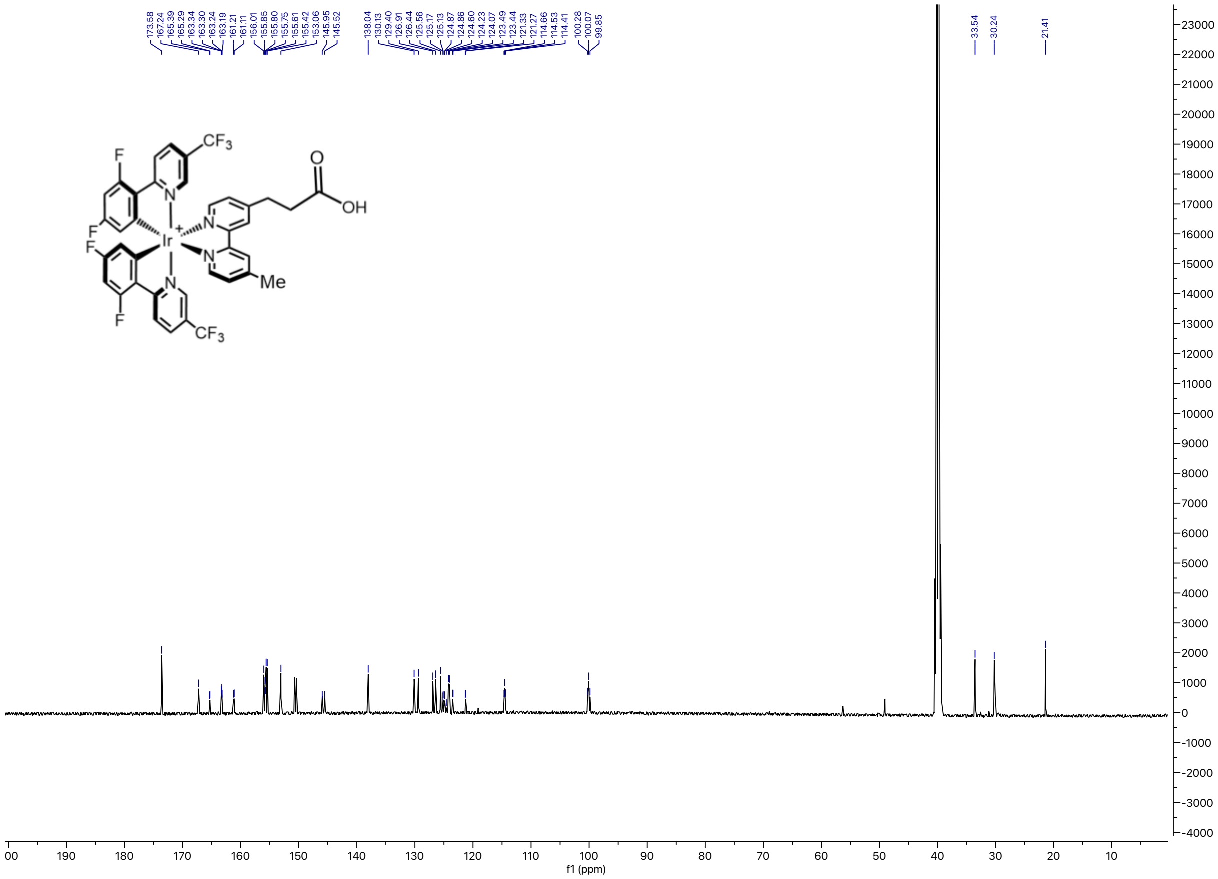

#### Ir-$CO_2$H: $^{19}$F NMR

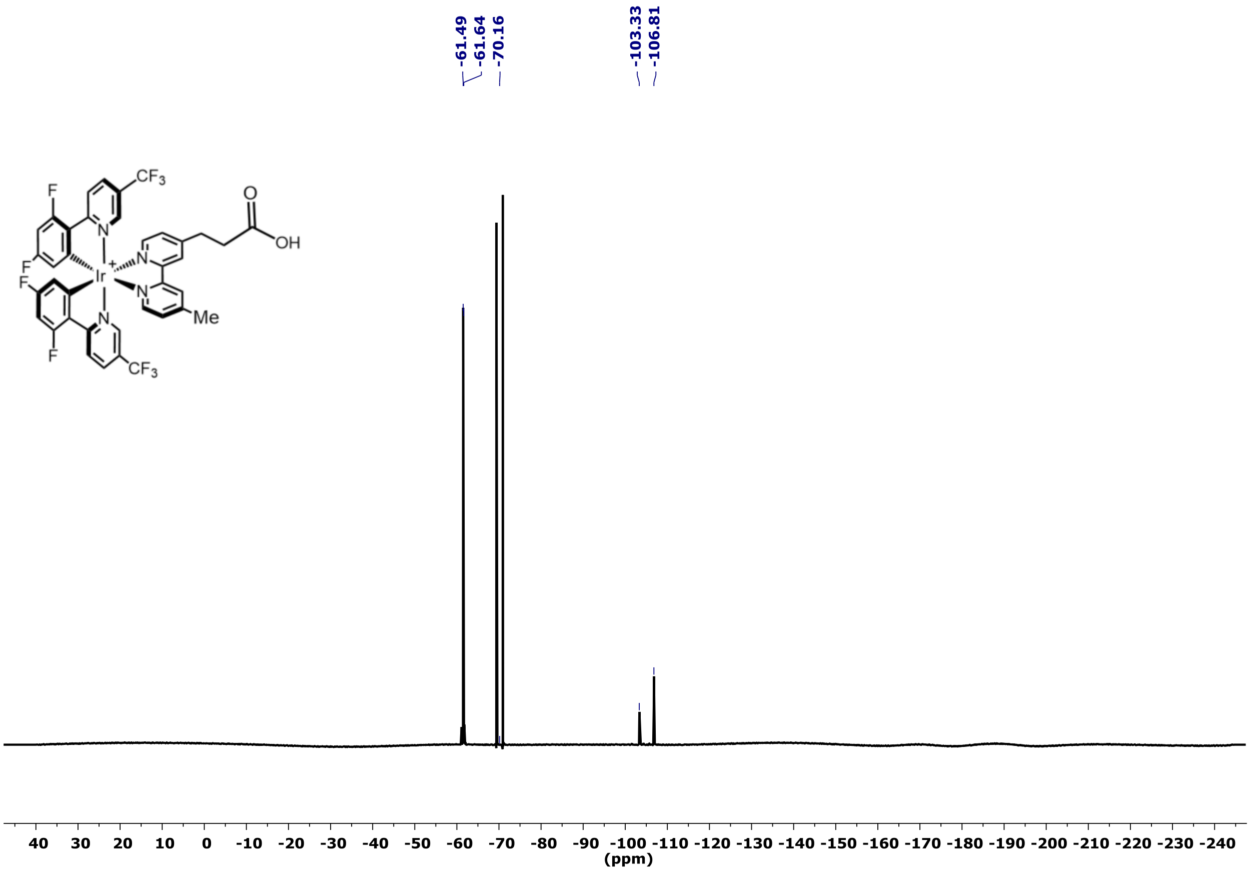

### Ir-$CO_2$H: [1]H-[1]H COSY NMR

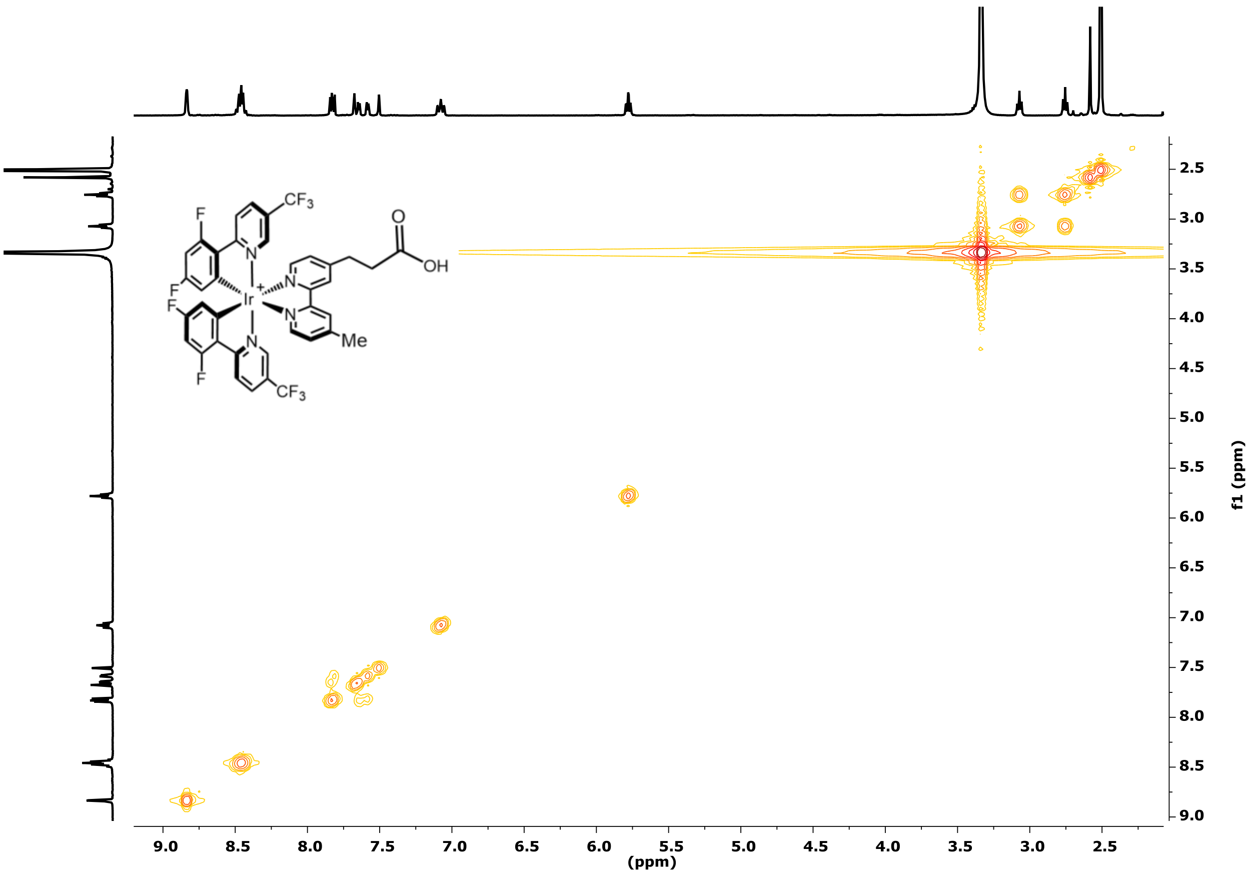

### Ir-PEG3-NHBoc: $^{1}H$ NMR

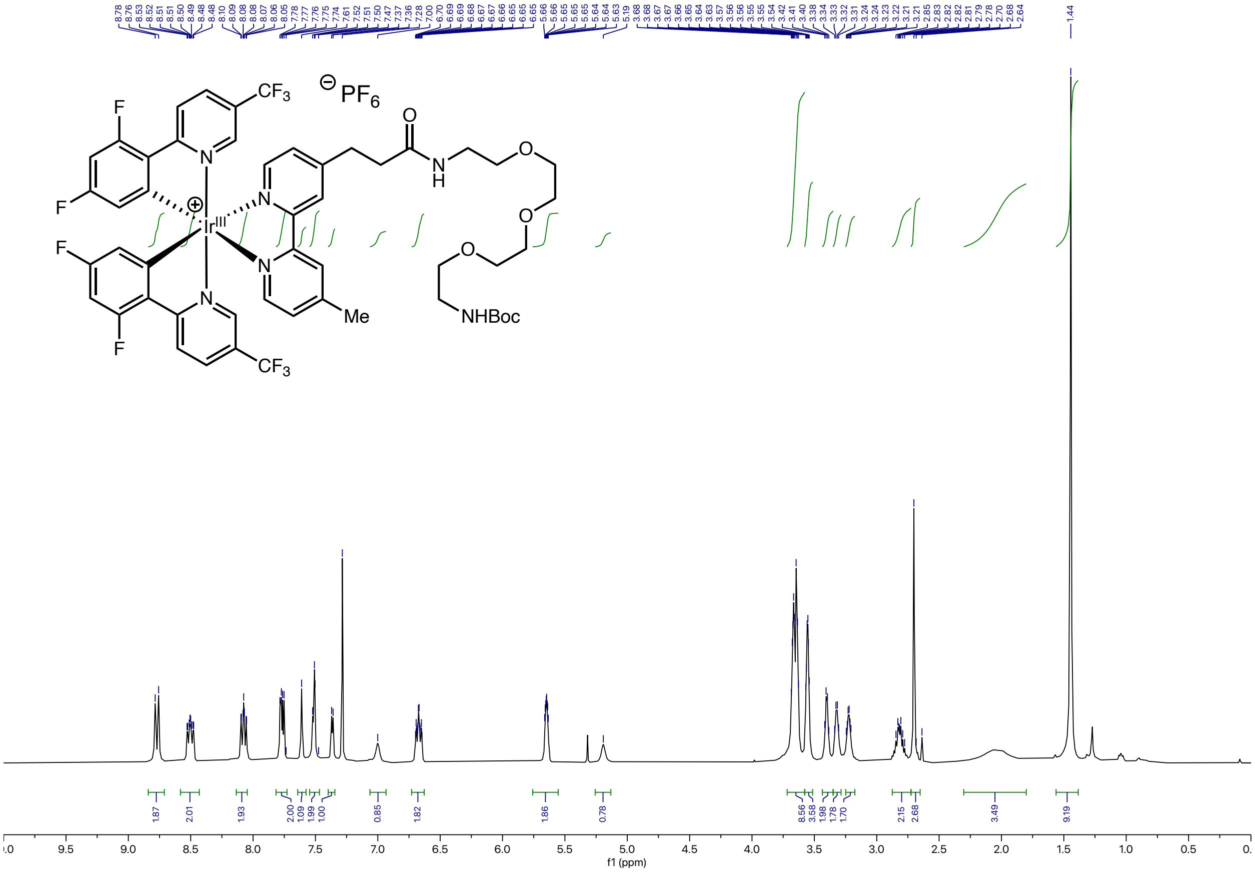

### Ir-PEG3-NHBoc: $^{13}C$ NMR

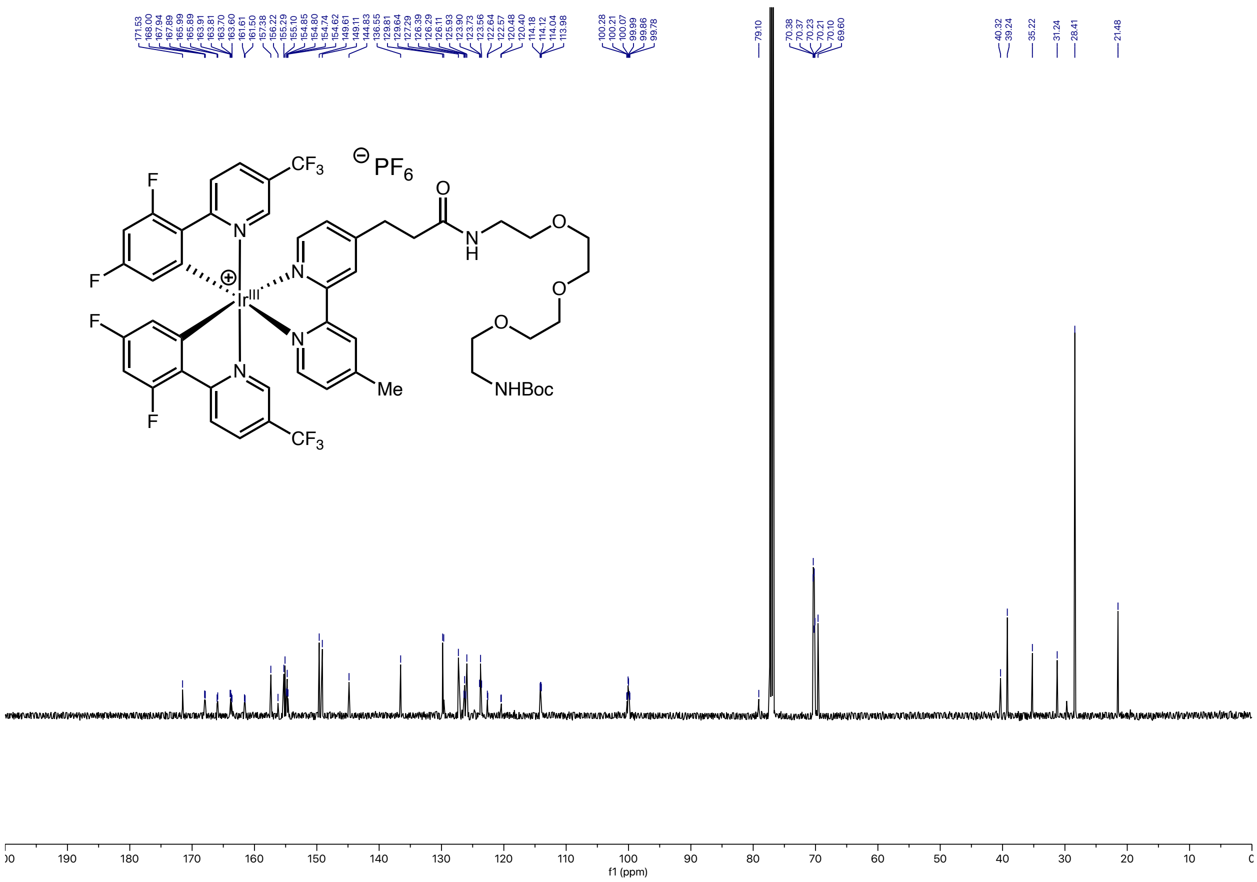

### Ir-PEG3-NHBoc: $^{19}F$ NMR

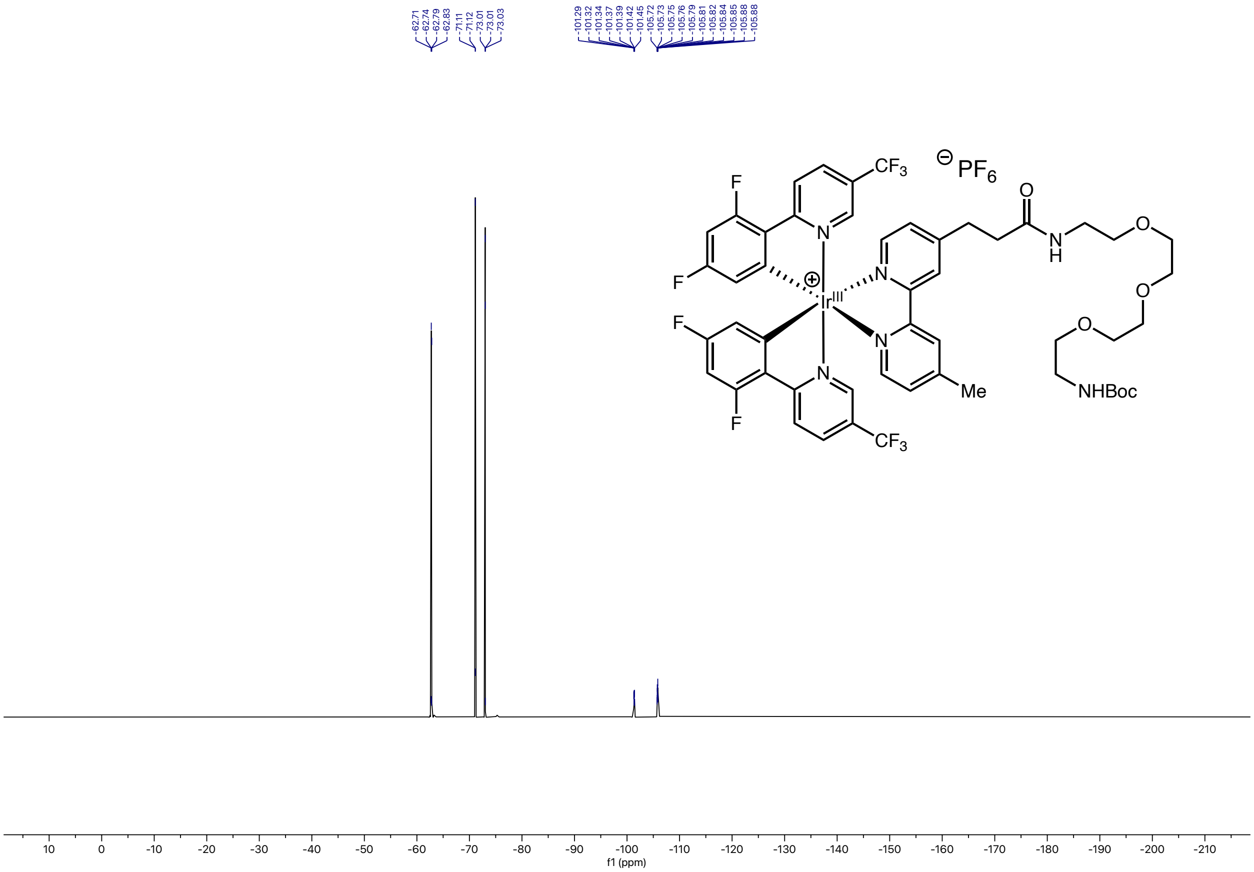

### Ir-G2-NHEt: $^1$H NMR

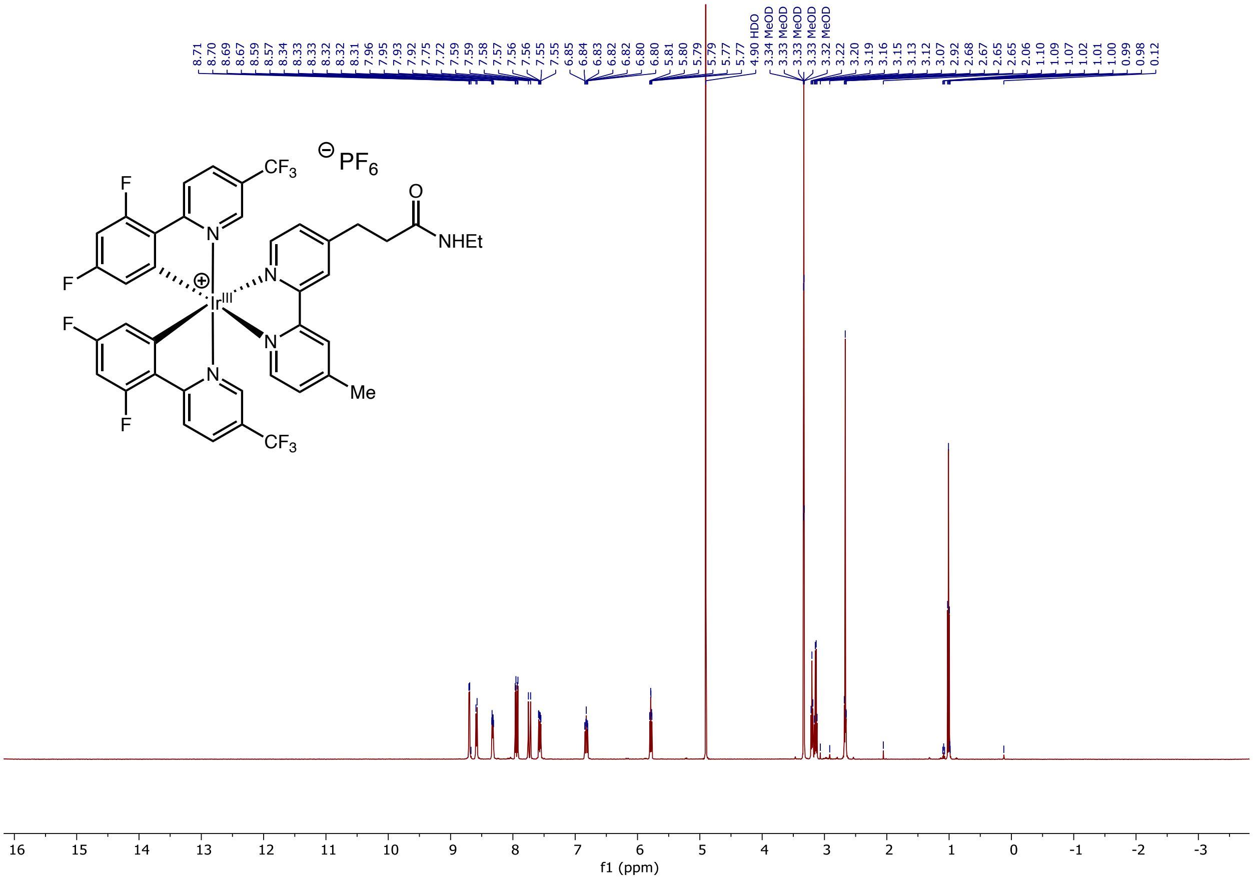

### Ir-G2-NHEt: $^{13}$C NMR

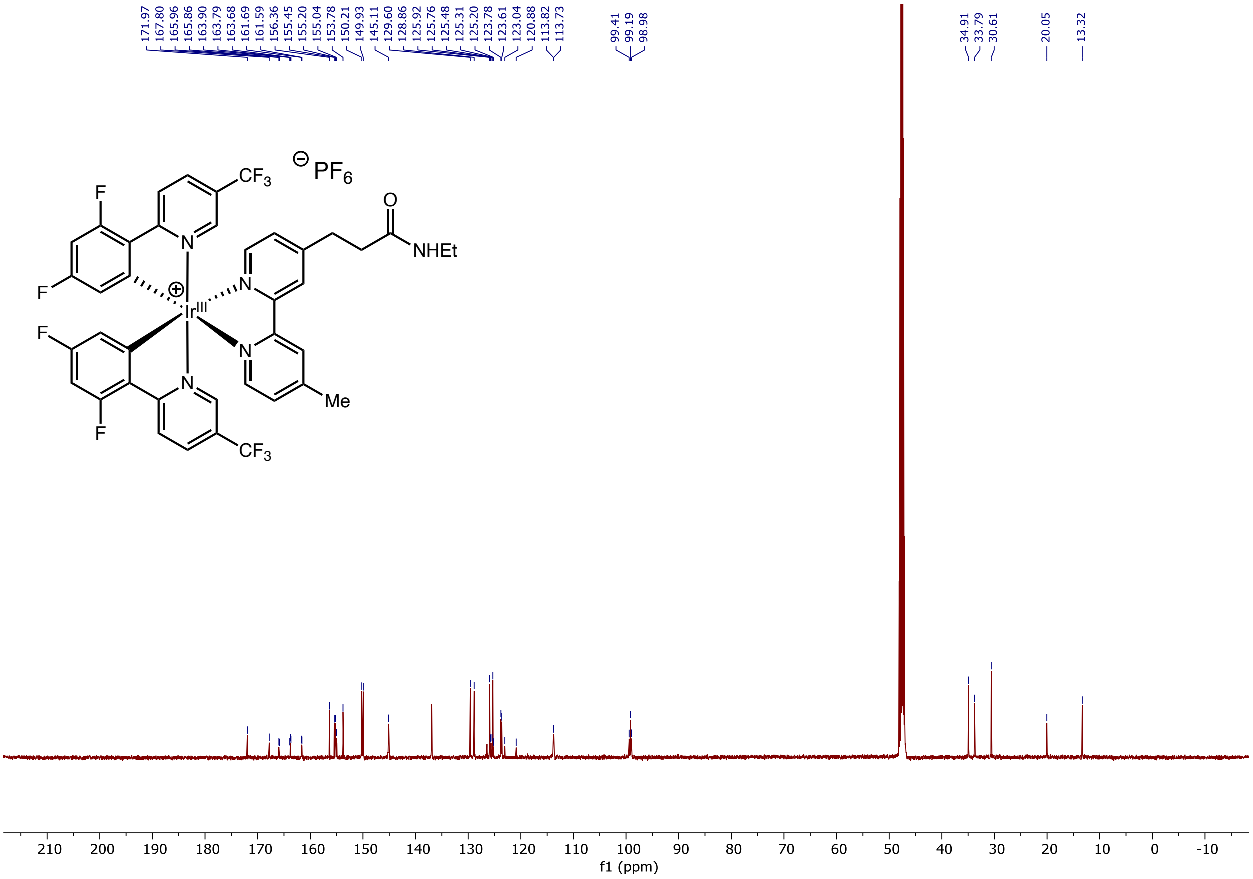

**Ir-G2-NHEt: ¹⁹F NMR**

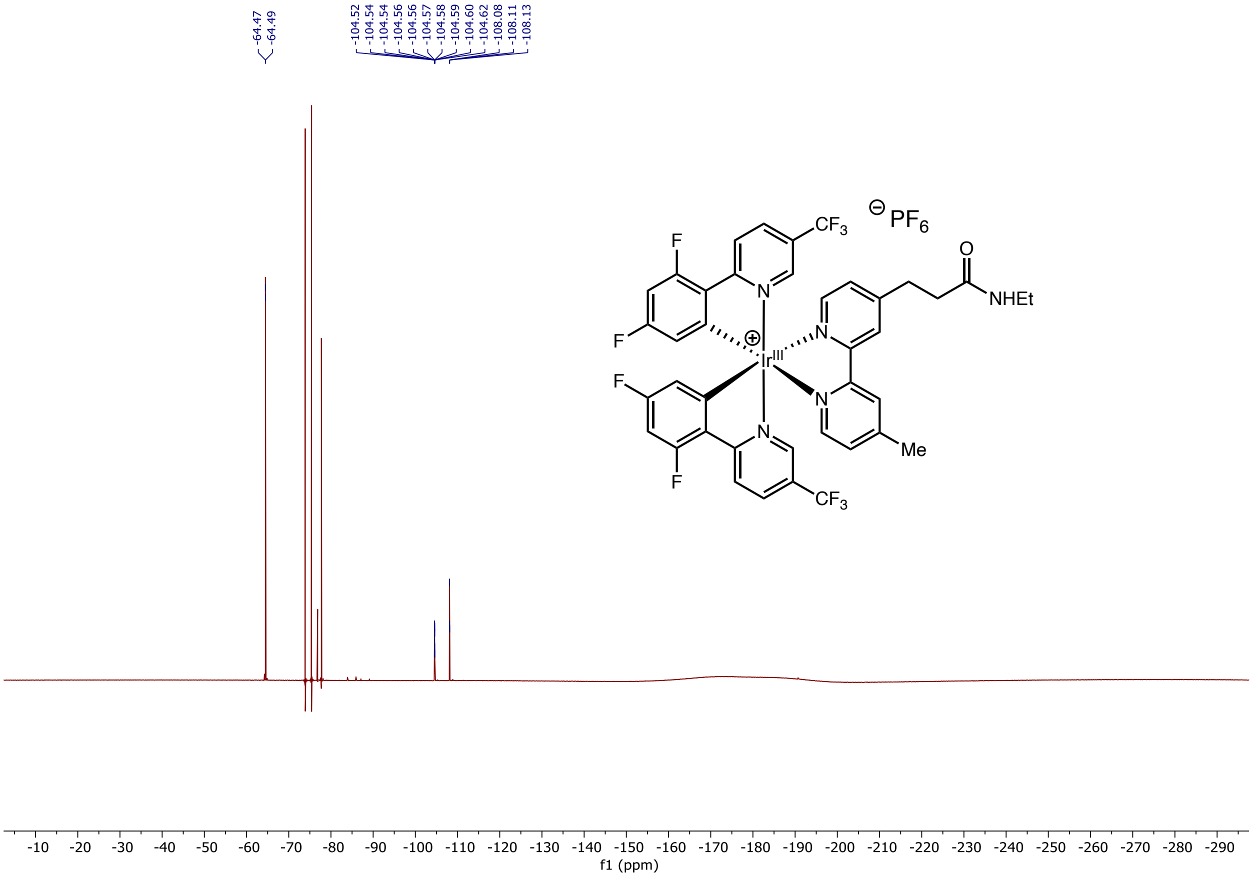

#### Ir-PEG3-CO$_2$H: $^1$H NMR

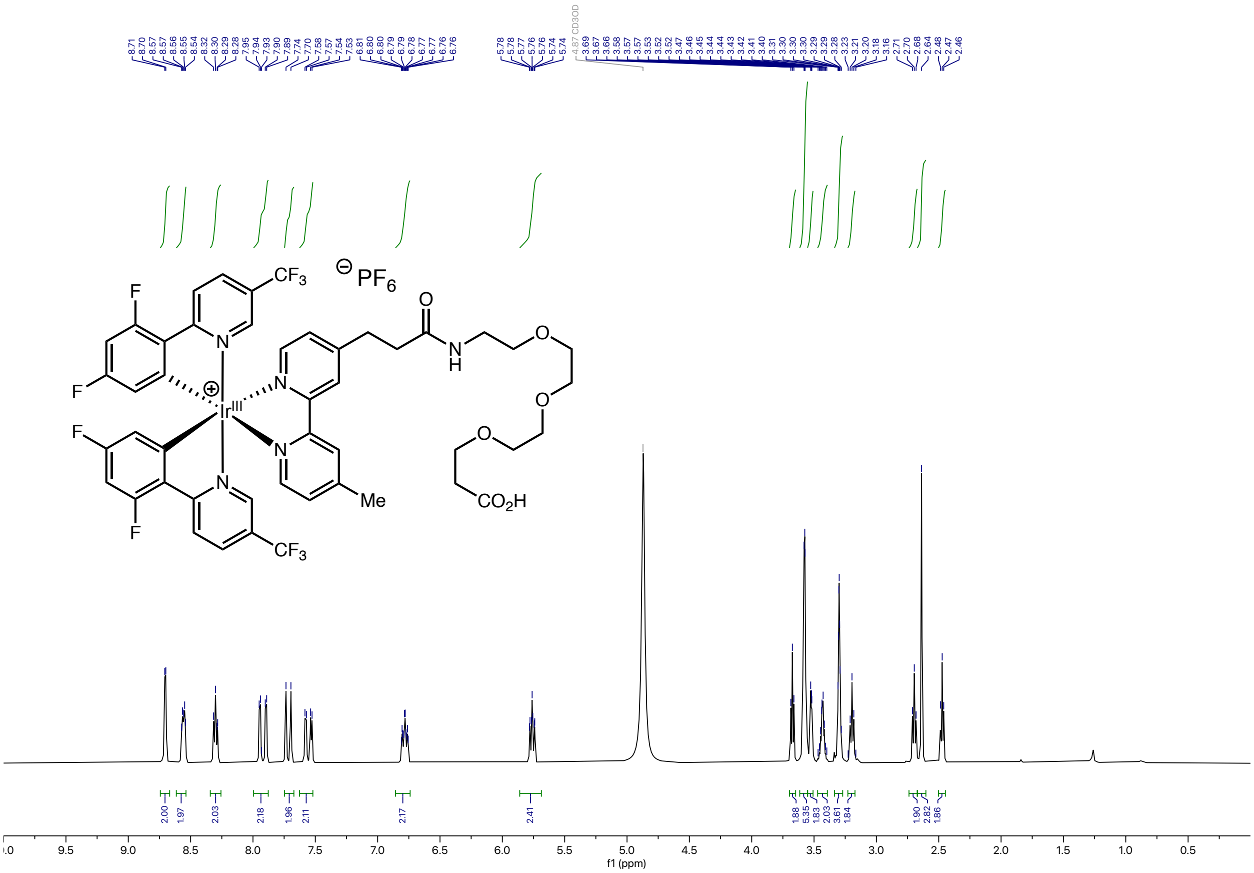

### Ir-PEG3-CO₂H: ¹³C NMR

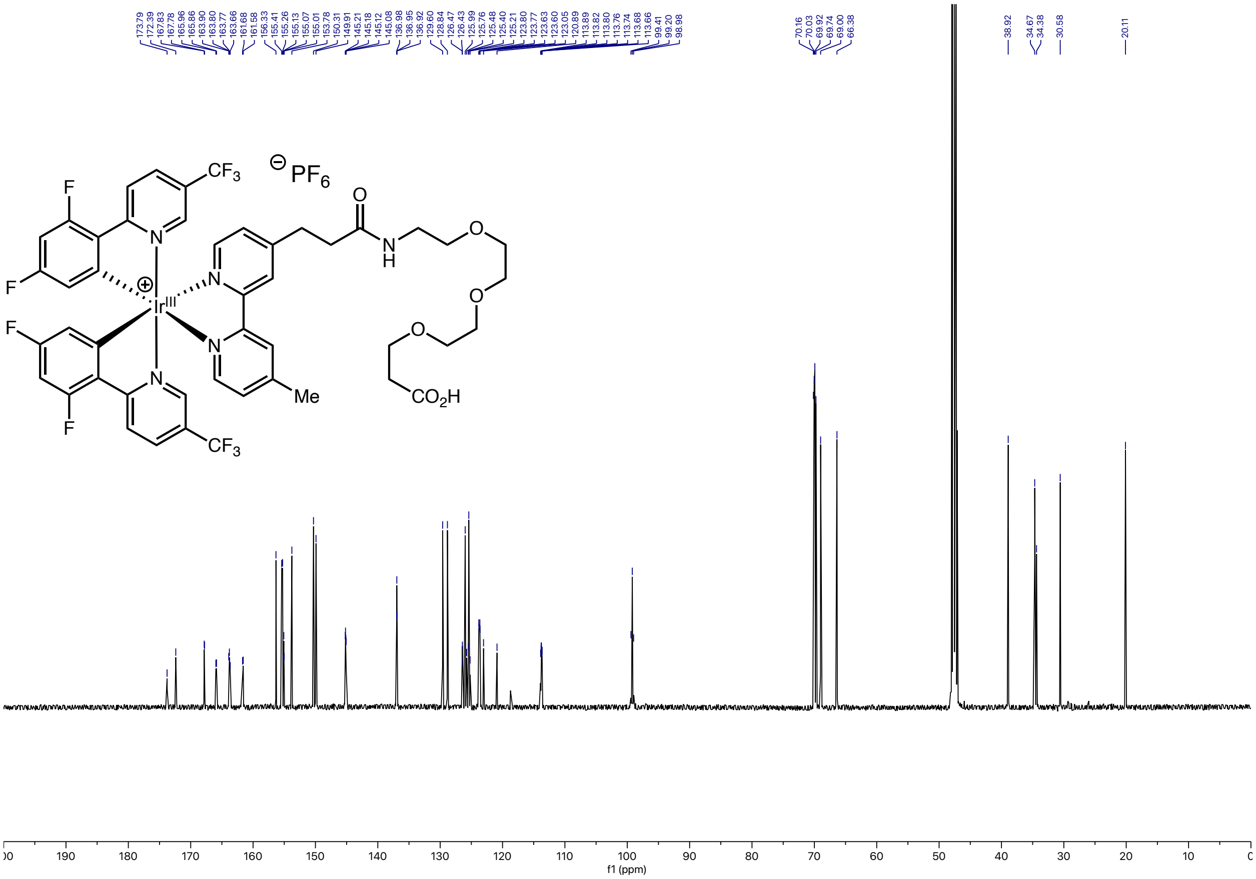

### Ir-PEG3-$CO_2H$: $^{19}F$ NMR

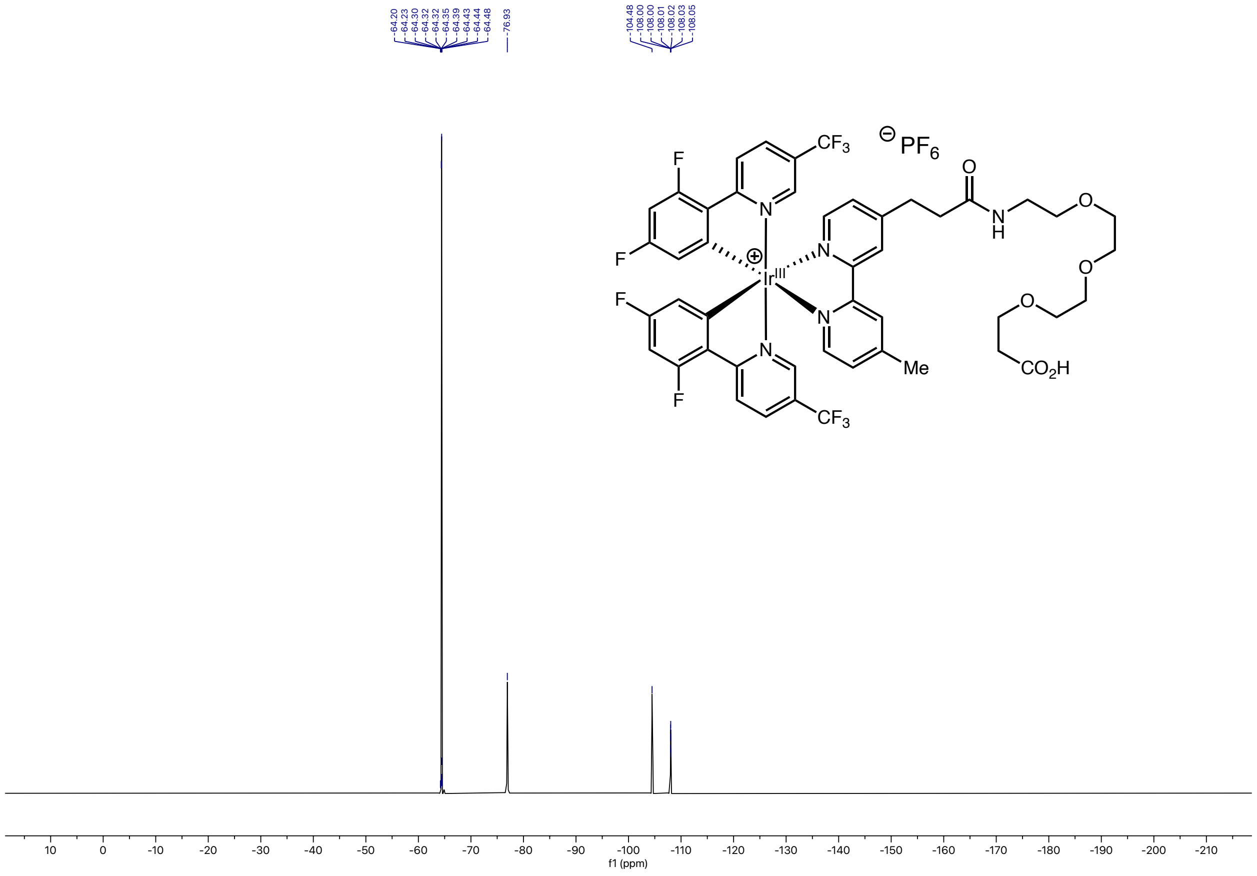

### Ir-PEG4-$C_6H_{12}$Cl: [1]H NMR

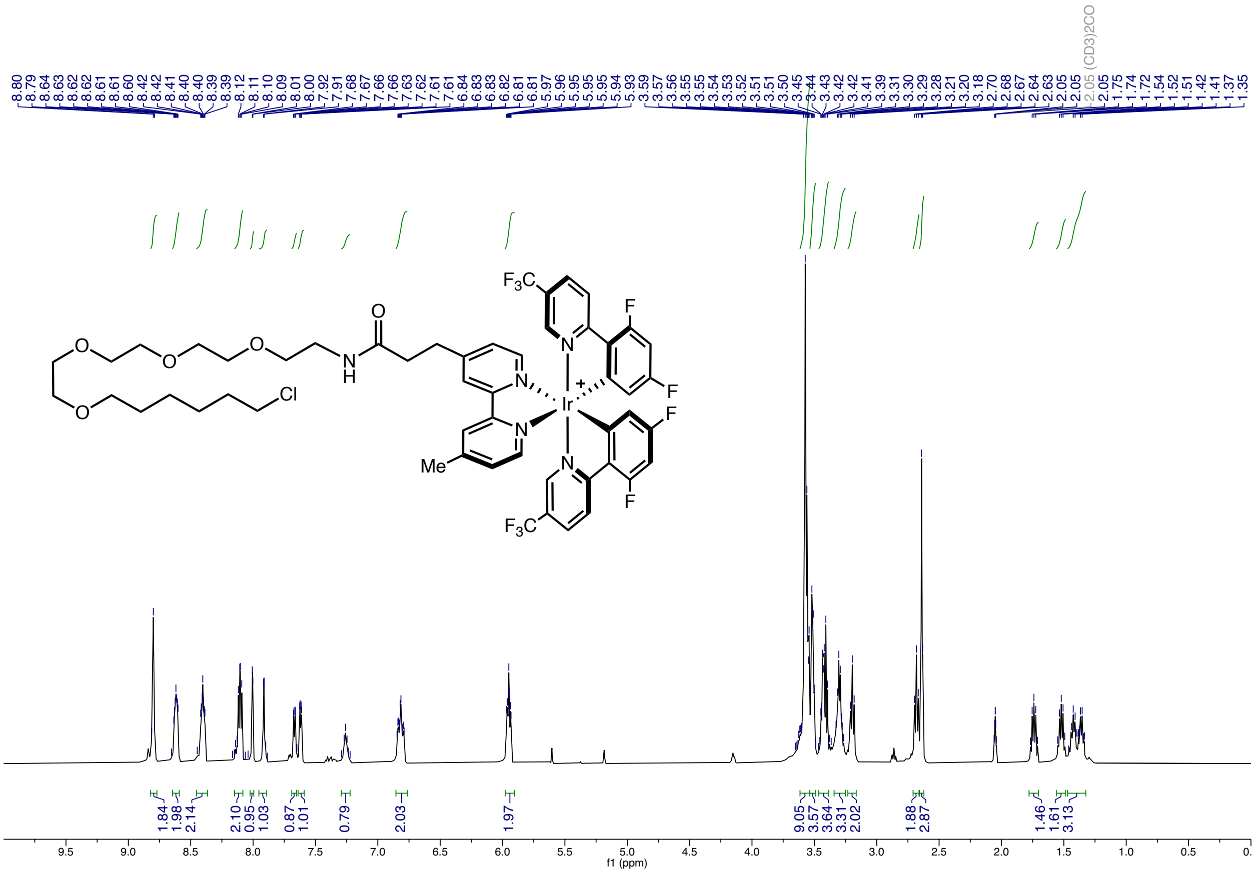

### Ir-PEG4-$C_6H_{12}$Cl: $^{13}$C NMR

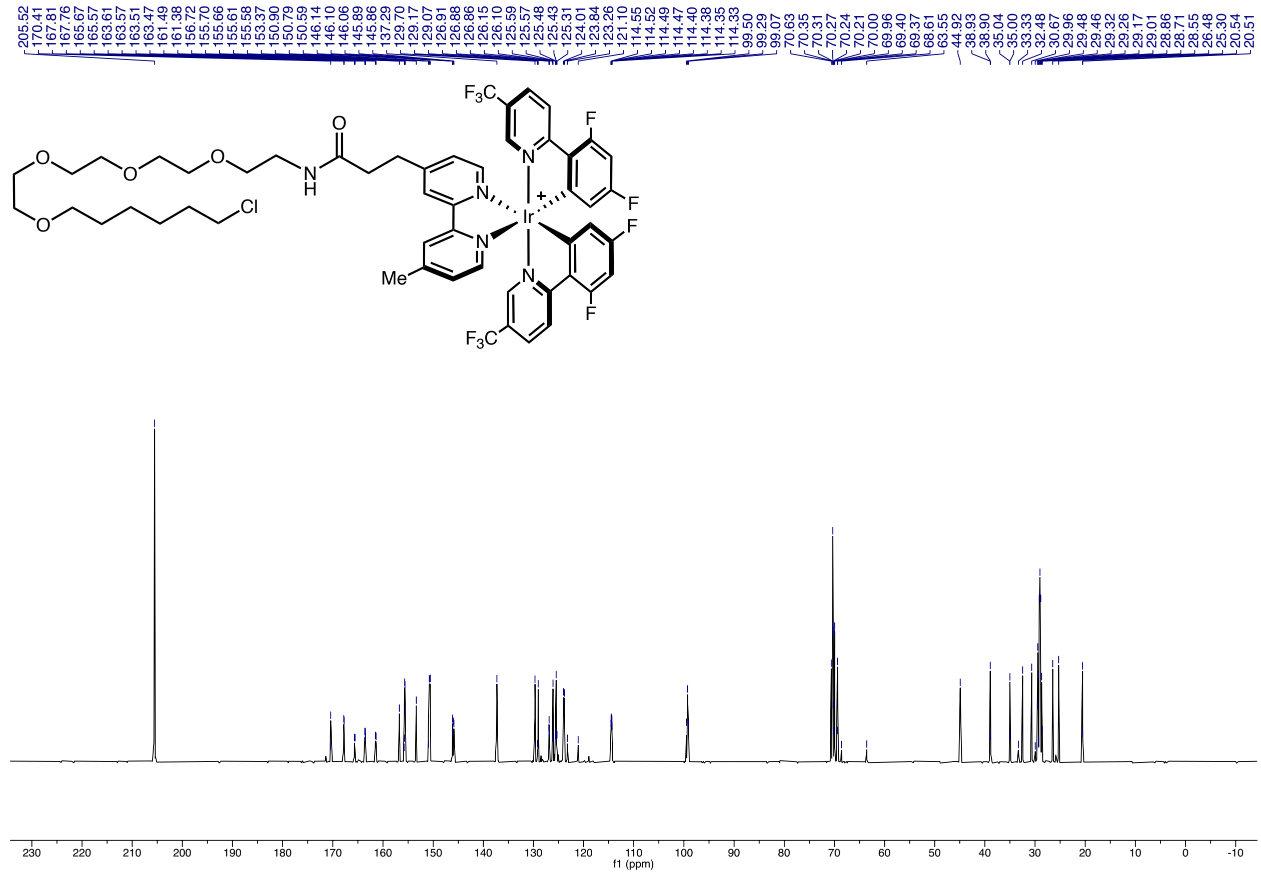

### Ir-PEG4-$C_6H_{12}$Cl: $^{19}$F NMR

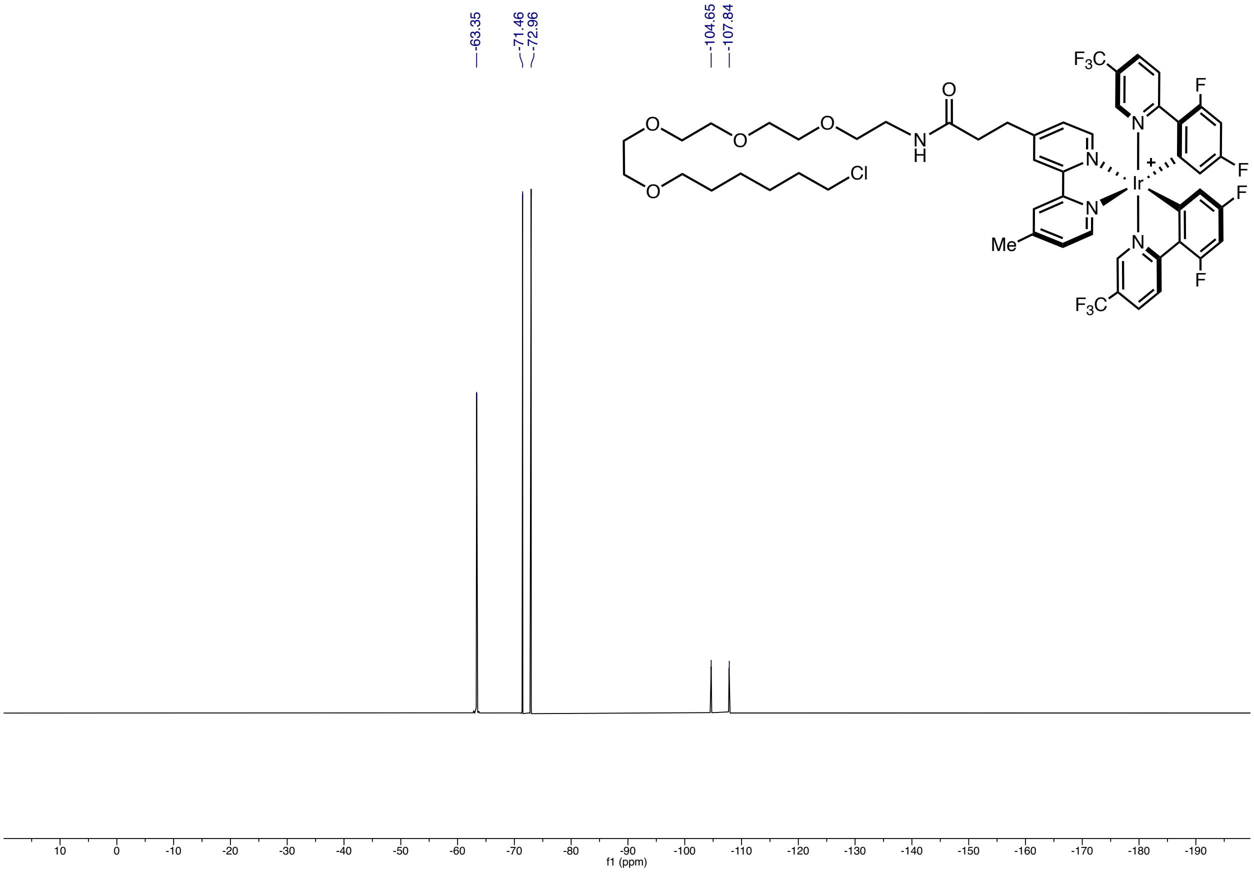

### JQ-1-PEG3-NHBoc: $^{1}H$ NMR

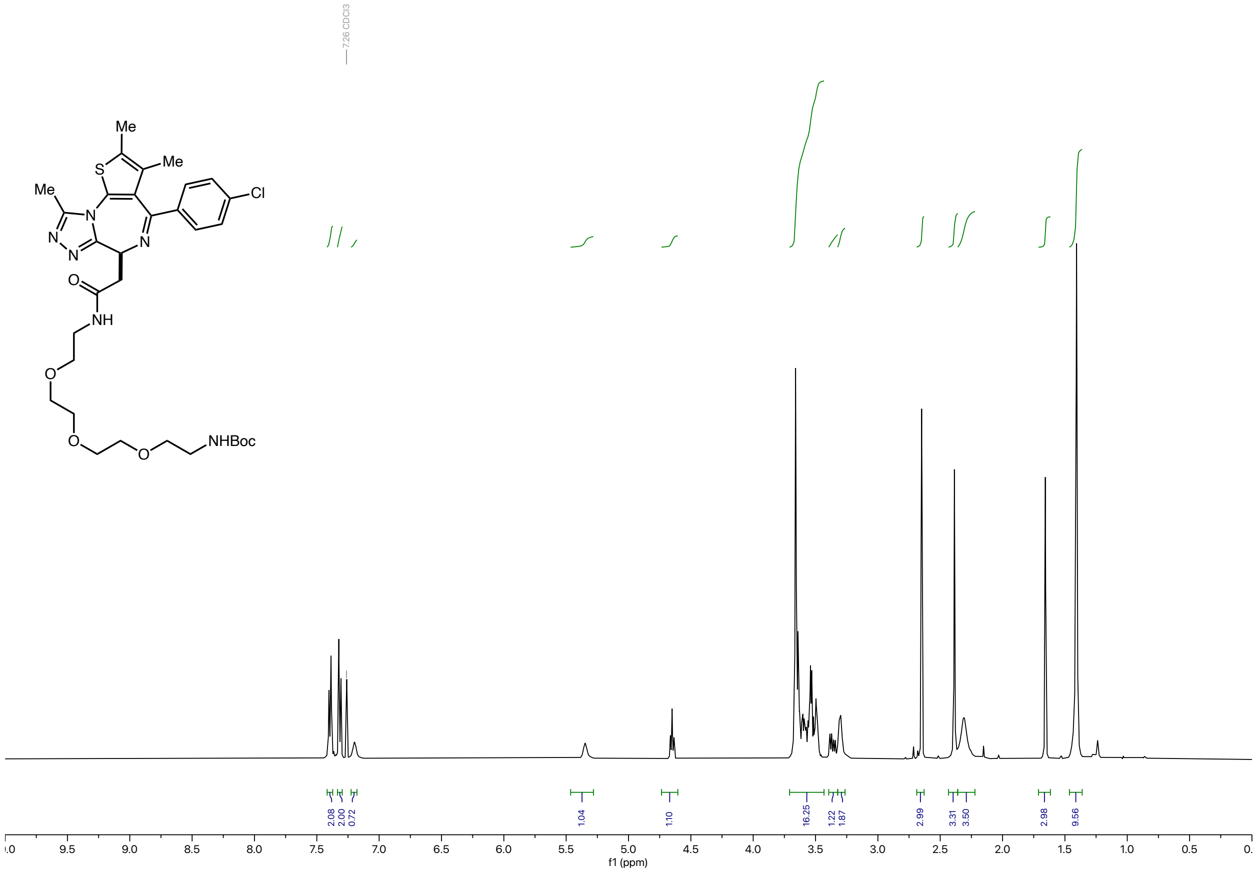

### JQ-1-PEG3-NHBoc: $^{13}C$ NMR

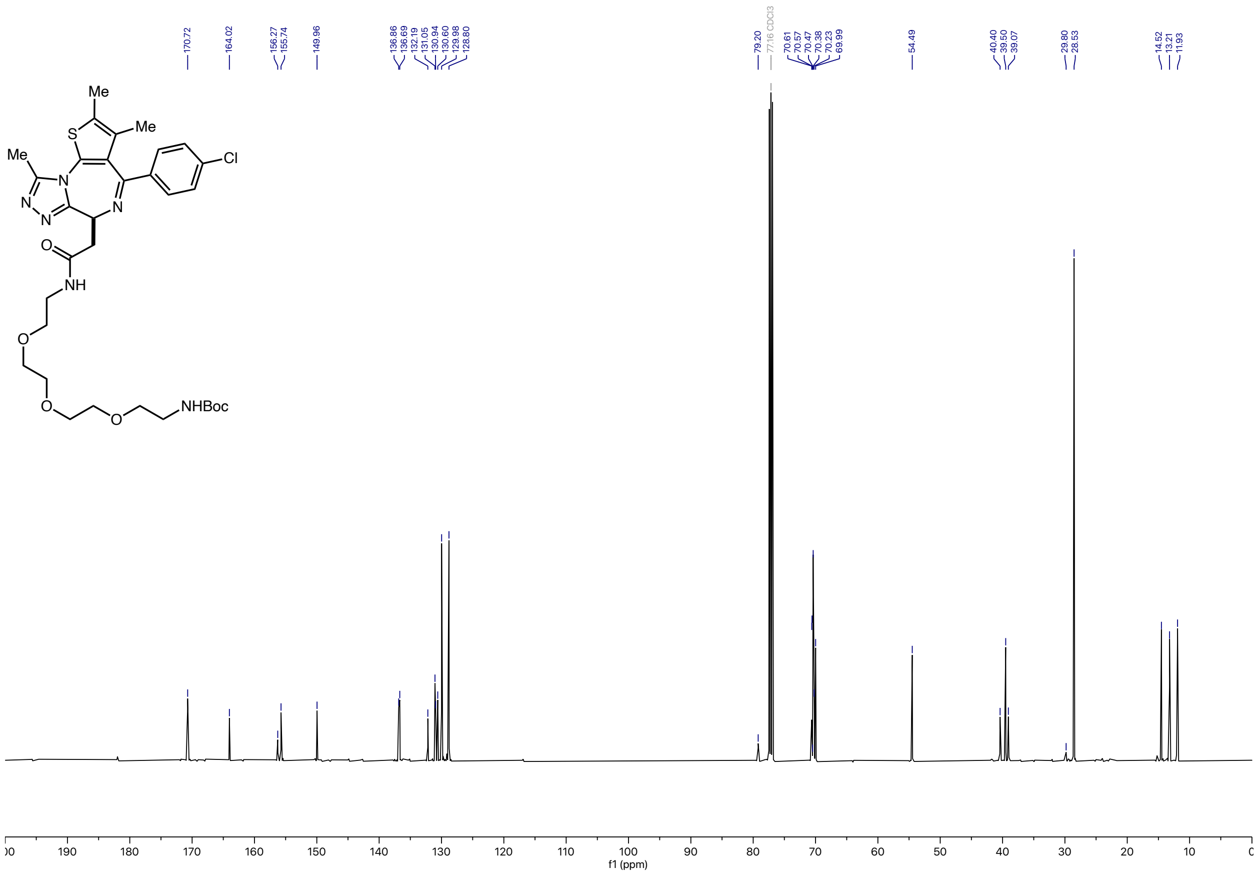

### JQ-1-PEG3-Ir: $^{1}$H NMR

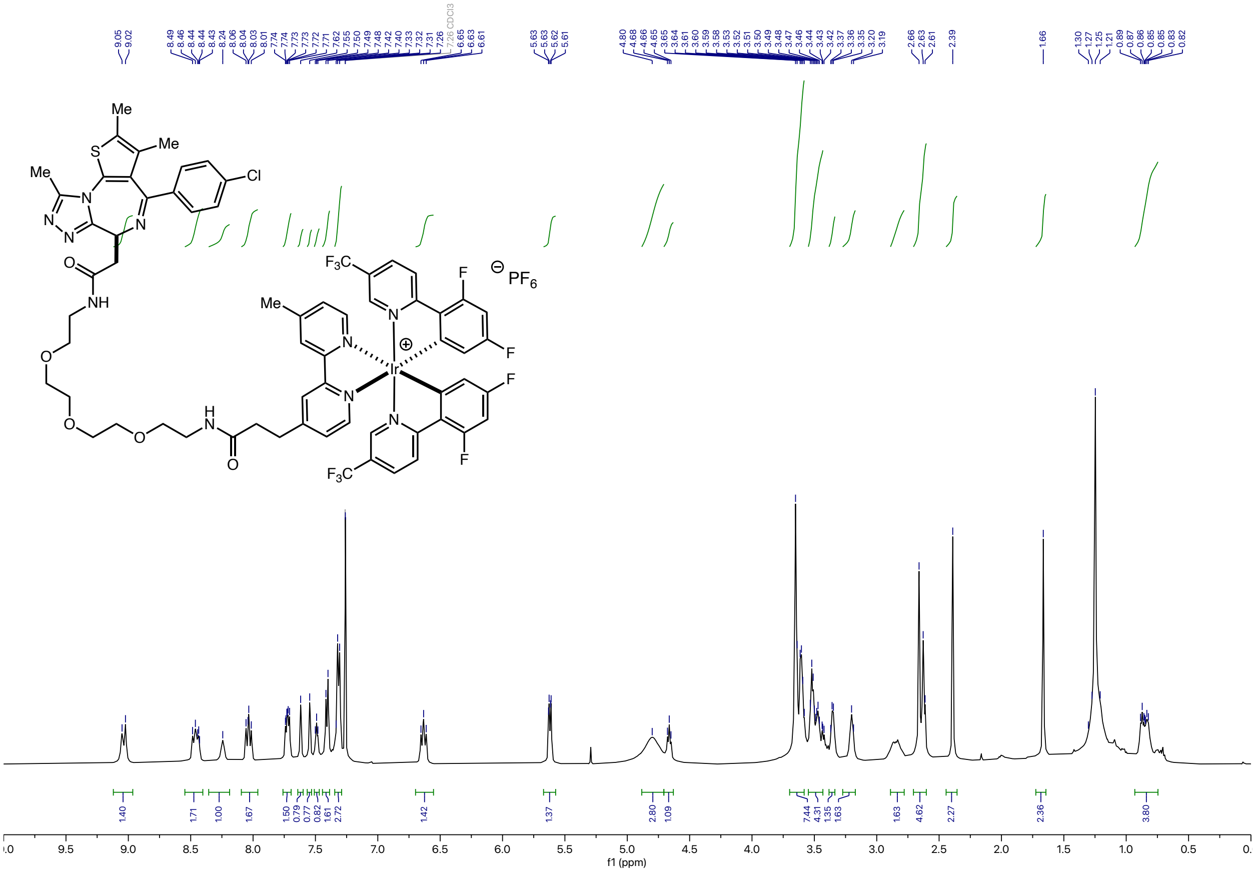

### JQ-1-PEG3-Ir: $^{13}C$ NMR

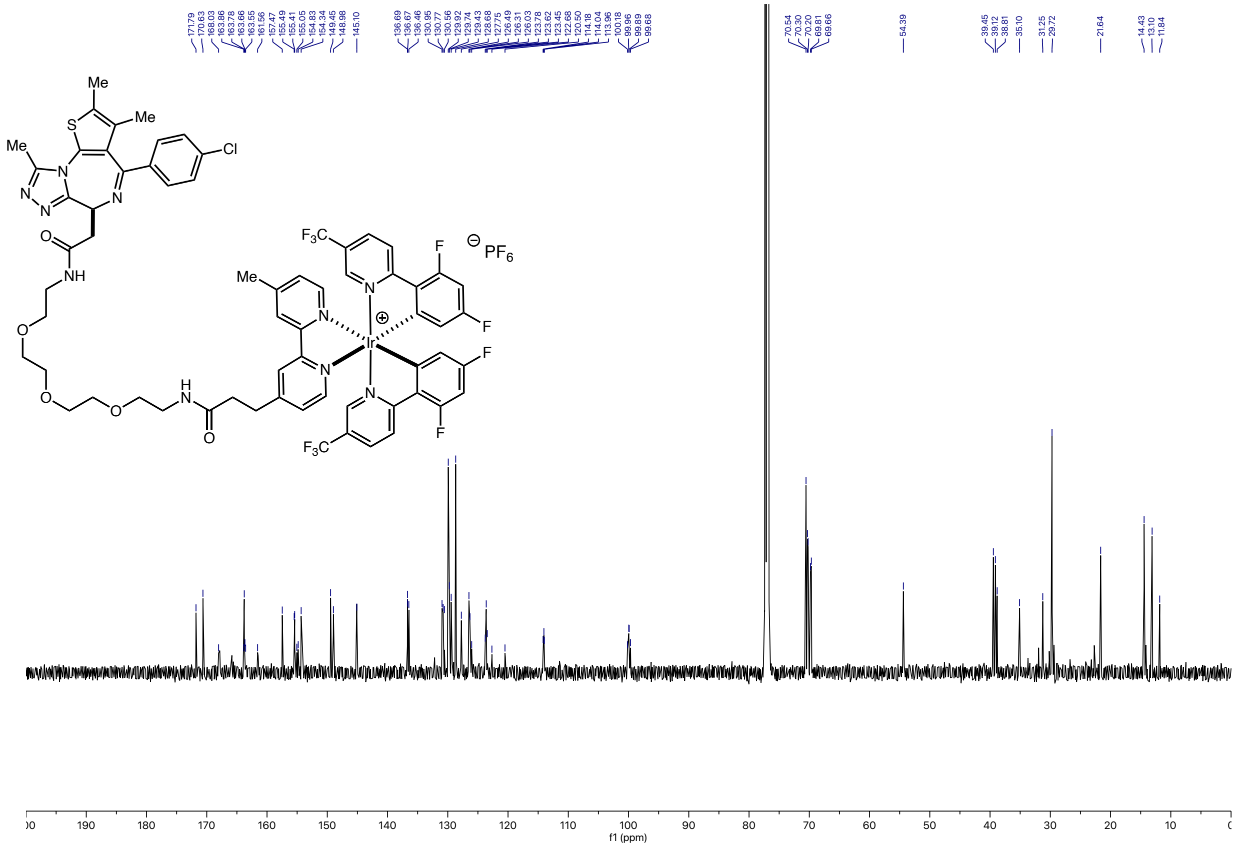

### JQ-1-PEG3-Ir: $^{19}$F NMR

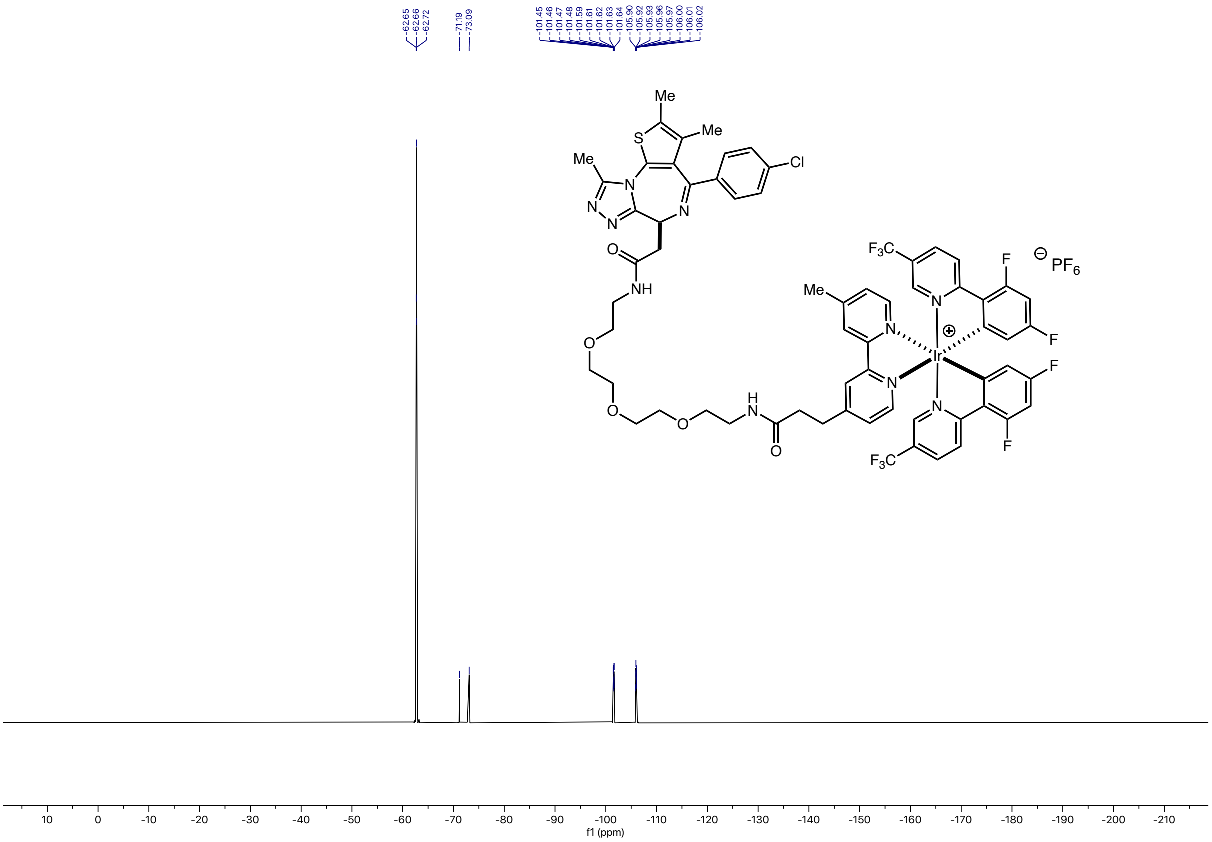

**JQ-1-PEG3-N$_3$: $^1$H NMR**

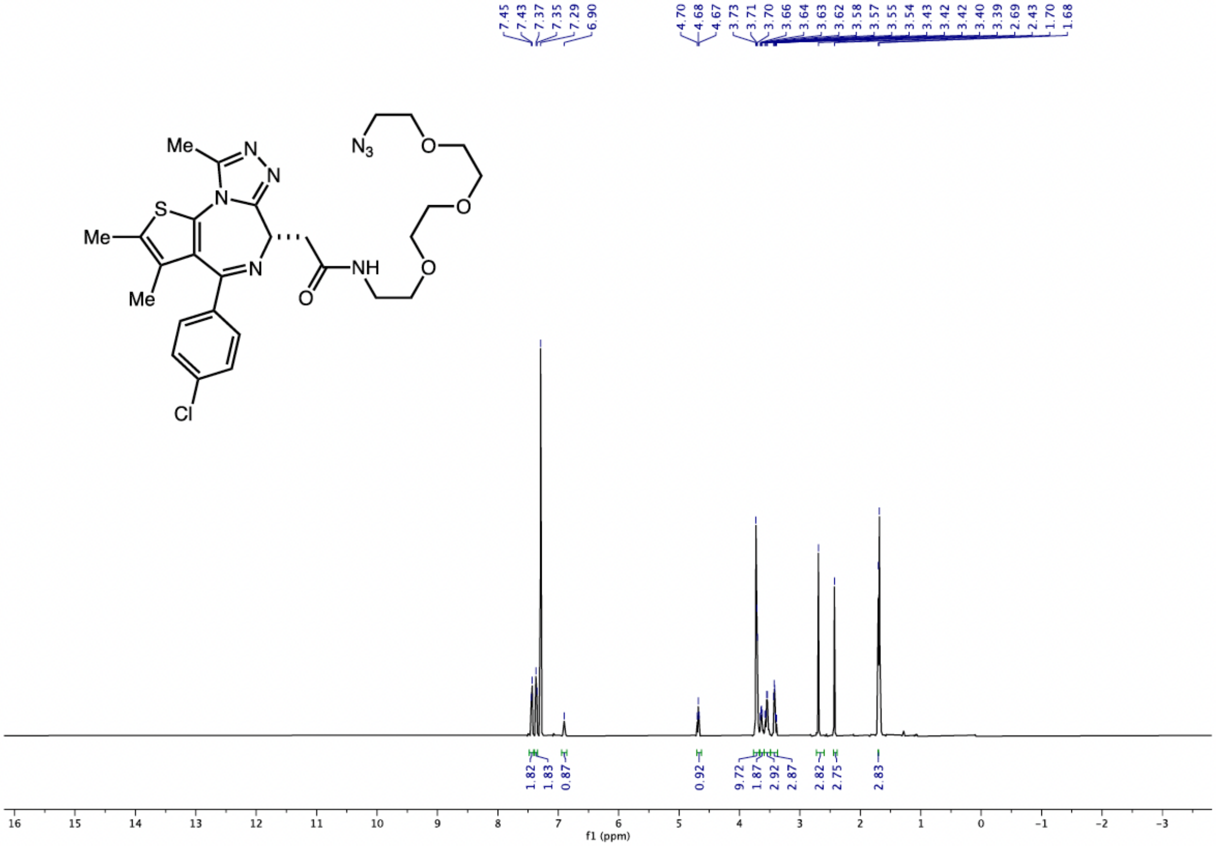

JQ-1-PEG3-$N_3$: $^{13}C$ NMR

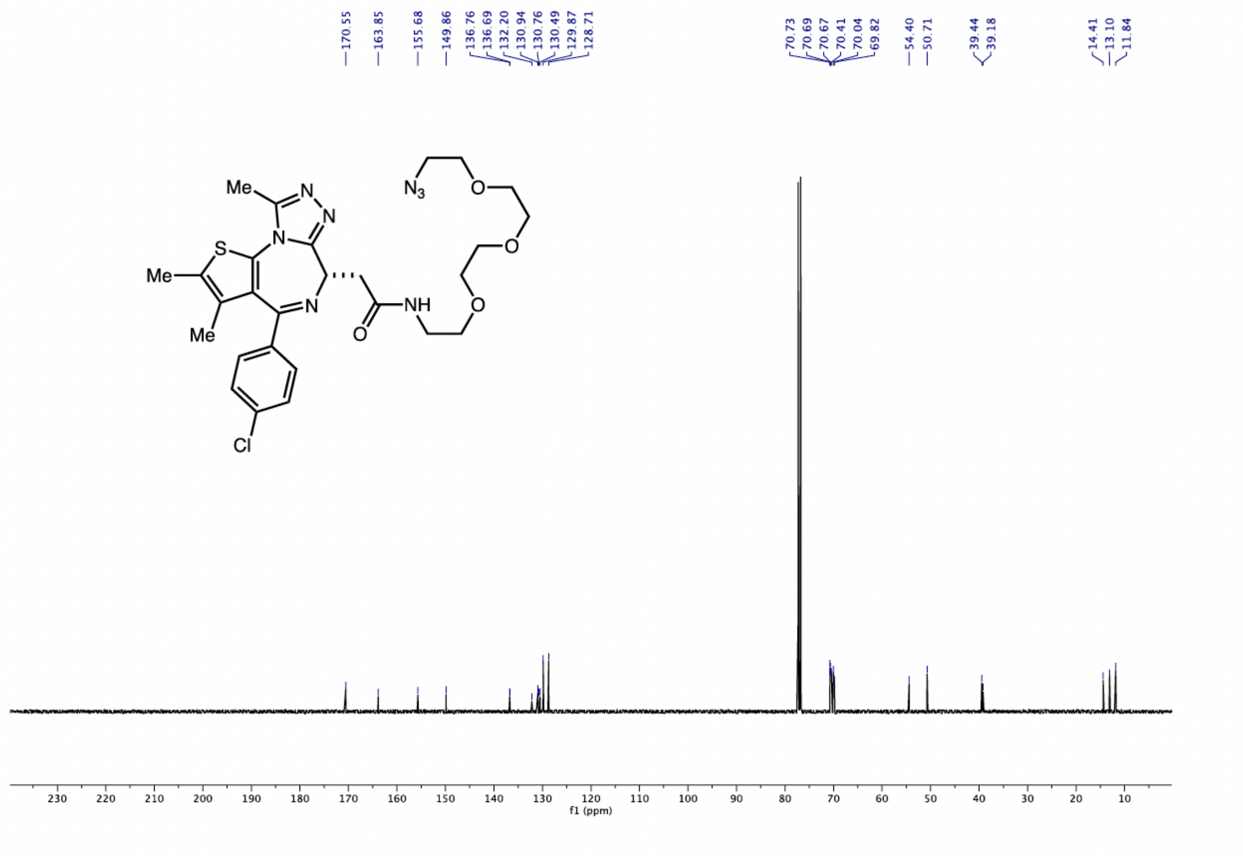

### JQ-1-PEG3-G1-Ir: $^{1}H$ NMR

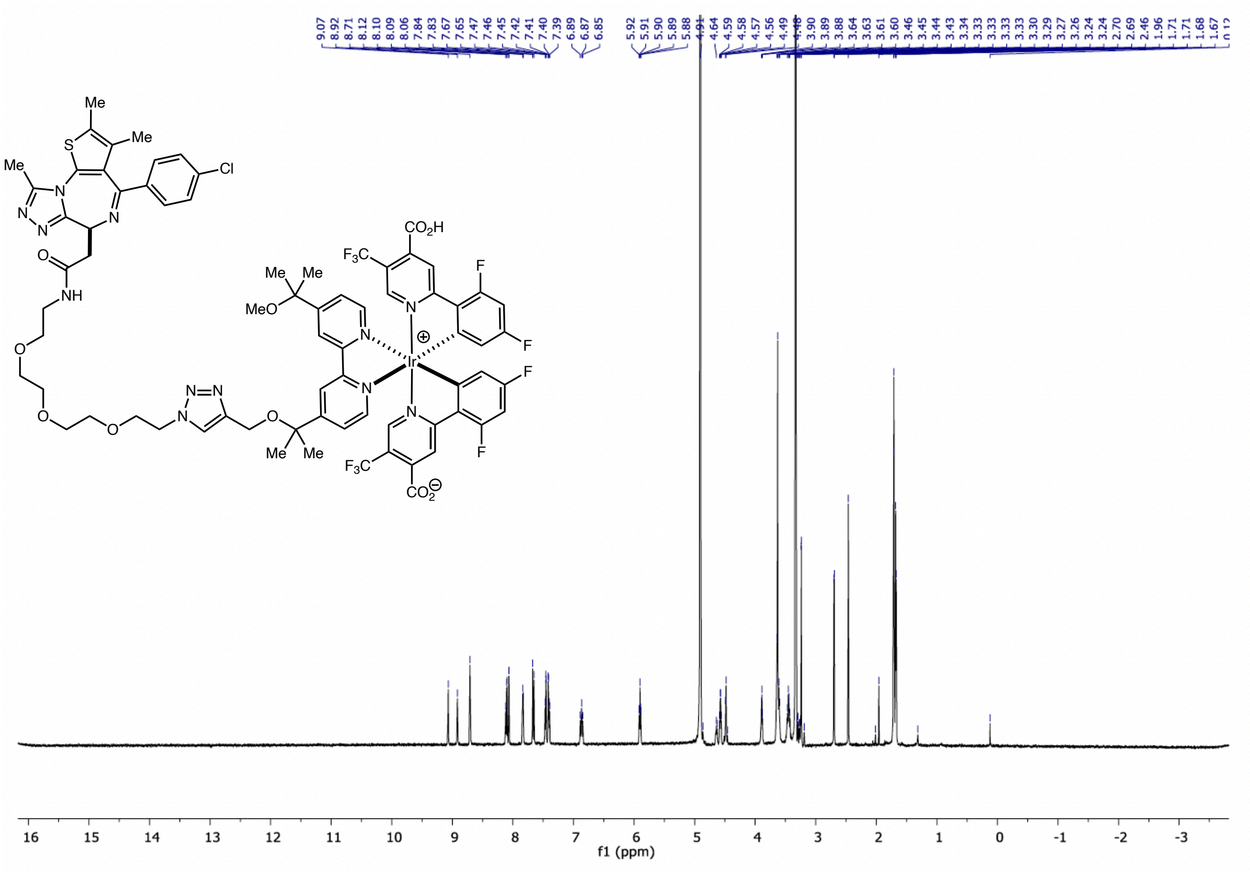

**JQ-1-PEG3-G1-Ir: $^{13}$C NMR**

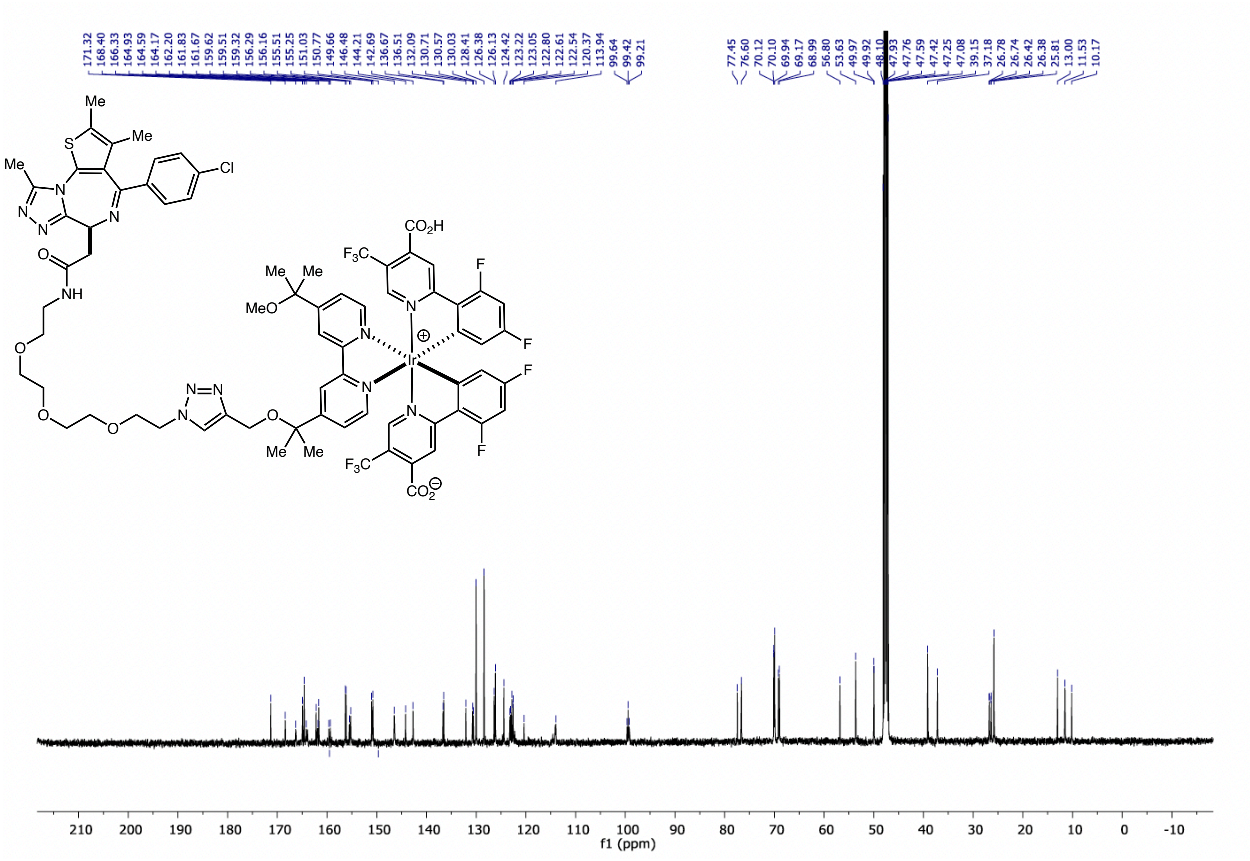

#### JQ-1-PEG3-G1-Ir: $^{19}F$ NMR

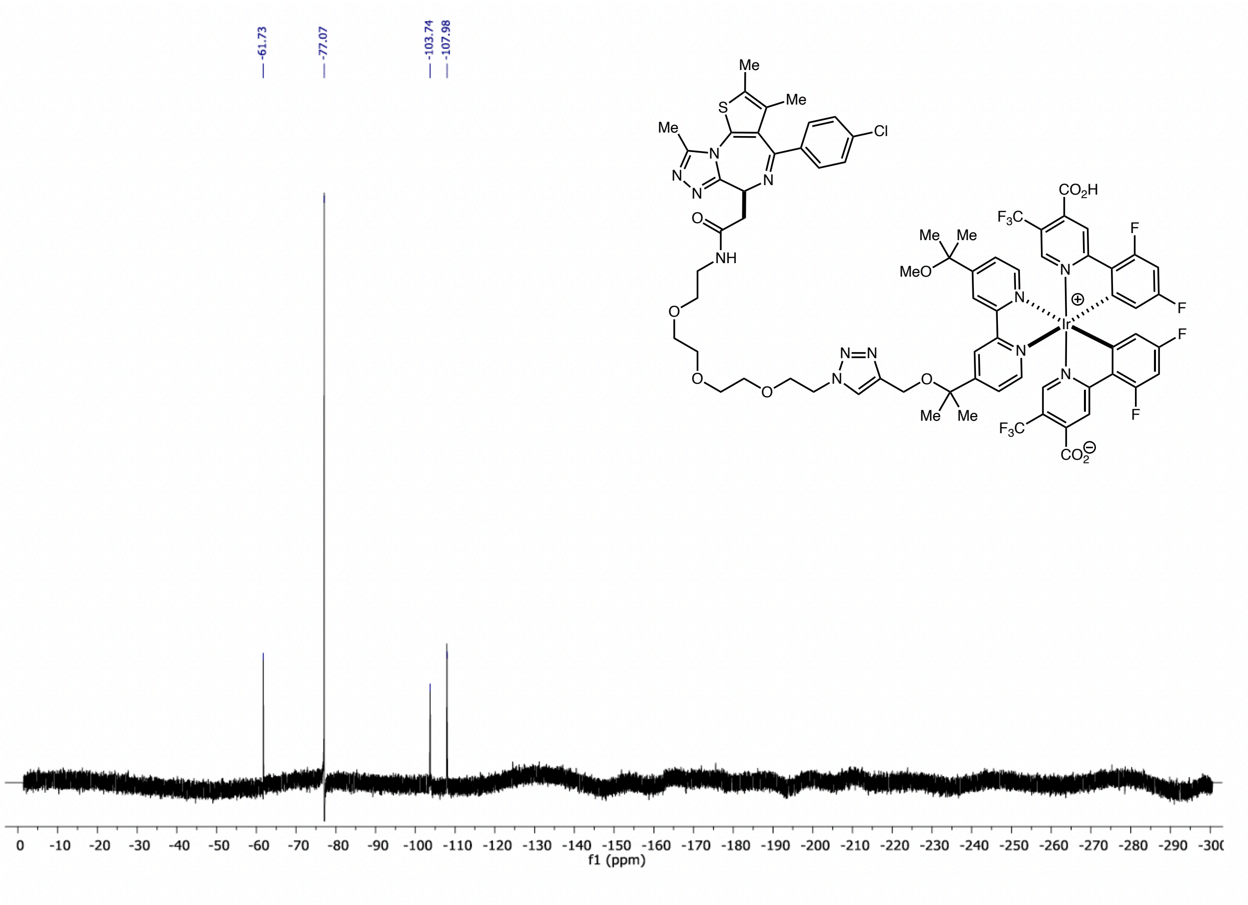

### JQ-1-Dz-Alkyne: $^1$H NMR

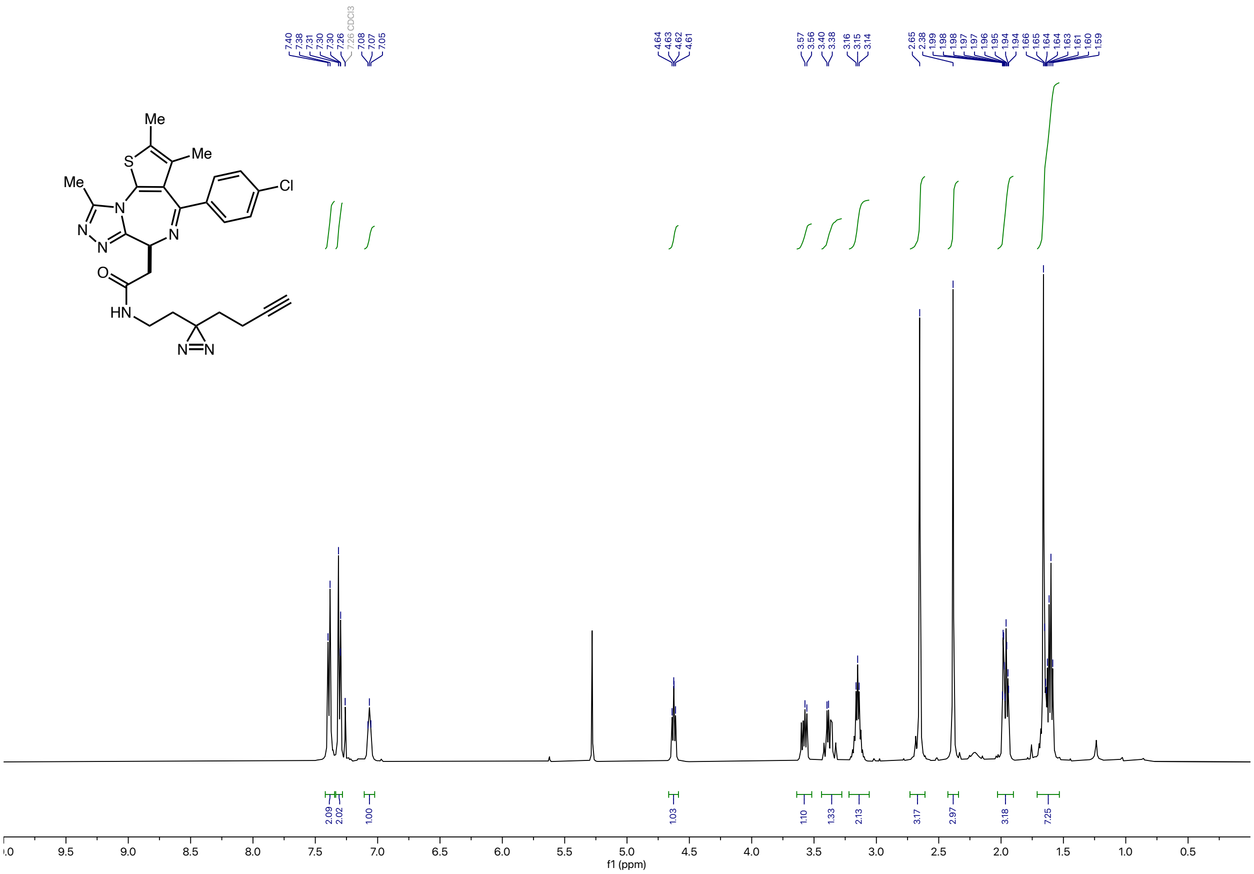

### JQ-1-Dz-Alkyne: $^{13}C$ NMR

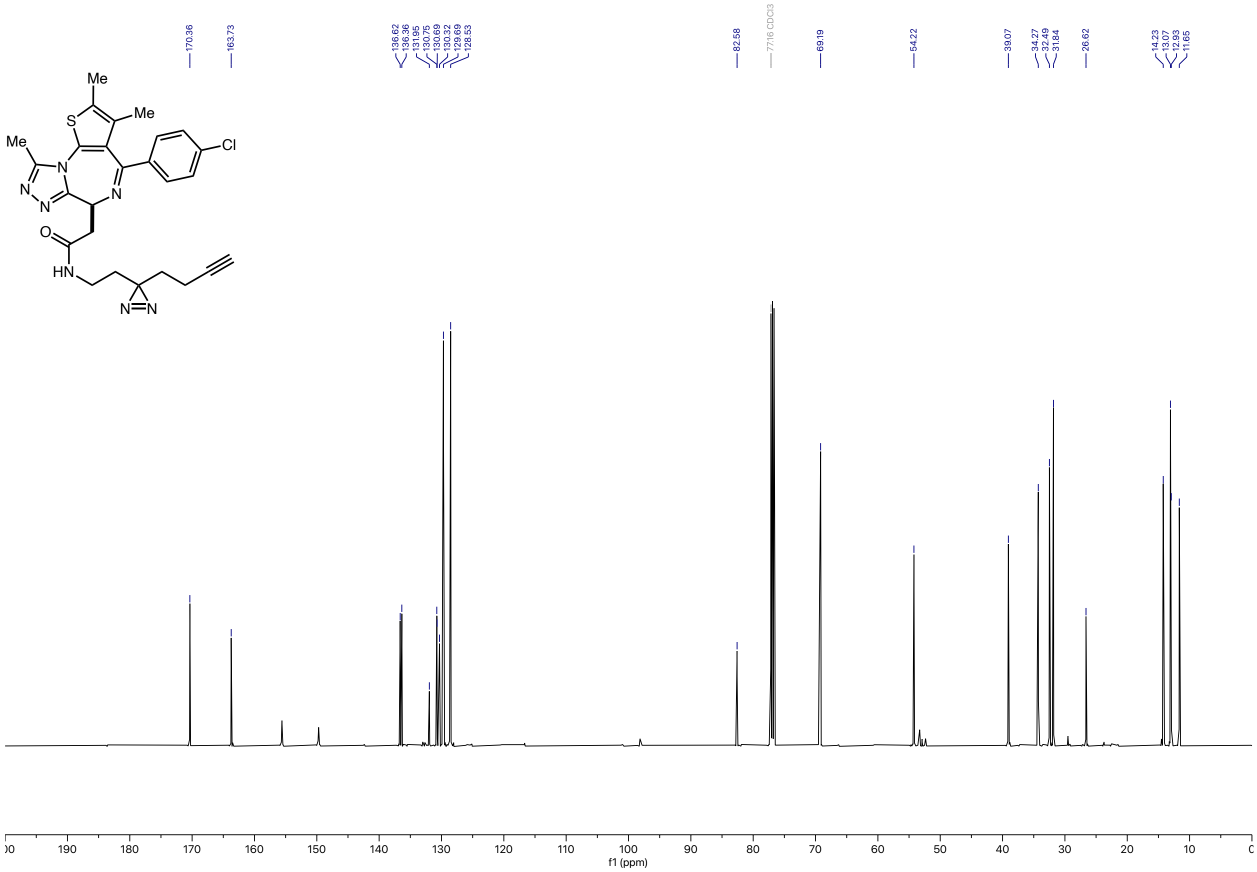

### Deshep-dasatinib-PEG5-NH$_2$: $^1$H NMR

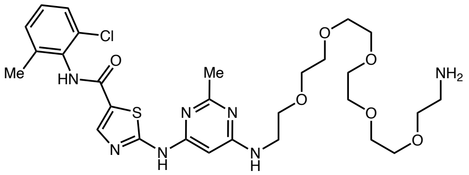

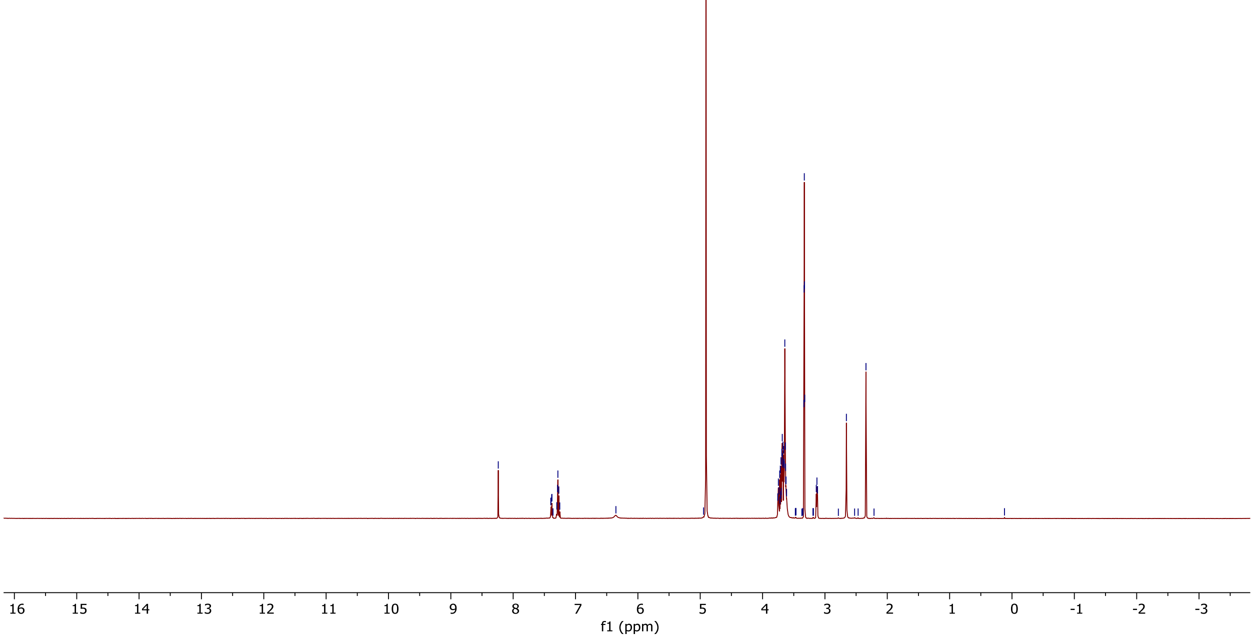

### Deshep-dasatinib-PEG5-NH$_2$: $^{13}$C NMR

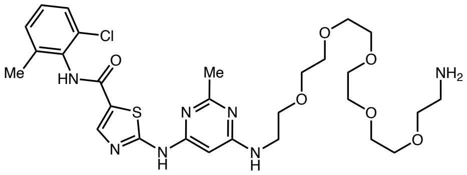

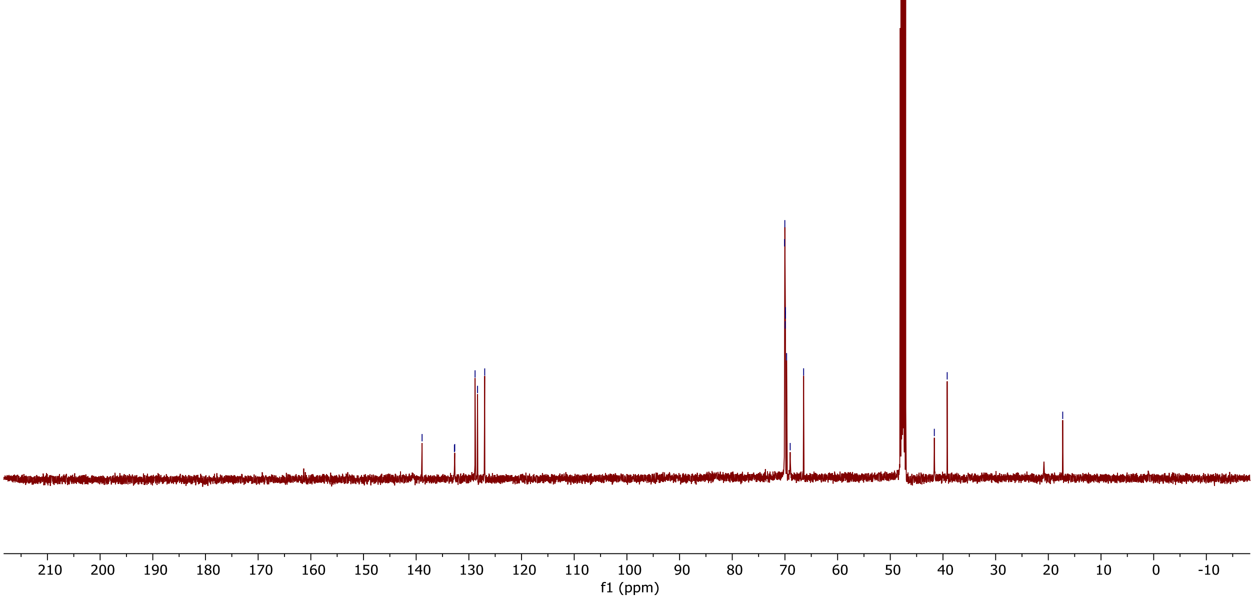

### Deshep-dasatinib-PEG5-Ir: $^{1}H$ NMR

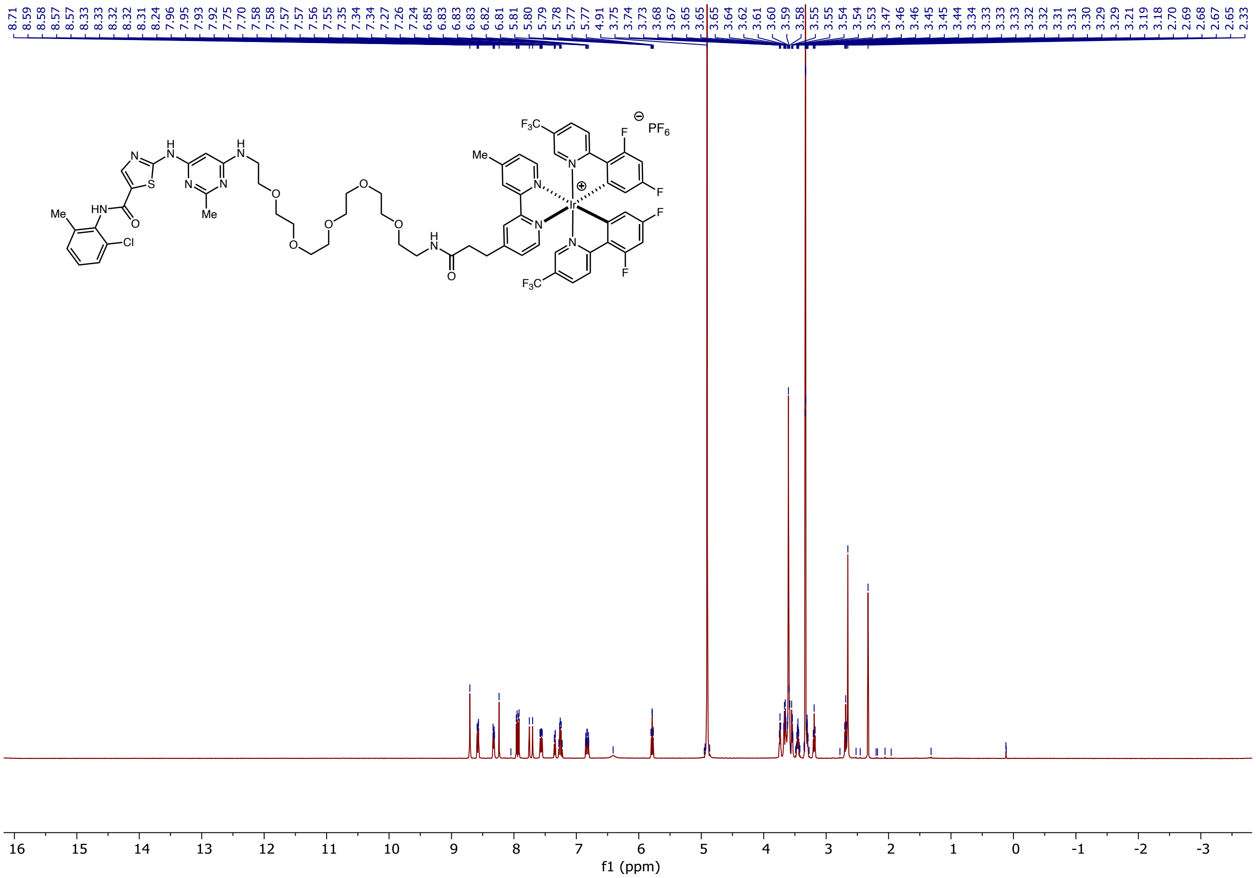

#### Deshep-dasatinib-PEG5-Ir: $^{13}C$ NMR

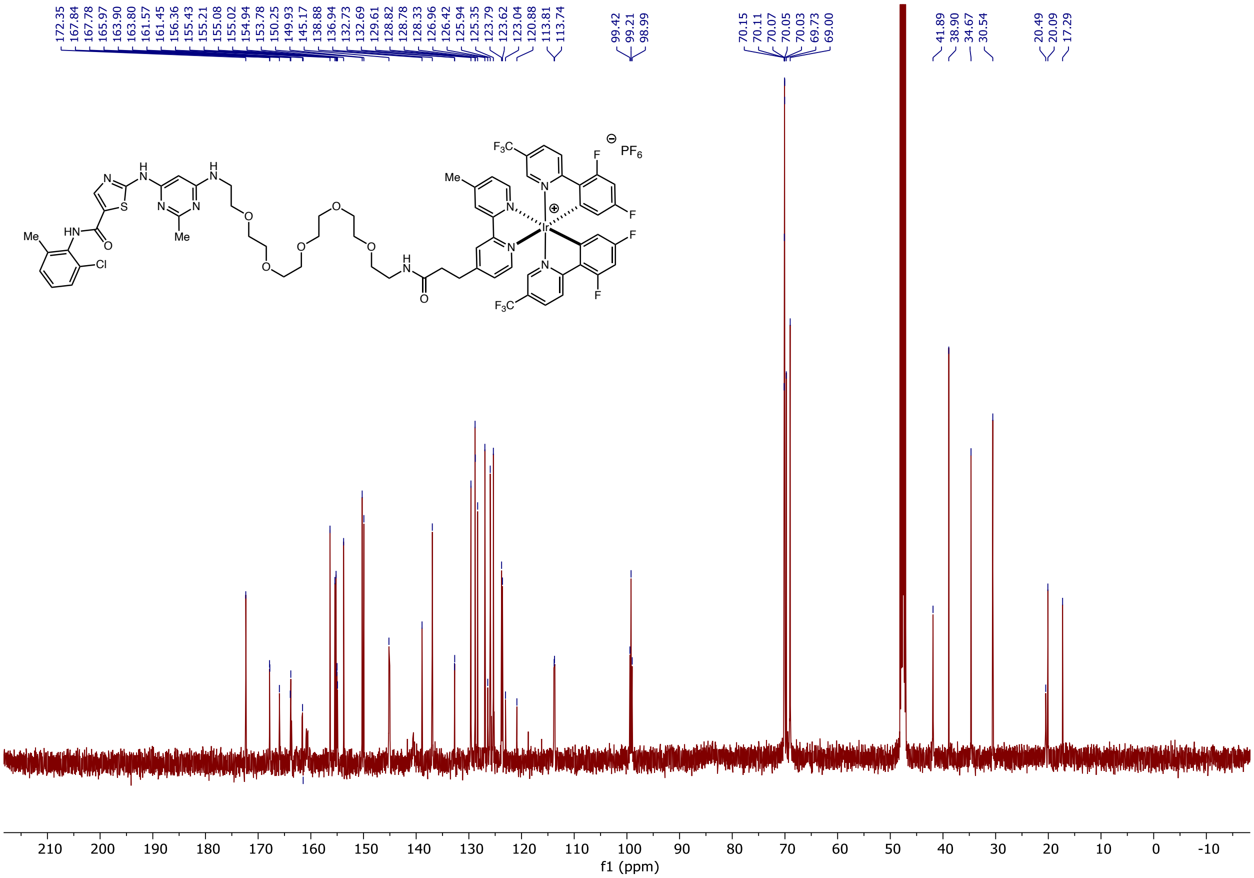

### Deshep-dasatinib-PEG5-Ir: $^{19}$F NMR

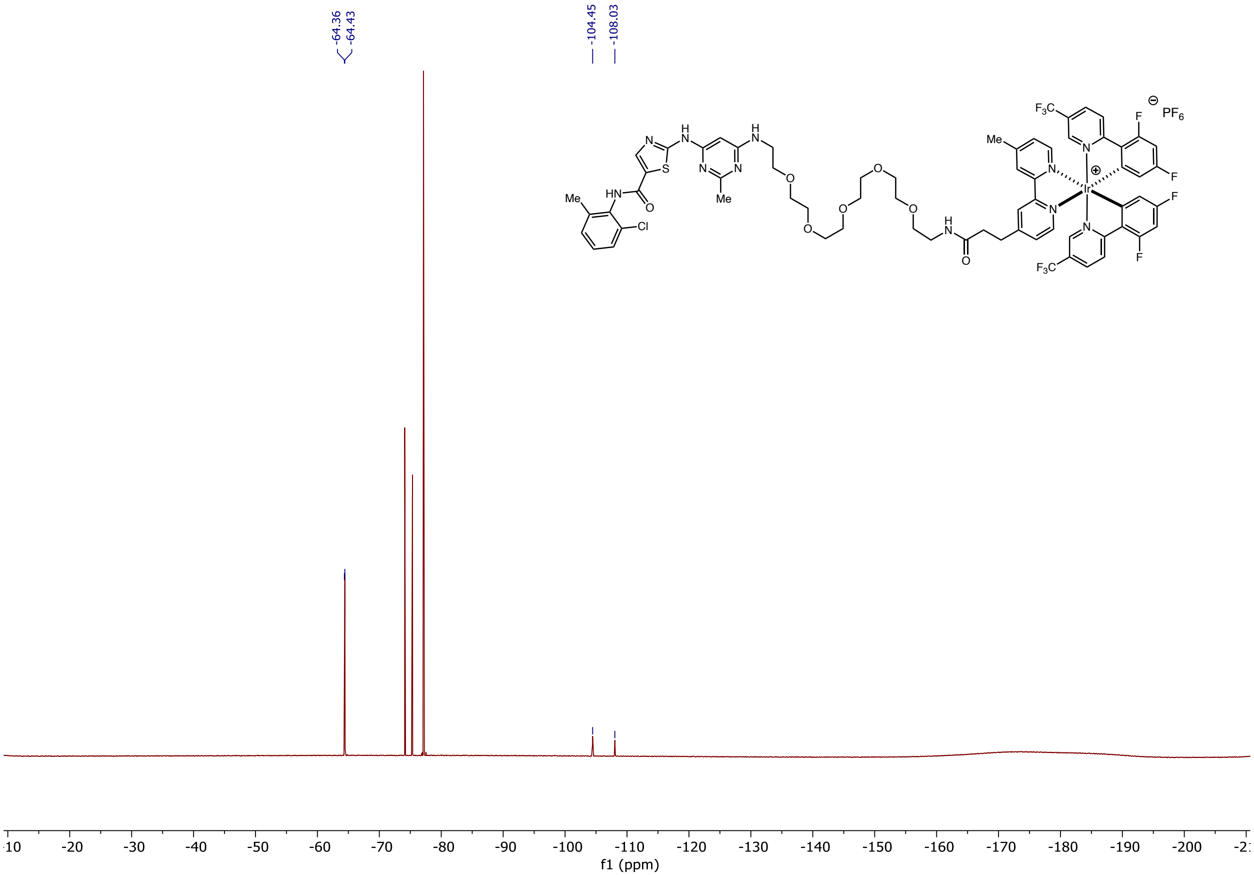

### Dasatinib-PEG3-Ir: $^{1}H$ NMR

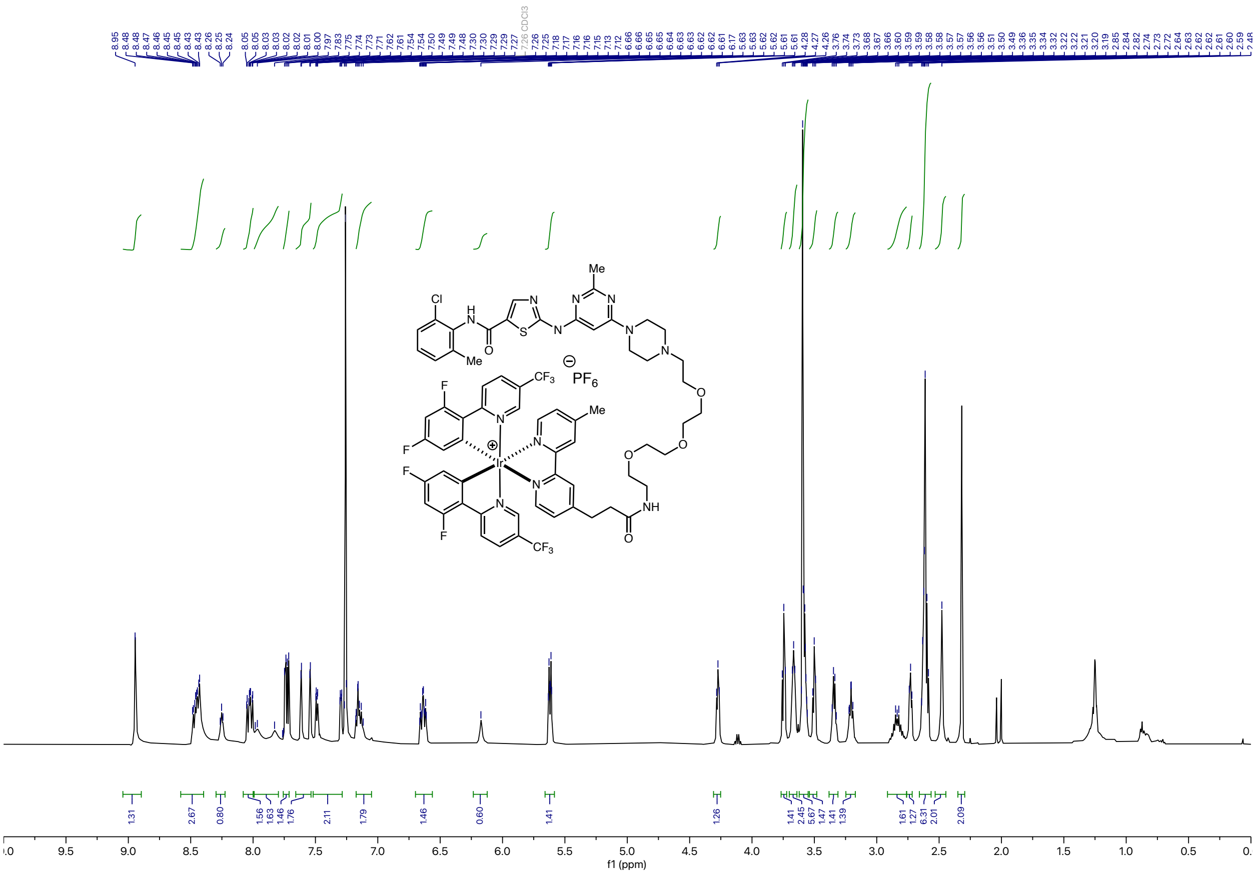

### Dasatinib-PEG3-Ir: $^{13}$C NMR

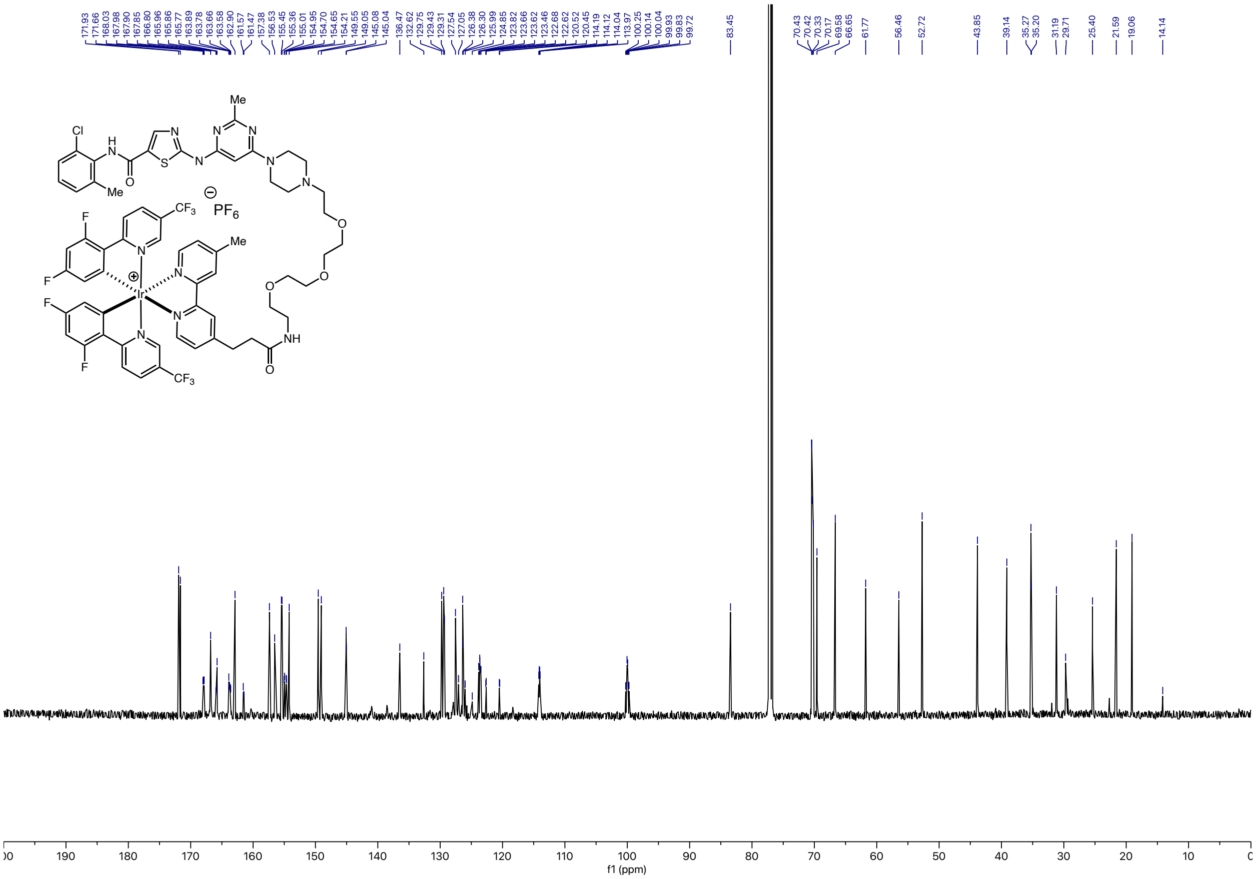

### Dasatinib-PEG3-Ir: $^{19}$F NMR

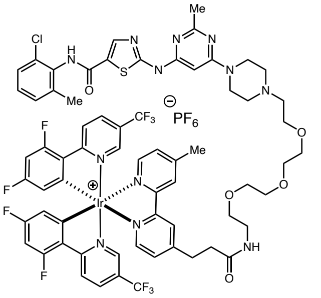

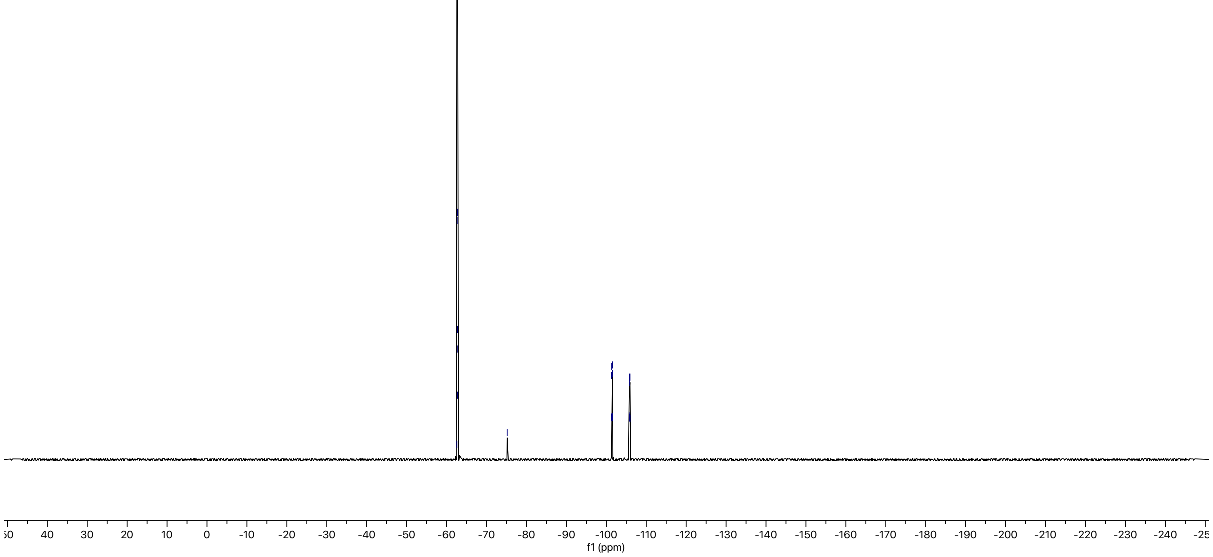

### Dasatinib-Dz-Alkyne: $^1$H NMR

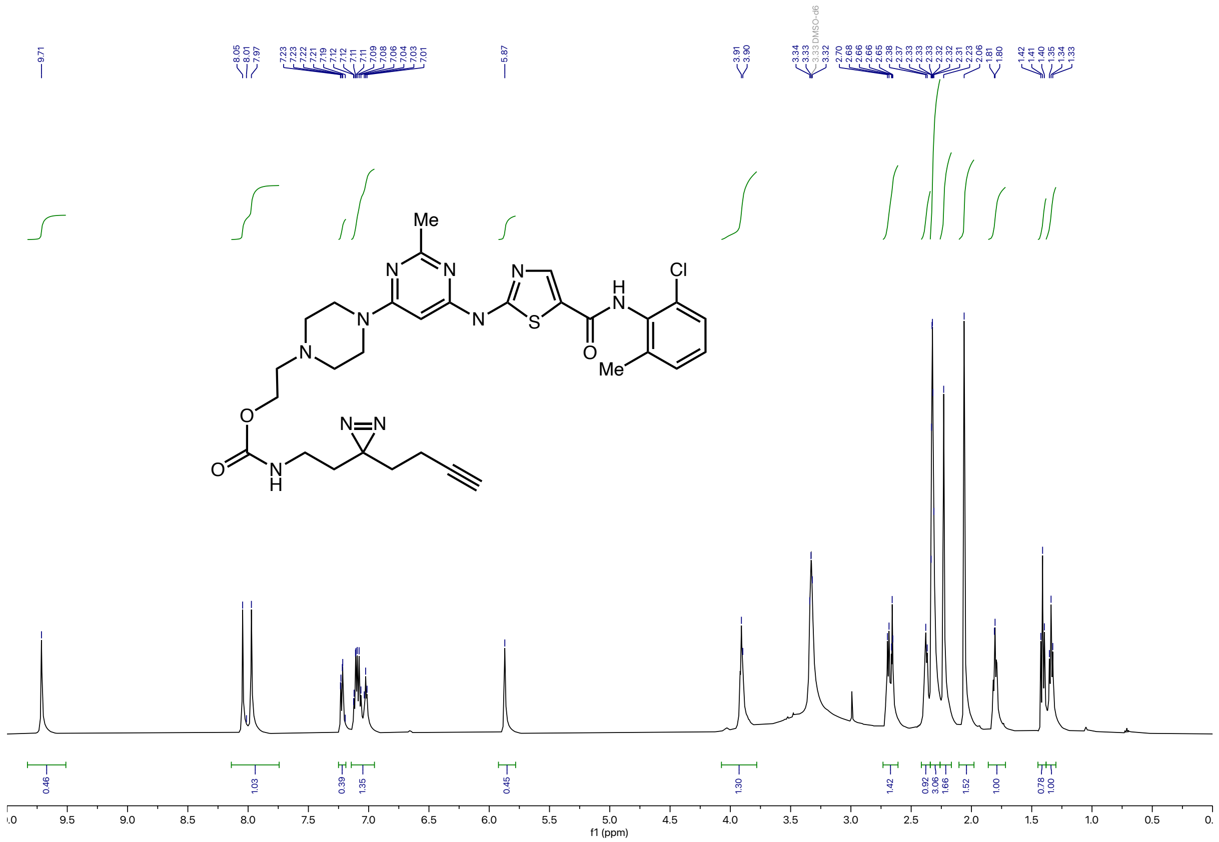

#### Dasatinib-Dz-Alkyne: $^{13}C$ NMR

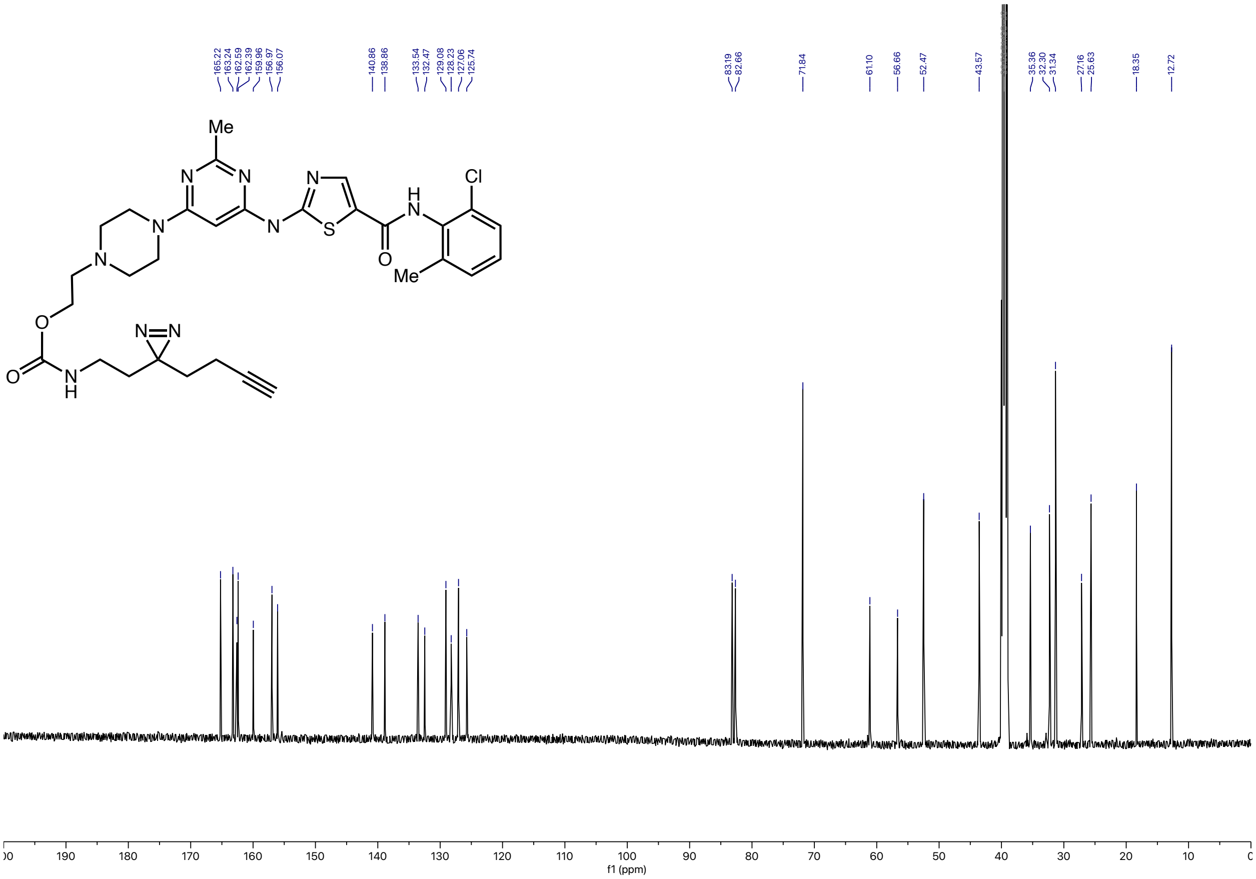

### Paclitaxel-G2-Ir: [1]H NMR

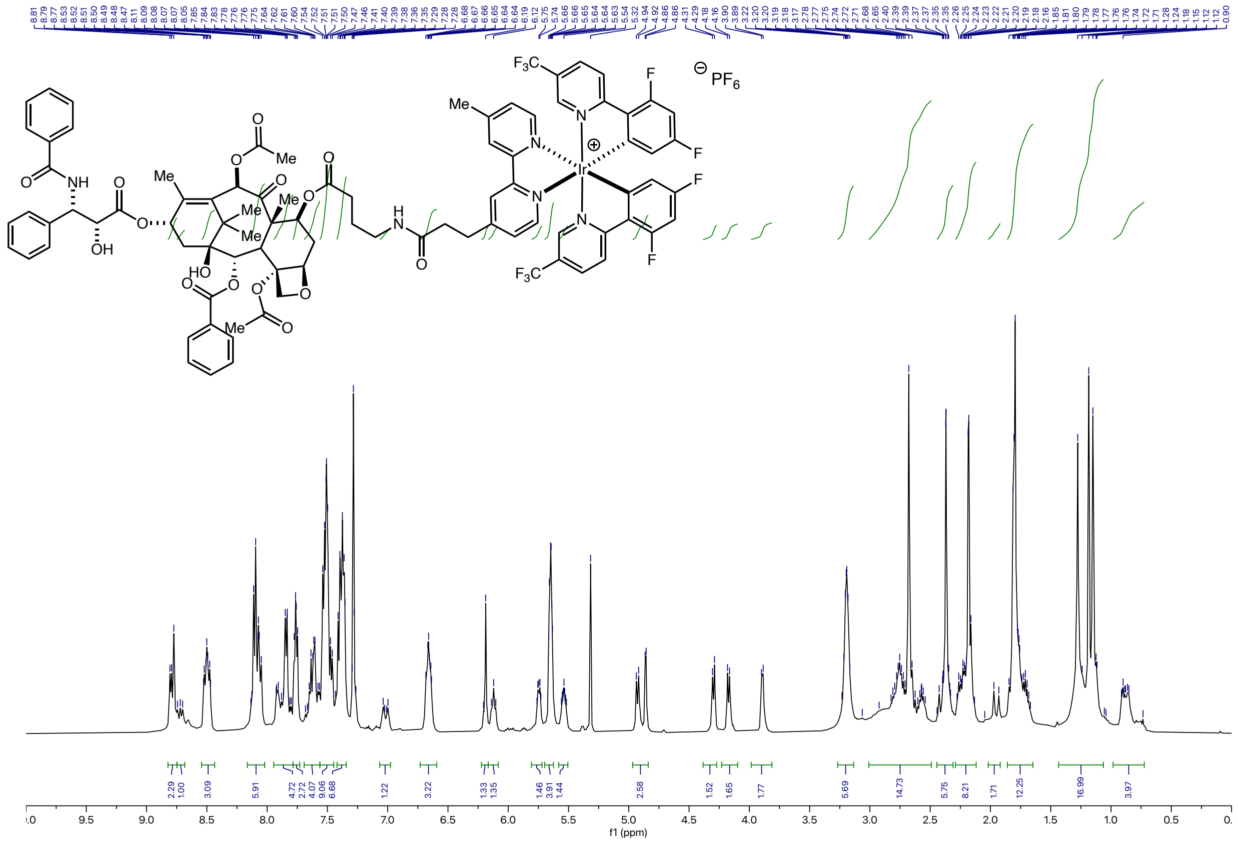

### Paclitaxel-G2-Ir: $^{13}C$ NMR

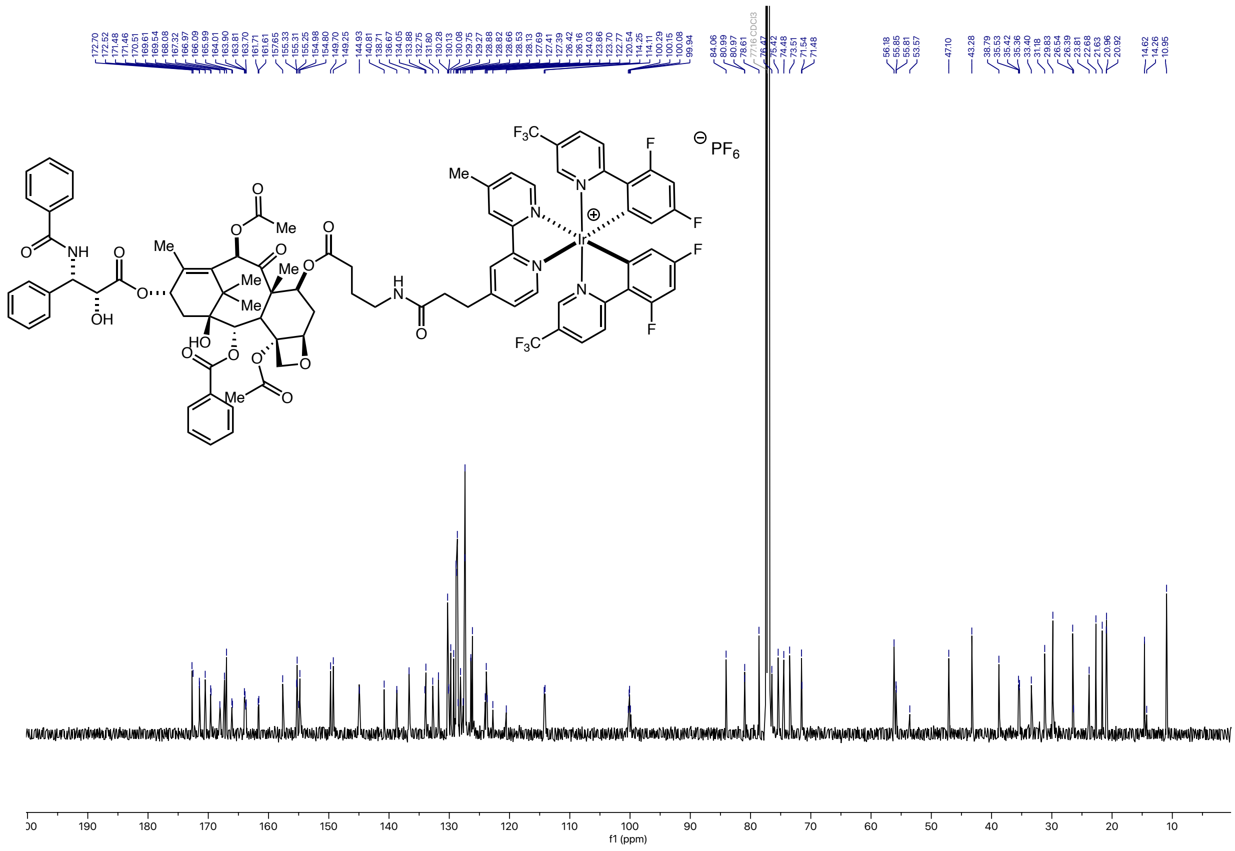

### Paclitaxel-G2-Ir: $^{19}$F NMR

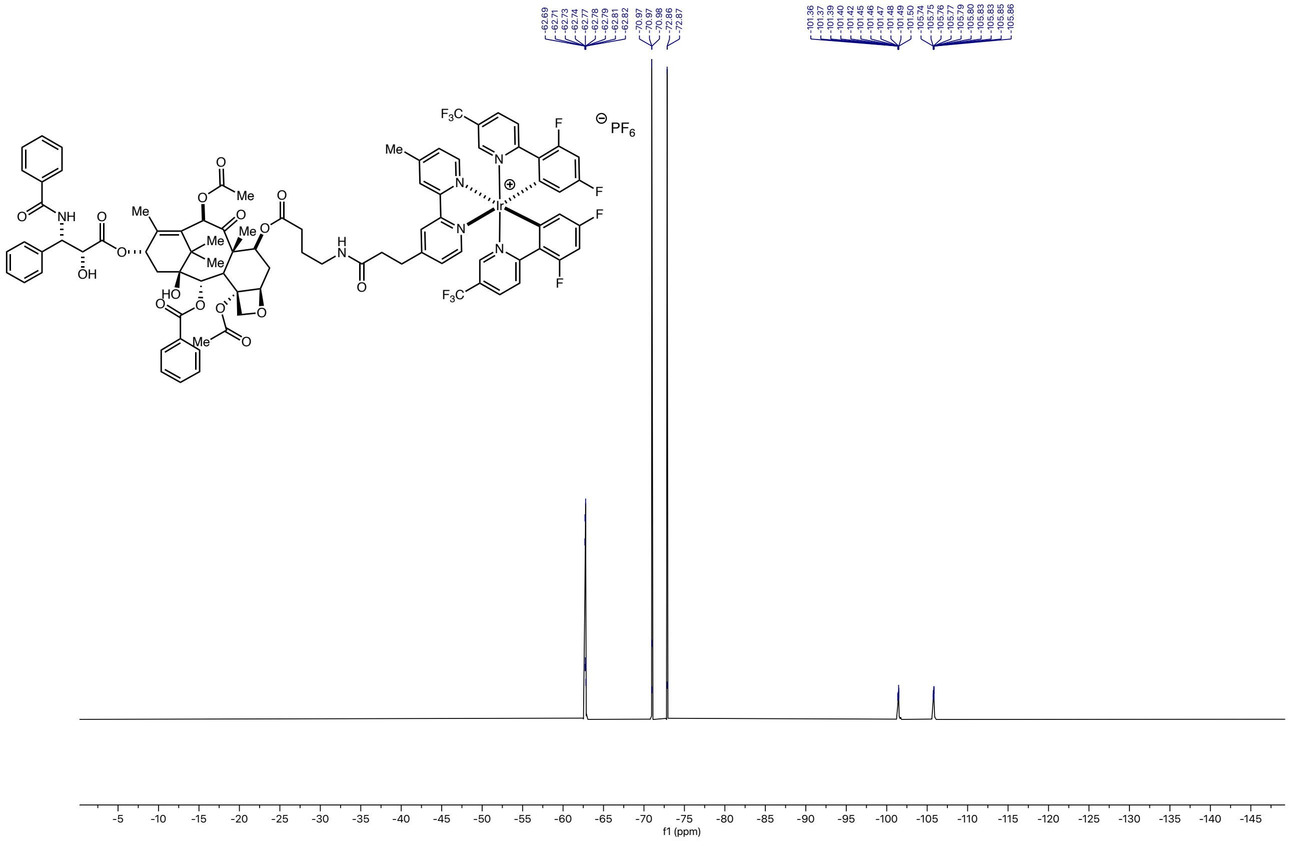

**(5aR,6S,6aS)-3-((4'-(2-(2-(2-azidoethoxy)ethoxy)ethoxy)-4,6-difluoro-2',6'-dimethyl-[1,1'-biphenyl]-3-yl)methoxy)-5,5a,6,6a-tetrahydrocyclopropa[4,5]cyclopenta[1,2-c]pyridine-6-carboxylic acid: MK8666-$N_3$**

**MK8666-$N_3$: $^1$H NMR**

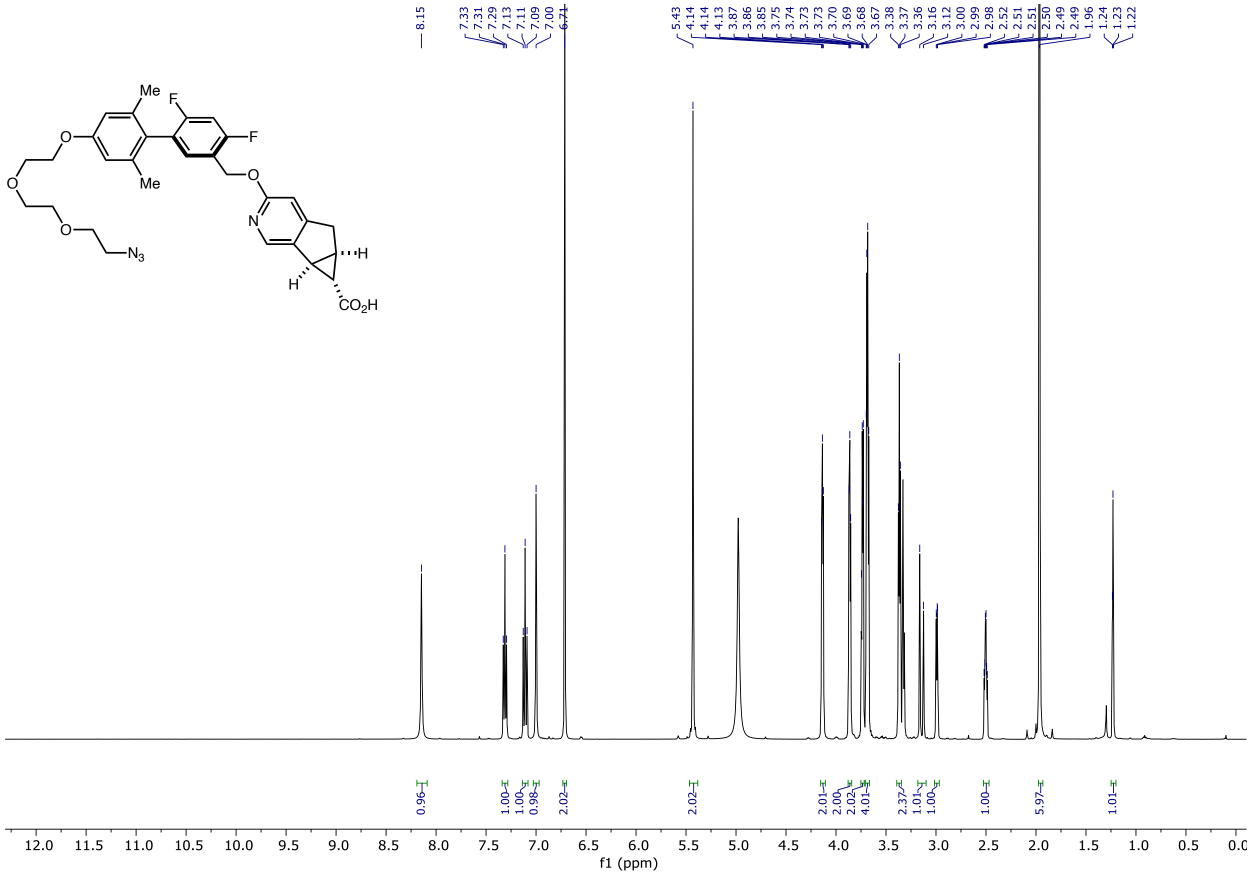

### MK8666-$N_3$: $^{13}C$ NMR

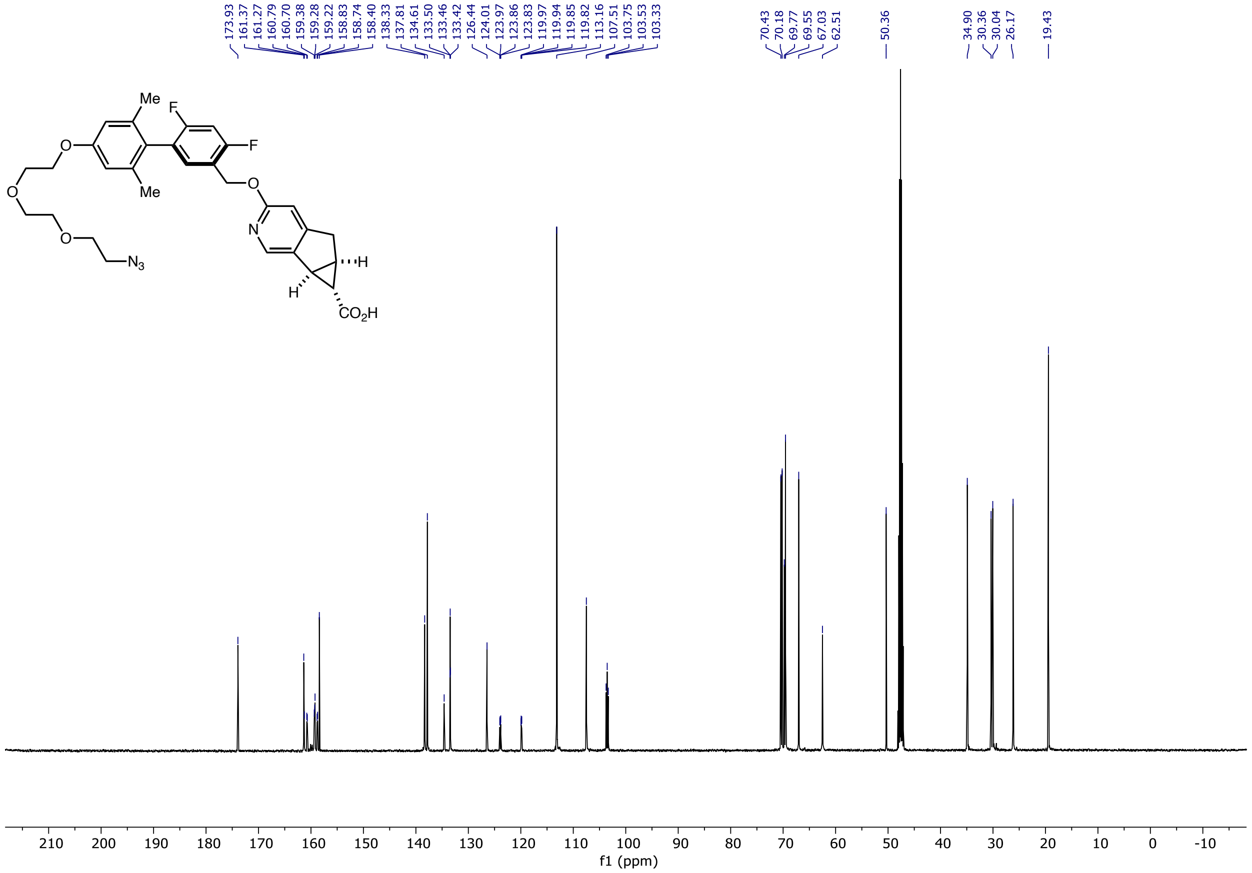

### MK8666-$N_3$: $^{19}$F-NMR

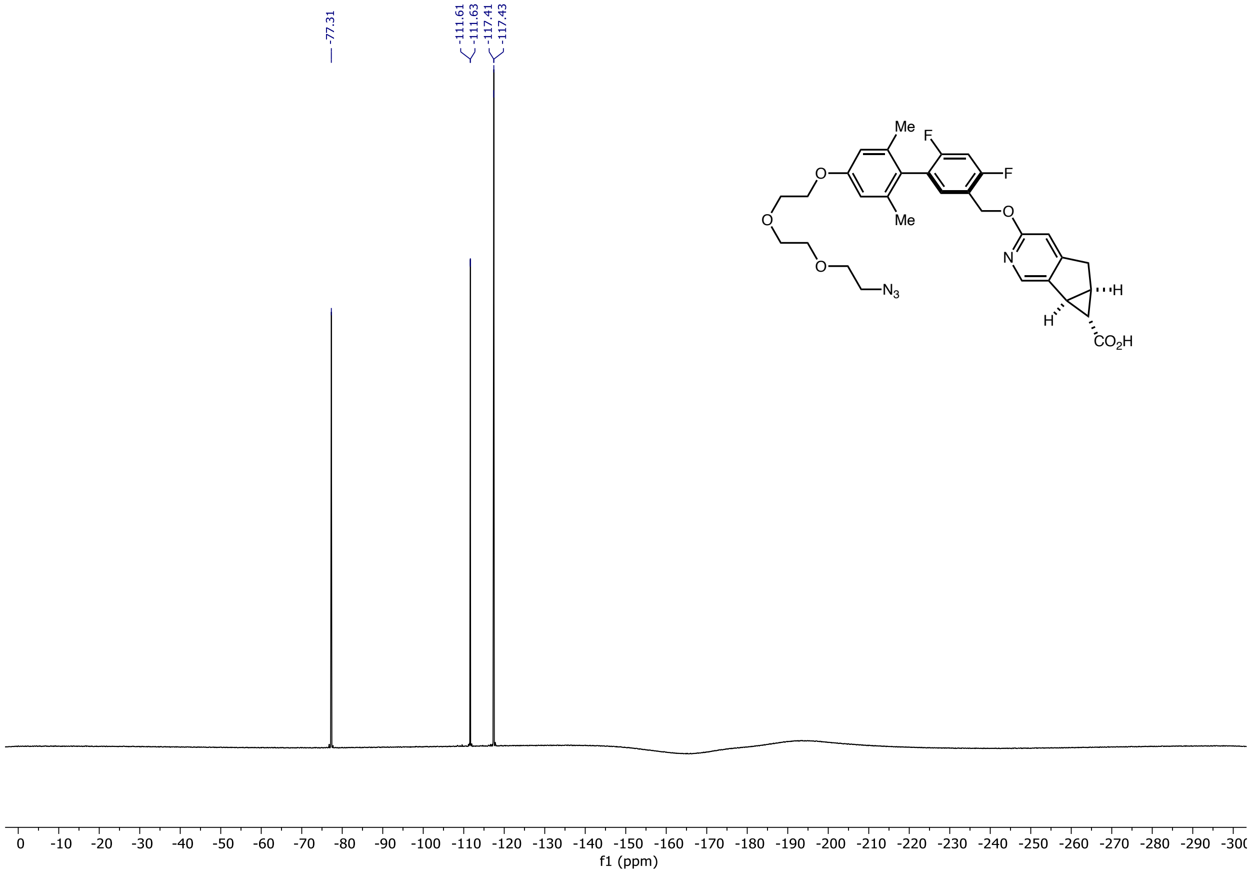

**(5aR,6S,6aS)-3-((4'-(2-(3-(but-3-yn-1-yl)-3H-diazirin-3-yl)ethoxy)-4,6-difluoro-2',6'-dimethyl-[1,1'-biphenyl]-3-yl)methoxy)-5,5a,6,6a-tetrahydrocyclopropa[4,5]cyclopenta[1,2-c]pyridine-6-carboxylic acid: MK8666-Dz**

**MK8666-Dz: $^{1}$H NMR**

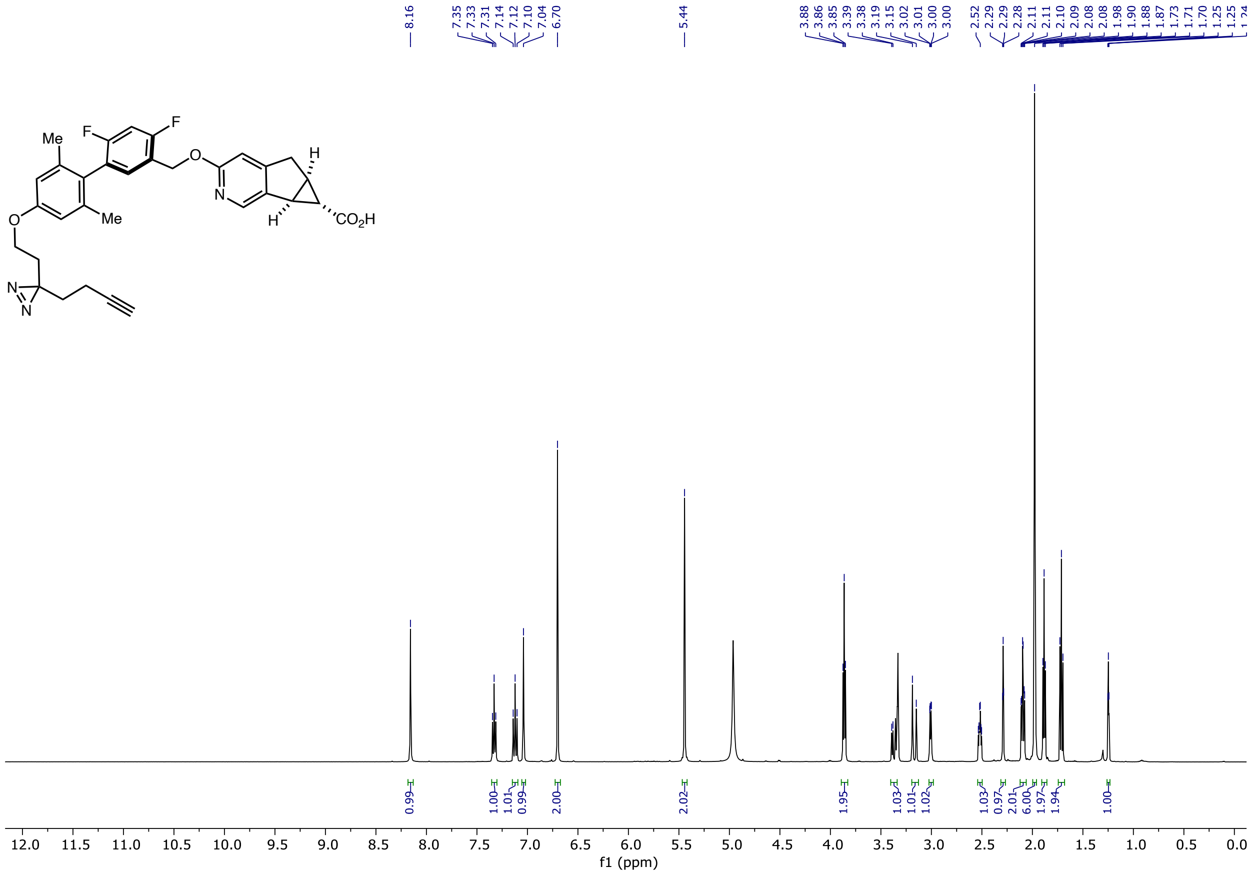

### MK8666-Dz: $^{13}$C NMR

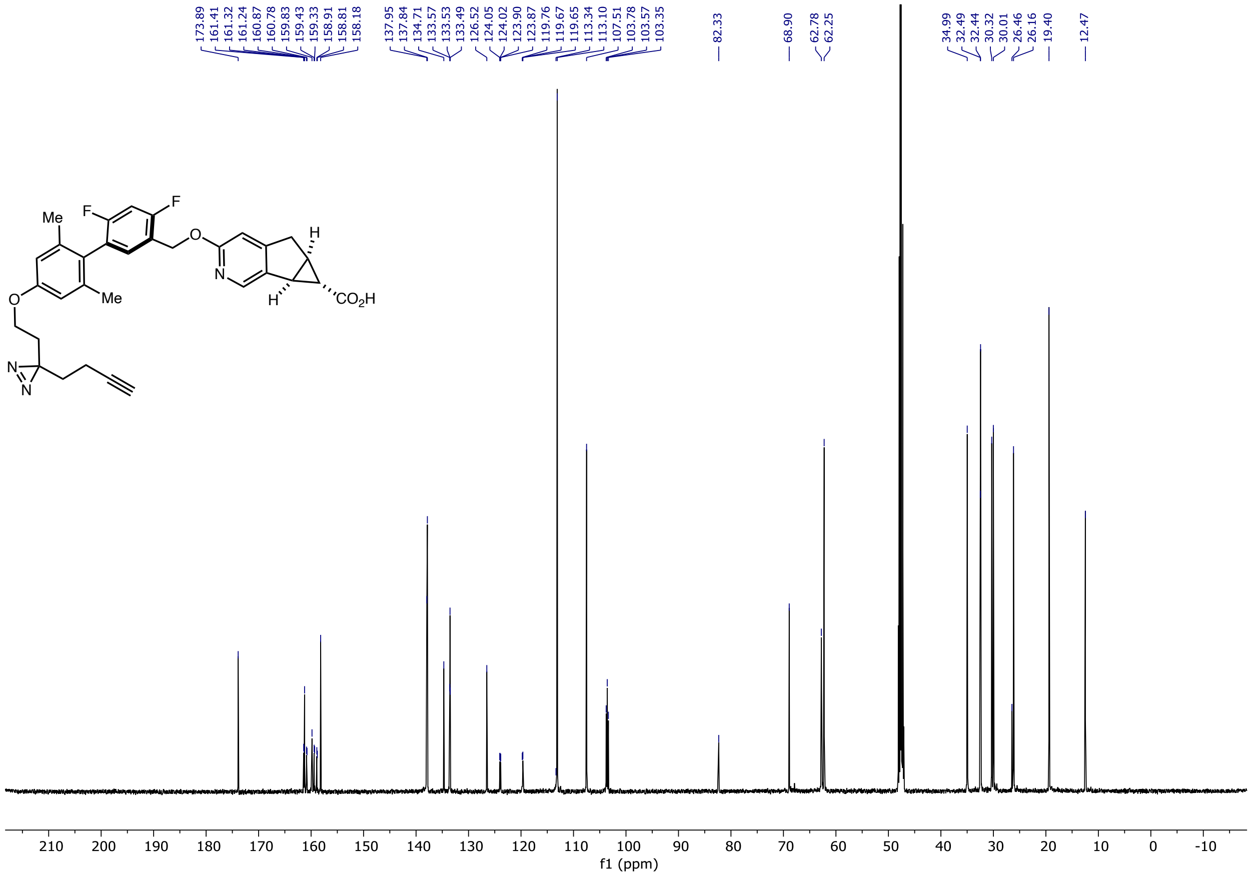

### MK8666-Dz: $^{19}$F-NMR

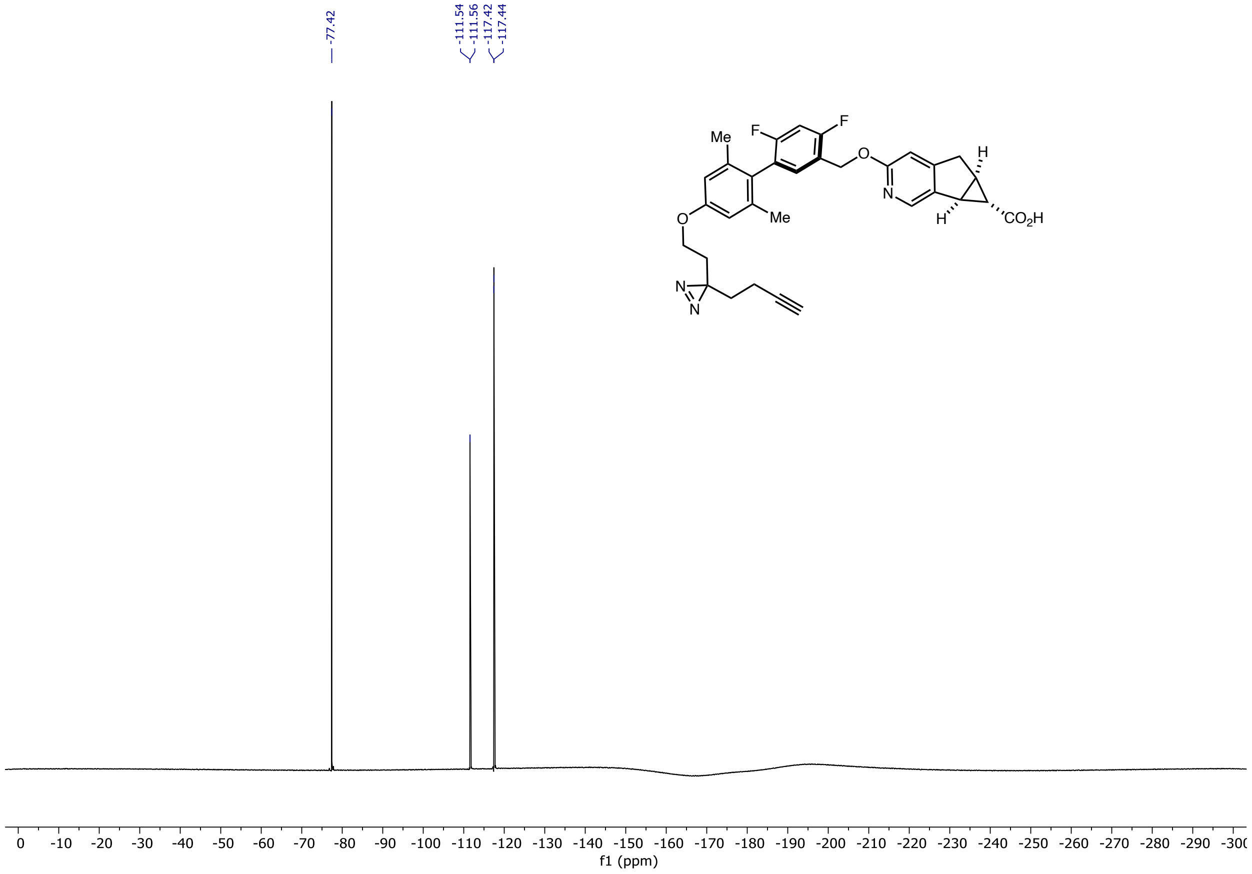

**(5aR,6S,6aS)-3-((4,6-difluoro-2',6'-dimethyl-4'-(2-(3-(2-(1-(13-oxo-17-((3aS,4S,6aR)-2-oxohexahydro-1H-thieno[3,4-d]imidazol-4-yl)-3,6,9-trioxa-12-azaheptadecyl)-1H-1,2,3-triazol-4-yl)ethyl)-3H-diazirin-3-yl)ethoxy)-[1,1'-biphenyl]-3-yl)methoxy)-5,5a,6,6a-tetrahydrocyclopropa[4,5]cyclopenta[1,2-c]pyridine-6-carboxylic acid: MK8666-Dz-Biotin**

#### MK8666-Dz-Biotin: [1]H NMR

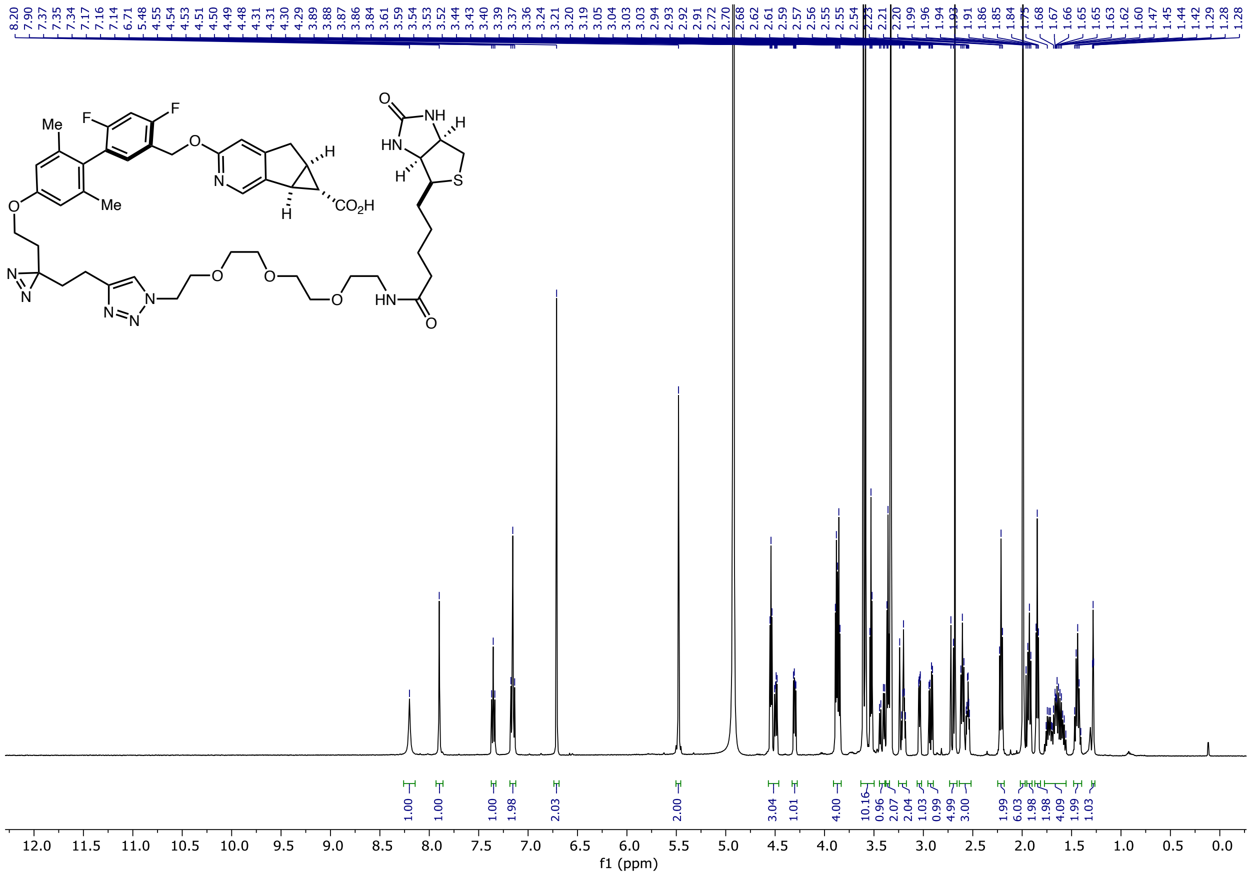

### MK8666-Dz-Biotin: $^{13}C$ NMR

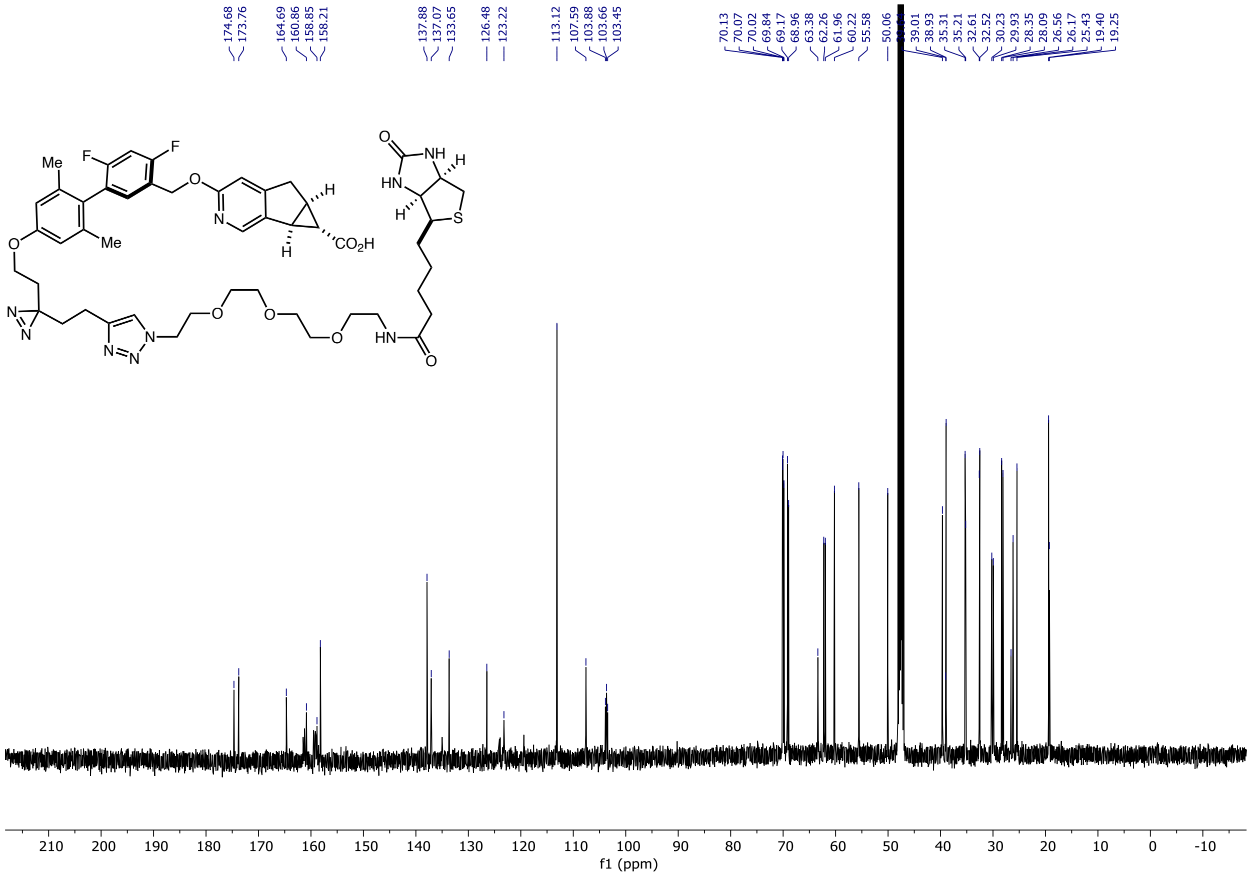

### MK8666-Dz-Biotin: $^{19}$F-NMR

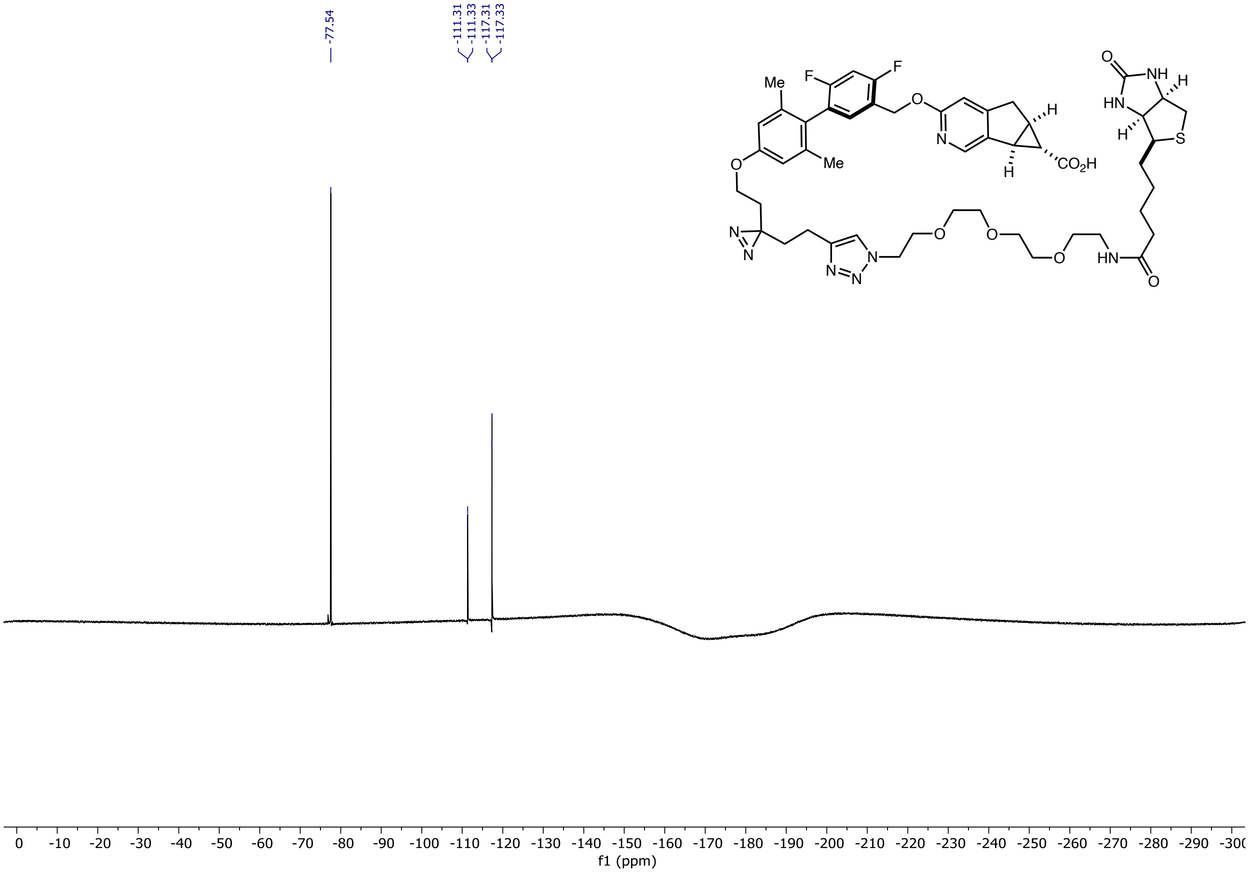

**5-((2-(2-(2-(2-(4-(2-(3-(2-((5'-((((5aR,6S,6aS)-6-carboxy-5,5a,6,6a-tetrahydrocyclopropa[4,5]cyclopenta[1,2-c]pyridin-3-yl)oxy)methyl)-2',4'-difluoro-2,6-dimethyl-[1,1'-biphenyl]-4-yl)oxy)ethyl)-3H-diazirin-3-yl)ethyl)-1H-1,2,3-triazol-1-yl)ethoxy)ethoxy)ethoxy)ethyl)carbamoyl)-2-(6-(dimethylamino)-3-(dimethyliminio)-3H-xanthen-9-yl)benzoate: MK8666-Dz-TAMRA**

#### MK8666-Dz-TAMRA: $^{1}$H NMR

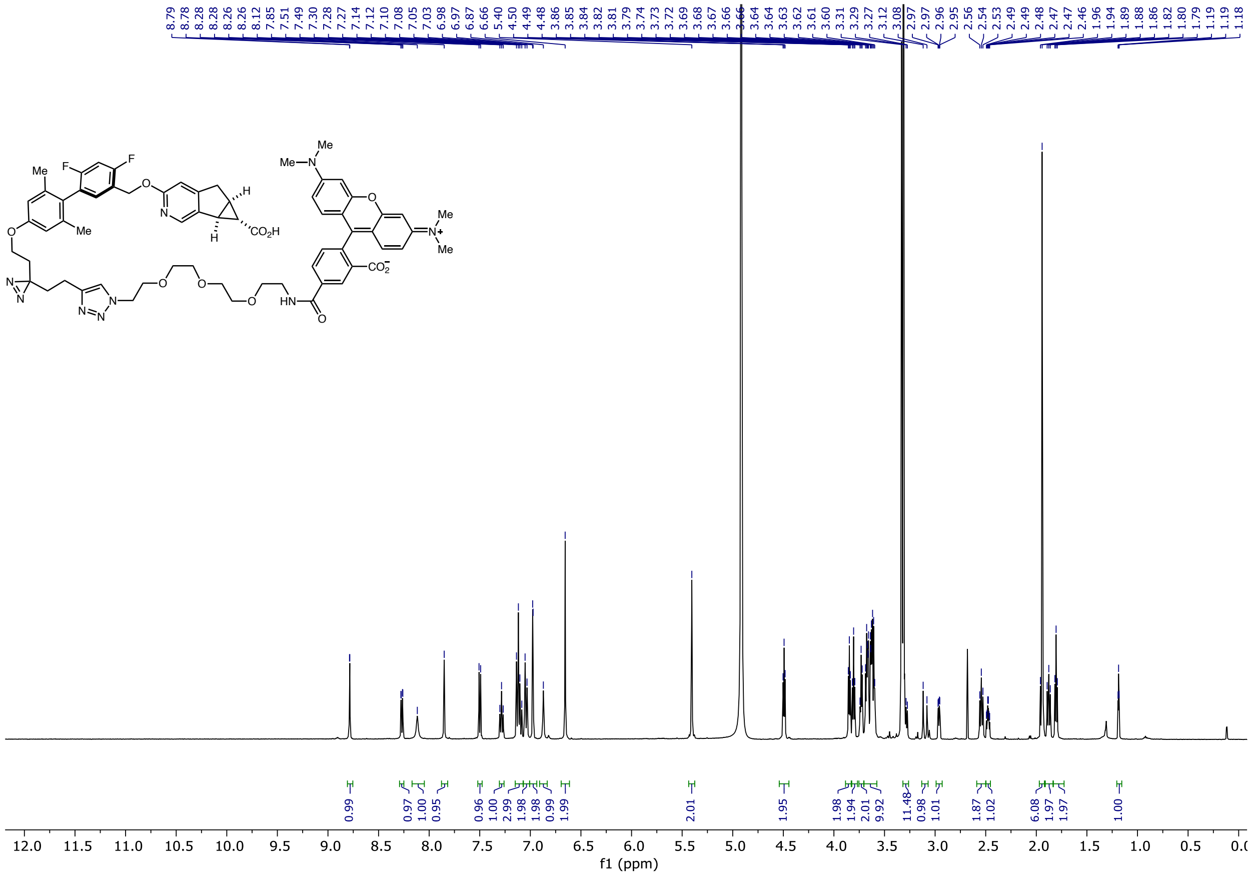

### MK8666-Dz-TAMRA: $^{13}C$ NMR

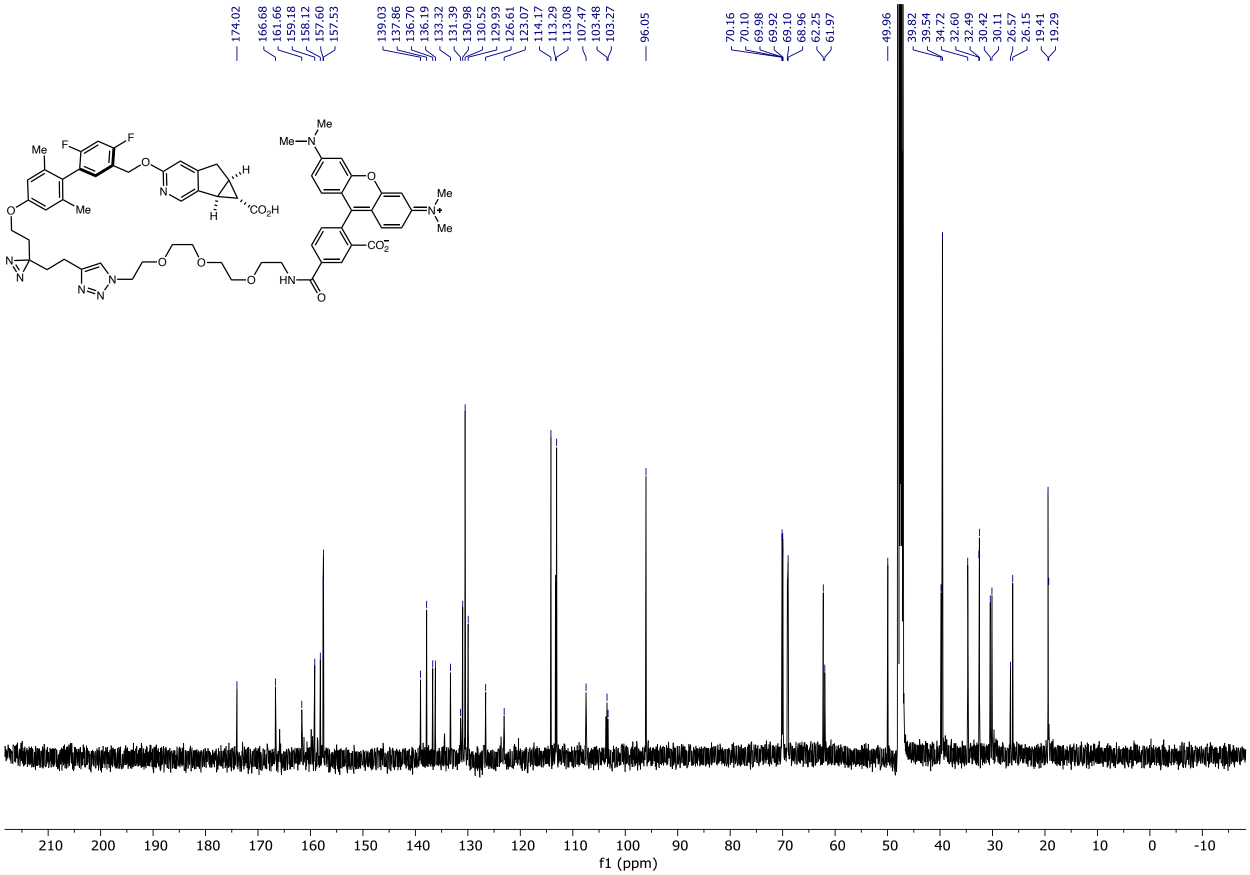

### MK8666-Dz-TAMRA: $^{19}$F-NMR

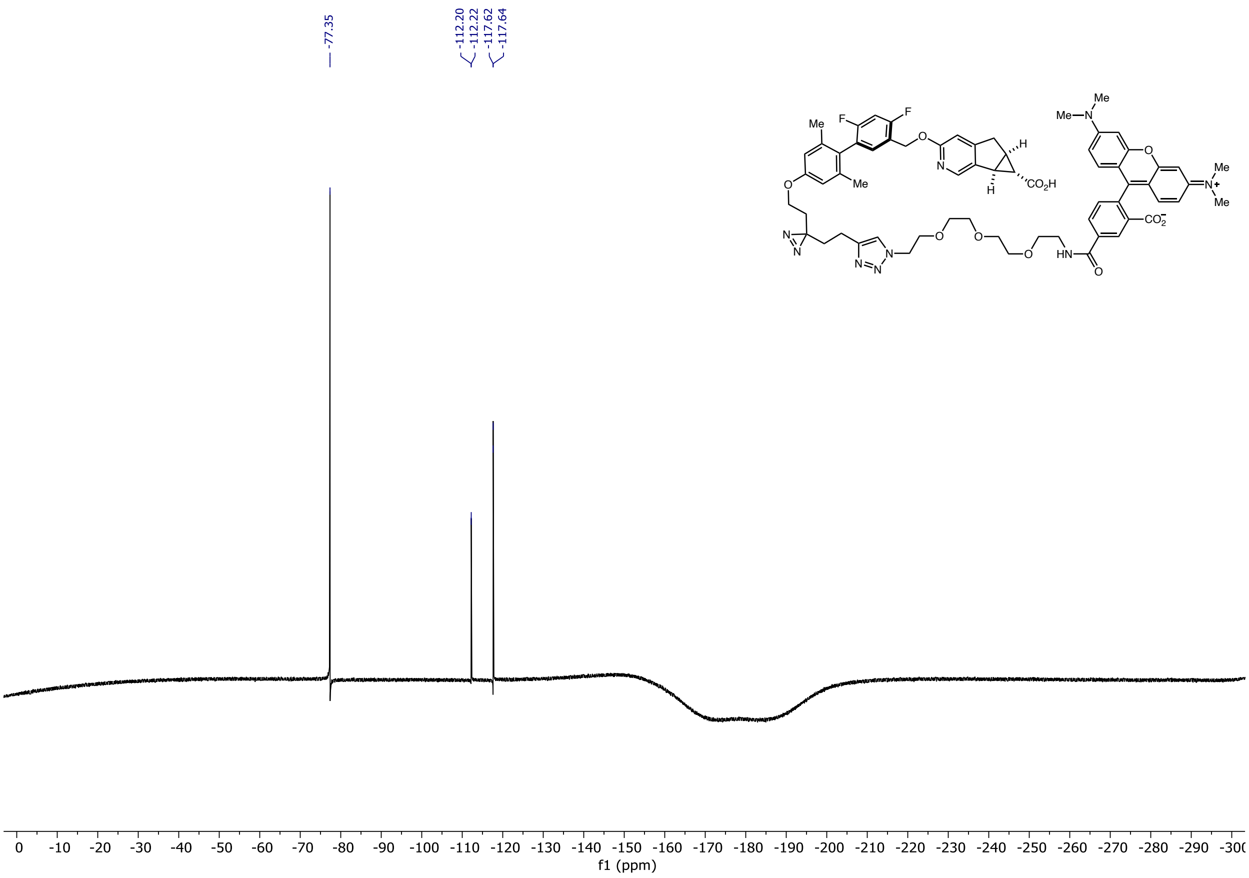

### MK8666-Ir: $^1$H NMR

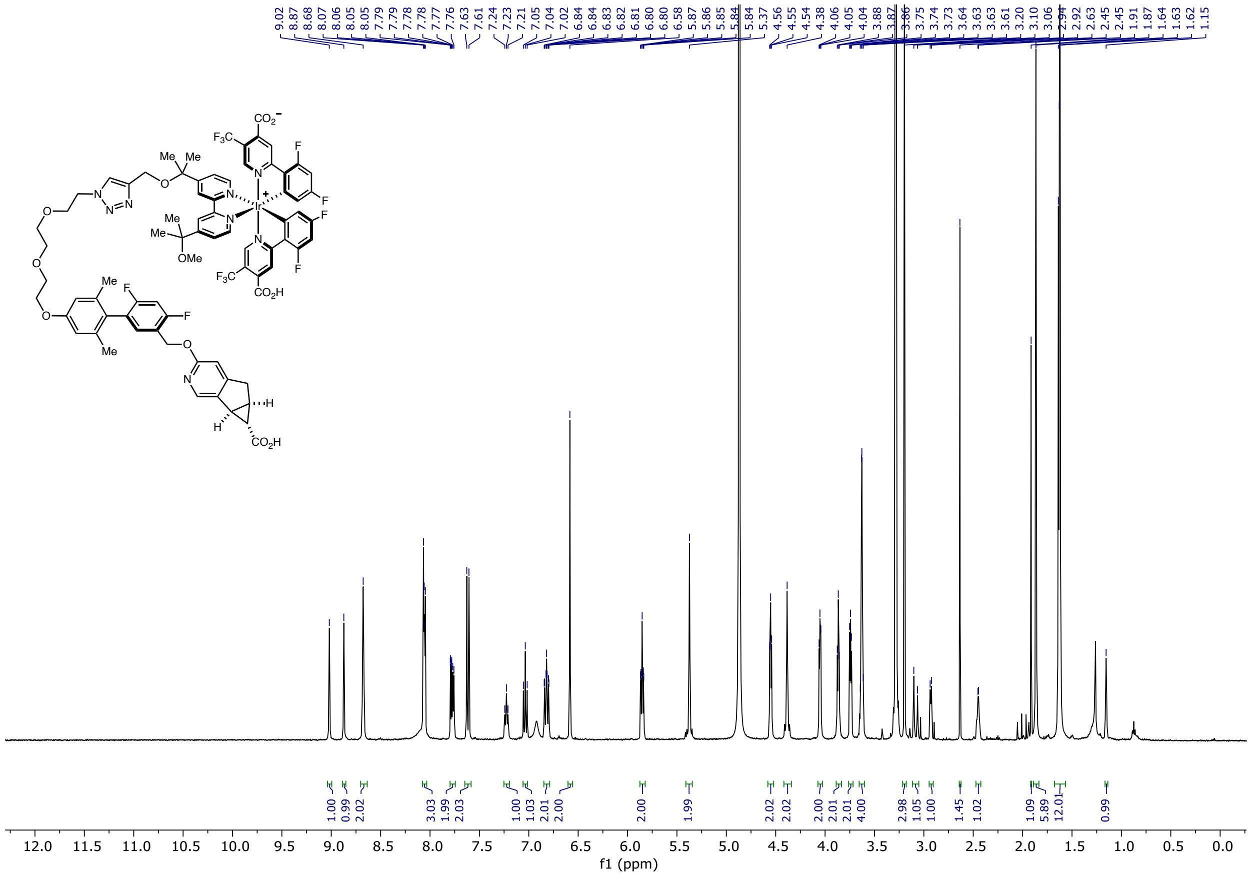

### MK8666-Ir: $^{13}$C NMR

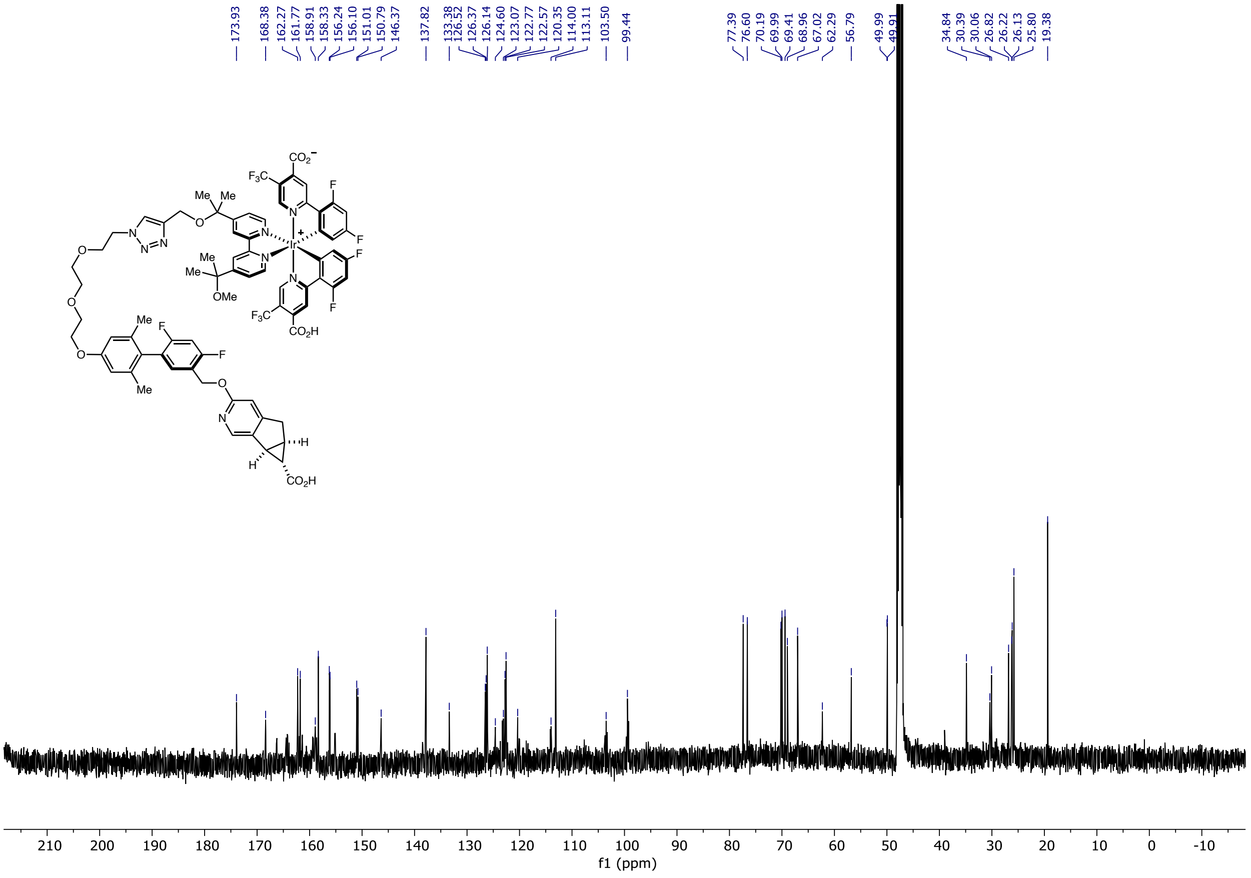

### MK8666-Ir: $^{19}$F-NMR

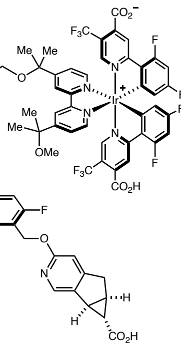

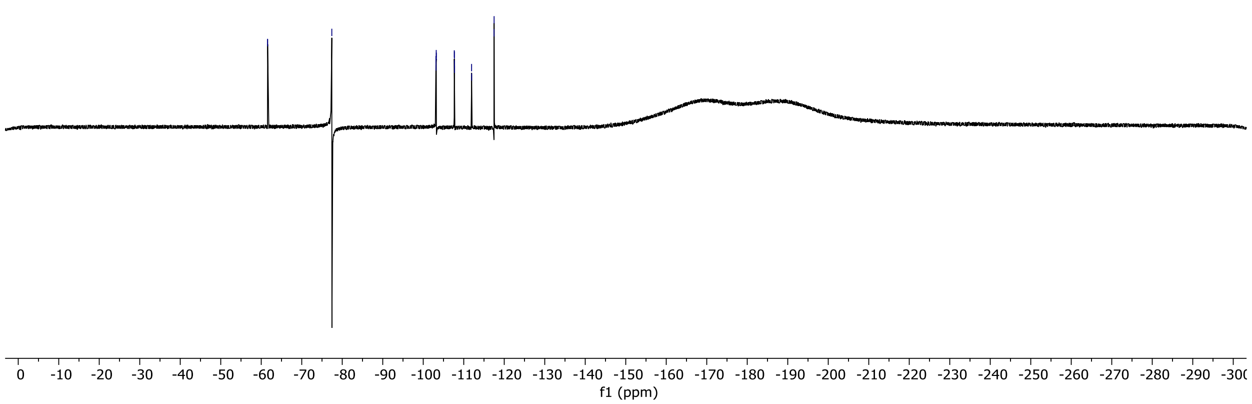

#### 3-(3-(but-3-yn-1-yl)-3H-diazirin-3-yl)-N-(4-(2-hydroxyethyl)benzyl)propenamide: $^{1}$H NMR

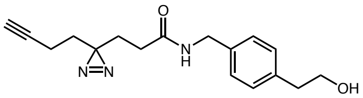

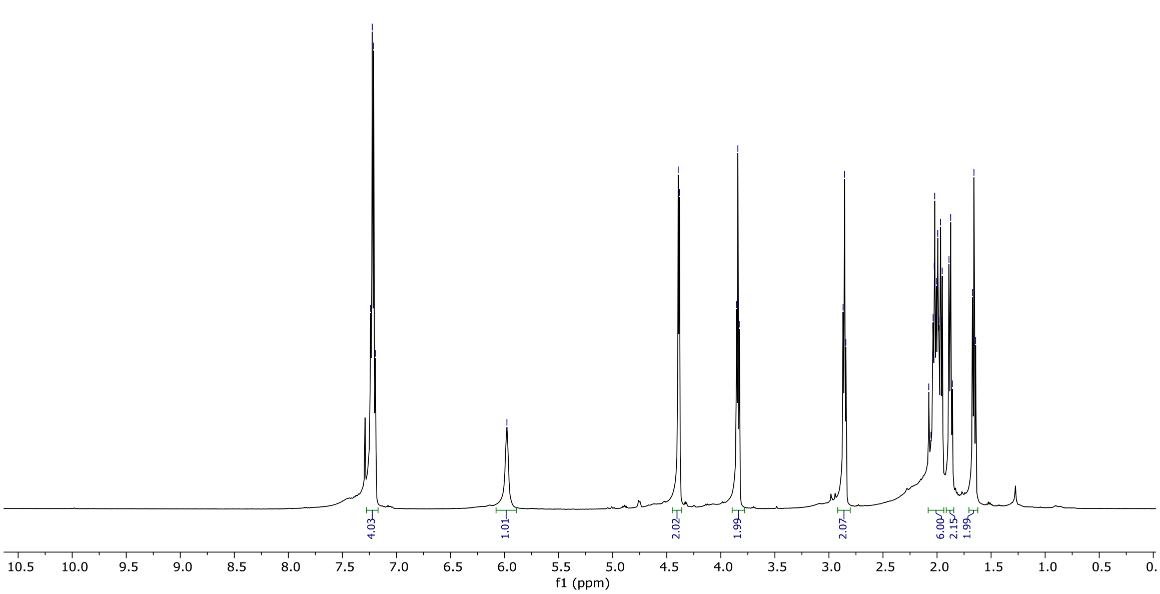

### A2aR-Yao, isomer

#### A2a-Dz, isomer [1]H NMR

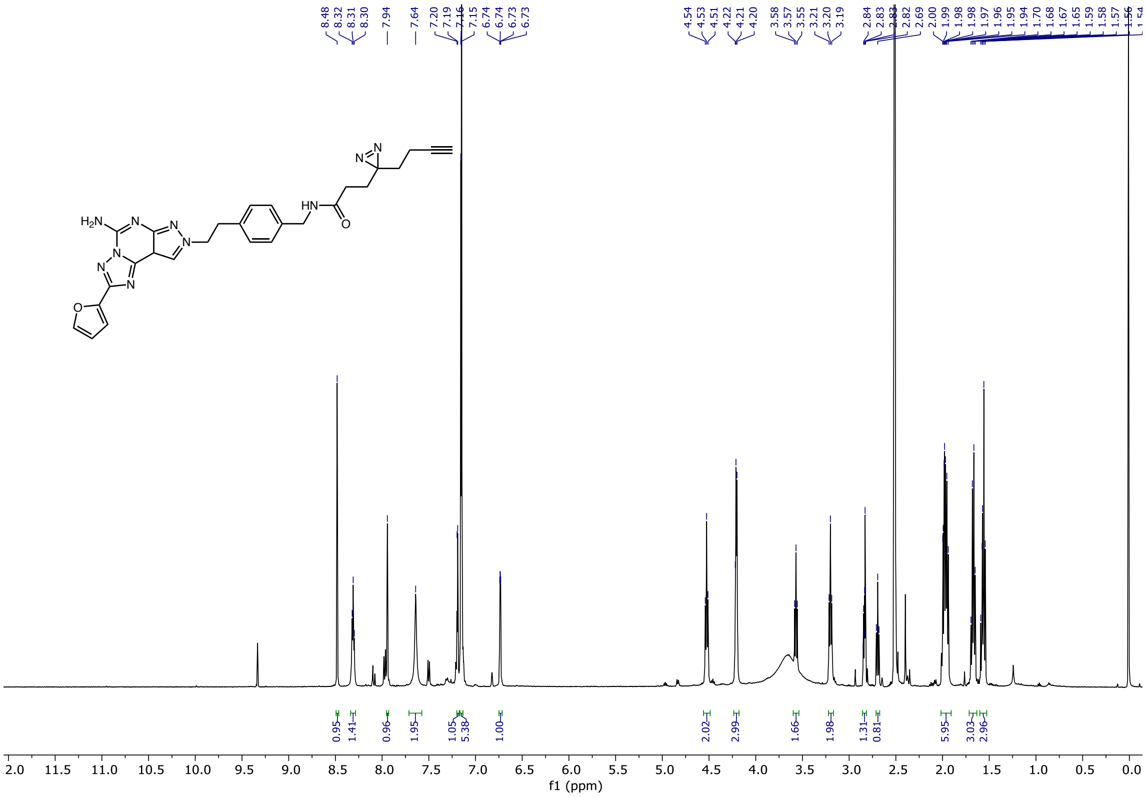

# A2aR-Dz

#### A2aR-Dz: $^{1}$H NMR

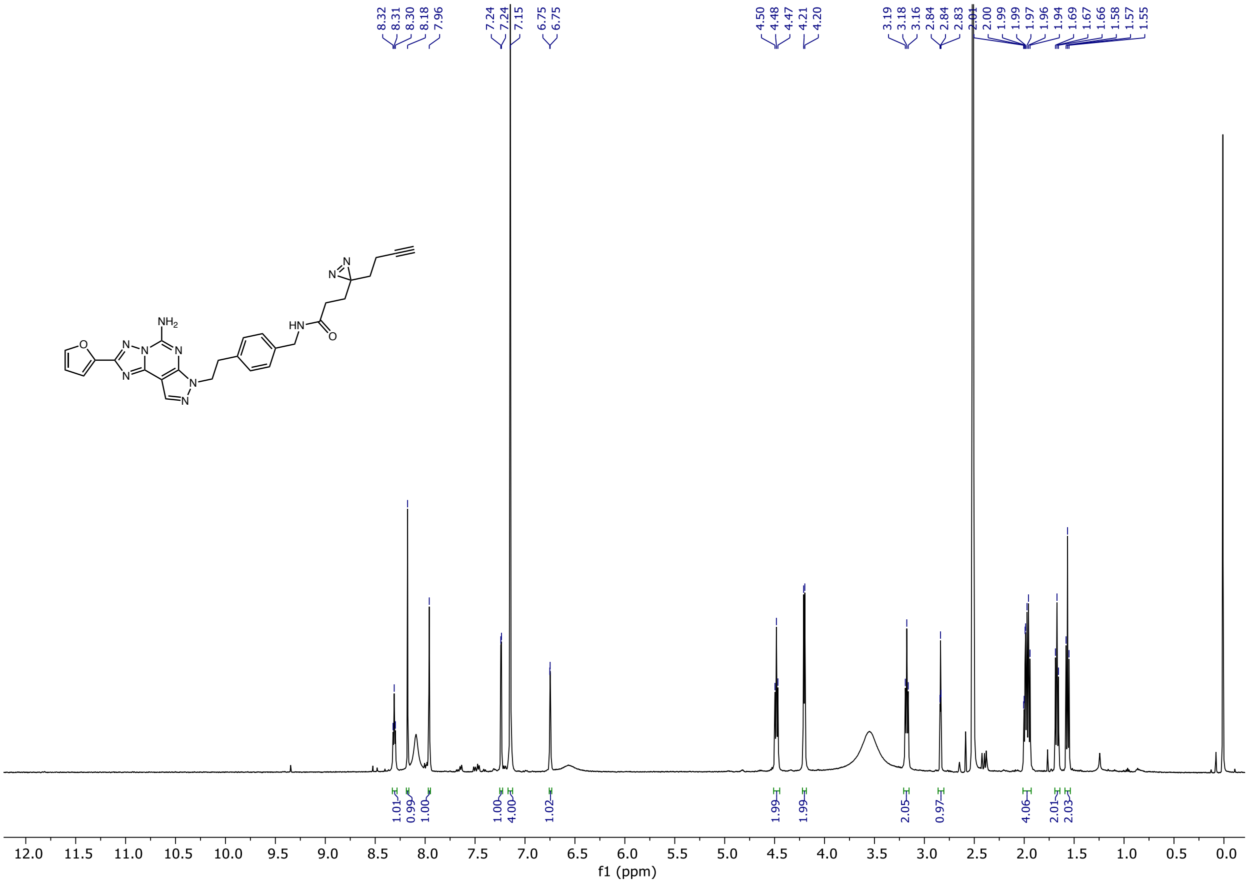

**A2aR-Dz: $^{13}$C NMR**

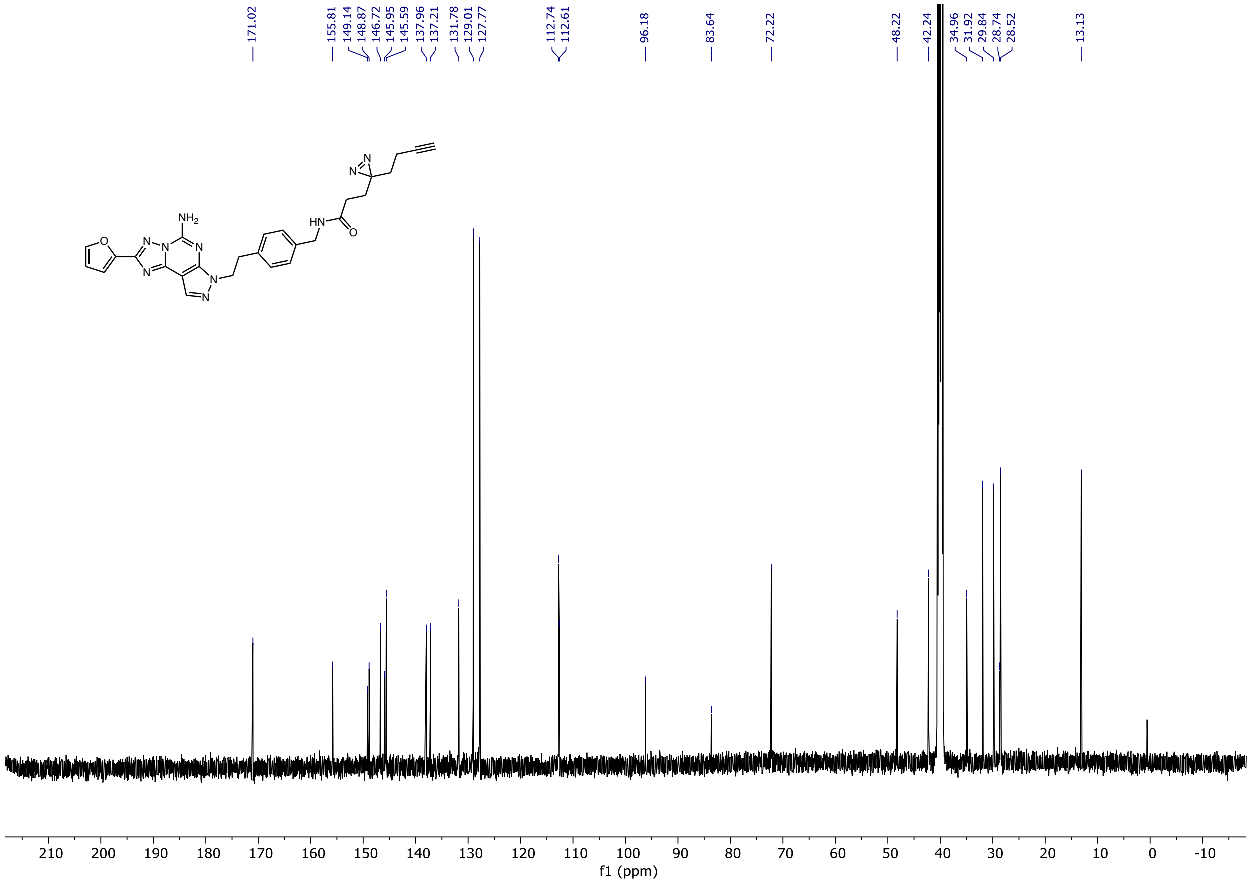

### 3-(2-(2-(2-azidoethoxy)ethoxy)ethoxy)-N-(4-(2-hydroxyethyl)benzyl)propenamide: $^1$H NMR

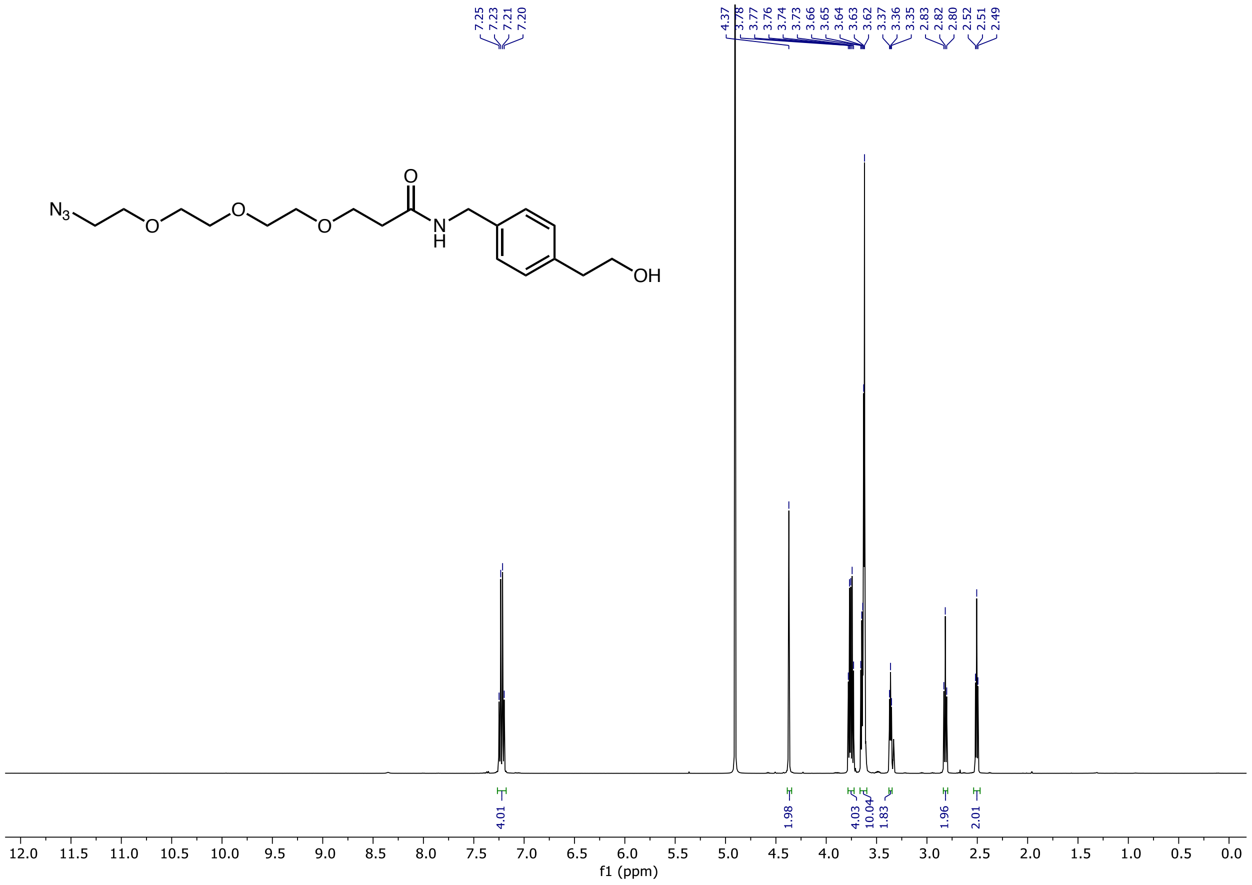

### A2aR-$N_3$, isomer $^1$H NMR

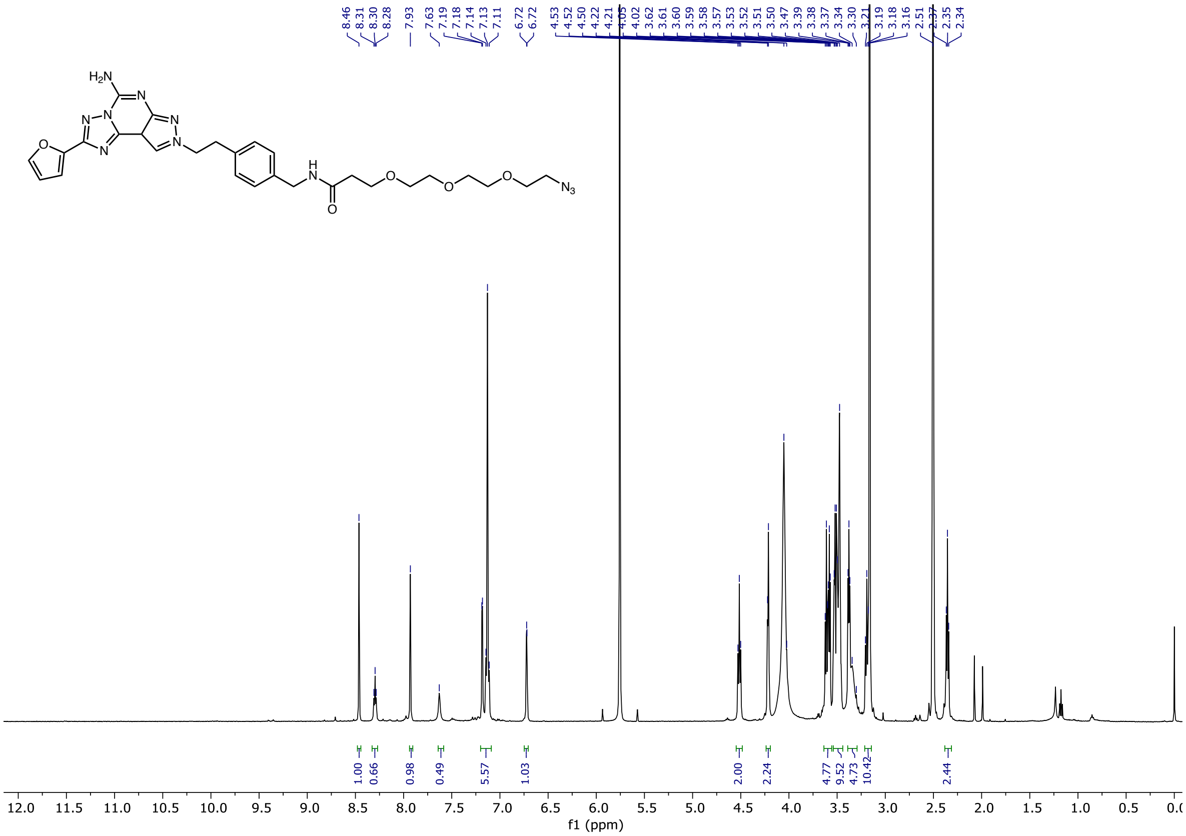

### A2aR-$N_3$: $^1$H NMR

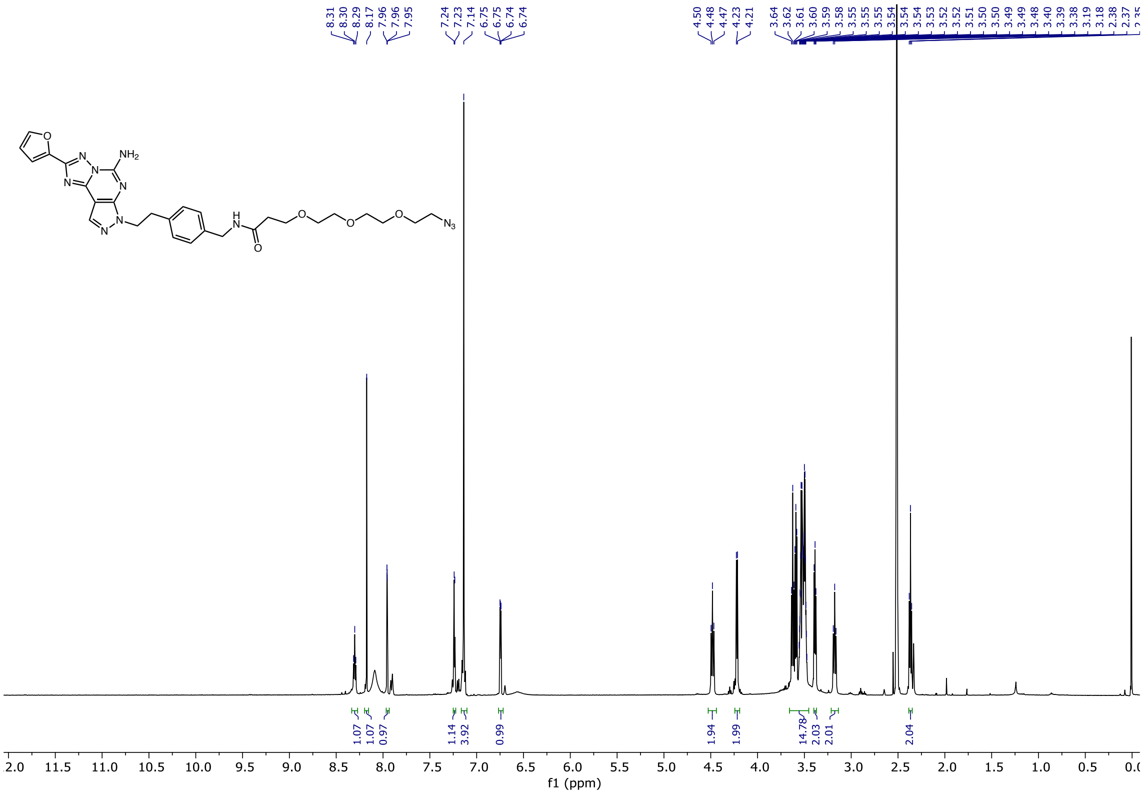

### A2aR-$N_3$: $^{13}C$ NMR

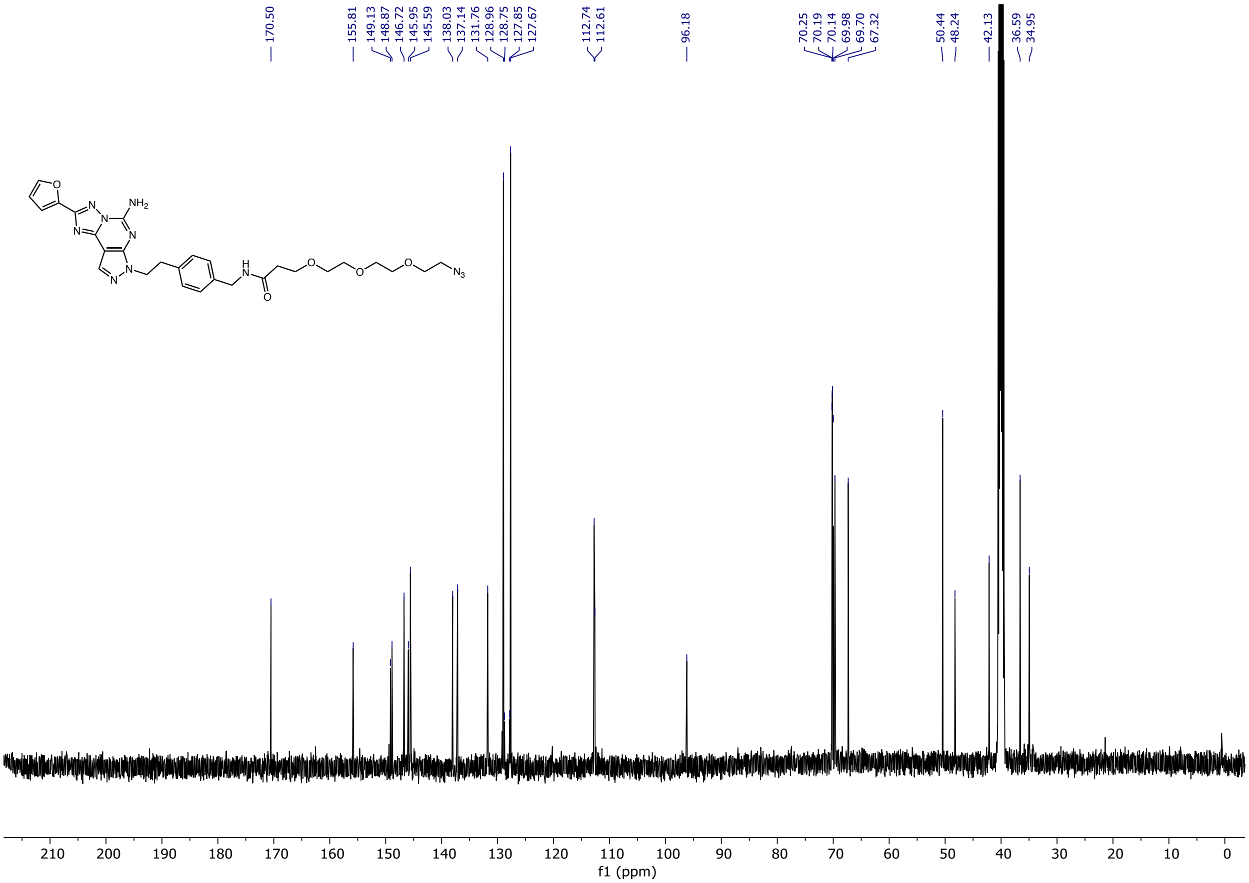

**A2aR-Yao-biotin**: $^{1}$H NMR

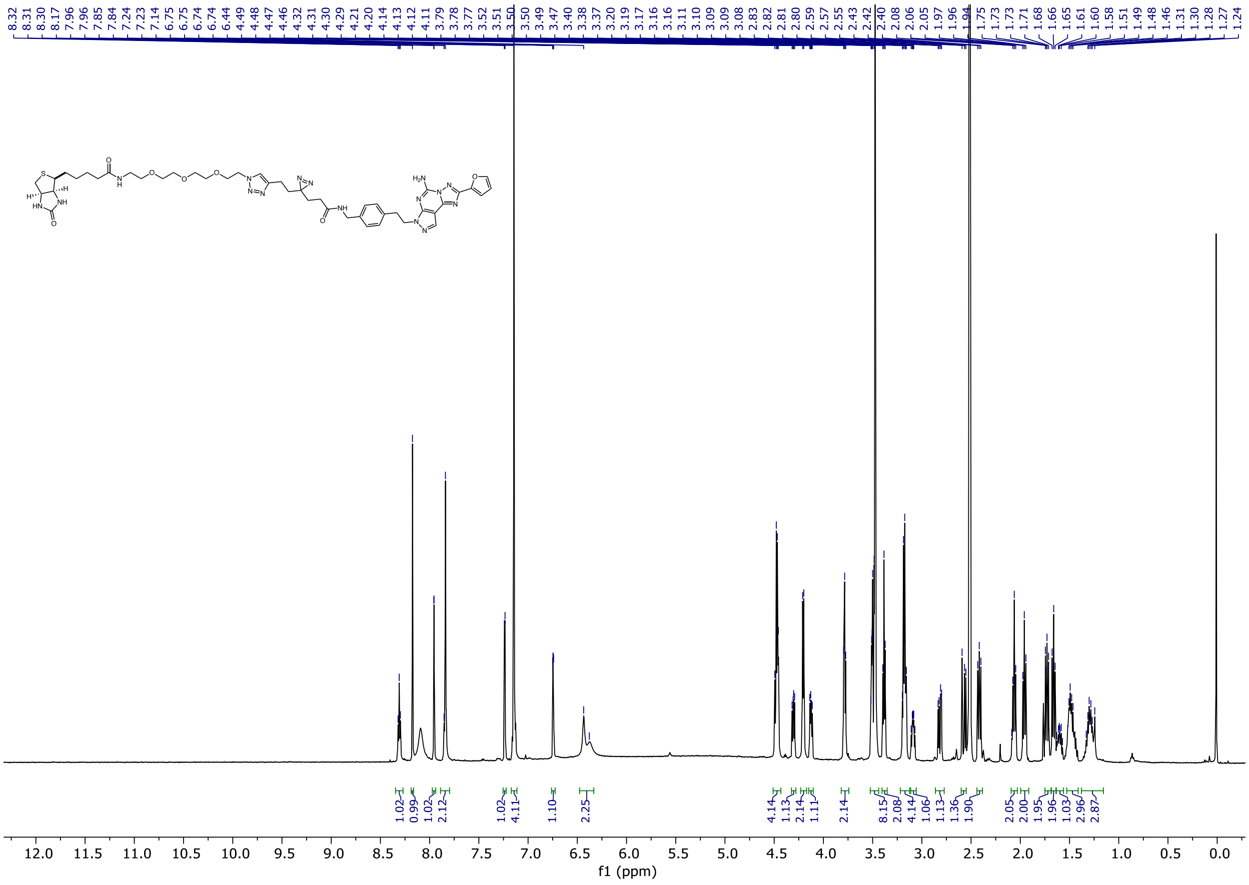

### A2aR-Yao-biotin: $^{13}C$ NMR

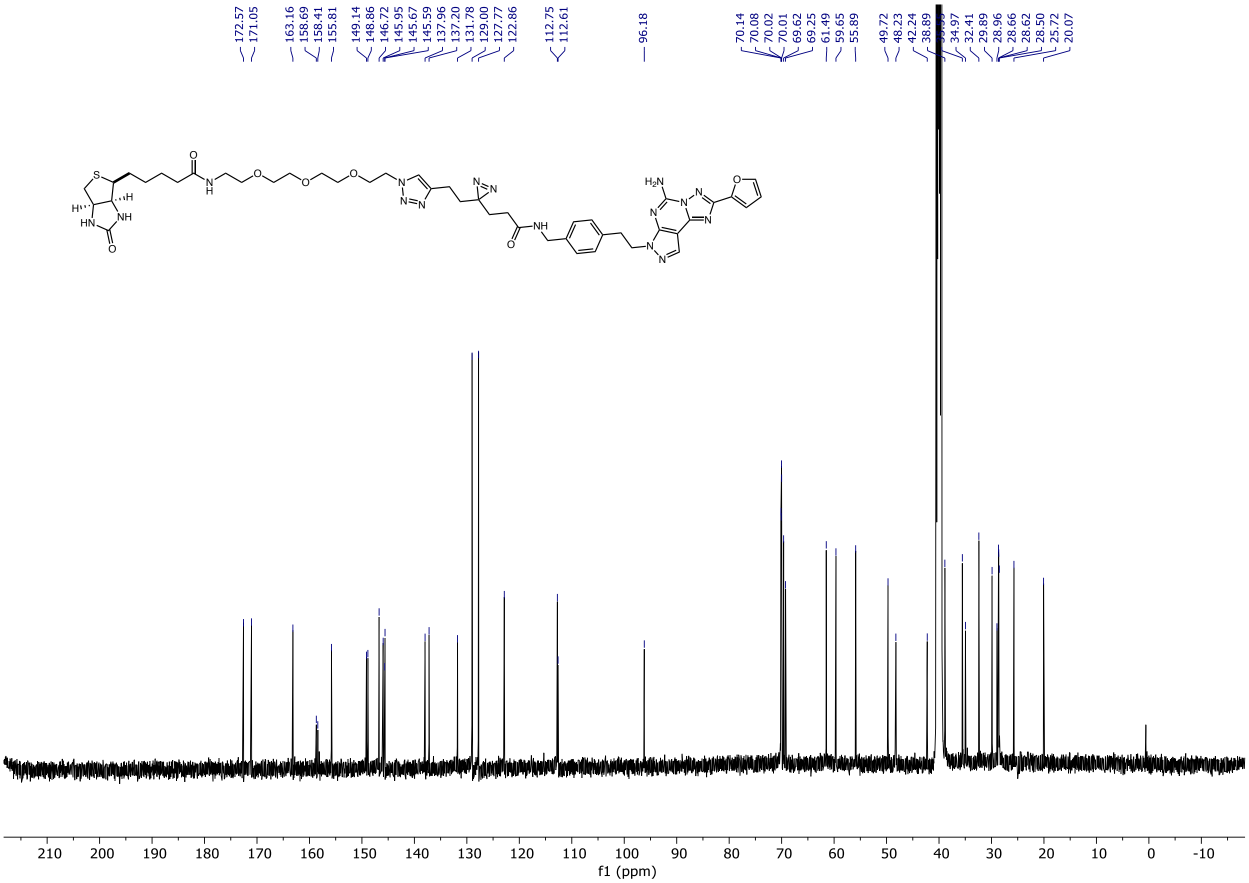

### A2aR-Yao-TAMRA: ¹H NMR

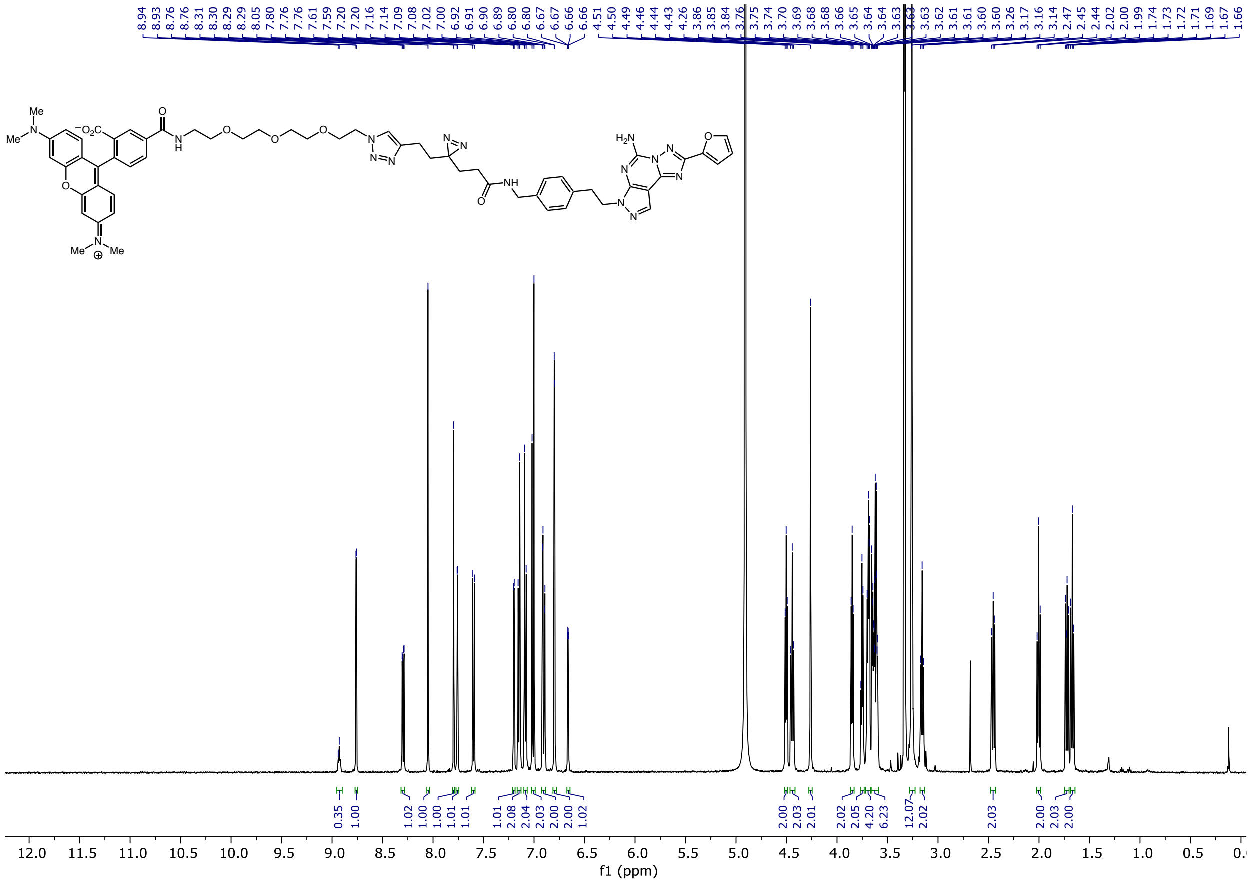

### A2aR-Yao-TAMRA: $^{13}C$ NMR

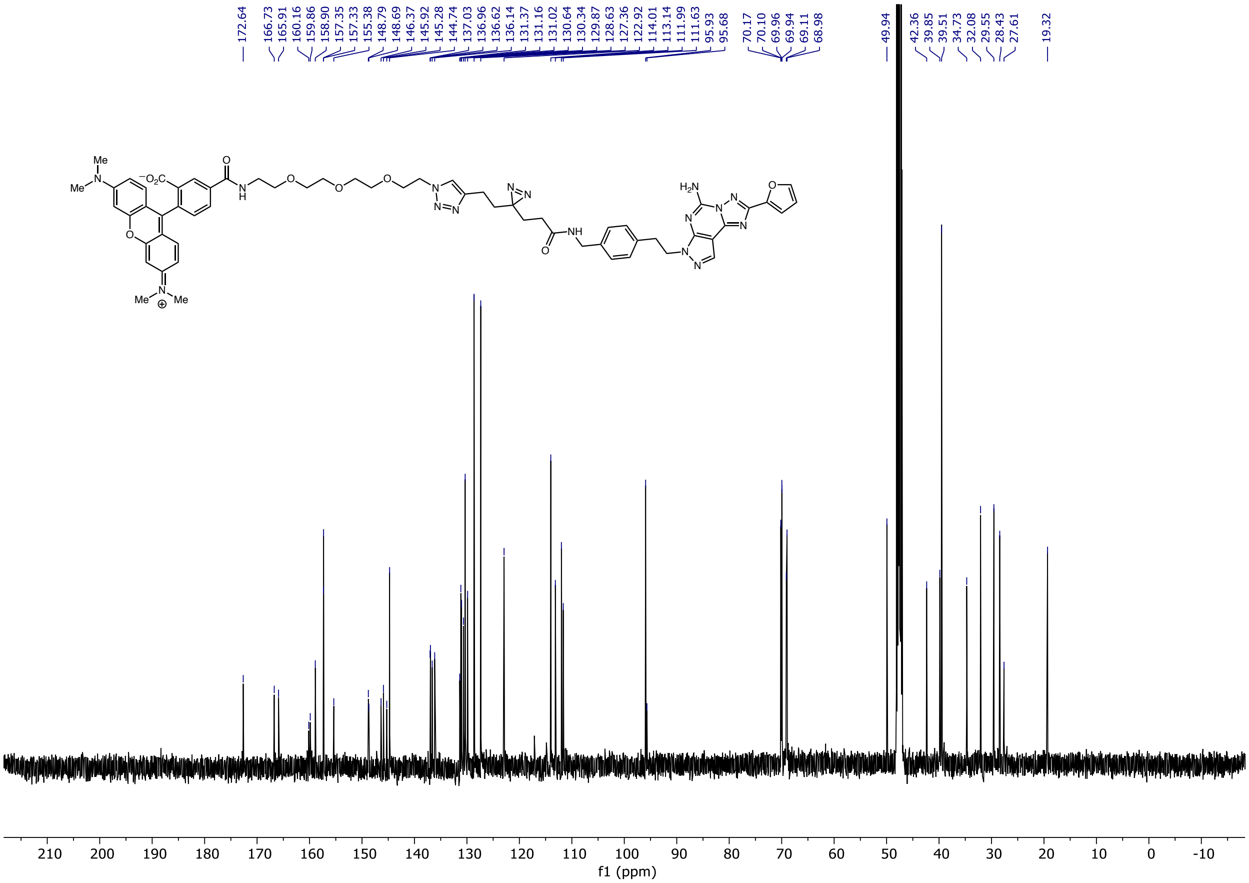

#### A2aR-Ir: $^{1}$H NMR

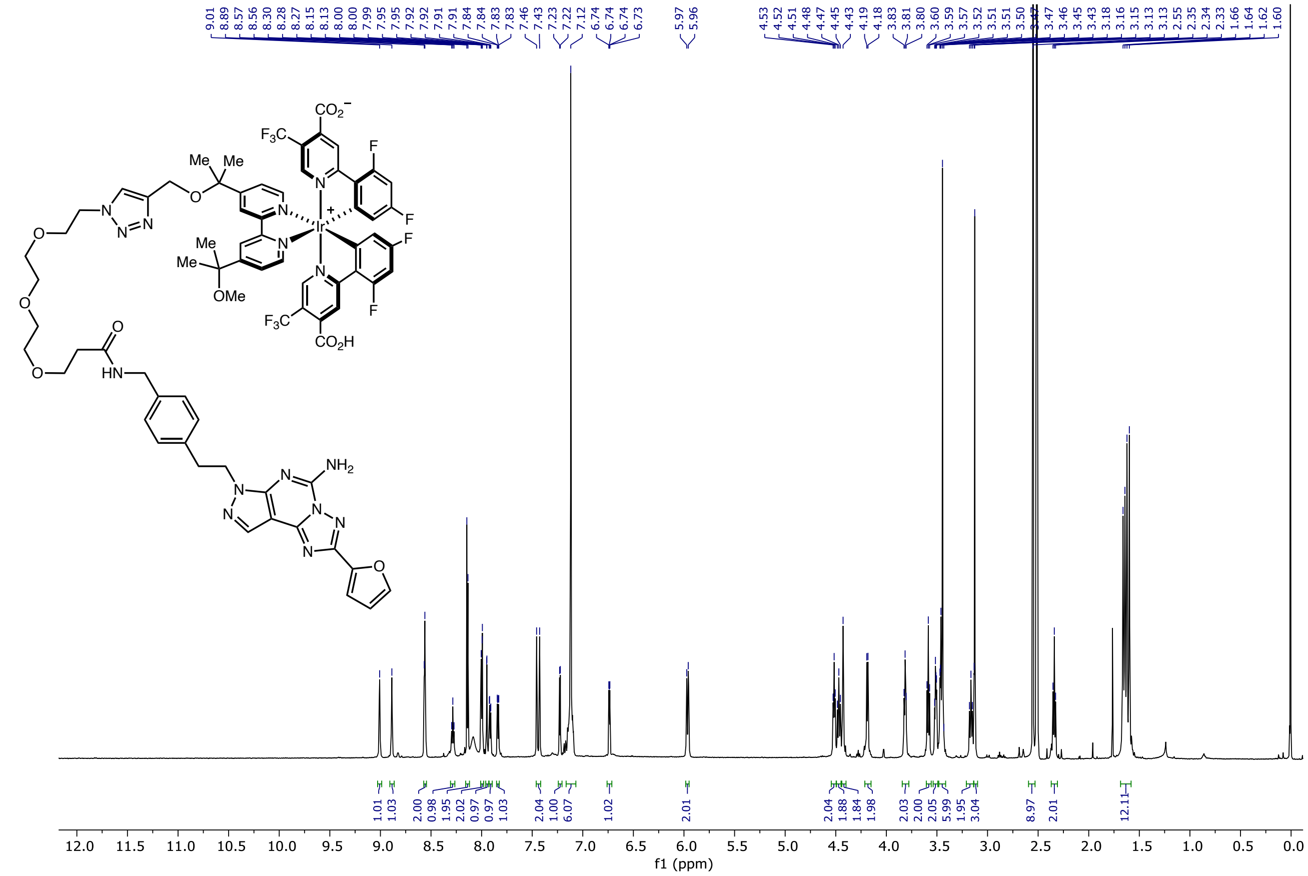

### A2aR-Ir: $^{13}C$ NMR

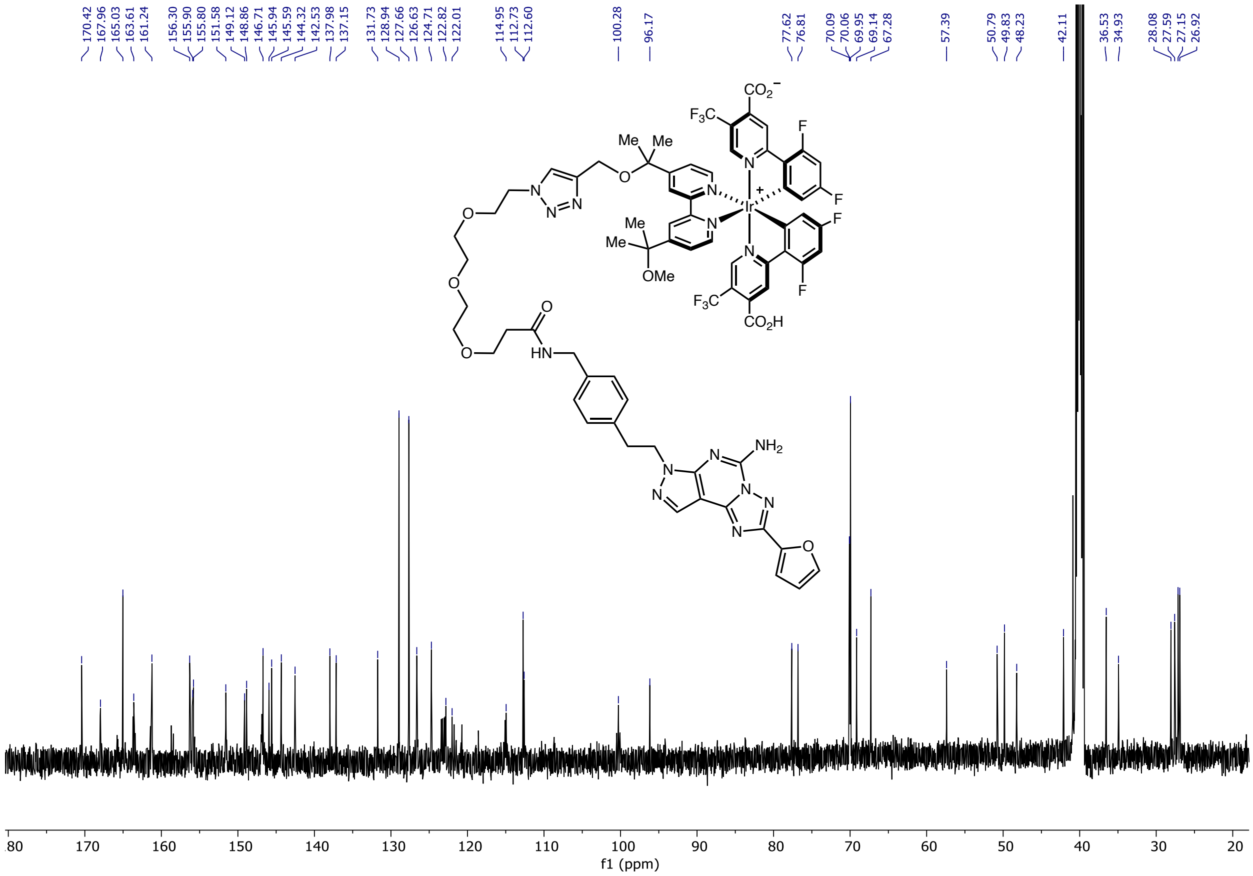

### A2aR-Ir: $^{19}$F NMR

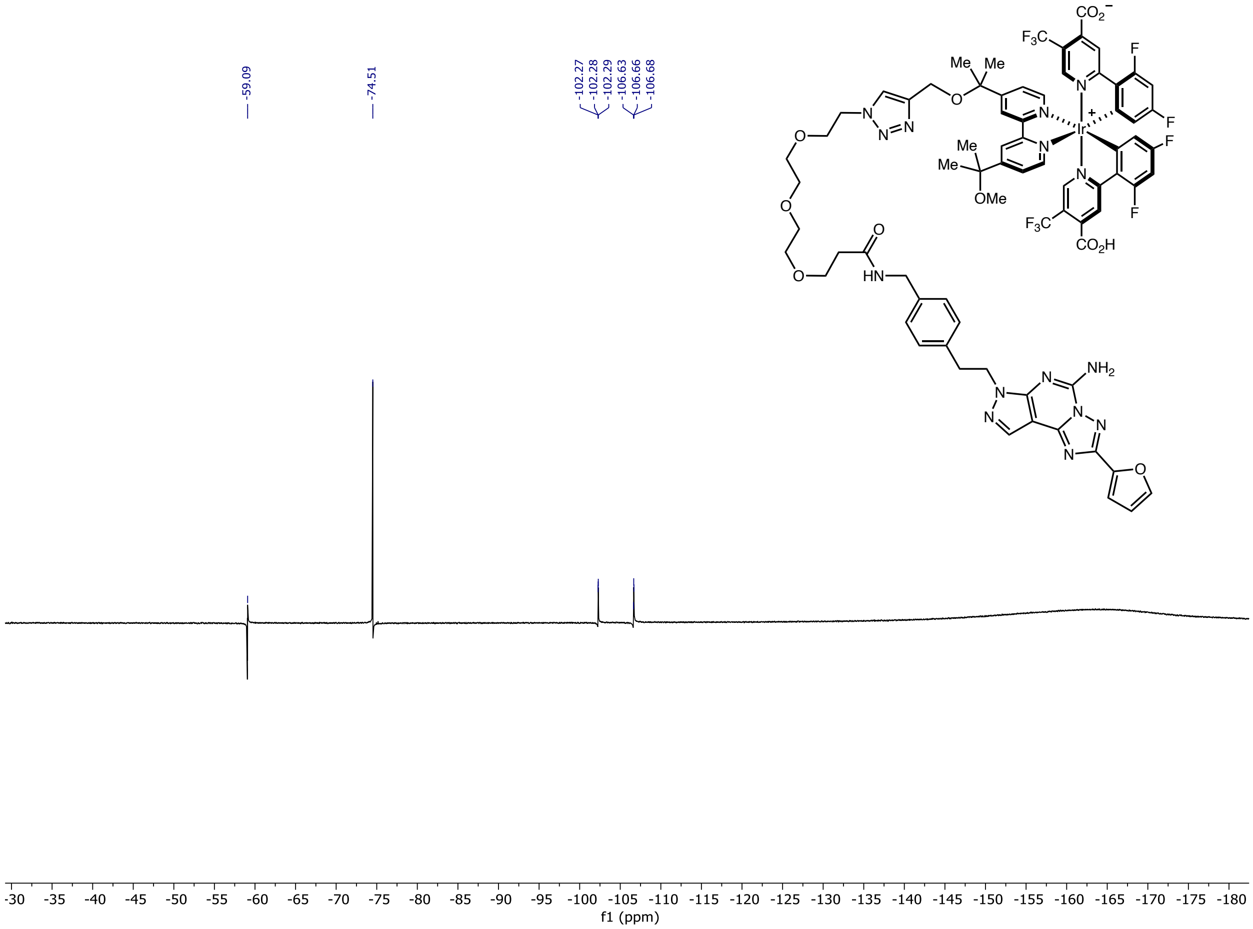
